# Drug resistance evolution in HIV in the late 1990s: hard sweeps, soft sweeps, clonal interference and the accumulation of drug resistance mutations

**DOI:** 10.1101/548198

**Authors:** Kadie-Ann Williams, Pleuni Pennings

## Abstract

The goal of this paper is to provide examples of evolutionary dynamics of HIV within patients who are treated with antiretrovirals. We hope that the figures in this paper will be used in evolution and population genetics classes. We show a wide variety of patterns, specifically: soft sweeps, hard sweeps, softening sweeps and hardening sweeps, simultaneous sweeps, accumulation of mutations and clonal interference.

## 2 Introduction

The evolution of drug resistance in HIV has been studied extensively by clinical researchers and evolutionary biologists alike (Clutter et al., 2016; Pennings, 2013; Reiss et al., 1988; Wensing et al., 2015). Research on drug resistance in HIV started in the 1980s when the first available HIV drug (AZT) turned out to be extremely vulnerable to drug resistance evolution (Mayers et al., 1992; Reiss et al., 1988) Many changes in HIV treatments in the two decades between the development of AZT (Fischl et al., 1987) and the development of single-pill combination treatments in 2006 (De Clercq, 2006) were aimed at slowing down the evolution of drug resistance. Drug resistance evolution on modern combination therapies has become quite uncommon. For example, a large study that compared the “quad” pill (based on an intergrase inhibitor) with Atripla (based on an non-nucleoside RT inhibitor) found that in both arms of the trial drug resistance developed in only 2% of patients in the first year of treatment (Sax et al., 2012). In evolutionary biology, HIV drug resistance is often used as a key example for the importance of evolutionary processes in textbooks and articles. With the increasing interest in evolutionary medicine (Nesse et al., 2009), it is likely that more and more students in science and medicine will learn about evolutionary processes through examples from HIV.

Drug resistance in HIV provides great examples of evolutionary phenomena such as selective sweeps (Messer and Neher, 2012; Pennings et al., 2014), standing genetic variation (Li et al., 2011; Paredes et al., 2010; Pennings, 2012) and mutation-selection balance (Theys et al., 2018; Zanini et al., 2017). Even though HIV has been the subject of intensive research and studied by evolutionary biologists for decades, we are still often limited by the availability of high-quality data. Specifically, datasets consisting of multiple sequences at multiple time points are uncommon for HIV. Notable exceptions are the datasets on virus populations in untreated patients described by Shankarappa et al. (1999) and the dataset by Zanini et al. (2015). Another exception, and the only one that focuses on treated patients, is the dataset from Bacheler et al. (2000), which we use in this paper. We have used this dataset for two previous studies (Pennings et al., 2014; Theys et al., 2018). In this manuscript, we show the many different patterns of drug resistance evolution observed in the Bacheler dataset. We will give an overview of the main patterns that can be observed in the evolutionary dynamics in HIV populations in patients on multi-drug treatments. These examples can be used to illustrate important evolutionary processes, such as selective sweeps and clonal interference, that have previously been illustrated by examples from computer simulations or more indirect evidence. Note that this article is meant to illustrate, not to quantify or analyze. We focus on showing the best examples of several phenomena, rather than, for example, trying to determine exactly how often a certain phenomenon occurs.

## 3 Methods

We used sequences from a dataset collected by Bacheler et al. (2000), a study that focused on patients in three clinical trials of different treatments, all based on Efavirenz (a non-nucleoside reverse transcriptase inhibitor, NNRTI) in combination with nucleoside reverse transcriptase inhibitors (NRTI) and/or protease inhibitors (PI). Some patients in these trials were initially prescribed monotherapy, which almost always lead to drug resistance, and some patients had previously been treated with some of the drugs, so their viruses were already resistant to some components of the treatment. Viral loads in these patients were typically not suppressed, which made it possible to sequence samples even during therapy. The samples were cloned and Sanger-sequenced so that we multiple independent viral sequences from each time point. We have previously used part of this dataset to study soft and hard selective sweeps (Pennings et al., 2014) and to study fitness costs of transition mutations (Theys et al., 2018).

We focused on 118 patients with at least two sequences for at least two sampling dates, and at least 5 sequences in total. This left us with a median of 24 sequences and 4.5 timepoints per patient. Sequences were 984 nucleotides long and were composed of the 297 nucleotides that encode the HIV protease protein and the 687 that encode the beginning of RT. Sequences were retrieved from Genbank under accession numbers AY000001 to AY003708.

Drug resistance mutations were used according to the HIV Drug Resistance Database (https://hivdb.stanford.edu/). For each patient, we visualized all observed resistance-related mutations and the codon they are part of. In addition, for the more extensive visualizations, we showed all sites that were polymorphic – as long as a variant was observed at least twice (i.e., we excluded singleton mutations from the visualizations). We then inspected all of the visualizations to determine whether there were convincing examples of different types of sweeps, accumulation resistance mutations or clonal interference. We considered evidence of clonal interference to be when at least one resistance mutation first goes up in frequency and then down in frequency, while at the same time another resistance mutation that is present on a different genetic background (in linkage disequilibrium) increases in frequency and reaches (near) fixation in the population. For the patients in which we found evidence for clonal interference we also created Muller plots (Muller, 1932) for the sites involved in clonal interference. All analysis was done in R (R Development Core Team (2011)) and code is available on github https://github.com/PenningsLab/ClonalInterferenceHIV.git.

## 4 Hard selective sweeps

We first focus on selective sweeps (Maynard-Smith and Haigh, 1974; Berry et al., 1991), which are very common in the Bacheler dataset. In molecular evolutionary biology, a hard selective sweep is the pattern that results when a single new mutation occurs in the population (de novo) and fixes quickly in the population due to strong selective pressure Maynard-Smith and Haigh (1974); Hermisson and Pennings (2017). In the example of patient 13, we see that a the K103N mutation in reverse trsnscriptase (RT) fixes in the population some time after day 0 (beginning of multi-drug therapy) and before day 161 (see Figure 1). This A→T mutation at the third position of codon 103 leads to an amino acid change (K103N) which is known to confer drug resistance to NNRTI drugs (Bacheler et al., 2000). Mutations in the surrounding area can either fix or become lost. Here, in patient 13, five synonymous (green) and two non-synonymous (pink) mutations fix along with the resistance mutation, whereas several synonymous and non-synonymous mutations that were seen at day 0 are no longer seen at day 161. The result is a strongly reduced amount of genetic variation at day 161 when compared to day 0. This pattern is known as a classic or hard selective sweep, as predicted by Maynard-Smith and Haigh (1974). We have previously described and analyzed selective sweeps involving K103N in the Bacheler data set (Pennings et al., 2014).

**Figure 1:**
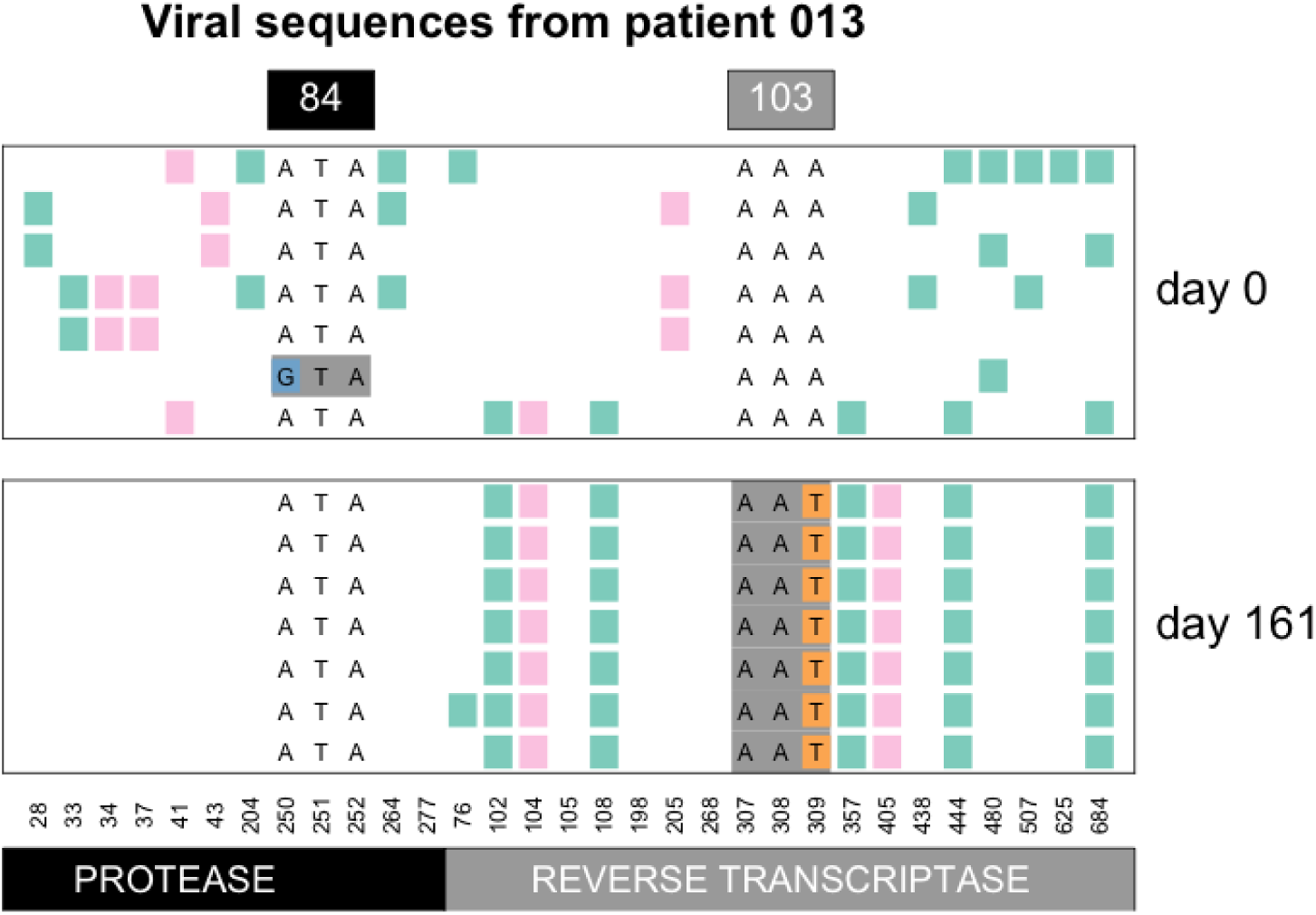
Hard sweep pattern in patient 13. The AAT codon of the K103N mutation, which confers resistance to NNRTI drugs fixes between day 0 and day 161.

Seeing the pattern of a hard sweep in HIV may be surprising to some because HIV is often seen as a pathogen with high mutation rates and large population sizes (e.g., Boltz et al. (2012); Coffin (1995)), which should favor soft sweeps (Hermisson and Pennings, 2017). There is indeed a possibility that the sweep in patient 13 was initially a soft sweep and later “hardened” – we have evidence that this can happen and we will discuss hardening of sweeps in more detail later in the manuscript. If we take the observation of a hard sweeps at face value, it suggests that the influx of mutations in some patient viral populations is not as high as one may expect. This could also be due to the treatment working well and reducing the amount of viral replication that can happen (Feder et al., 2016). Also note that this selective sweep pattern (with linked mutations up to 180 nucleotides away fixing alongside the resistance mutation) shows that the recombination rate is not high enough to effectively unlink the resistance site from its surroundings during the selective sweep. The most common resistance mutation in the Bacheler data set is K103N in reverse transcriptase.

This mutation makes the virus highly resistant to the drugs Efavirenz and Nevirapine (according to the HIV Drug Resistance Database, this mutation “reduces Nevirapine and Efavirenz susceptibility by about 50 and 20-fold, respectively.” (Shafer, 2009)). In Figure 2 we show in how many patients specific, known resistance mutations are observed in the dataset. K103N is observed in more than 80% of the patients in the dataset.

**Figure 2:**
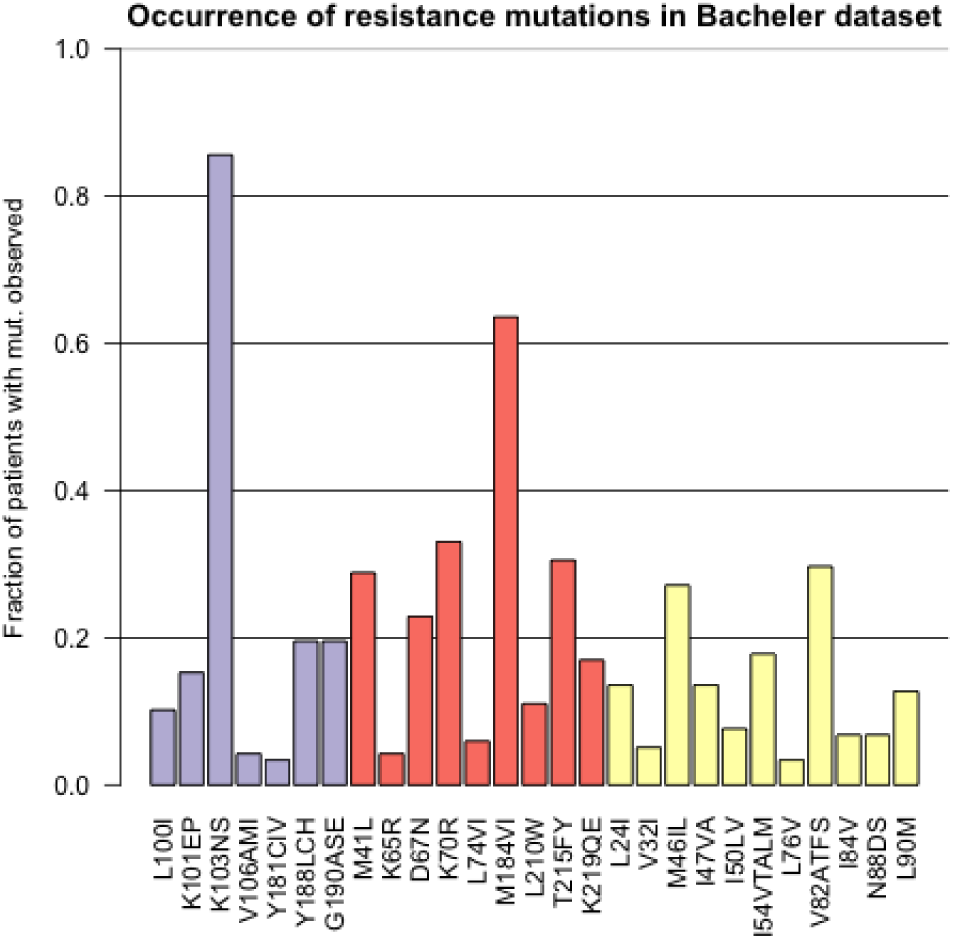
Prevalence of different resistance mutations in the Bacheler dataset with 118 patients. The purple bars are mutations that lead to NNRTI resistance, the red bars are mutations that lead to NRTI resistance and the yellow bars are mutations that lead to PI resistance.

There are two mutations that can create the K103N amino acid change: AAA→AAT and AAA→AAC. In patient 13 the AAT codon fixed (Figure 1), whereas in patient 66, the AAC codon fixed (Figure 3). Overall, we observed the AAC allele more often than the AAT allele. In 56 patients, we observed only the AAC allele, while in 4 patients we observed only the AAT allele. In 41 patients we observed both alleles (see also next section on soft sweeps from recurrent mutation) and in 17 patients, neither of the two resistance alleles are observed. It is unclear why the AAC allele is more common than the AAT allele, but this has been observed previously (Clutter et al., 2016; Palmer et al., 2006). The difference in mutation rates from A→ C and from A →T is minimal and cannot explain the difference in how often these mutations are observed in the viral sequences (Abram et al., 2010; Zanini et al., 2017). We have some evidence that the AAT allele is associated with an additional fitness cost (Tenorio, Fryer, Pennings, in preparation).

**Figure 3:**
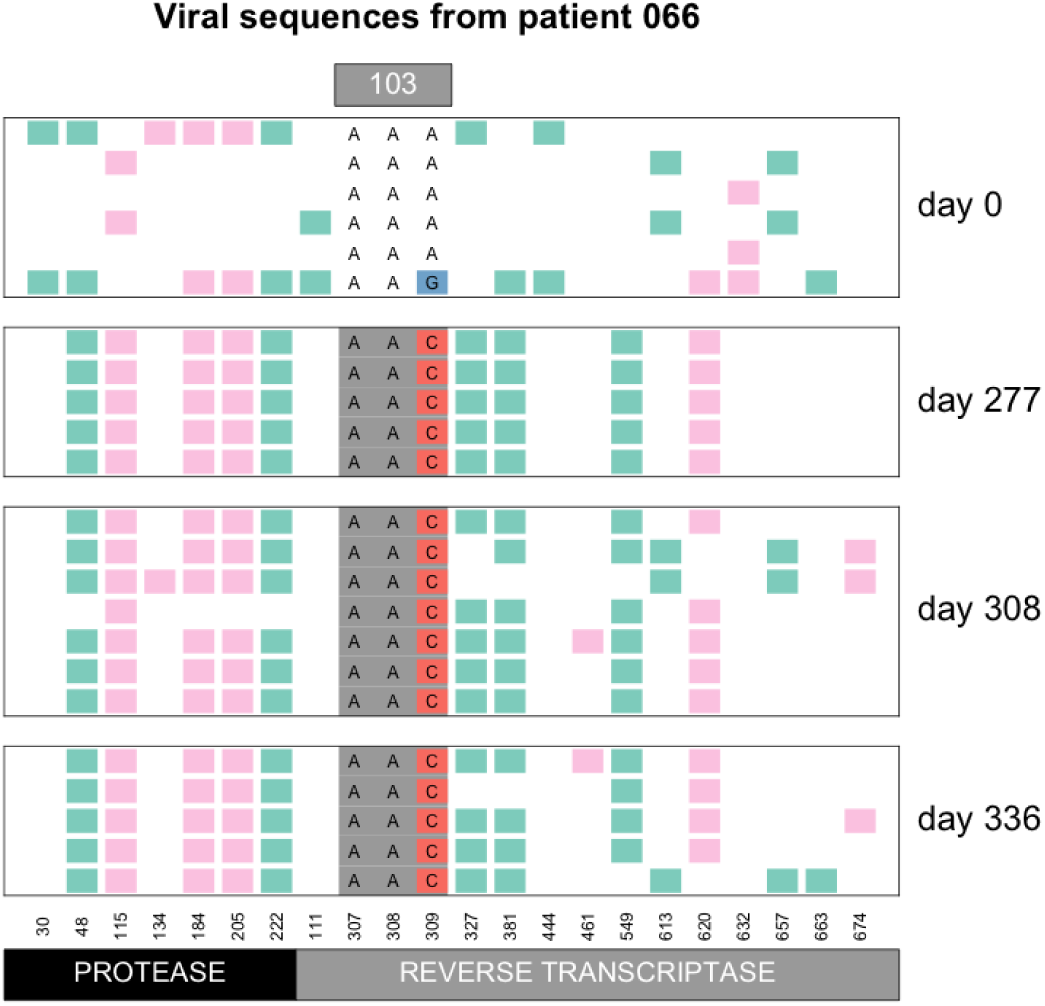
Hard sweep pattern in patient 66. The AAC codon of the K103N mutation, which confers resistance to NNRTI drugs, fixes between day 0 and day 277.

### 4.1 Sweeps at positions other than RT103

Even though K103N is the most common drug resistance mutation in the dataset, there are many other mutations that sweep and leave the classical sweep pattern of reduced variation. For example, in patient 21, mutation Y188L is not observed at day 0, but then it is seen in two of three sequences at day 55 and it is fixed at day 83 (Figure 19). The difference in genetic variation between day 0 and day 83 is striking. Note that Y188L requires two mutations (T→ C at position 1 and A →T at position 2 of the codon).

**Figure 4:**
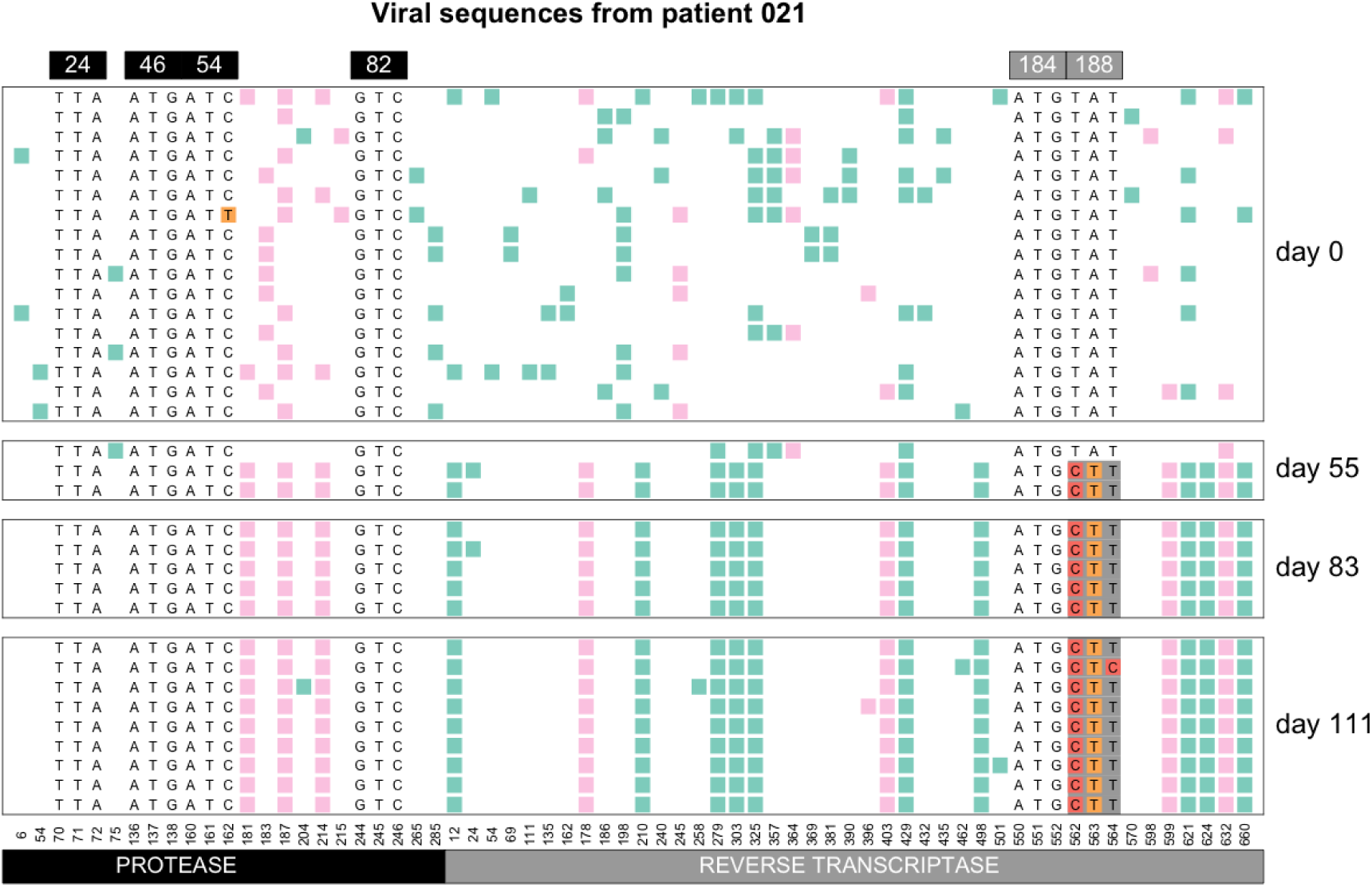
A hard selective sweep at codon 188 of reverse trsnscriptase. The wildtype codon is TAT and the mutant codon is CTT.

## 5 Soft sweeps from recurrent mutation

The commonly observed K103N mutation in reverse transcriptase can be created by an A →T mutation or an A→ C mutation. Both of these mutations are tranversion mutations and therefore have a fairly low mutation rate for HIV (mutation rate 5 *10*−*7 and 9 *10*−*7 according to Abram et al. (2010)). In earlier work, we have shown that both soft and hard sweeps occur at this position in the Bacheler dataset (Pennings et al., 2014). Hard sweeps were shown in patient 13 (Figure 1) and 66 (Figure 3). However, in one third of the patients (41 out of 118 patients, 35%) both alleles were observed, see Figure 5. When both codons (AAT and AAC) are present at position 103, this provides a clear example of a soft sweep from recurrent mutation (Pennings et al., 2014). We see this pattern in patient 81 (Figure 6).

**Figure 5:**
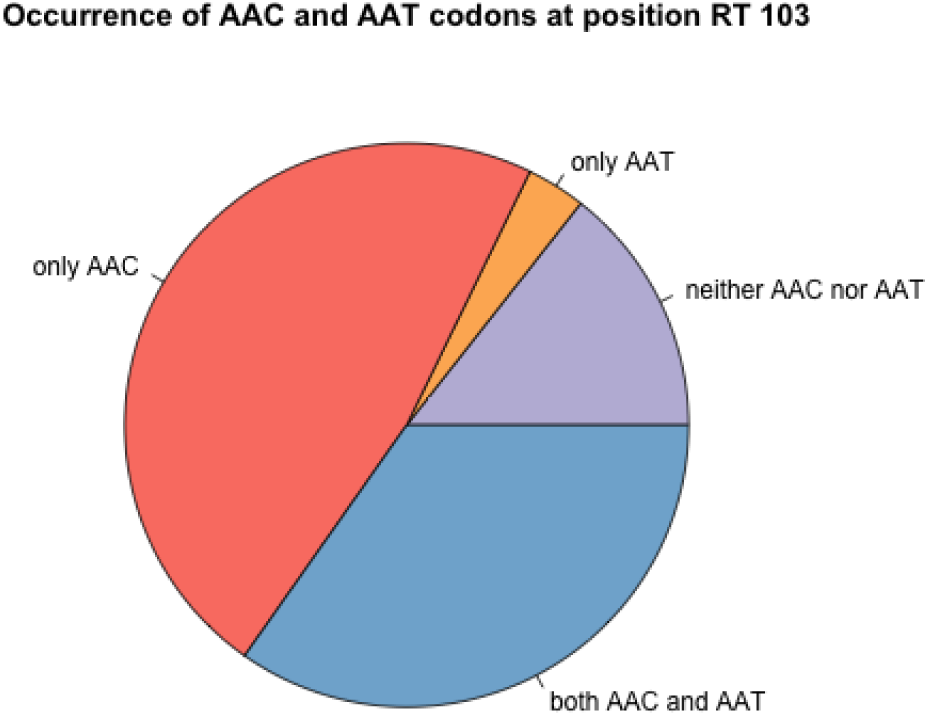
Occurrence of AAC and AAT codons at K103N in 118 patients. In 14% of the patients, we see neither AAC nor AAT, but only the WT codon (AAA) is observed. In 47% of the patients, we observe the resistance codon AAC, in 3% of the patients we see the resistance codon AAC and in 35% of the patients, we see both resistance codons. In all cases we may also see the WT codon, especially in the early time points.

**Figure 6:**
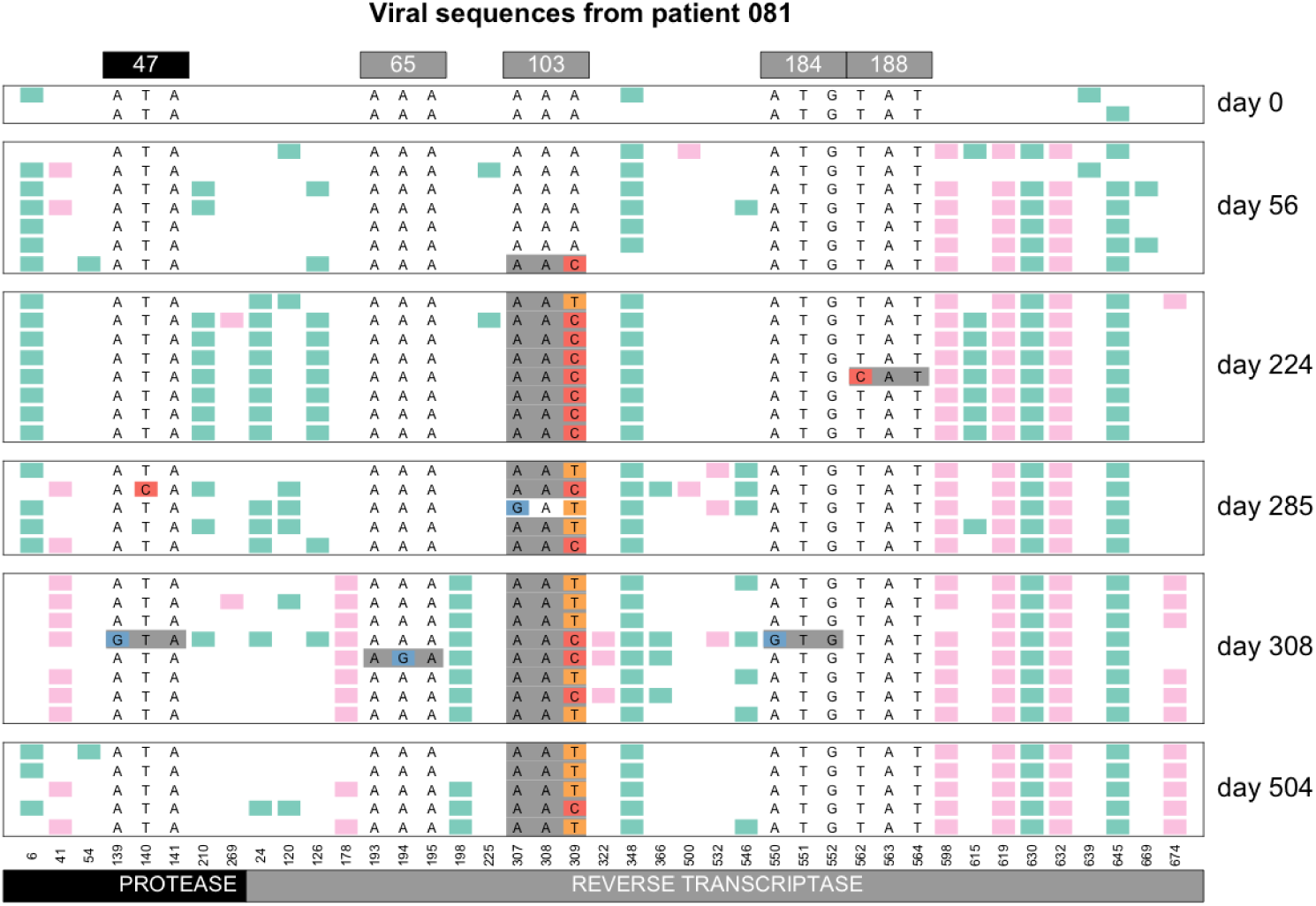
Soft sweep from recurrent mutation in patient 81. At codon 103, both the AAC and the AAT resistance codon are observed.

### Soft sweeps at M41L

Recurrent mutation soft sweeps also occur at other positions than RT103. Here, we look at recurrent mutation soft sweeps at RT41. At this position, the M41L mutation happens through ATG→TTG or ATG →CTG. M41L confers resistance to NRTI drugs such as AZT and Tenofovir. Both alleles are seen in patient 133 (Figure 7). At the second sampling timepoint (day 280) only one of the alleles is observed, possibly due to a sweep at position K103N between day 0 and 280.

**Figure 7:**
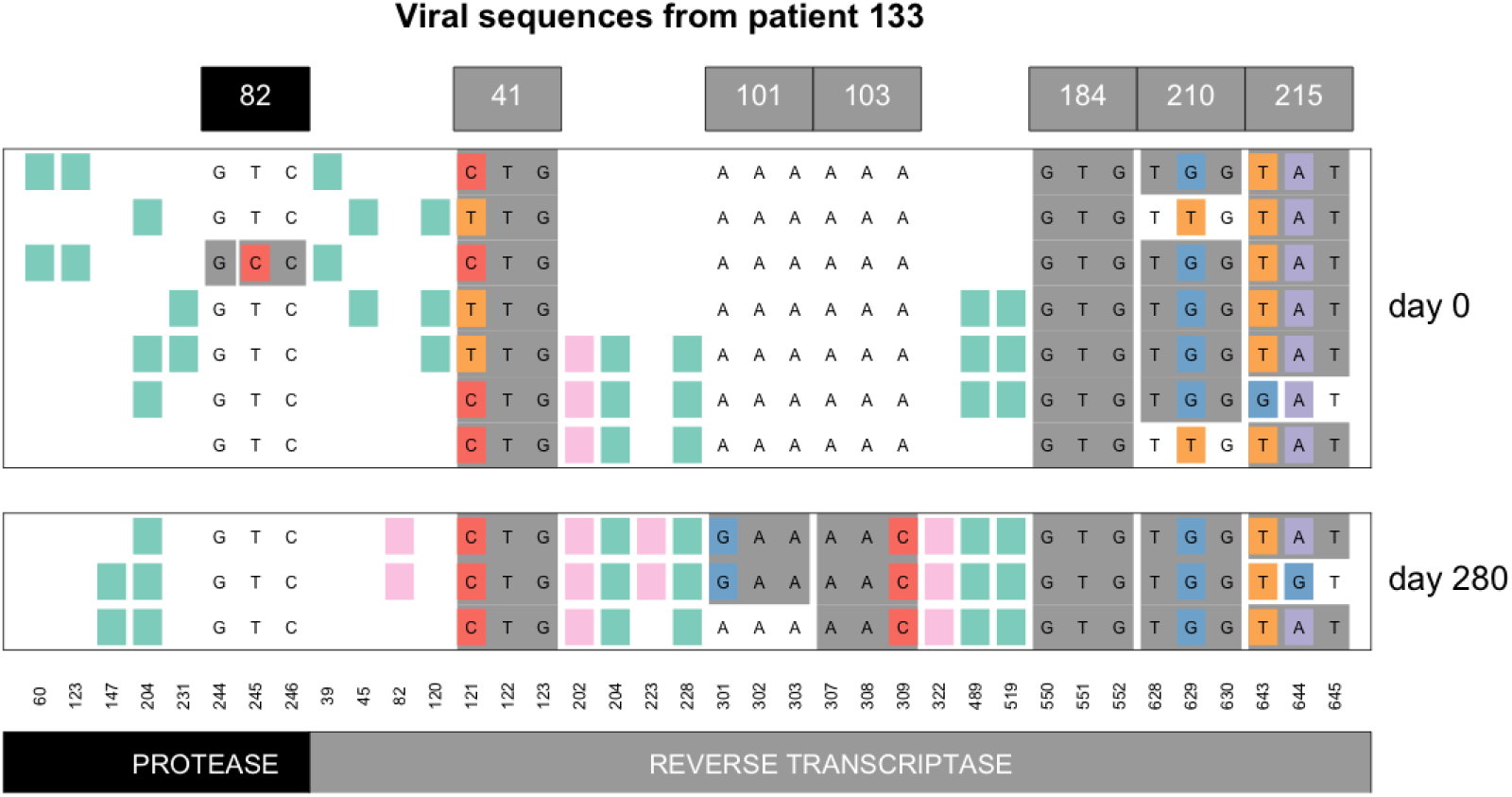
Soft sweep from recurrent mutation at M41L in patient 133.

The M41L mutation is not as common in the Bacheler dataset as K103N, and when it is observed, it is frequently already present on day 0 (as seen in Figure 7). Just like for the K103N mutation, we wanted to determine which of the two alleles is more common in the dataset at the M41L mutation. For M41L we see more balanced numbers of the two codons (CTC and TTC, Figure 8), which suggests that these codons are equal or nearly equal in fitness.

**Figure 8:**
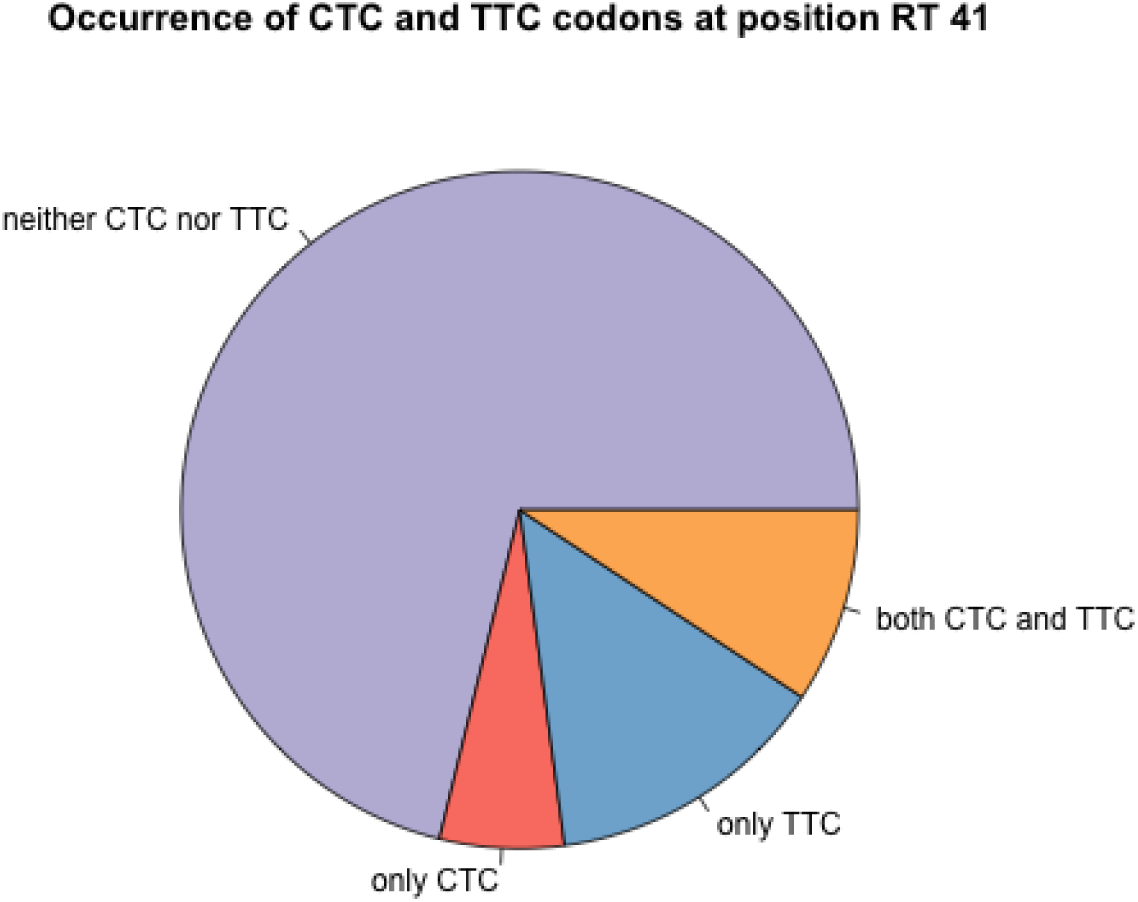
Occurrence of the CTC and TTC alleles at position RT41. In 71% of the patients, we see neither CTC nor TTC, but only the WT codon (ATC) is observed. In 6% of the patients, we observe the resistance codon CTC, in 14% of the patients we see the resistance codon TTC and in 9% of the patients, we see both resistance codons. In all cases we may also see the WT codon, especially in the early time points.

A recurrent mutation soft sweep is also seen at position RT188 in patient 26 (supplementary figure for patient 26).

### “Hardening” of soft sweeps

In 2014, Wilson and colleagues described a scenario where sweeps can start off being soft (due to recurrent mutation) but then become hard due to population bottlenecks. This phenomenon was named the “hardening” of soft sweeps (Wilson et al., 2014). In several patients, we observe patterns that are consistent with a hardening of soft sweeps. Here we show two examples of the hardening of soft sweeps from recurrent mutation in patient 24 (Figure 9) and in patient 11 (Figure 10). In these patients, both resistance codons at position 103 are present initially, but one of the two alleles then vanishes, as the other fixes. This hardening of a sweep is fairly common in the dataset (e.g., we also see it in patients 5, 55, 57, 87 and 170 at position RT103 and in patients 94 and 133 for position RT41, see supplementary figures). The hardening may be due to demographic bottlenecks, as described in the original paper by Wilson et al., but it could also be due to hitchhiking when a new beneficial mutation occurs. For example, in patient 11, the G →A reversal mutation in RT184 (recreating the WT at that position) may be a beneficial mutation that spreads, such that the AAC allele at RT103 hitchhikes to fixation due to the sweep at 184 (Figure 10)

**Figure 9:**
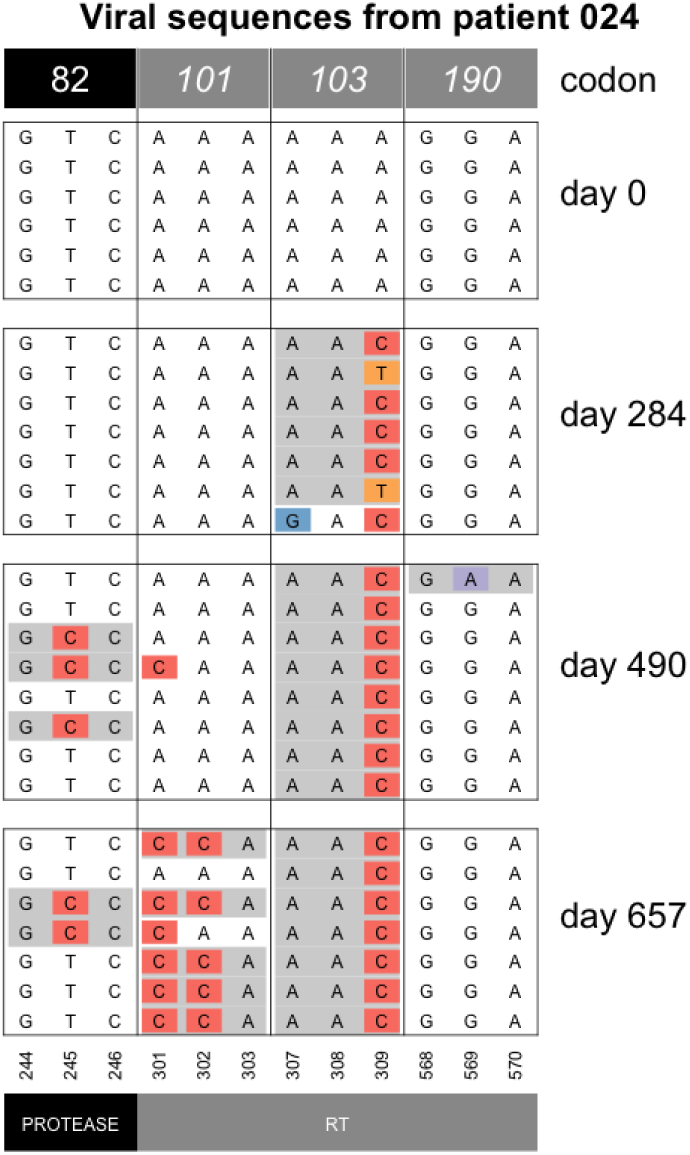
Hardening of a soft sweep at K103N in patient 24.

**Figure 10:**
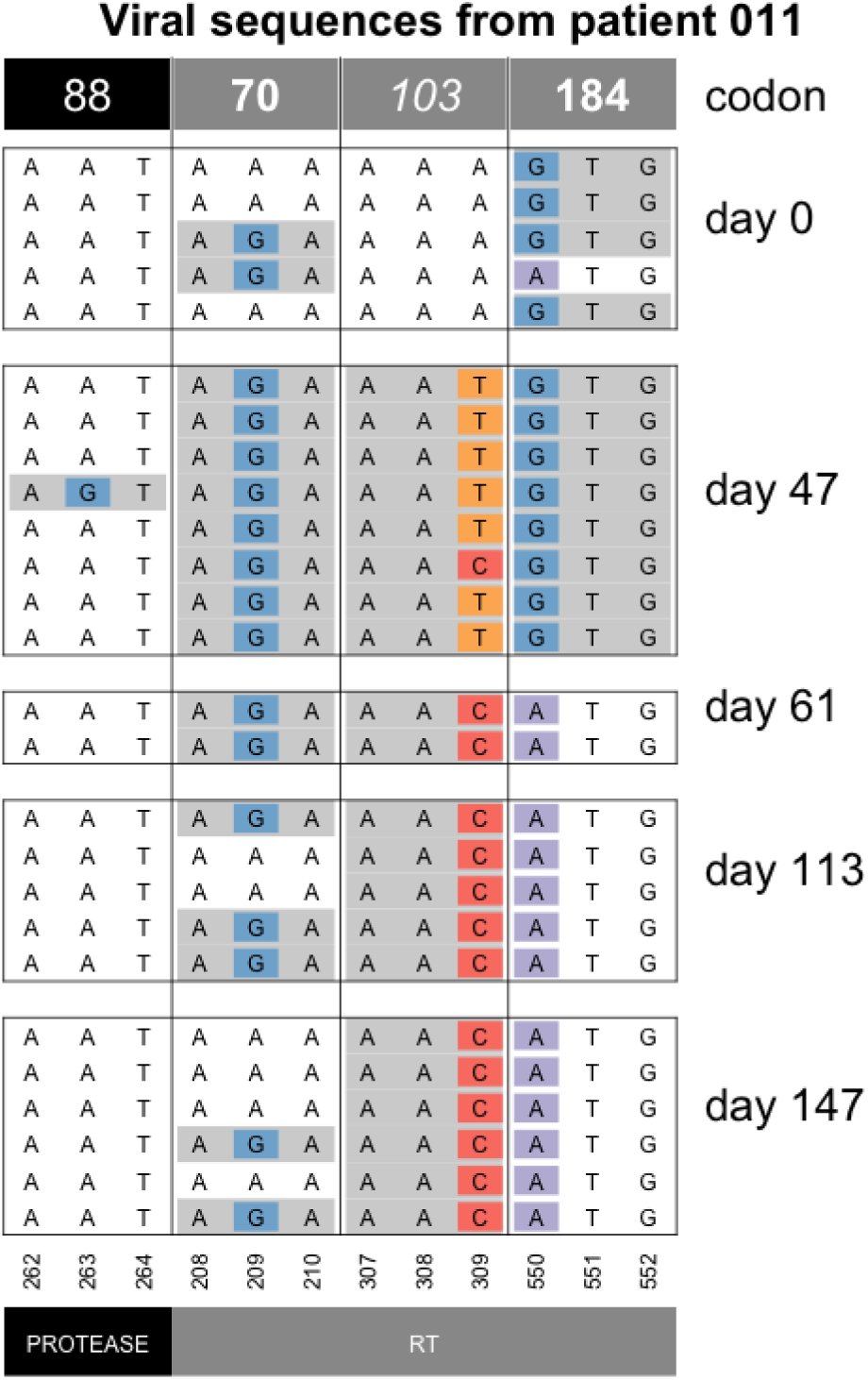
Hardening of soft sweep at K103 in patient 11.

### “Softening” of hard sweeps

We also see several cases of “softening” of hard sweeps. In these cases, a single mutation is initially seen to reach a high frequency or even fix in the population, but the alternative allele appears in later samples (see, for example, patient 73, Figure 11). As far as we are aware, this phenomenon has not been observed before, but unpublished work (Alison Feder, Dmitri Petrov, Joachim Hermisson and Pleuni Pennings, in preparation), predicts that this pattern can occur when selection and migration are both strong and sampling is done from a single compartment (or deme, subpopulation). What happens is that different beneficial mutations can occur and reach high frequencies in different compartments, but soon afterwards, migration mixes genetic material from the different compartments, so that other beneficial mutations appear in the sampled compartment. In patient 73, the AAC allele is seen fixed on day 56, presumably because the allele fixed in the blood compartment, which is where the samples in this study are from. From day 113, however, the other allele (AAT) is also present. Presumably, this happened because the AAT allele fixed in one or several other compartments, and the AAT alleles that are observed from day 113 are migrants from one or more other, unsampled, compartments. The phenomenon of “softening” of hard sweeps is seen three times in the current dataset. Twice, we see the AAC allele first (patient 73 (Figure 11) and 70 (Figure 12), and one time we see the AAT allele first (patient 93, see Figure 13).

**Figure 11:**
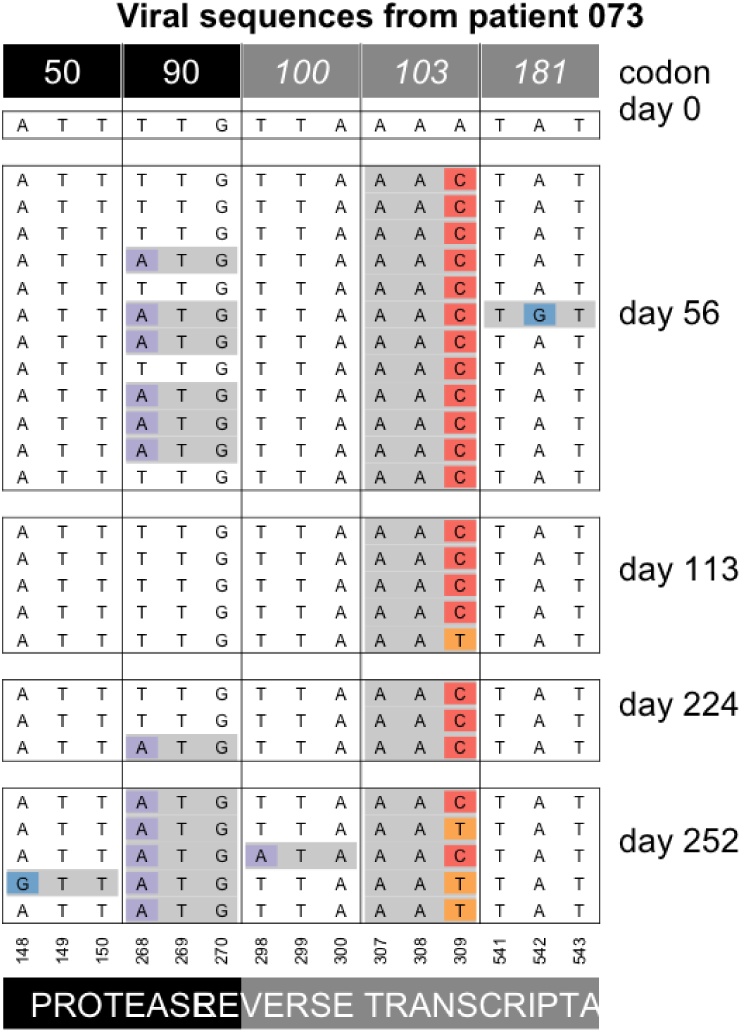
Softening of a hard sweep at K103N in patient 73.

**Figure 12:**
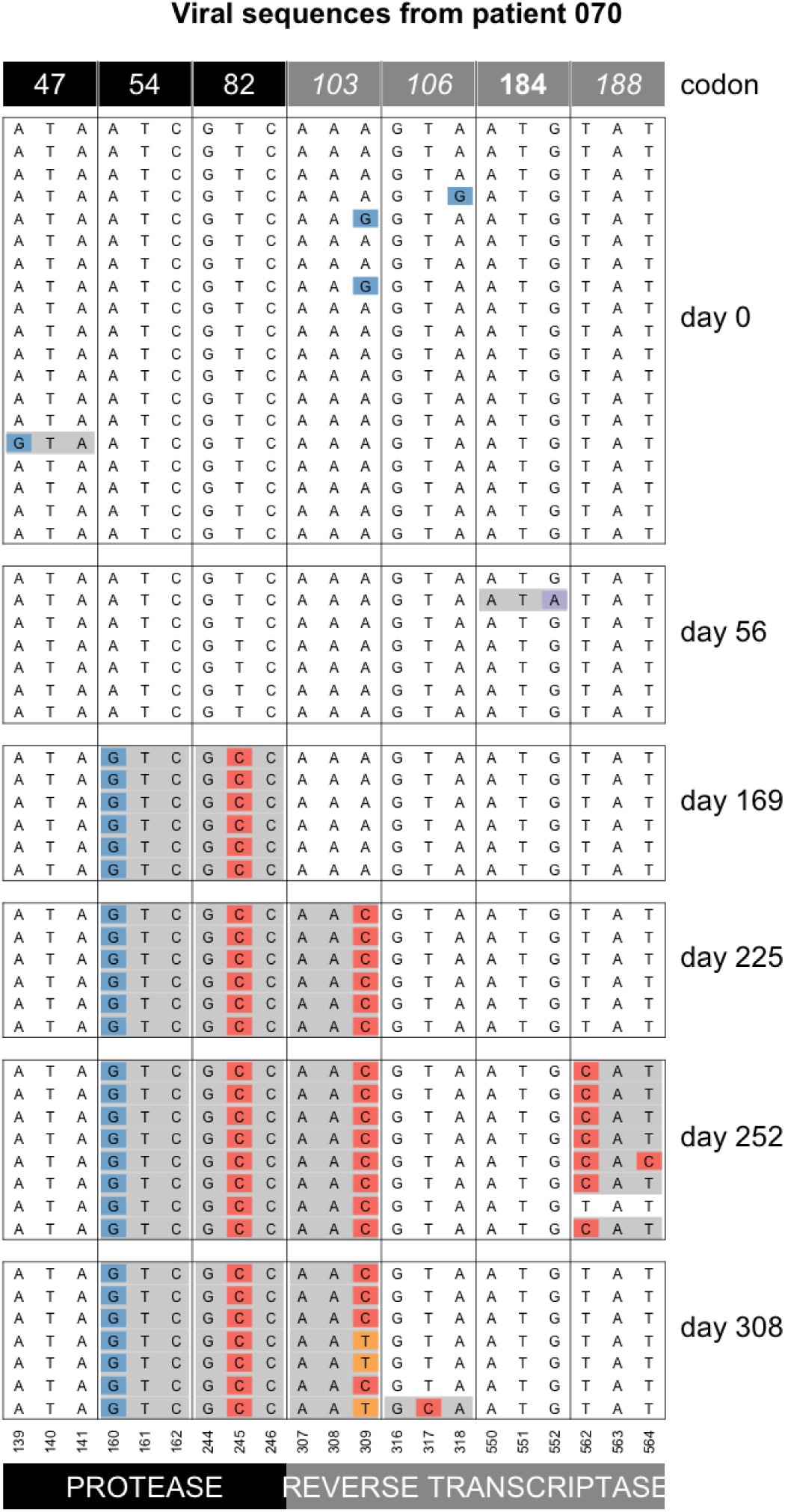
Softening of a hard sweep at K103N in patient 70.

**Figure 13:**
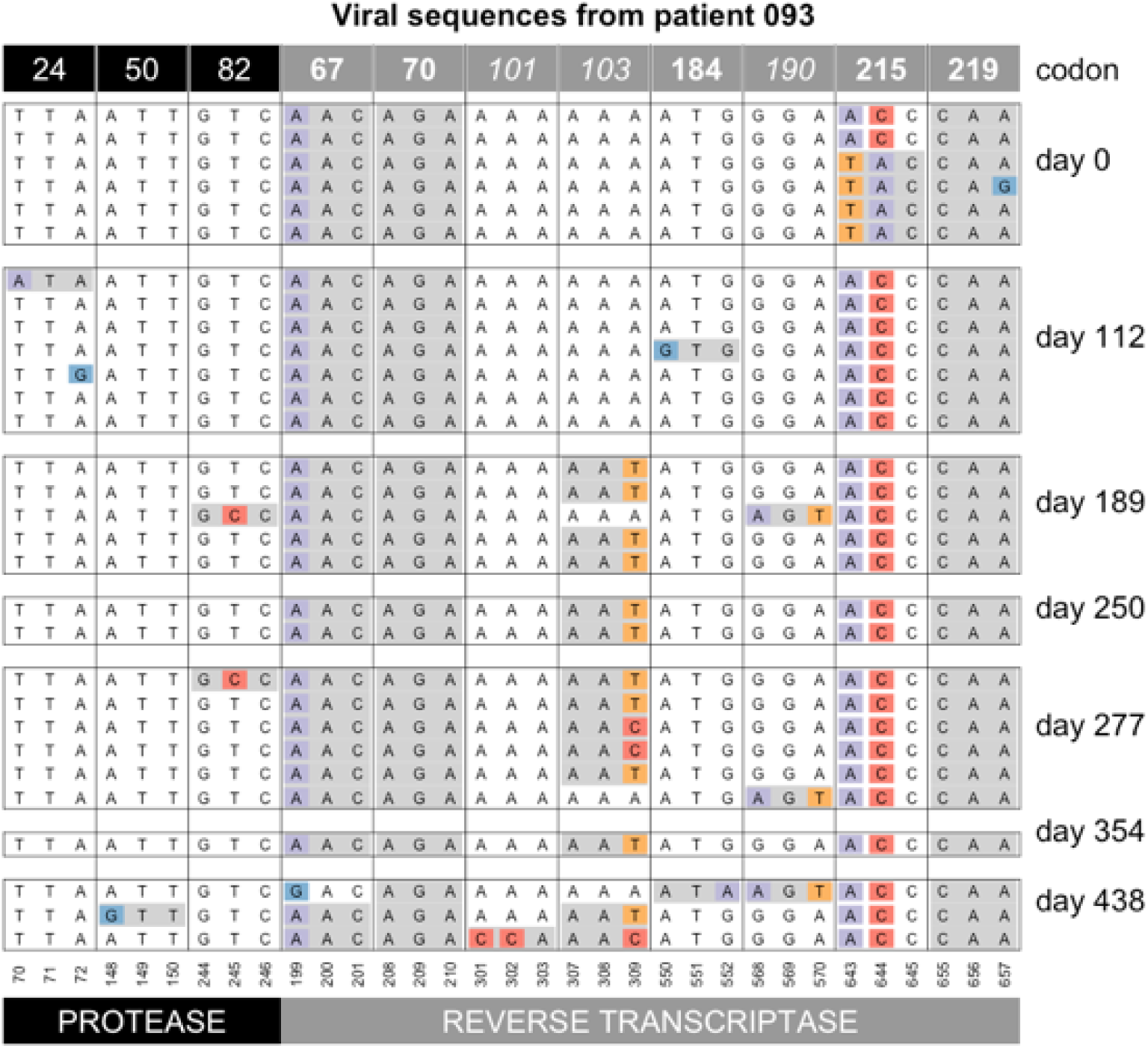
Softening of a hard sweep at K103N in patient 93.

## 6 Sweeps at multiple sites

### Two mutations sweep simultaneously

In a previous paper that was also based on data from the Bacheler dataset, we argued that most sweeps involve a single beneficial mutation (Pennings et al., 2014). It should be noted that even if this is strictly true, sampling may not be done often enough to observe each individual sweep. For example, in patient 70 (Figure 12, resistance mutations at protease codons 54 and 82 are fixed in the population on day 169, even though they were absent in the previous sample (day 56). The data cannot tell us whether these two mutations fixed at the same time or one after the other. However, in the current dataset, we also have two examples of mutations that can be observed to sweep simultaneously. In both cases, we see the simultaneous sweep before it reaches fixation. In patient 84, mutations M46I and L76V in protease are not seen at day 0 (Figure 14), but on day 83, they both appear on 5 out of 7 sequences in perfect linkage disequilibrium. Later, on day 112, they are both fixed.

**Figure 14:**
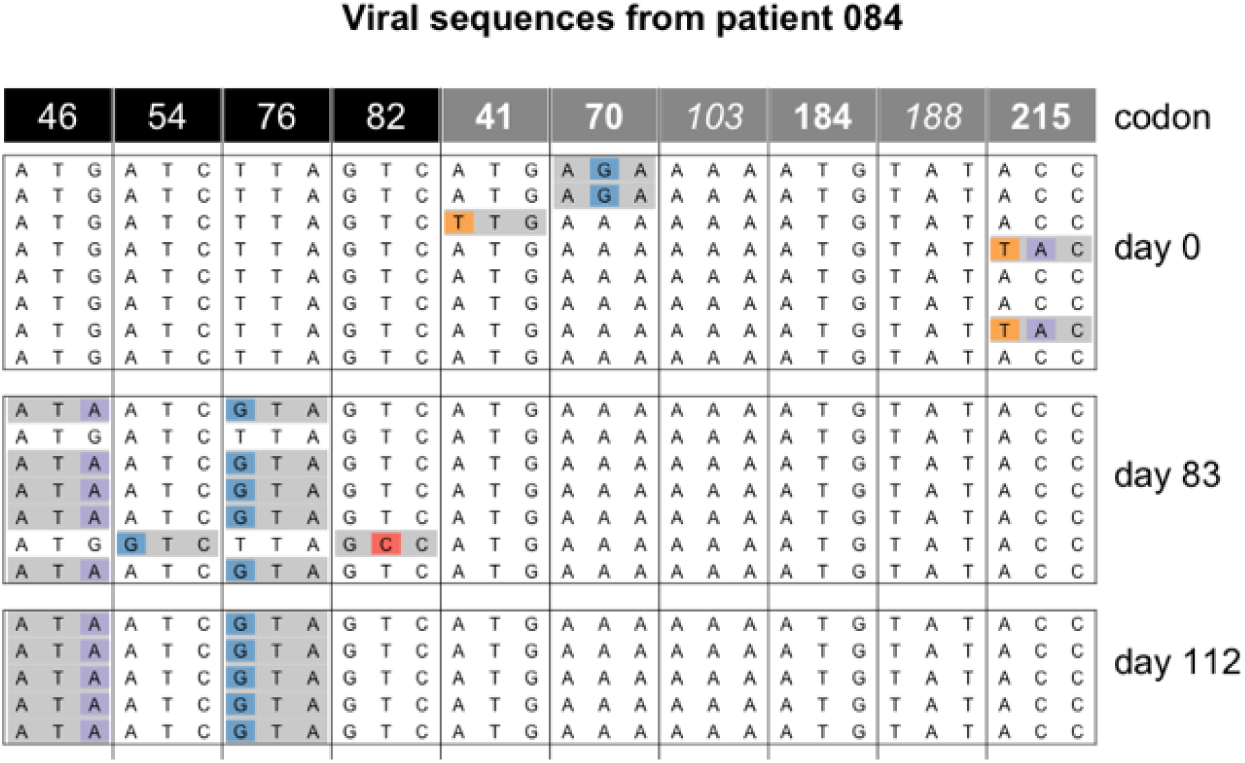
Simultaneous sweep of M46I and L76V in protease in patient 84.

The second example of a simultaneous sweep is seen in patient 100 (Figure 15). Mutations D67N and K70R appear to be sweeping simultaneously in perfect linkage disequilibrium, though in this case we don’t observe the start of the sweep. These two examples of simultaneous sweeps are important because they show that it is possible for multiple mutations to sweep simultaneously even if this is not the most common mode of adaptation. Note that in both cases, the two mutations involved confer resistance to one class of drugs (two PI mutations in the first example and two NRTI mutations in the second example), and they are mutations that are often seen to occur together, presumably because of an epistatic interaction between them.

**Figure 15:**
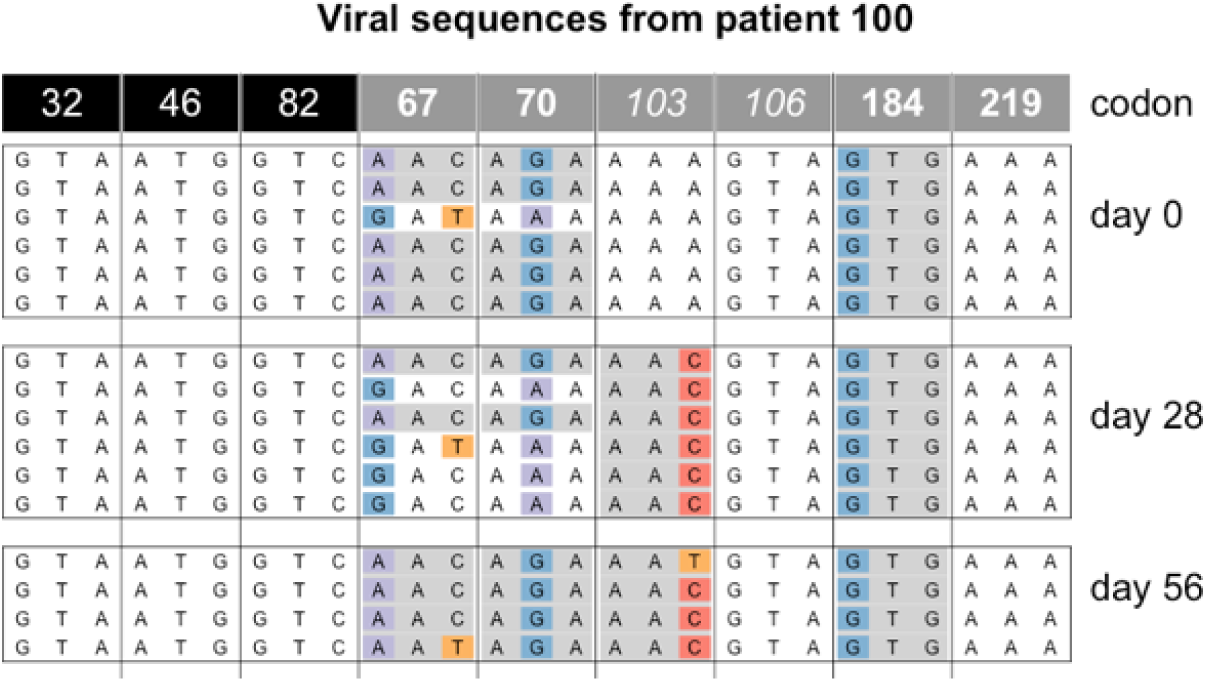
Simultaneous sweep at D67N and K70R in patient 100.

### Accumulation of resistance mutations

We see several examples of several sweeps happening one after the other, leading to an accumulation of resistance mutations in the genome, a pattern that is sometimes called periodic selection (Atwood et al., 1951). For example, in patient 94, the day 0 sample shows resistance only at position 41, but in the two years that this patient was followed four new mutations are acquired by the virus (Table 1 and Figure 16). All sweeps are seen as complete sweeps, except the resistance mutation at codon 215 which is seen in 6 out of 7 sequences at the last time point (day 676).

**Table 1:** Accumulation of resistance mutations in patient 94.

| Patient 94 |  |
| --- | --- |
| Day | DRMs |
| 0 | 41 |
| 186 | 41, 103 |
| 242 | 41, 103, 82 |
| 270 | 41, 103, 82 |
| 375 | 41, 103, 82, 54 |
| 403 | 41, 103, 82, 54 |
| 487 | 41, 103, 82, 54 |
| 676 | 41, 103, 82, 54, 215 |

**Figure 16:**
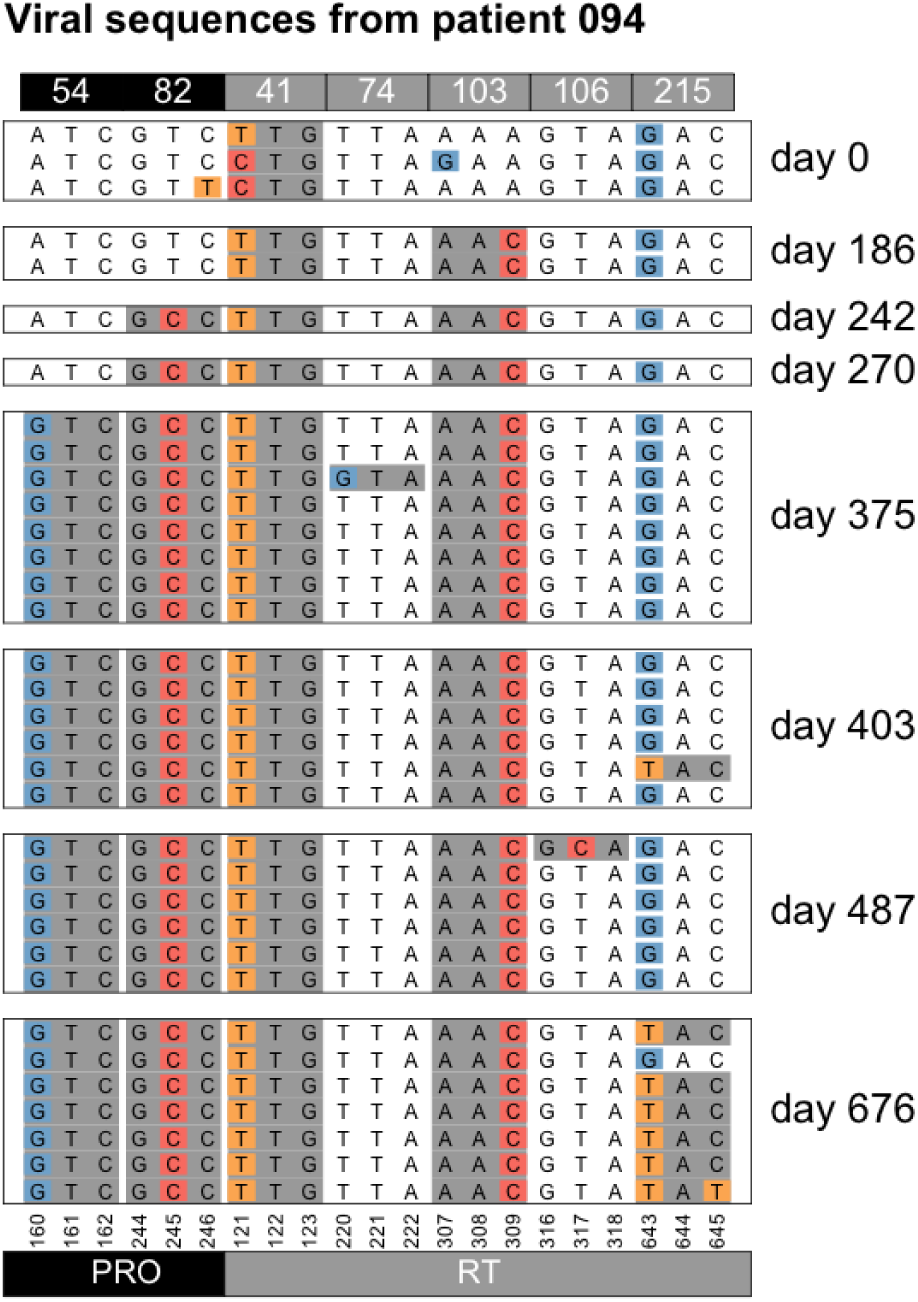
Accumulation of resistance mutations in patient 94.

**Figure 17:**
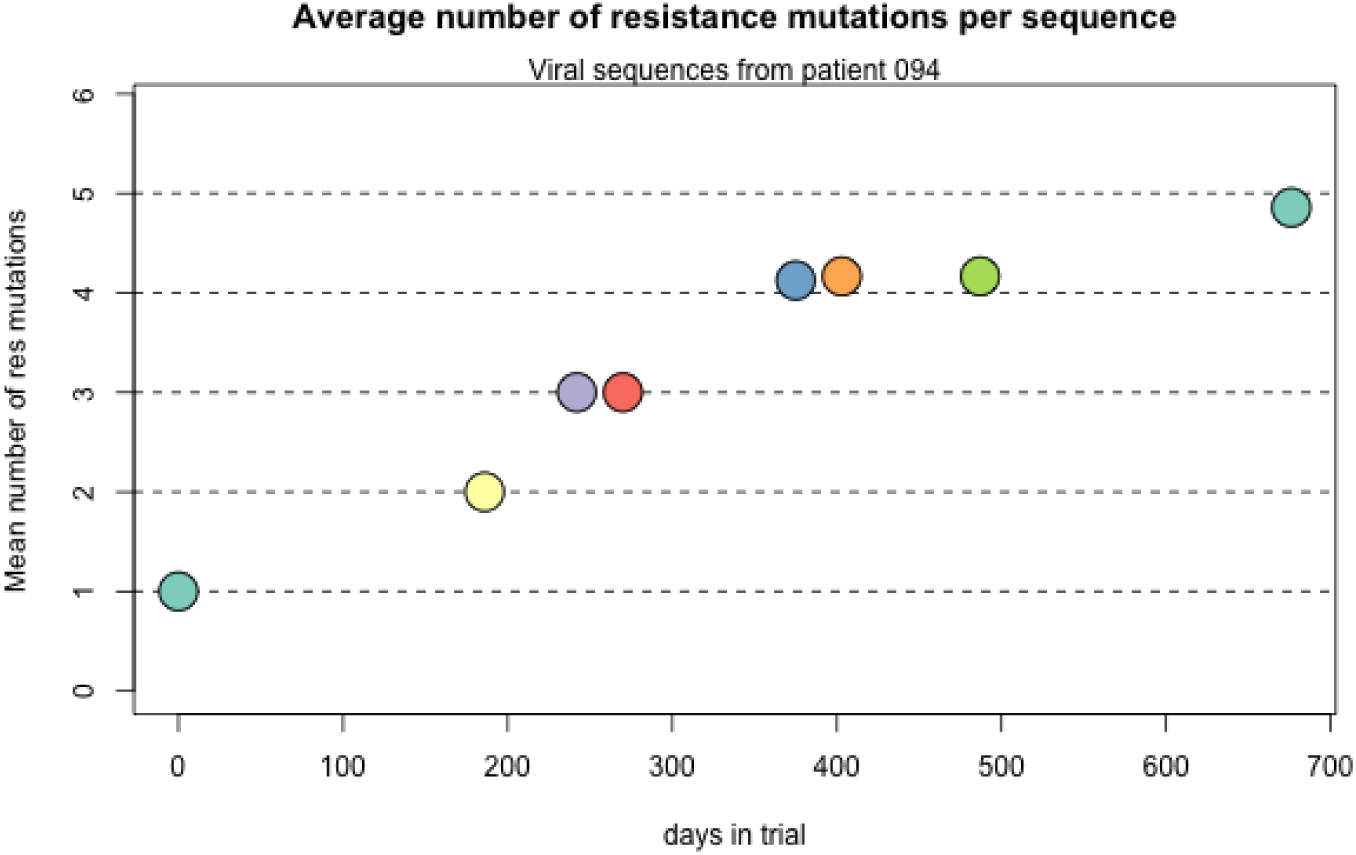
Accumulation of resistance mutations in patient 94.

Data from patient 21 provides another example of accumulation of resistance mutations (Figures 18 and 19 and Table 2). In this case however, there are two time points where two mutations are observed as fixed at the same time (PI46 and PI82 at day 229 and RT24 and RT184 at day 489). We cannot know whether these sweeps were simultaneous or whether we simply missed the time point where one was fixed, but not the other. In addition, at the last time point, a mutation at PI54 appears in perfect linkage disequilibrium with the mutation at PI24. Clonal interference may play a role here, such that either the mutation at 24 or the mutation at 54 can fix, but not both.

Data from patient 77 provide a third example of a patient whose viral population accumulates mutations (Figures 20 and 21 and Table 3).

**Table 2:** Accumulation of resistance mutations in patient 21.

| Patient 21 |  |
| --- | --- |
| Day | DRMs |
| 0 | — |
| 55 | 188 (66%) |
| 83 | 188 (100%) |
| 111 | 188 (100%) |
| 229 | 188 (100%), 46, 82 |
| 312 | 188 (100%), 46, 82 |
| 489 | 188 (100%), 46, 82, 24, 184 |
| 670 | 188 (100%), 46, 82, 24 (33%), 184, 54 (66%) |

**Table 3:**
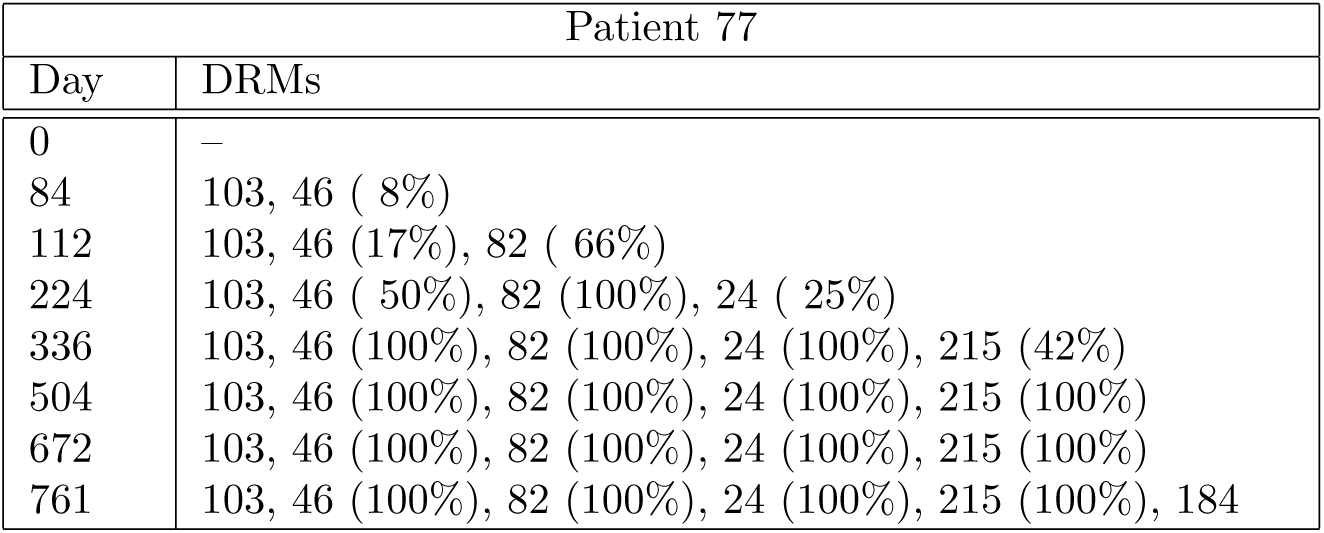
Accumulation of resistance mutations in patient 77.

| Patient 77 |  |
| --- | --- |
| Day | DRMs |
| 0 | — |
| 84 | 103, 46 ( 8%) |
| 112 | 103, 46 (17%), 82 ( 66%) |
| 224 | 103, 46 ( 50%), 82 (100%), 24 ( 25%) |
| 336 | 103, 46 (100%), 82 (100%), 24 (100%), 215 (42%) |
| 504 | 103, 46 (100%), 82 (100%), 24 (100%), 215 (100%) |
| 672 | 103, 46 (100%), 82 (100%), 24 (100%), 215 (100%) |
| 761 | 103, 46 (100%), 82 (100%), 24 (100%), 215 (100%), 184 |

**Figure 18:**
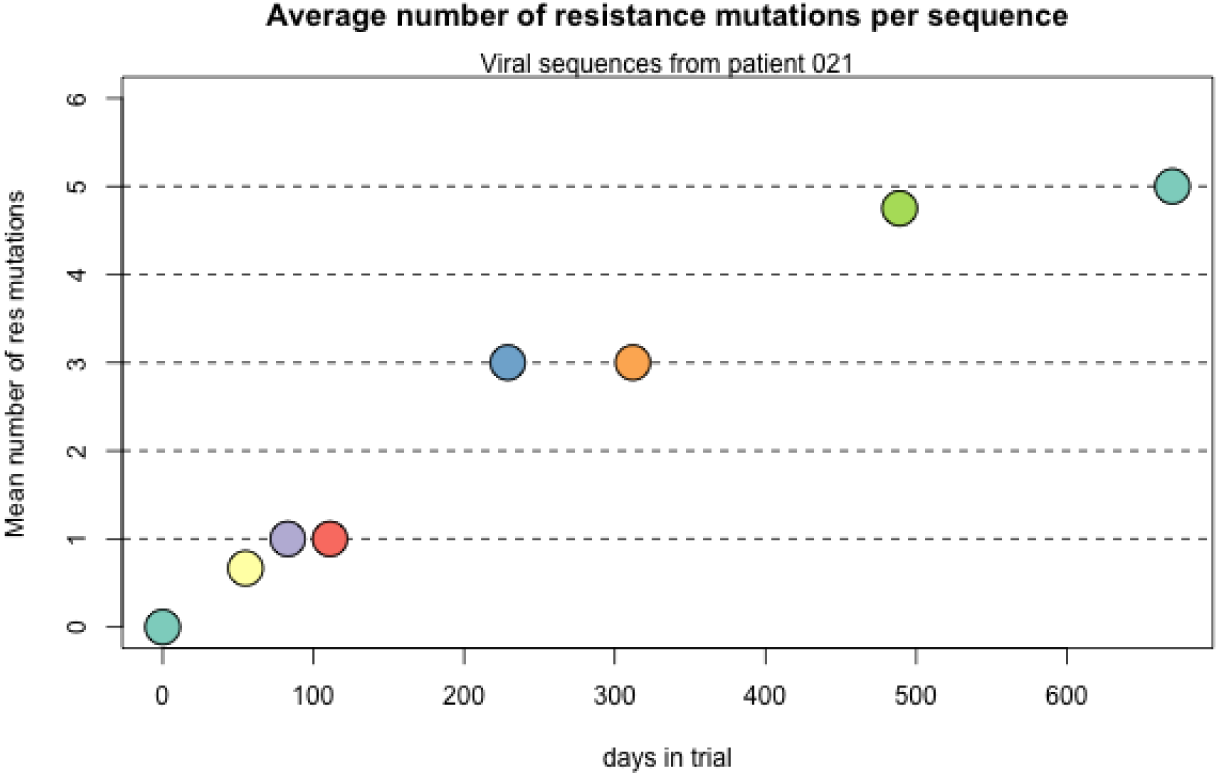
Accumulation of resistance mutations in patient 21.

**Figure 19:**
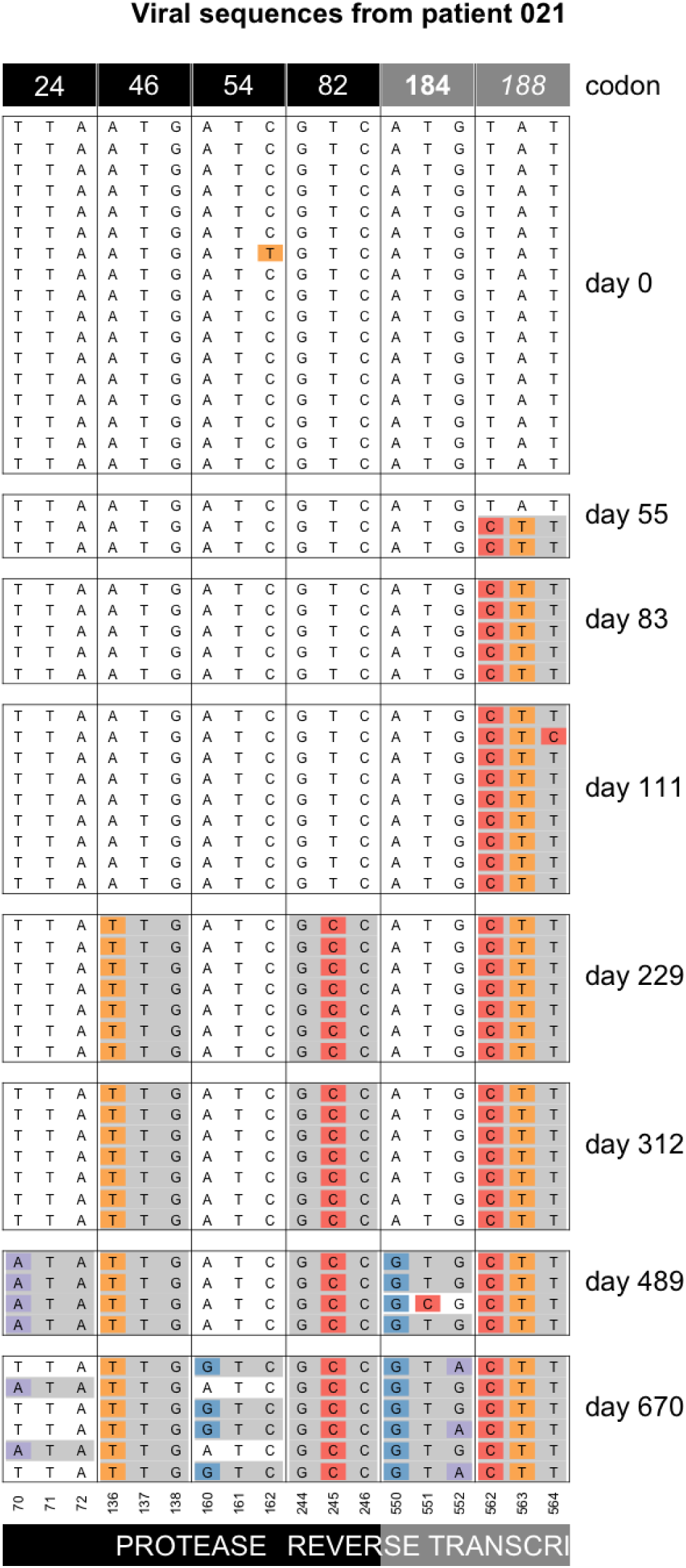
Accumulation of resistance mutations in patient 21.

**Figure 20:**
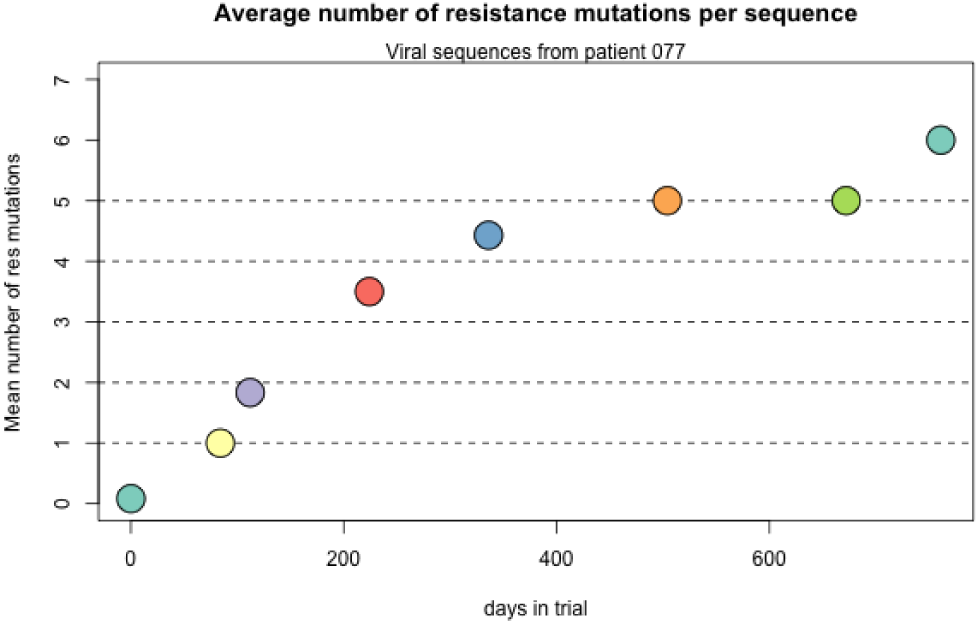
Accumulation of resistance mutations in patient 77.

**Figure 21:**
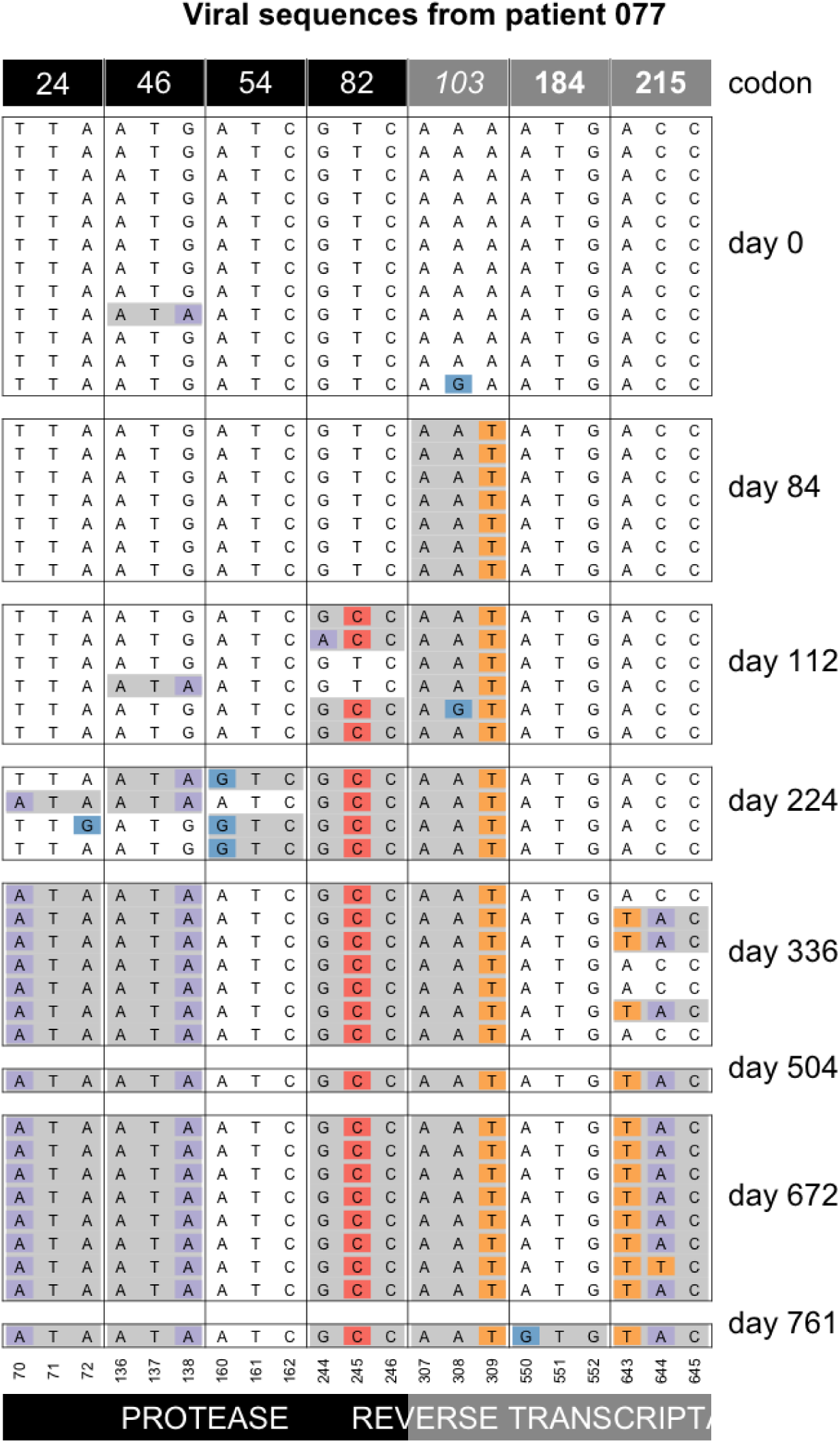
Accumulation of resistance mutations in patient 77.

## 7 Clonal interference

Clonal interference is the effect when multiple beneficial mutations occur in a population, but are not able to recombine onto the same background. They compete with each other because only one or the other can fix. Given that soft sweeps from recurrent mutation happen (which means that the rate of input of new mutations in the population must be fairly high) and that simultaneous sweeps happen (suggesting that mutations can stay linked), it appears likely that clonal interference also happens in adapting HIV populations. We show Muller plots (Muller, 1932) or evolvograms (Angelova et al., 2018) for several examples, where we think that clonal interference is occurring.

### Patient 84, K103N loses against T188L

We have shown the first few timepoints for patient 84 previously (Figure 14). Here we show a Muller diagram (Figure 22) and the sequences for the later time points (Figure 22). Mutations T188L and K103N both lead to drug resistance against NNRTI drugs. Mutation T188L requires two transversion mutations. K103N requires a single transversion mutation. In this patient, the data suggest that K103N occurred first, but but the haplotype with T188L had higher fitness and ultimately fixed. One sequence is observed with both K103N and T188L. In Figure 22 M184V is also shown, which requires a single transition (A →G) and thus occurs de novo much more often as it has a higher mutation rate. M184V is fixed on both the K103N background and the T188L background.

**Figure 22:**
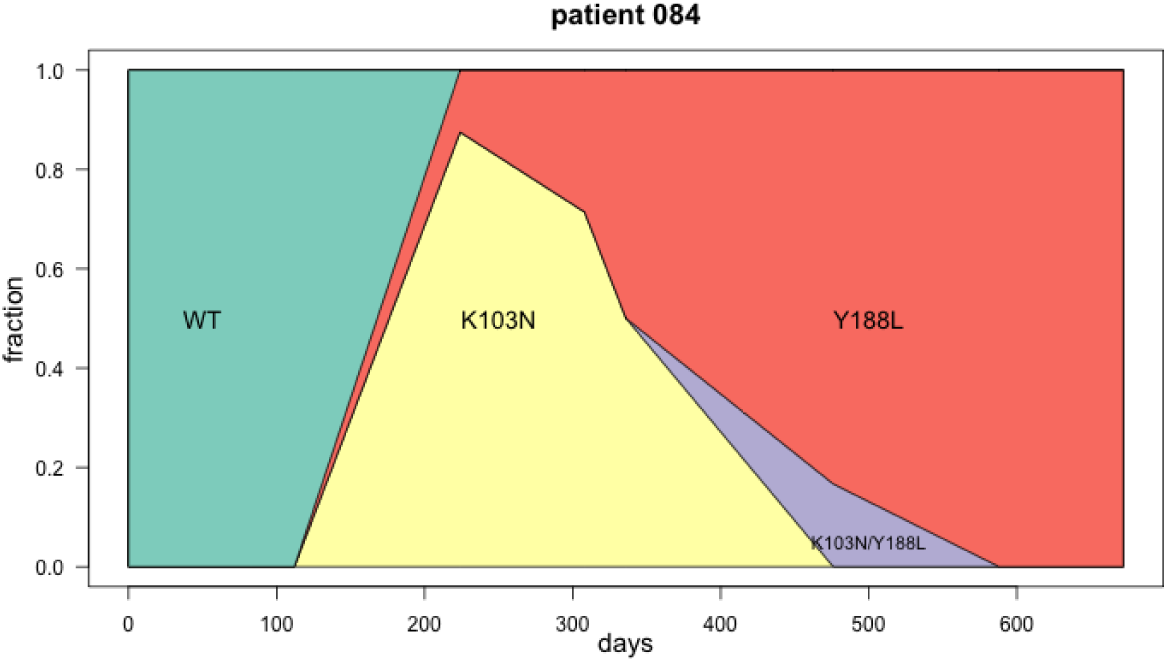
Muller plot showing clonal interference in patient 89.

**Figure 23:**
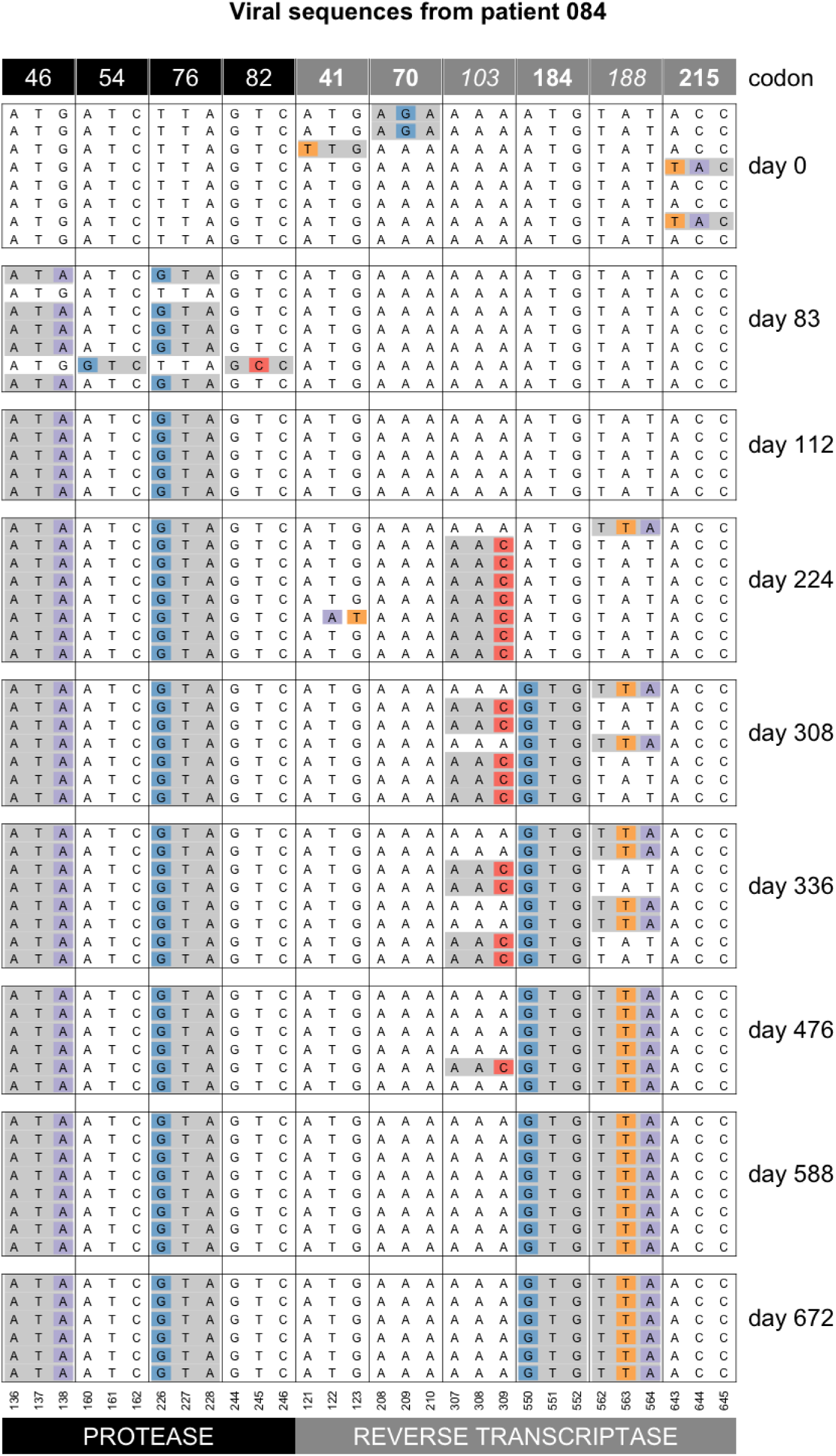
Sequence evolution in patient 84. At day 83, we see resistance mutations at positions 52 and 82 in protease. These mutations are then no longer observed at later time points. On days 224, 308, 336 and 476, we see mutation K103N in RT, it is in complete linkage disequilibrium with T188L. On day 588, T188L has fixed and K103 is no longer observed.

### Patient 26: M41L loses against D67N and K70R

In patient 26, mutation M41L appears and reaches a high frequency of 50%, but then is outcompeted by a genotype with D67N and K70R. M41L is part of the TAM1 pathway and D67N and K70R are part of the TAM2 pathway, and typically patients get mutations from one or the other TAM pathway, but not both.

### Patient 72: N88D loses against V82A and I54V

In patient 72, mutation N88D reaches a high frequency (75% at time point 112) and then disappears. Mutation V82A becomes associated with 46L and 54V and fixes in the population. See Figure 24.

**Figure 24:**
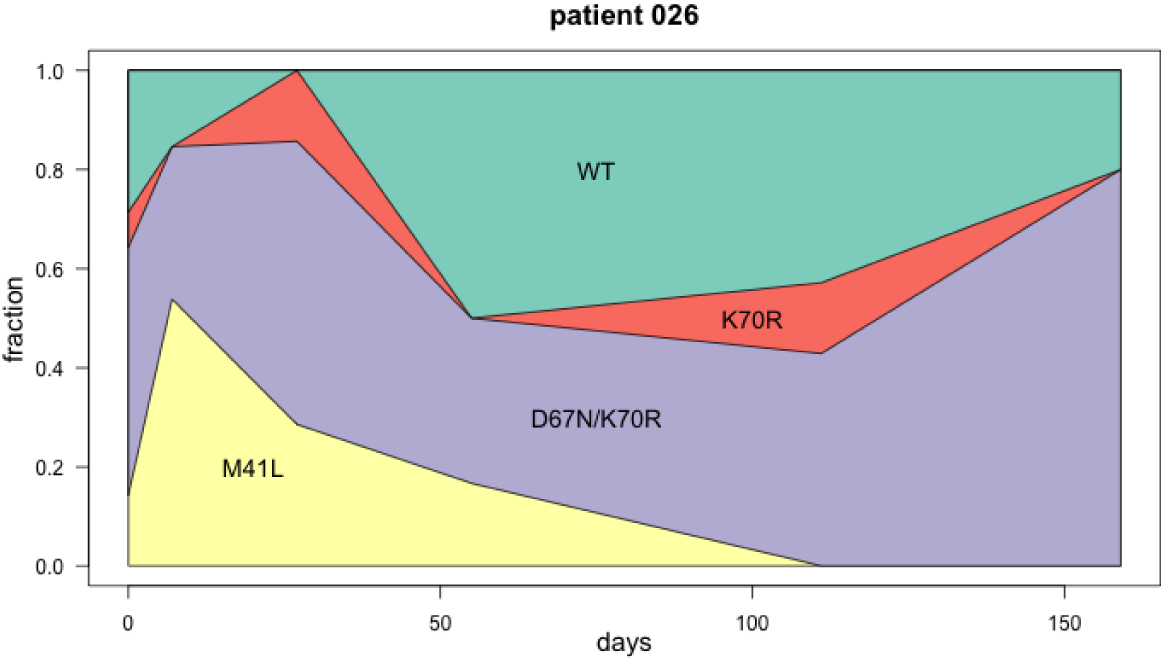
Muller plot showing clonal interference in patient 26.

**Figure 25:**
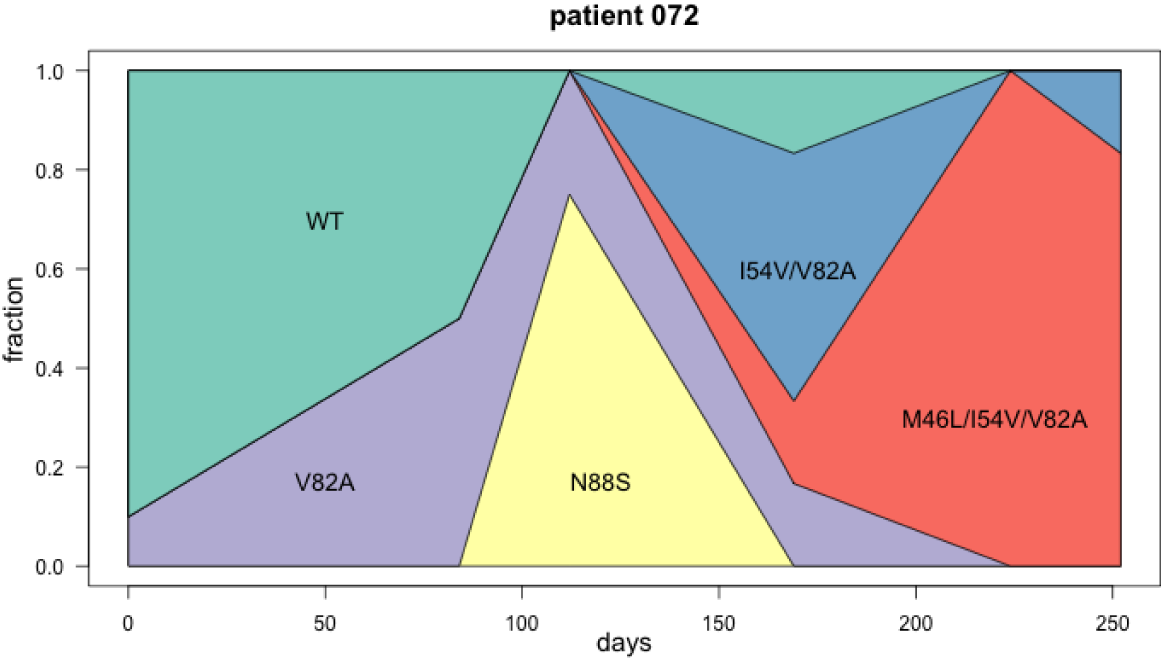
Muller plot showing clonal interference in patient 72.

### Patient 77, I54V loses against L24I

Mutation 54 loses against mutation 24. Both 54 and 24 exist on genotypes with 46, but the combination of 24 and 46 ultimately fixes in the population. Note that the mutation at 24 is the only transversion here, and is thus much rarer than the others. 24, 46, 54 and 82 all confer resistance to several PIs. See Figure 26.

**Figure 26:**
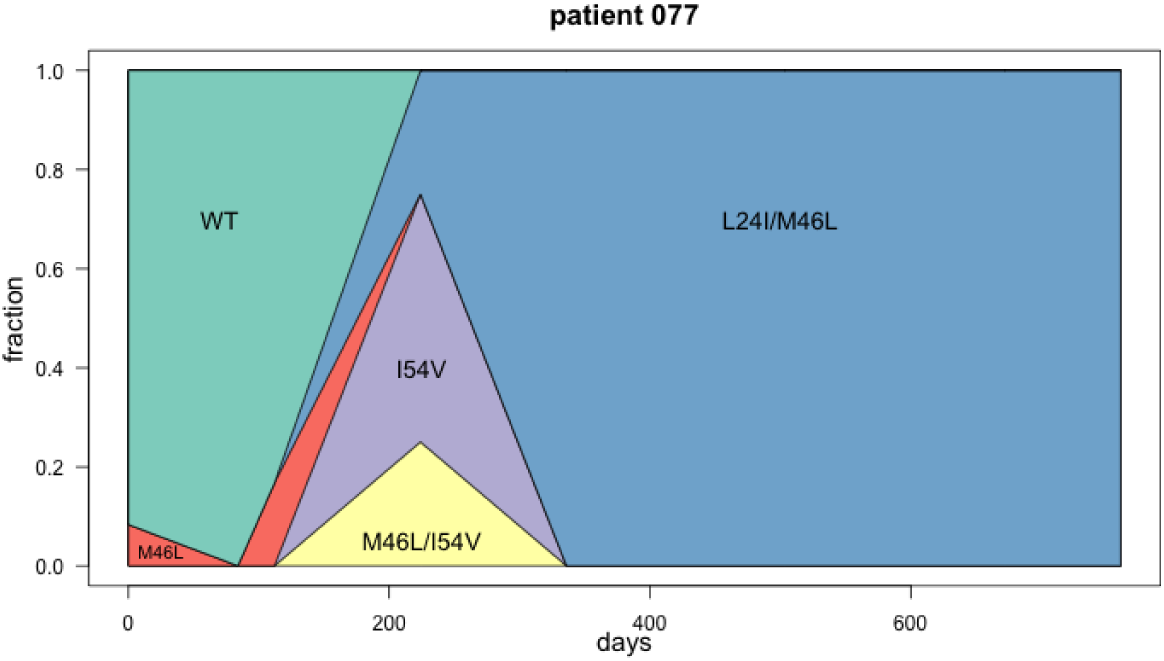
Muller plot showing clonal interference in patient 77.

### Patient 89, K103N loses against K101E

In this case the relevant mutations are NNRTI mutations K103N, G190S and K101E. In this patient, the ancestral codon at 190 is GGC (glycine) which means that the G190S mutation only requires a single G →A mutation to create AGC (Serine). This is uncommon because in most B subtypes, the 190 codon is encoded by GGA (glycine). The 190S mutation reduces susceptibility to Efavirenz by 50-fold (Liu and Shafer, 2006). K101E requires an A →G mutation (AAA →GAA). K103N requires a A →C or A→ T mutation (tranversions), which are much less common than A →G or G→A (transition mutations). We see that genotypes that have both 101E and 190S exist, just like genotypes that have 103N and 190S. There are no genotypes that combine 101E and 103N. This is likely because the mutation rates for 101E and 103N are lower (than for 190S) and the recombination rate between the two must be very low, given that they are only two codons apart. See Figure 27.

**Figure 27:**
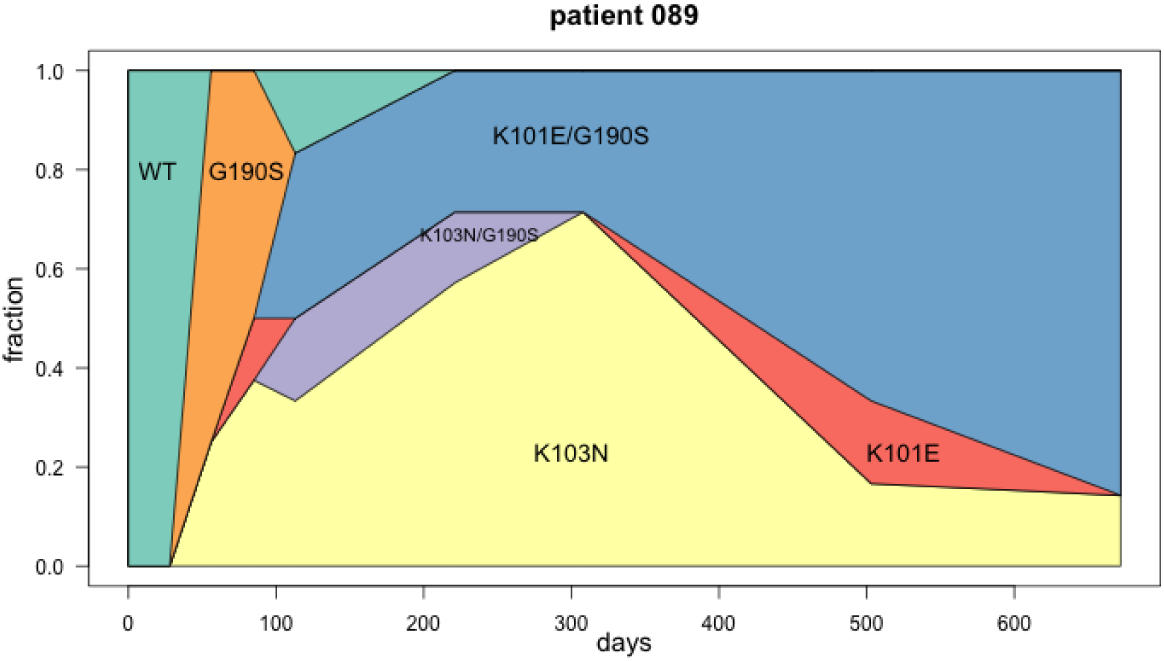
Muller plot showing clonal interference in patient 89.

### Patient 132, K101E loses against L100I

Mutations L100I and K101E occur in neighboring codons. One sequence is observed (at time point 111) that has both mutations. Ultimately, the mutation at 101 disappears and 100 fixes. Not shown here is that at time point 41, K103N fixes (see supplemental figure for patient 132). See Figure 28.

**Figure 28:**
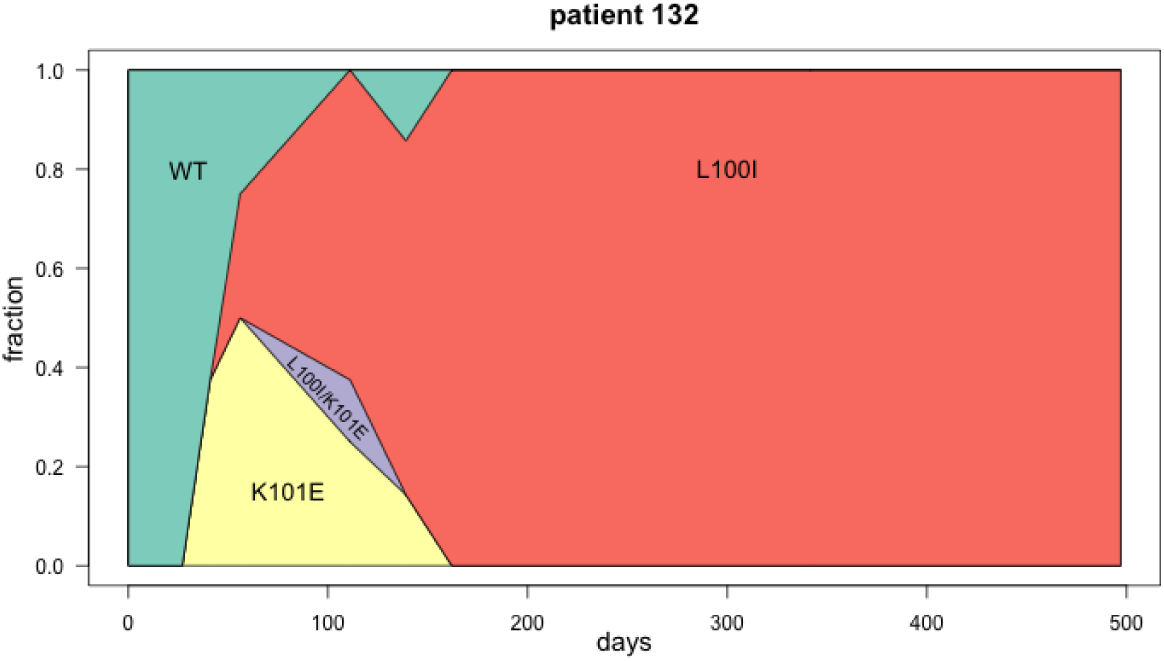
Muller plot showing clonal interference in patient 132.

## 8 Discussion

The goal of this paper is to provide examples of evolutionary dynamics of HIV within patients who are treated with antiretrovirals. We hope that the figures in this paper will be used in evolution and population genetics classes.

The data we used came from a dataset collected by Lee Bacheler and colleagues in the late 1990s. The patients were enrolled in several trials focused on the (then new) drug Efavirenz. The dataset contains sequences from 170 patients, but we focused on 118 patients with at least two sampling dates, and at least 5 sequences in total. We provide a supplement with figures showing all polymorphic nucleotides for all 118 patients. A caveat is that the treatment history of the patients is not always known. We have reason to believe that the annotation in the Genbank entries for the sequences is not always correct. Still, we included the treatment annotation in the supplemental figures.

In conclusion, we show a wide variety of patterns, specifically: soft sweeps, hard sweeps, softening sweeps and hardening sweeps, simultaneous sweeps, accumulation of mutations and clonal interference.

## Supporting information

Supplemental Figures for all 118 patients

## 9 Acknowledgements

We thank Noah Whiteman, Claudia Bank and Alison Feder for discussions.

