## Supplemental Figures for all 118 patients for "Drug resistance evolution in HIV in the late 1990s: hard sweeps, soft sweeps, clonal interference and the accumulation of drug resistance mutations"

#### Viral sequences from patient 001

( likely treated with indinavir then later switched to efavirenz combination therapy )

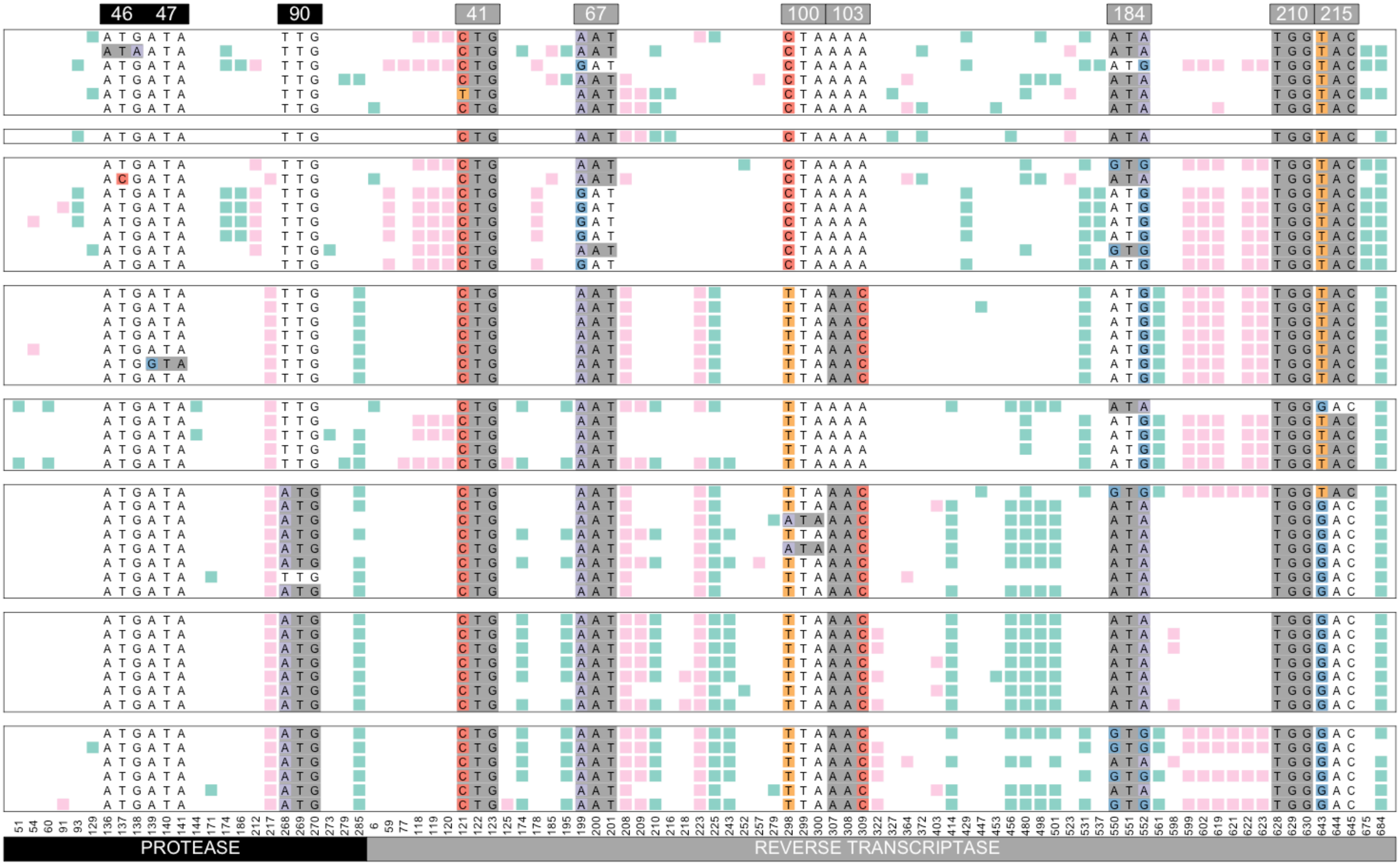


Williams, Pennings, 2018. Data from Bacheler et al 2000

#### Viral sequences from patient 003

( likely treated with indinavir + efavirenz )

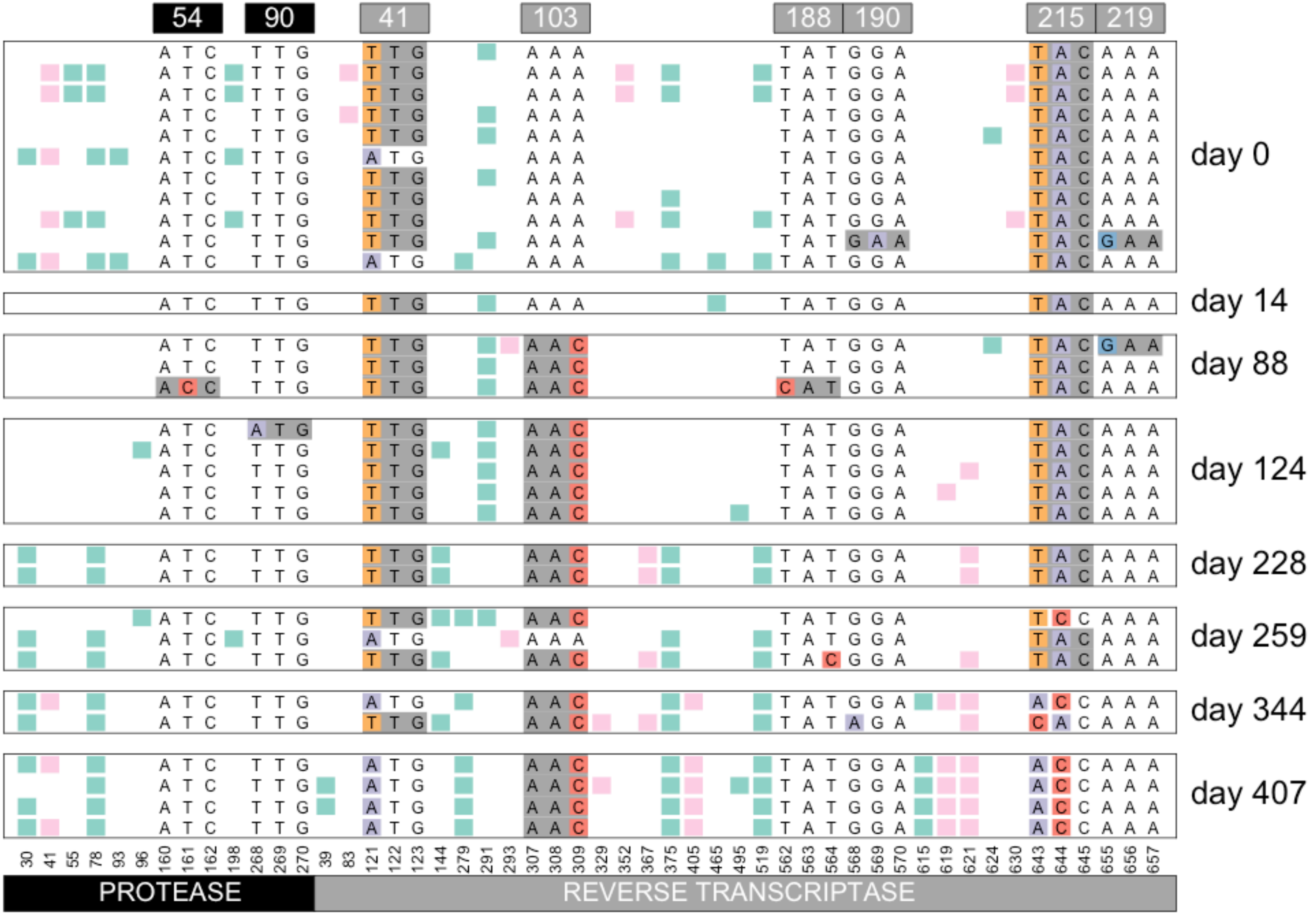


Williams, Pennings, 2018. Data from Bacheler et al 2000

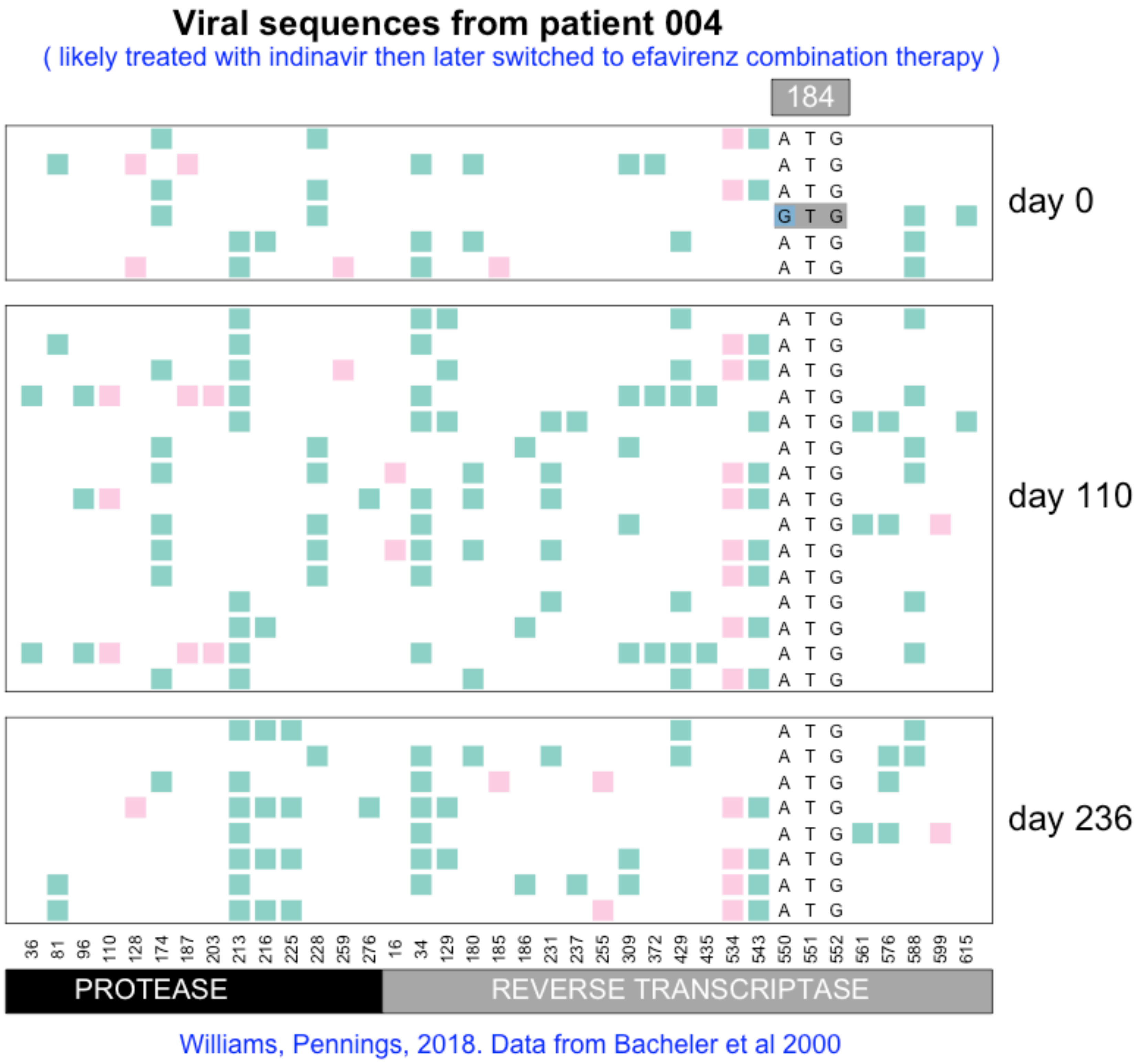

Viral sequences from patient 004
( likely treated with indinavir then later switched to efavirenz combination therapy )
184
day 0
day 110
day 236
36 81 96 110 128 174 187 203 213 216 225 228 259 276 16 34 129 180 185 186 231 237 255 309 372 429 435 534 543 550 551 552 561 576 588 599 615
PROTEASE
REVERSE TRANSCRIPTASE
Williams, Pennings, 2018. Data from Bacheler et al 2000

#### Viral sequences from patient 005

( likely treated with indinavir then later switched to efavirenz combination therapy )

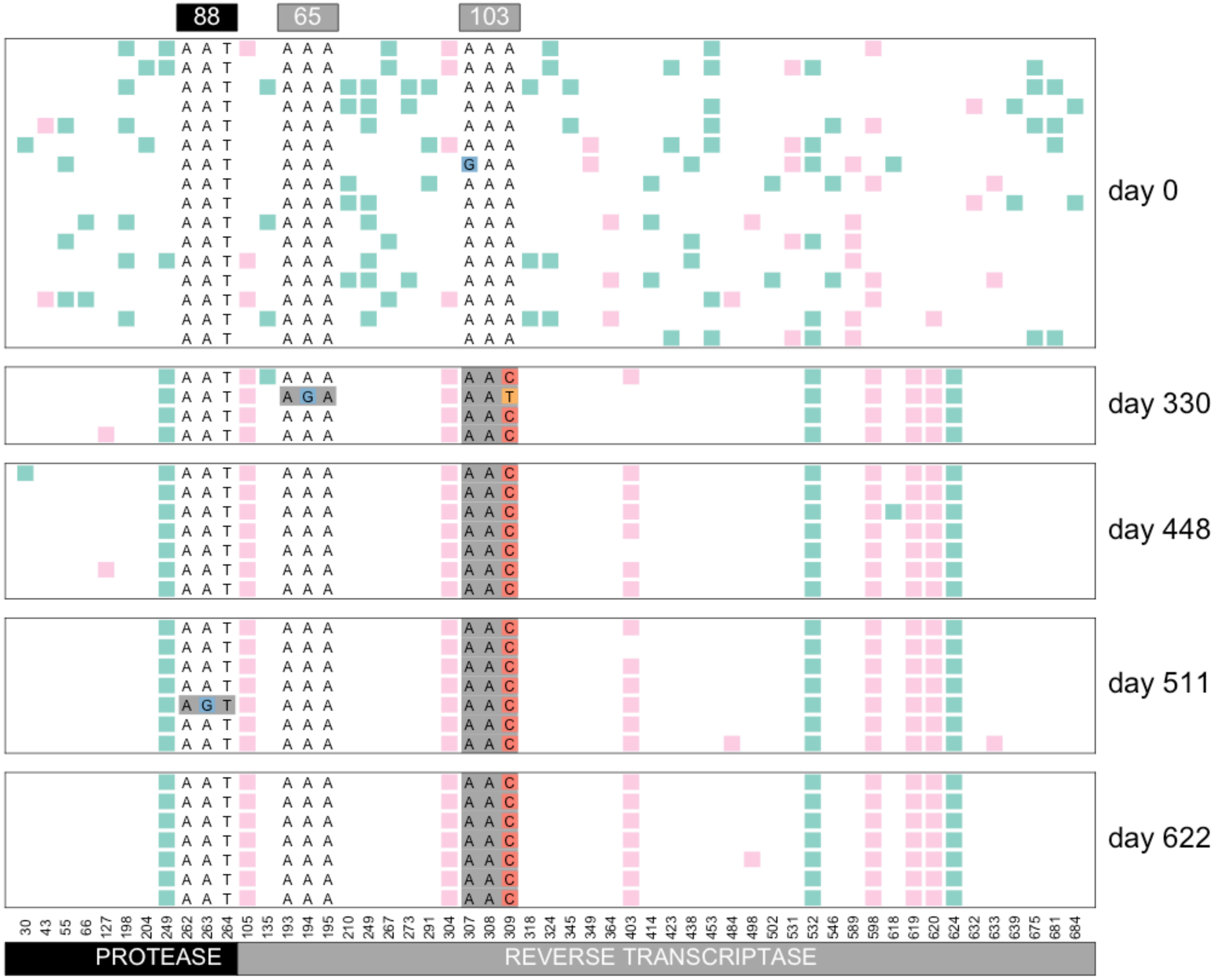


Williams, Pennings, 2018. Data from Bacheler et al 2000

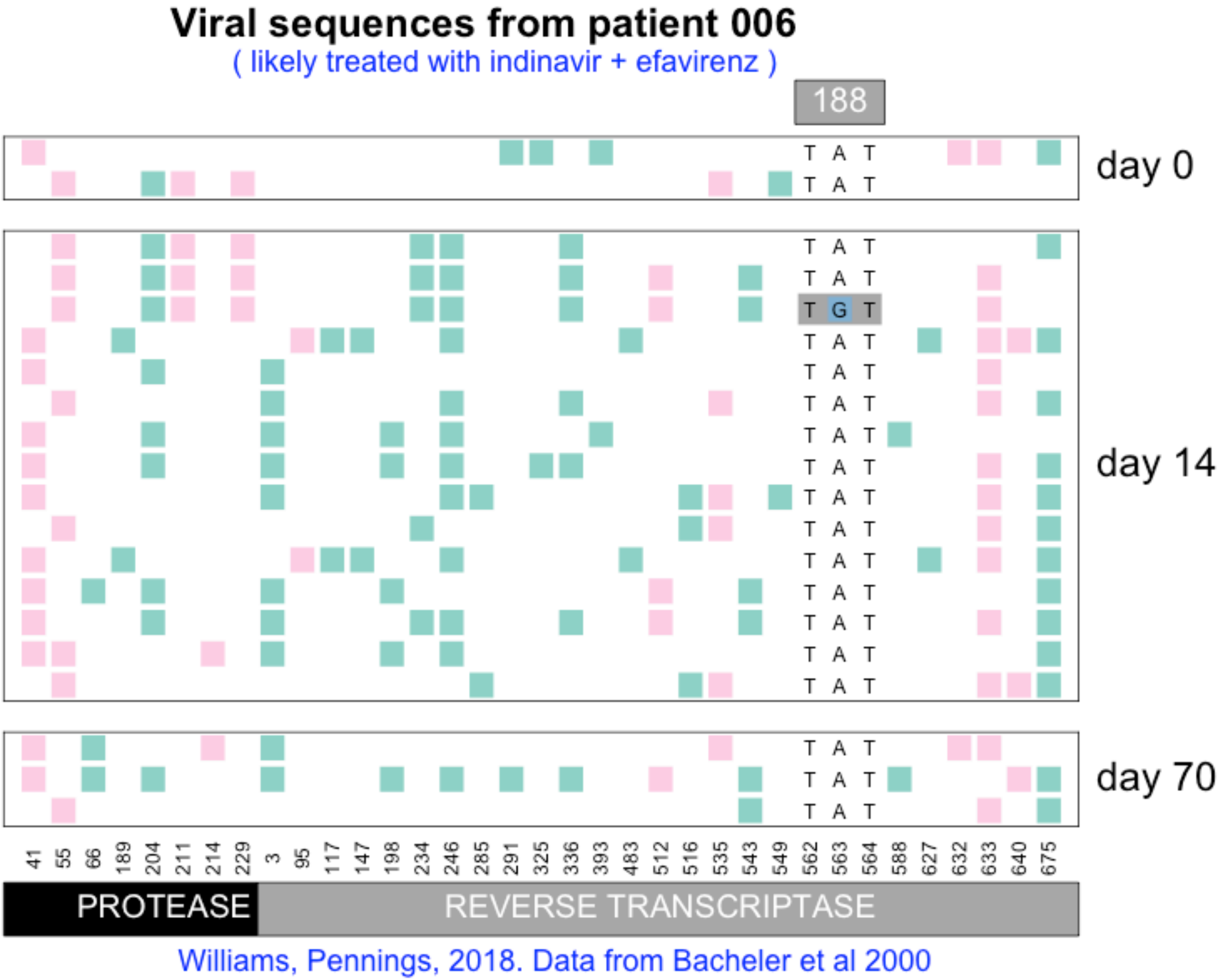

Viral sequences from patient 006
( likely treated with indinavir + efavirenz )
188
day 0
day 14
day 70
T A T
T G T
41 55 66 189 204 211 214 229 3 95 117 147 198 234 246 285 291 325 336 393 483 512 516 535 543 549 562 563 564 588 627 632 633 640 675
PROTEASE
REVERSE TRANSCRIPTASE
Williams, Pennings, 2018. Data from Bacheler et al 2000

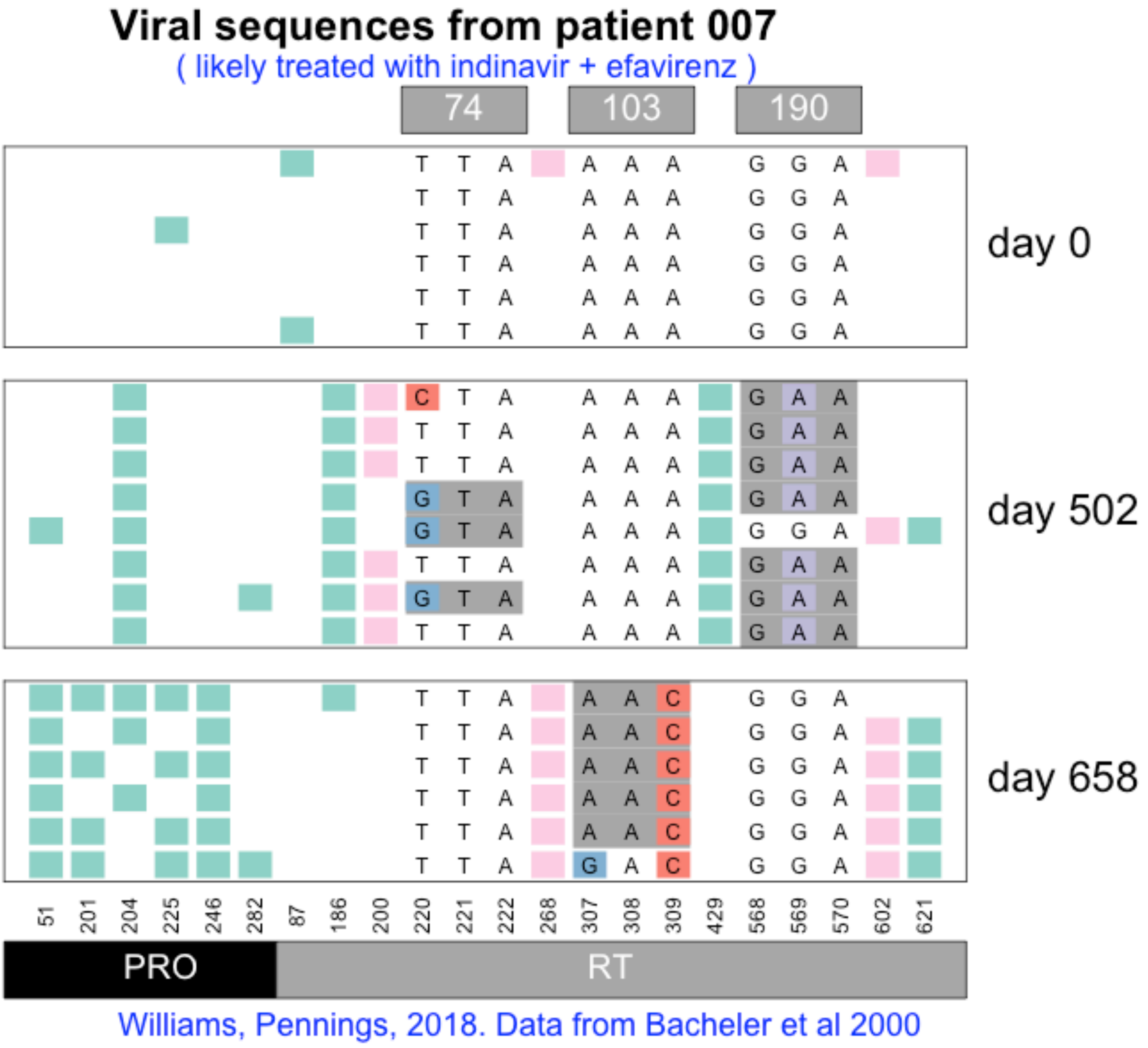
Viral sequences from patient 007
( likely treated with indinavir + efavirenz )
74
103
190
day 0
day 502
day 658
51
201
204
225
246
282
87
186
200
220
221
222
268
307
308
309
429
568
569
570
602
621
PRO
RT
Williams, Pennings, 2018. Data from Bacheler et al 2000

#### Viral sequences from patient 008

( likely treated with indinavir then later switched to efavirenz combination therapy )

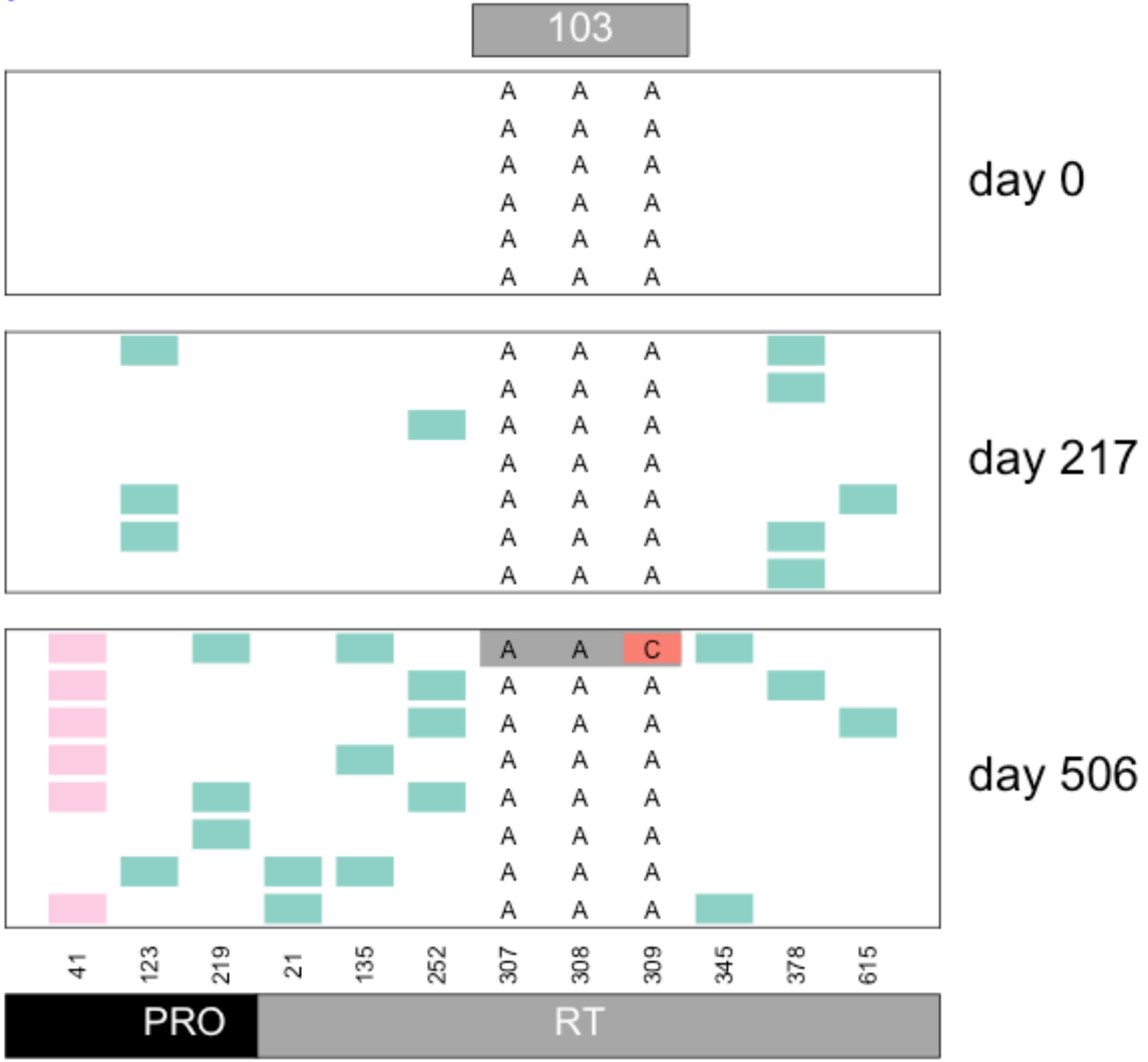


Williams, Pennings, 2018. Data from Bacheler et al 2000

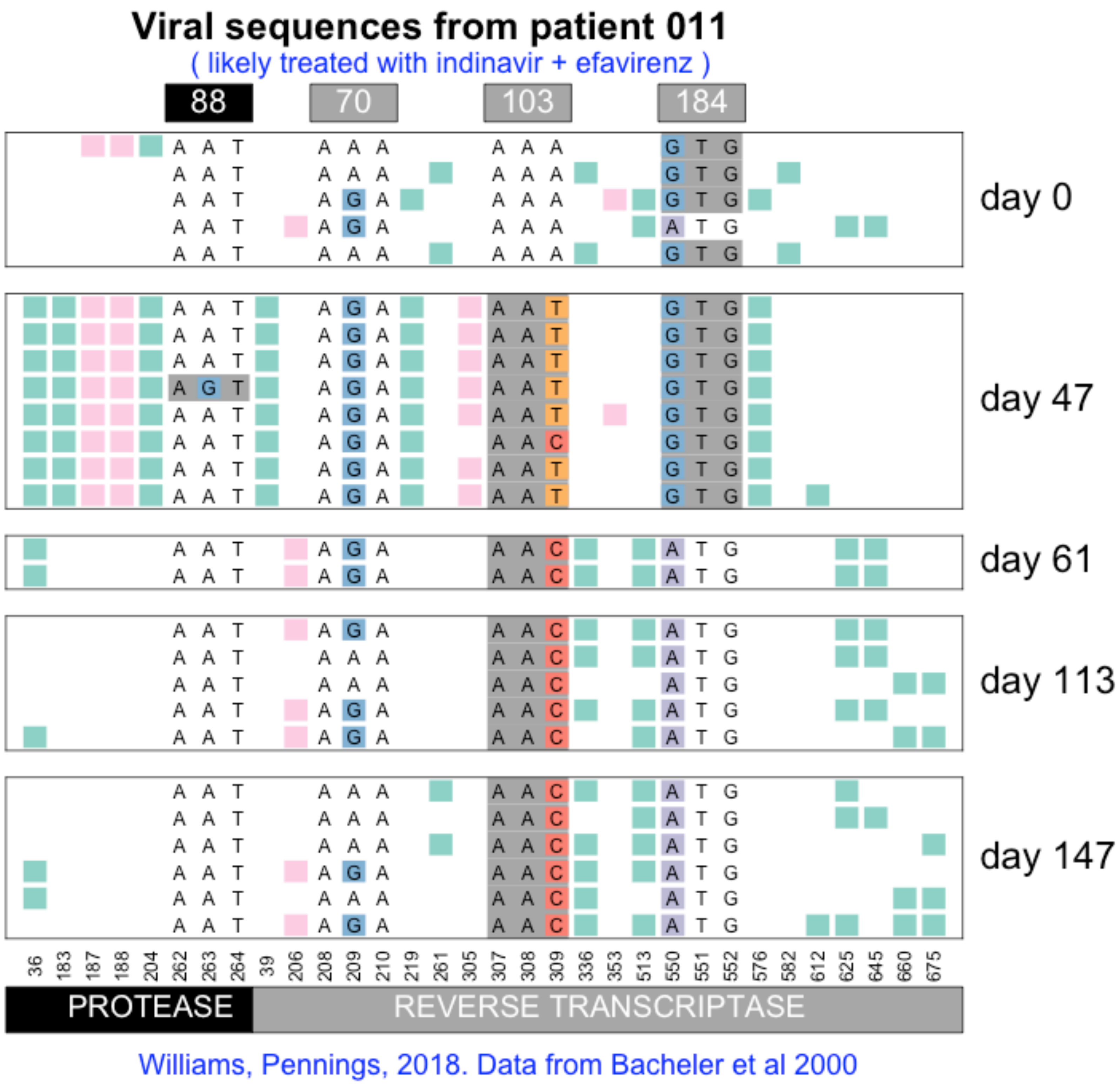
Viral sequences from patient 011
( likely treated with indinavir + efavirenz )
88
70
103
184
day 0
day 47
day 61
day 113
day 147
36 183 187 188 204 262 263 264 39 206 208 209 210 219 261 305 307 308 309 336 353 513 550 551 552 576 582 612 625 645 660 675
PROTEASE
REVERSE TRANSCRIPTASE
Williams, Pennings, 2018. Data from Bacheler et al 2000

### Viral sequences from patient 012

( likely treated with indinavir + efavirenz )

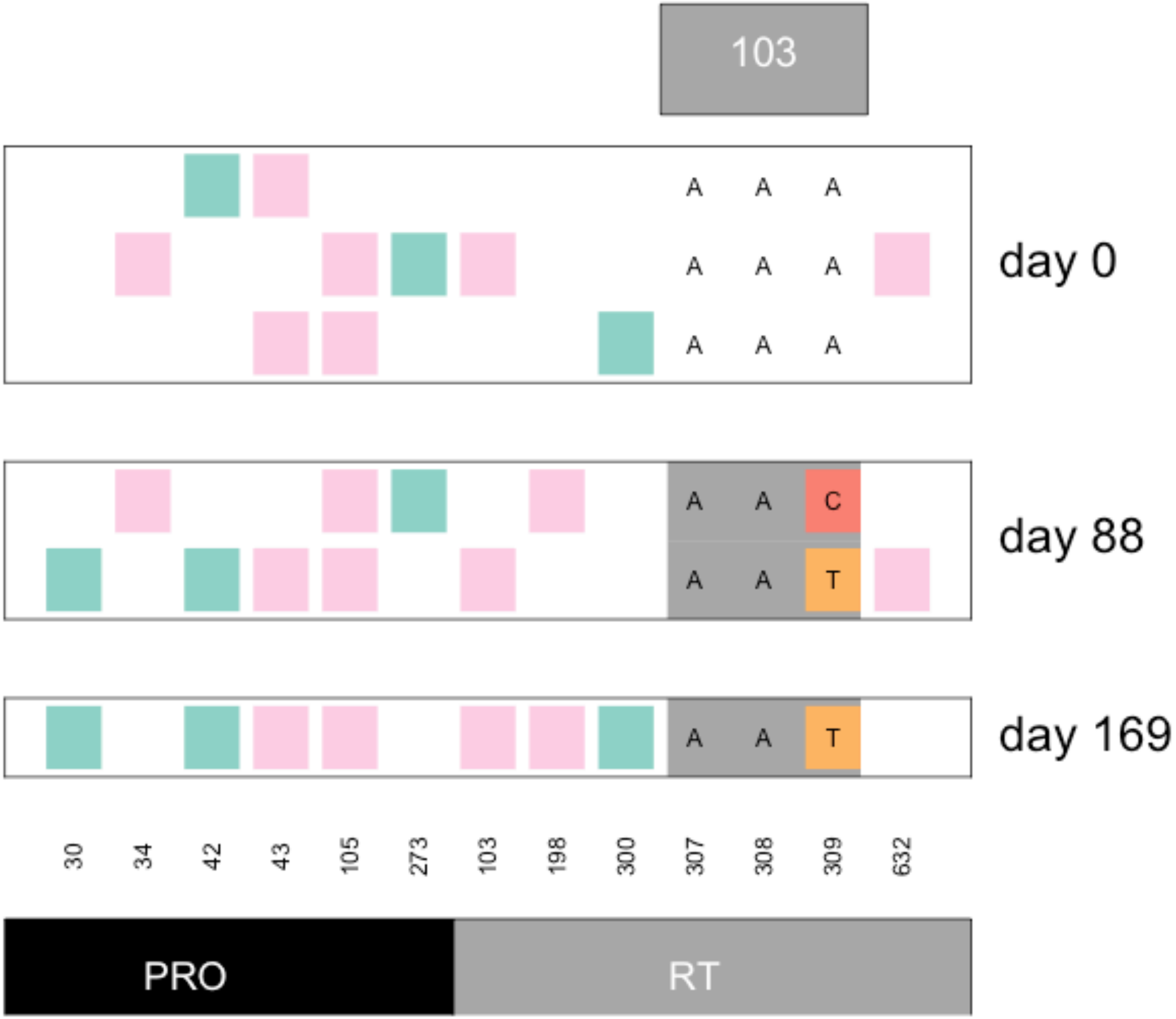


Williams, Pennings, 2018. Data from Bacheler et al 2000

**Viral sequences from patient 013**

( likely treated with indinavir + efavirenz )

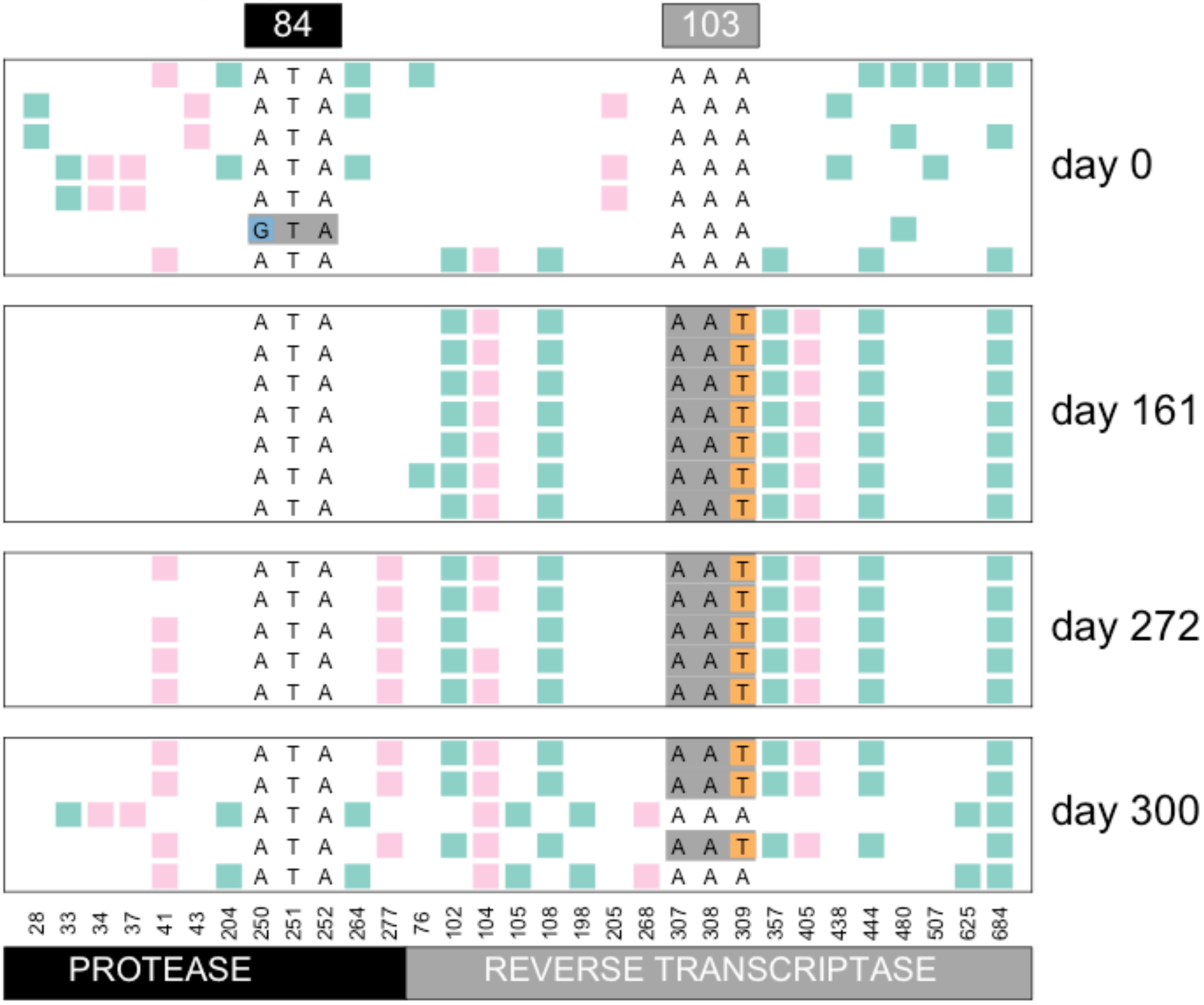


Williams, Pennings, 2018. Data from Bacheler et al 2000

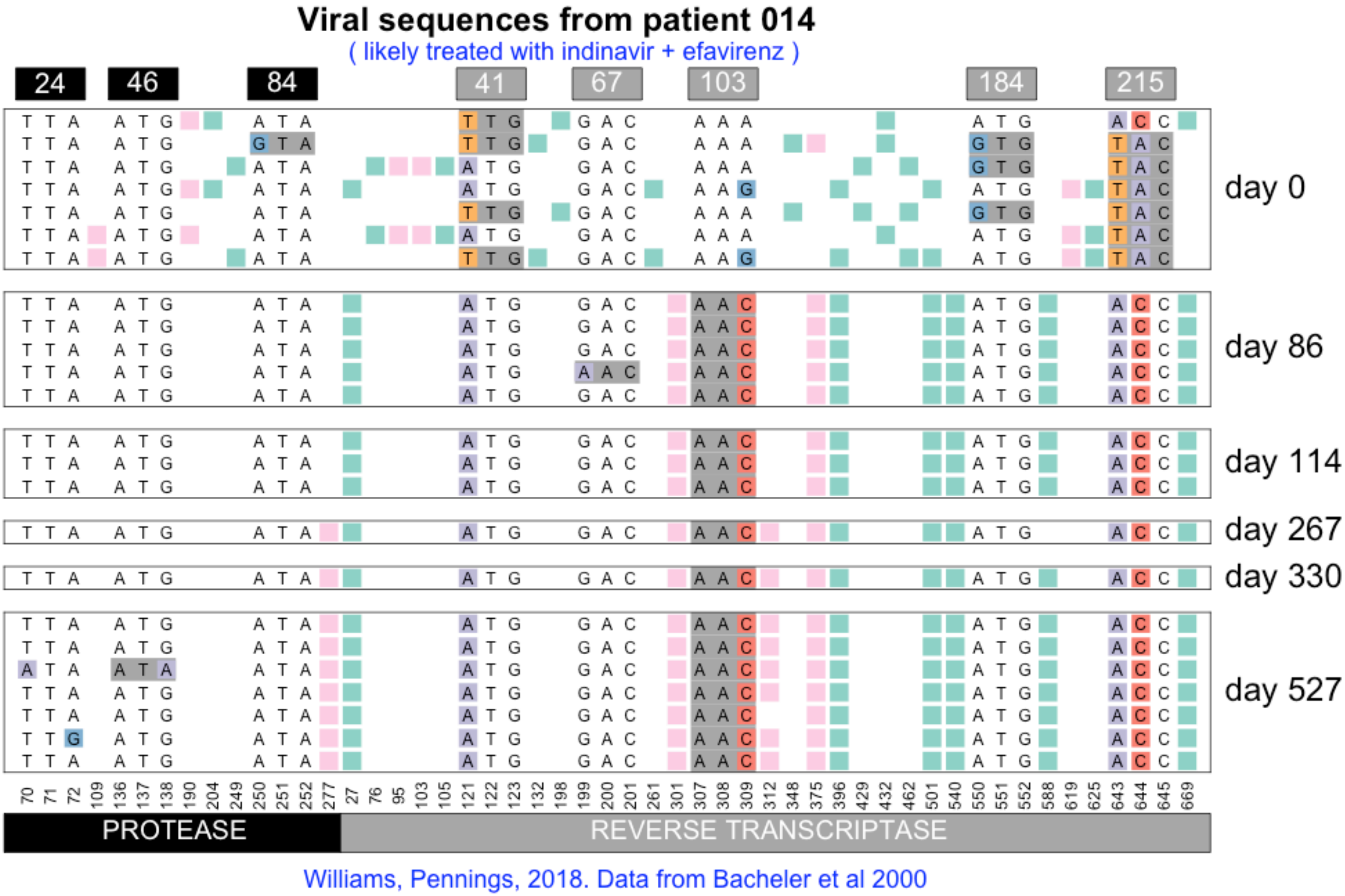

Viral sequences from patient 014
( likely treated with indinavir + efavirenz )
24
46
84
41
67
103
184
215
day 0
day 86
day 114
day 267
day 330
day 527
PROTEASE
REVERSE TRANSCRIPTASE
Williams, Pennings, 2018. Data from Bacheler et al 2000

#### Viral sequences from patient 015

( likely treated with indinavir + efavirenz )

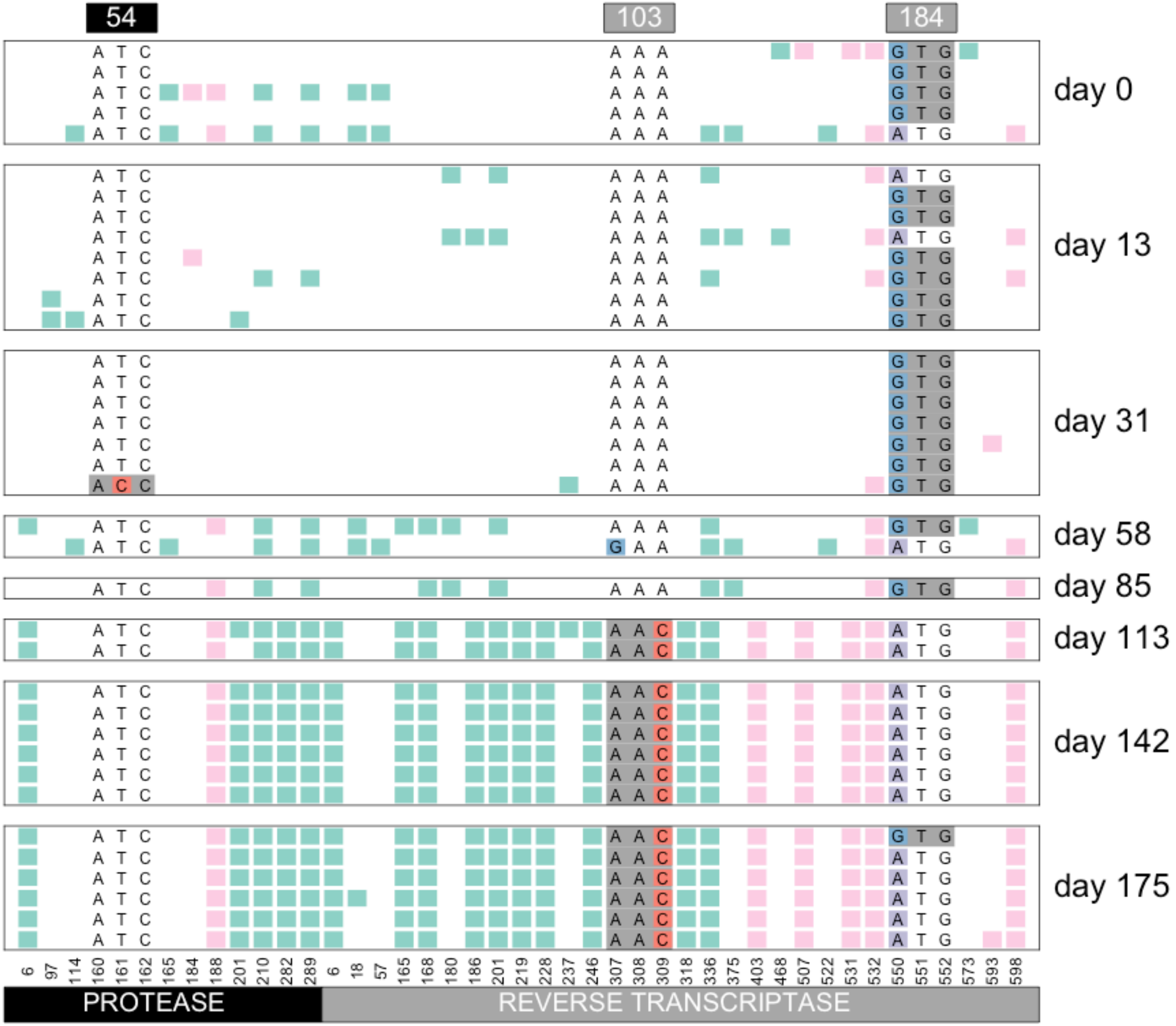


Williams, Pennings, 2018. Data from Bacheler et al 2000

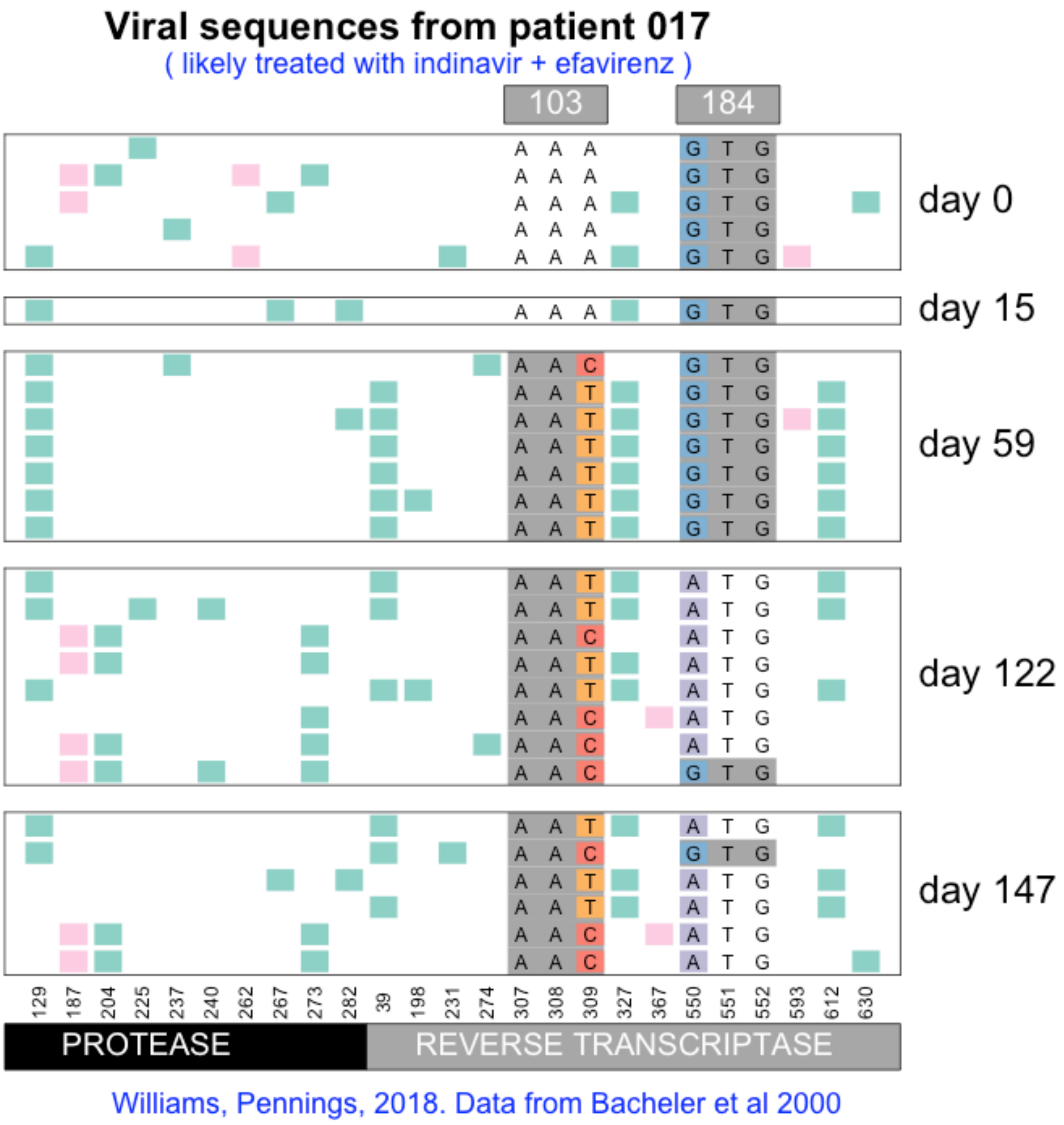

Viral sequences from patient 017
( likely treated with indinavir + efavirenz )
103
184
day 0
day 15
day 59
day 122
day 147
129 187 204 225 237 240 262 267 273 282 39 198 231 274 307 308 309 327 367 550 551 552 593 612 630
PROTEASE
REVERSE TRANSCRIPTASE
Williams, Pennings, 2018. Data from Bacheler et al 2000

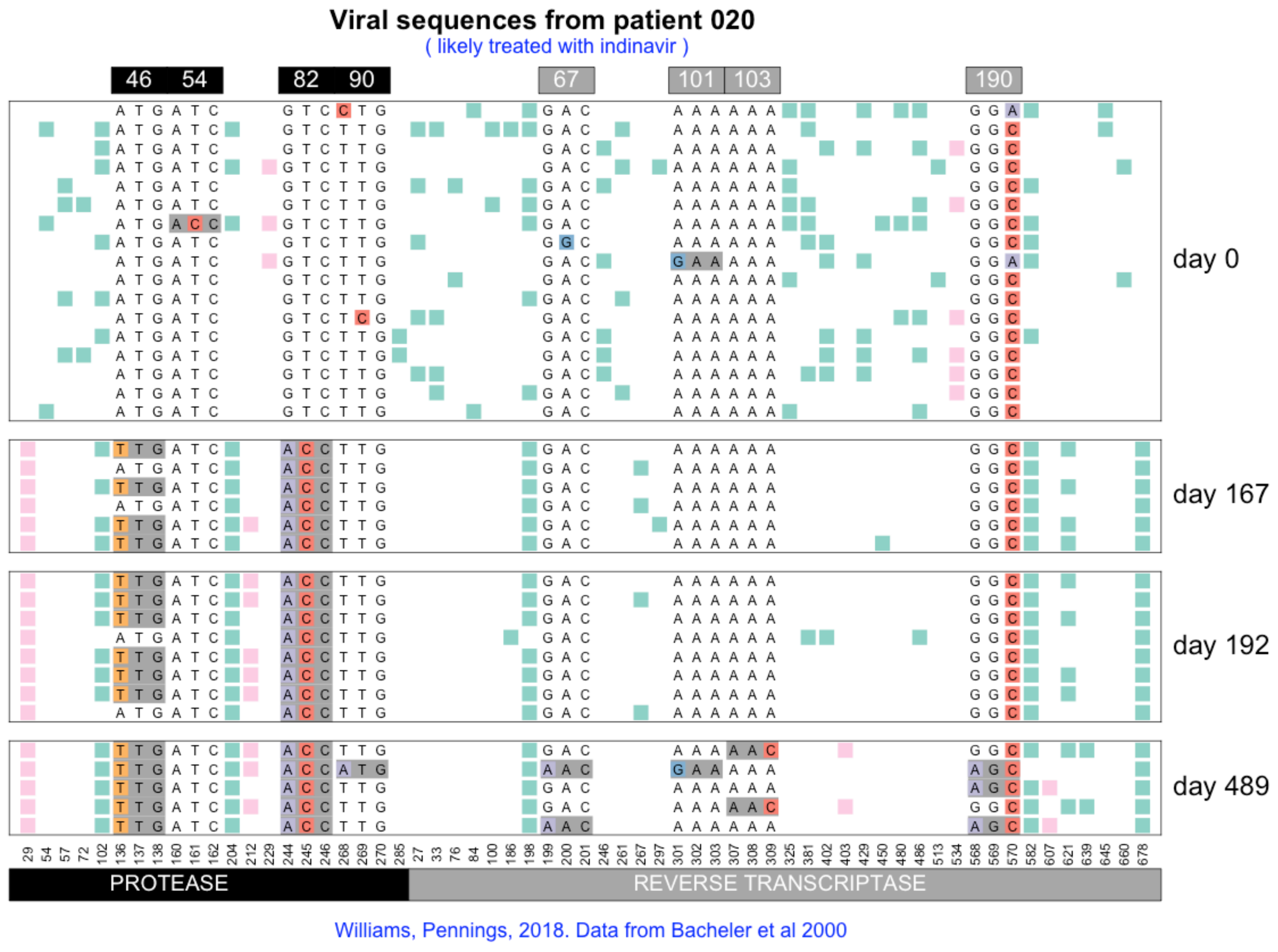

Viral sequences from patient 020
( likely treated with indinavir )
46
54
82
90
67
101
103
190
day 0
day 167
day 192
day 489
PROTEASE
REVERSE TRANSCRIPTASE
Williams, Pennings, 2018. Data from Bacheler et al 2000

#### Viral sequences from patient 021

( likely treated with indinavir + efavirenz )

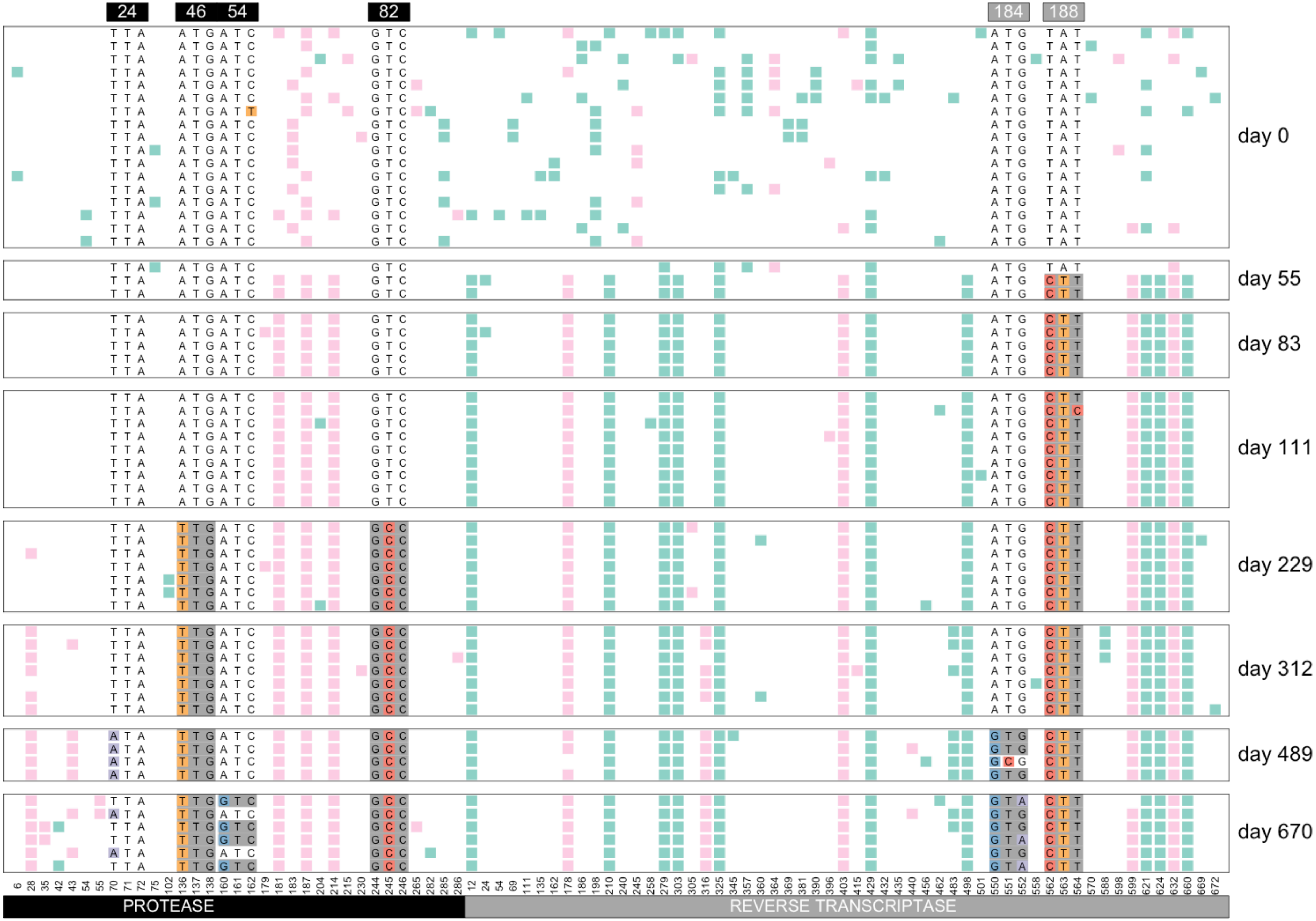


Williams, Pennings, 2018. Data from Bacheler et al 2000

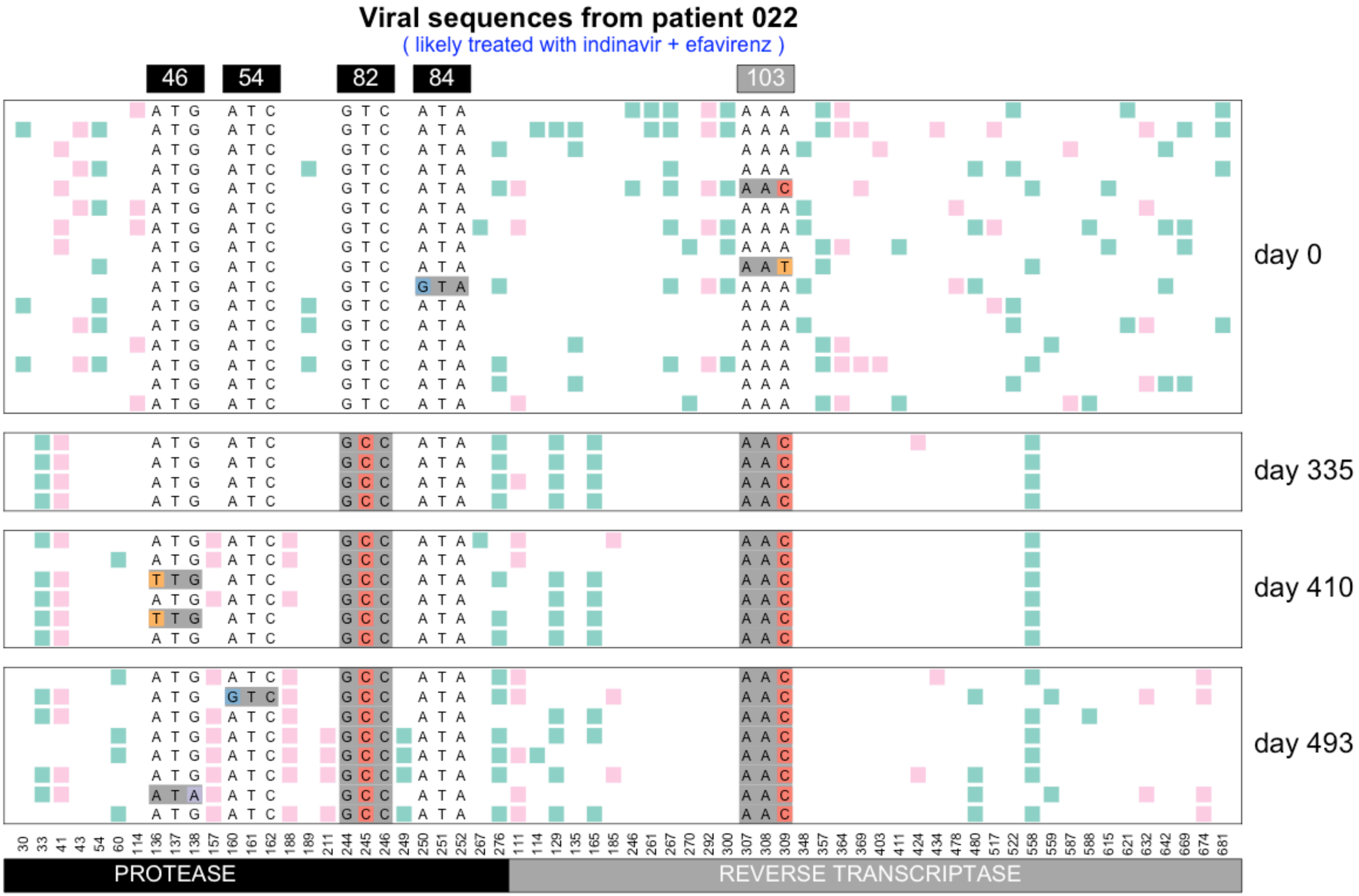


Williams, Pennings, 2018. Data from Bacheler et al 2000

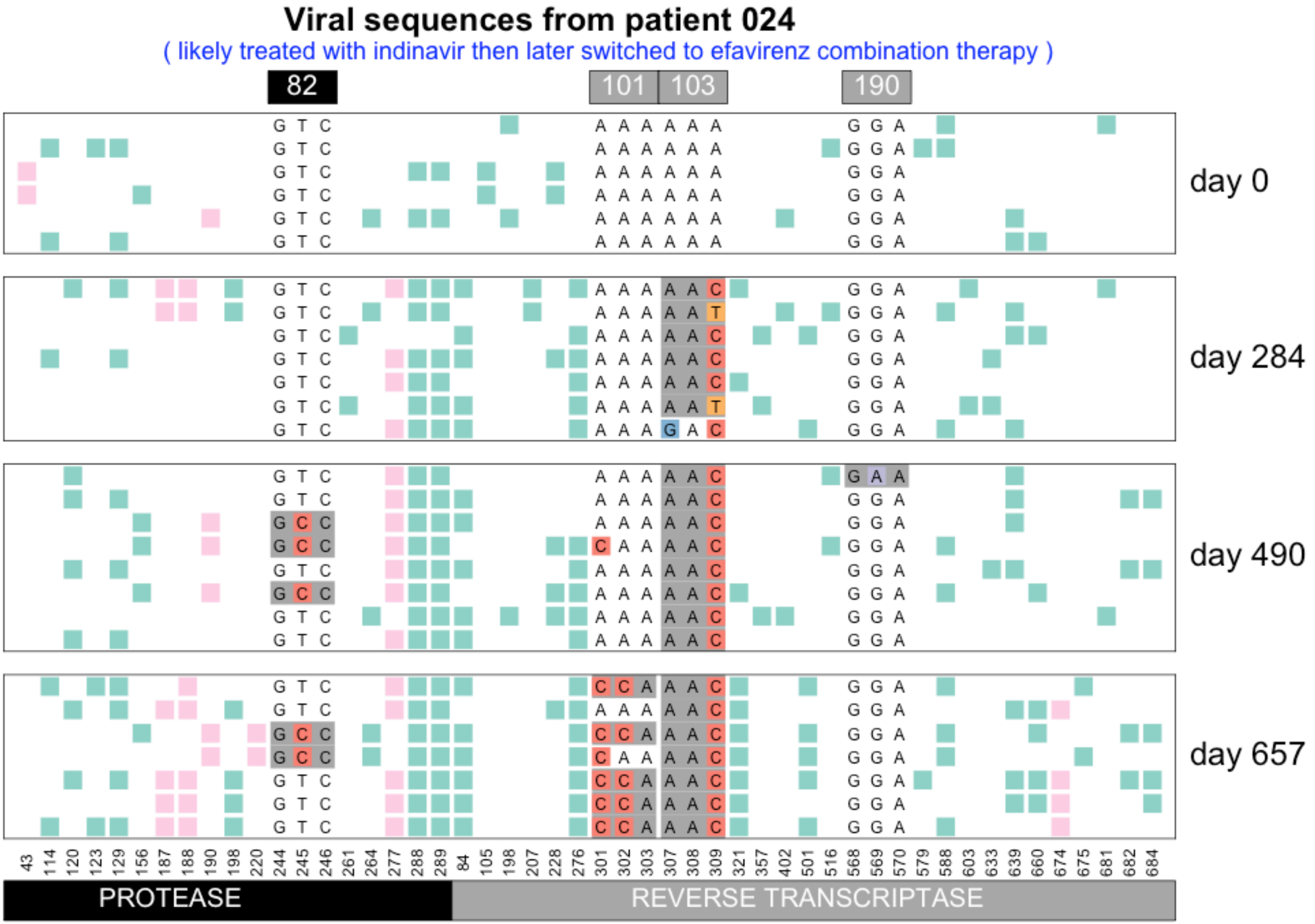


Williams, Pennings, 2018. Data from Bacheler et al 2000

### Viral sequences from patient 025

( likely treated with indinavir + efavirenz )

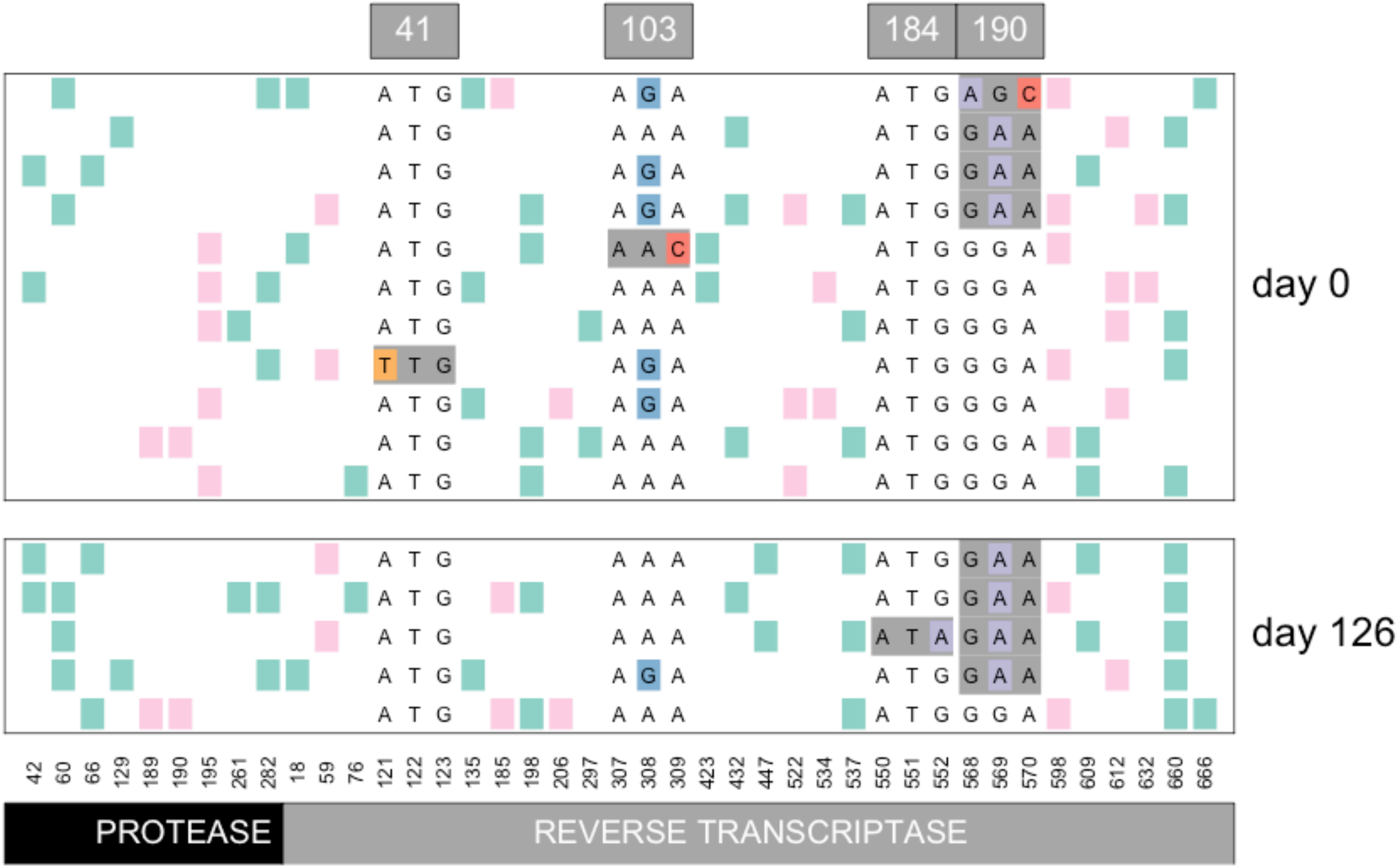


Williams, Pennings, 2018. Data from Bacheler et al 2000

#### Viral sequences from patient 026

( likely treated with indinavir + efavirenz )

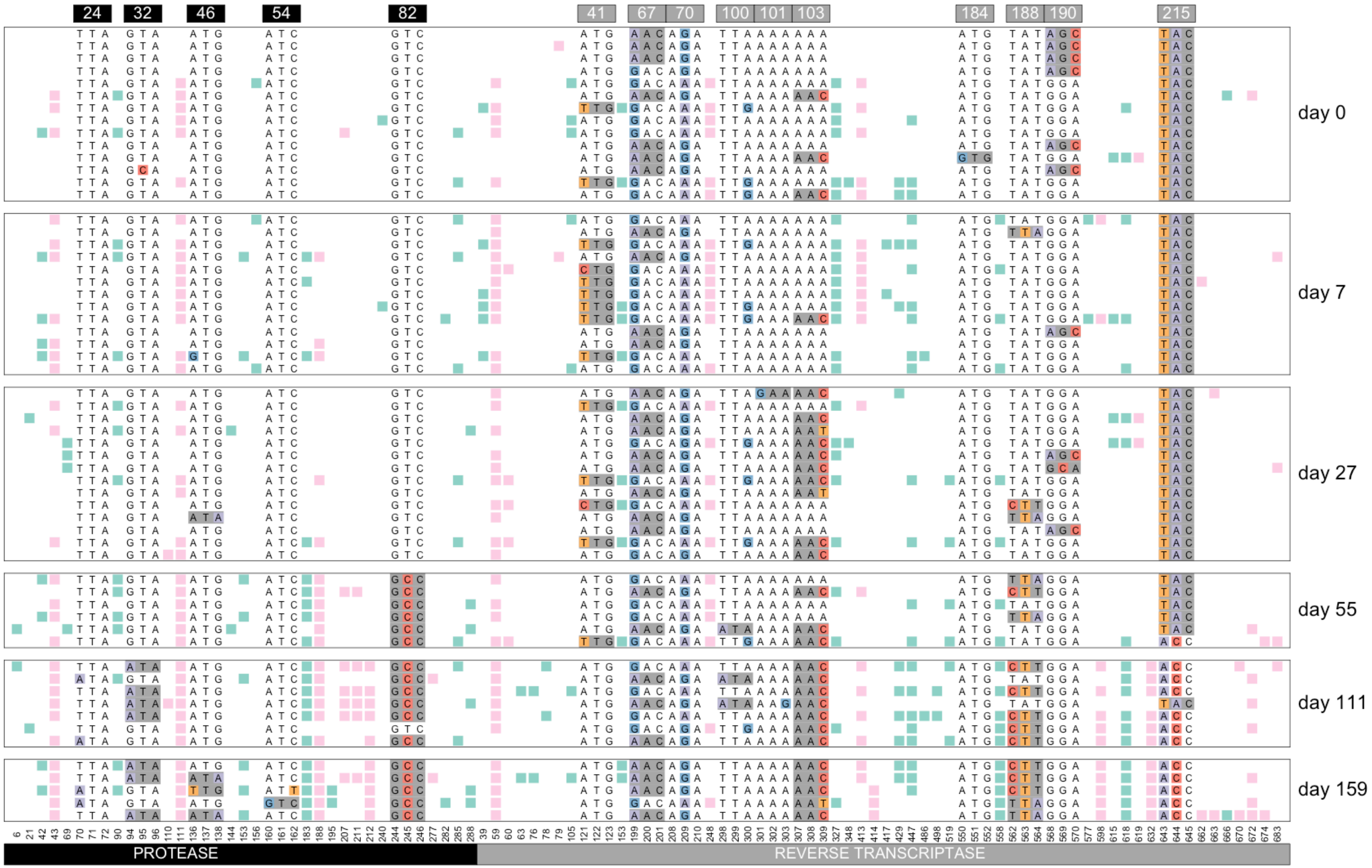


Williams, Pennings, 2018. Data from Bacheler et al 2000

#### Viral sequences from patient 028

( likely treated with indinavir + efavirenz )

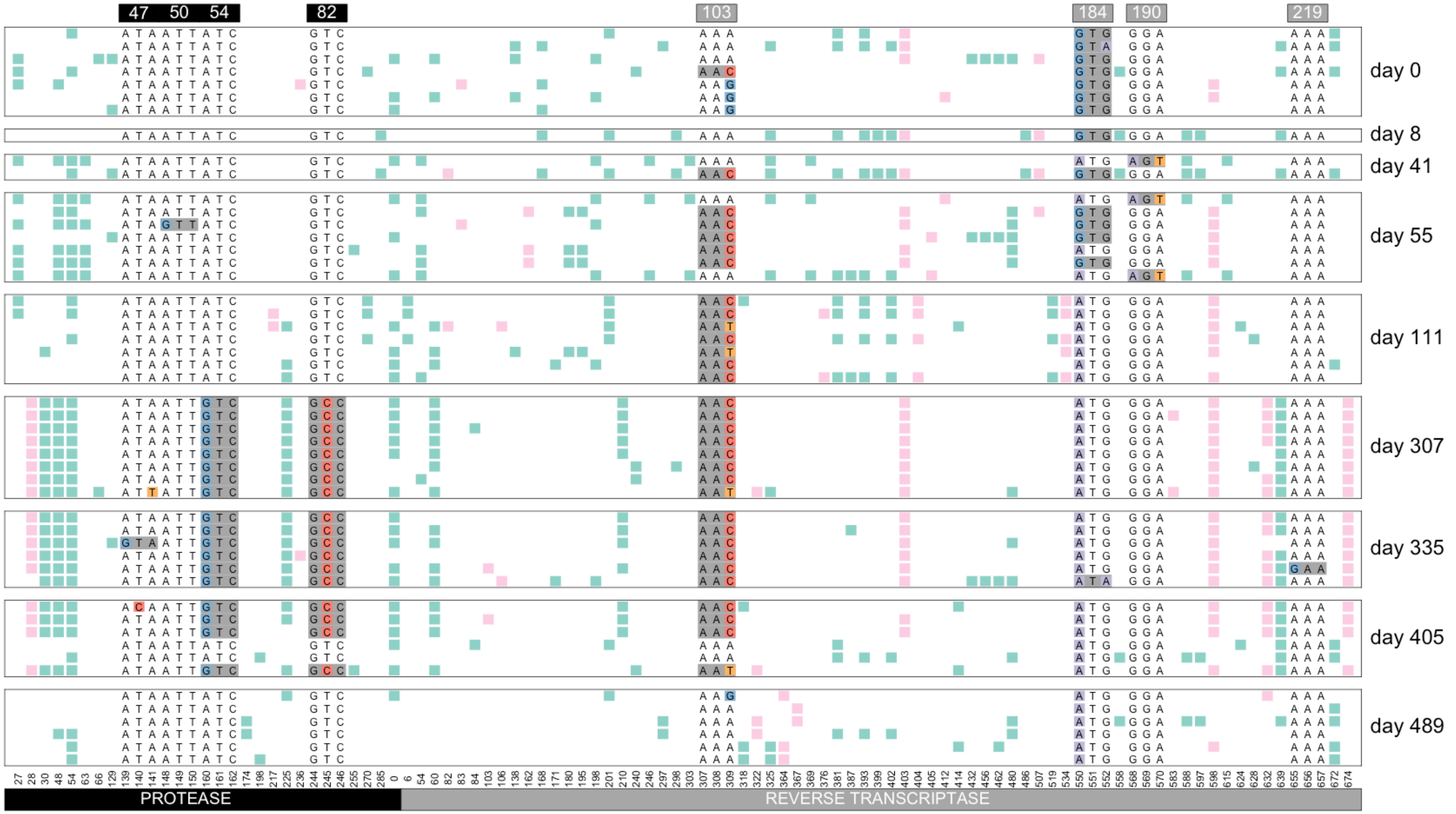


Williams, Pennings, 2018. Data from Bacheler et al 2000

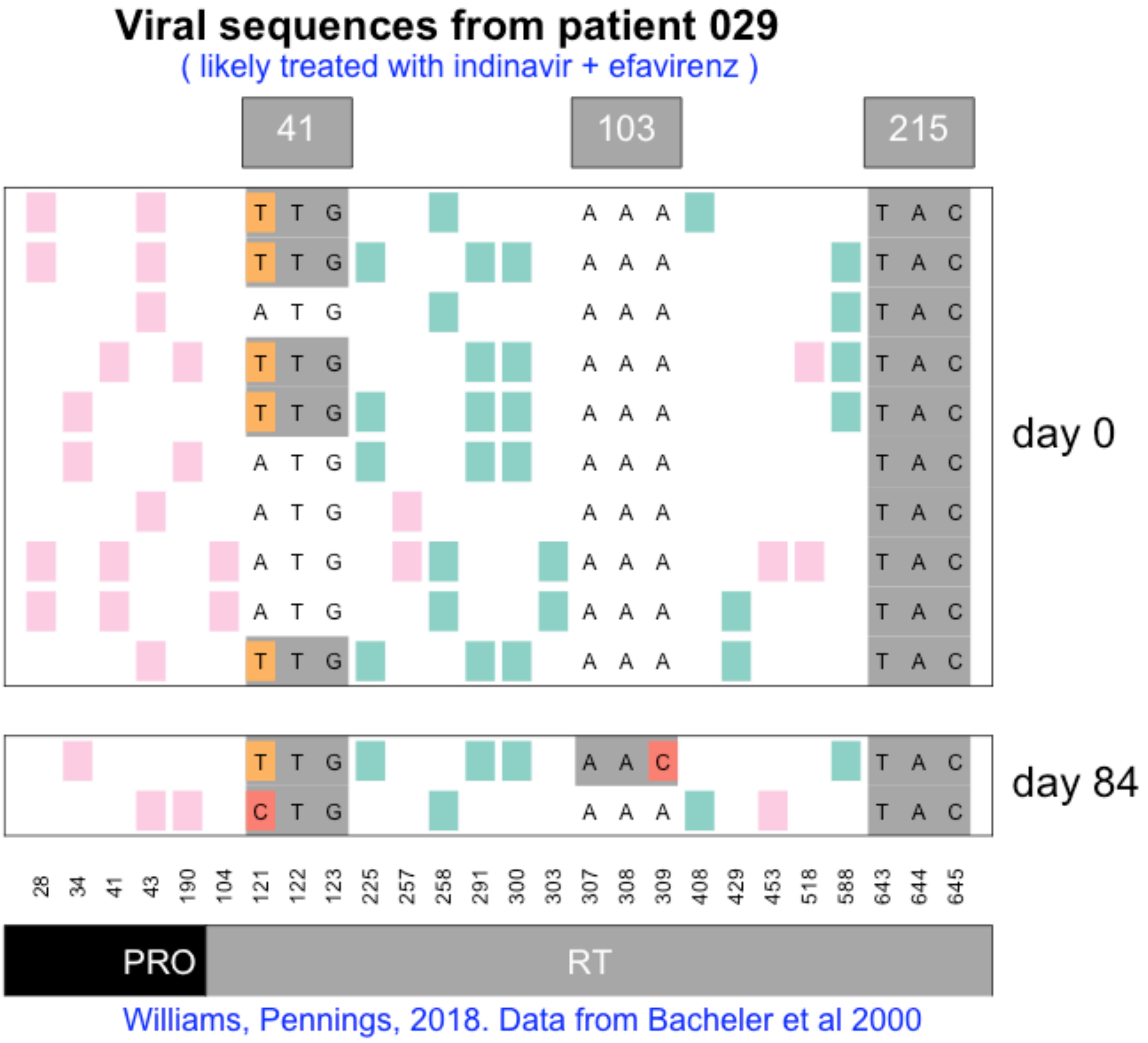

Viral sequences from patient 029
( likely treated with indinavir + efavirenz )
41
103
215
day 0
day 84
28 34 41 43 190 104 121 122 123 225 257 258 291 300 303 307 308 309 408 429 453 518 588 643 644 645
PRO
RT
Williams, Pennings, 2018. Data from Bacheler et al 2000

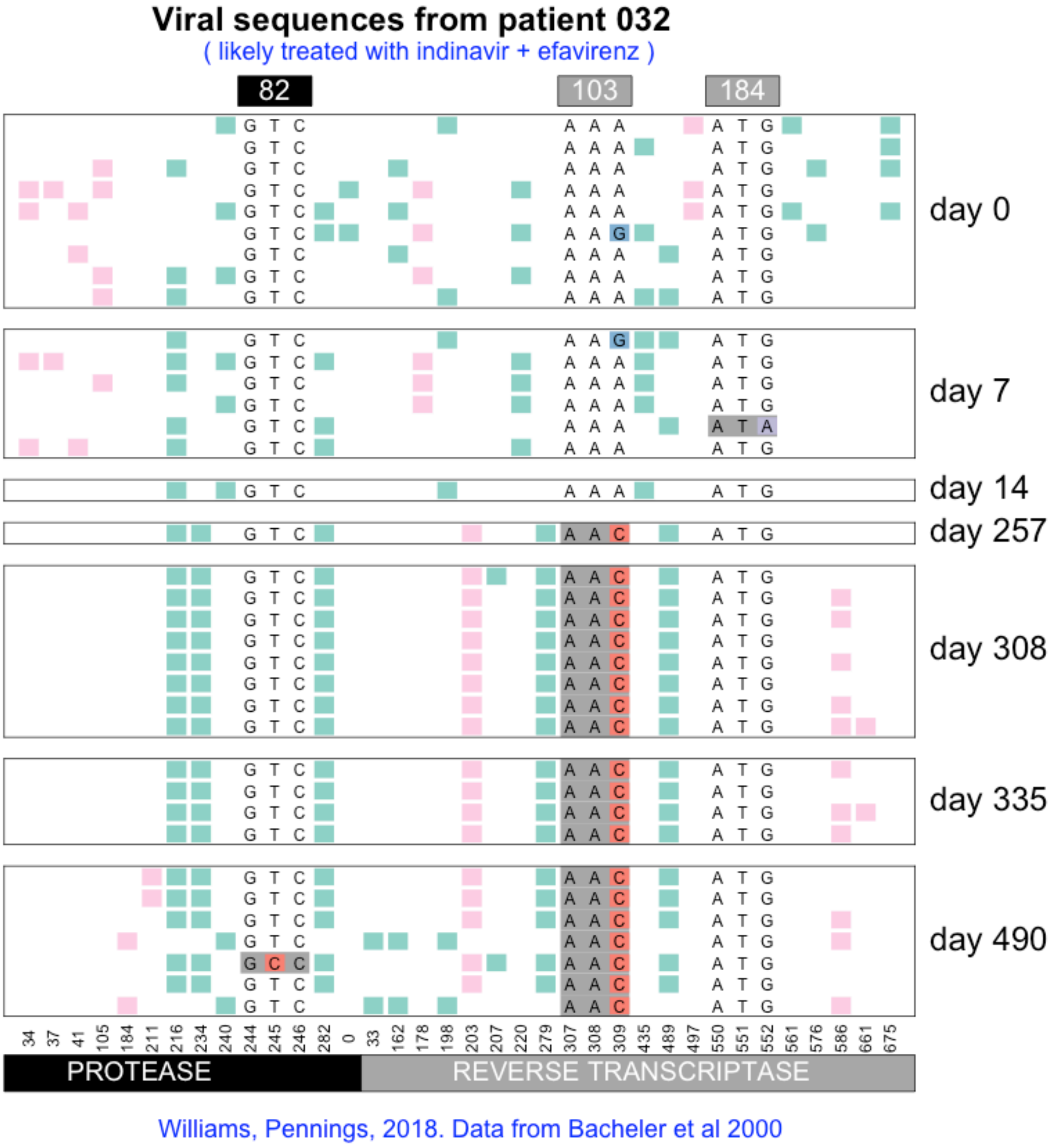
Viral sequences from patient 032
( likely treated with indinavir + efavirenz )
82
103
184
day 0
day 7
day 14
day 257
day 308
day 335
day 490
34 37 41 105 184 211 216 234 240 244 245 246 282 0 33 162 178 198 203 207 220 279 307 308 309 435 489 497 550 551 552 561 576 586 661 675
PROTEASE
REVERSE TRANSCRIPTASE
Williams, Pennings, 2018. Data from Bacheler et al 2000

### Viral sequences from patient 037

( likely treated with indinavir then later switched to efavirenz combination therapy )

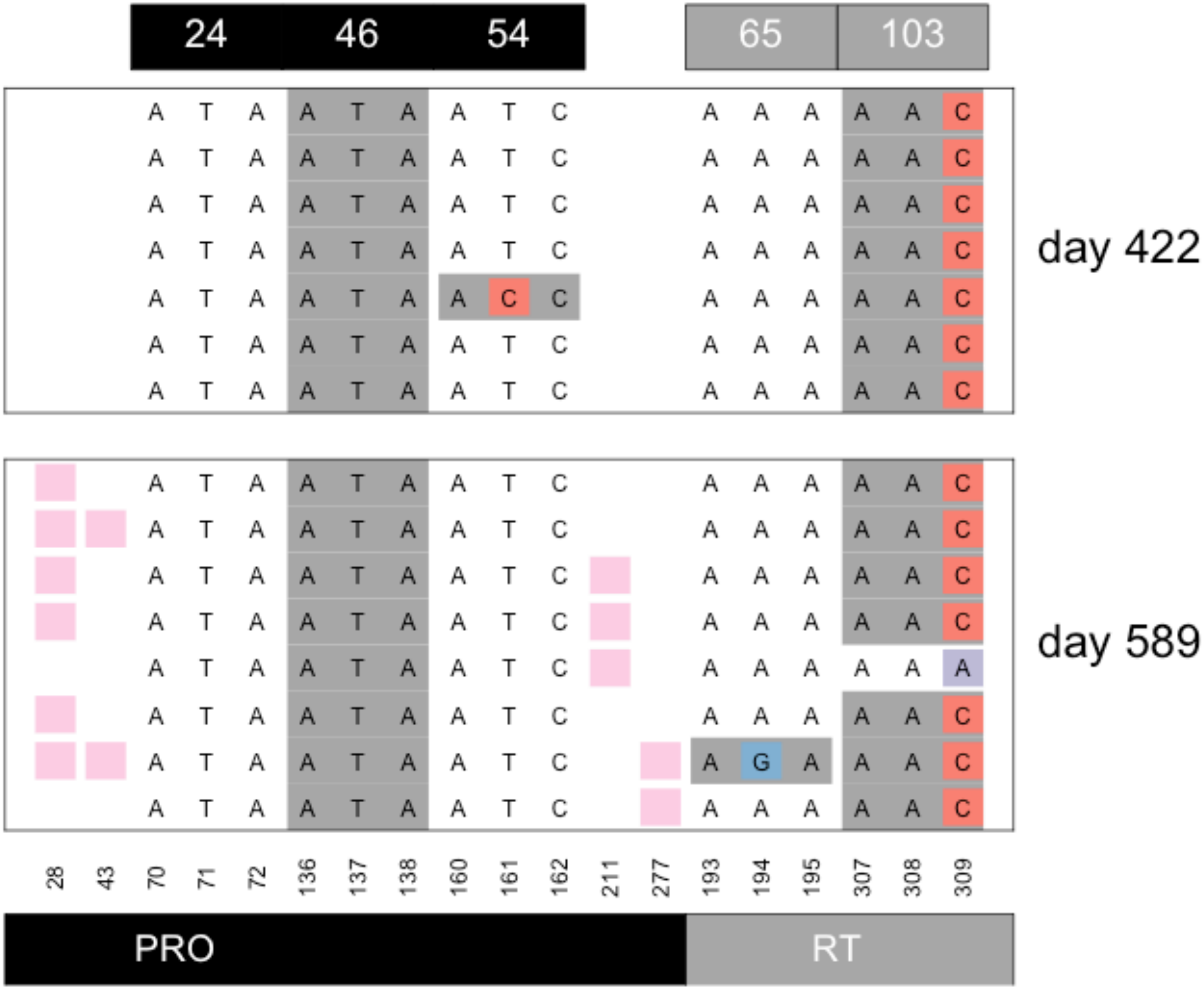


Williams, Pennings, 2018. Data from Bacheler et al 2000

#### Viral sequences from patient 039

( likely treated with indinavir then later switched to efavirenz combination therapy )

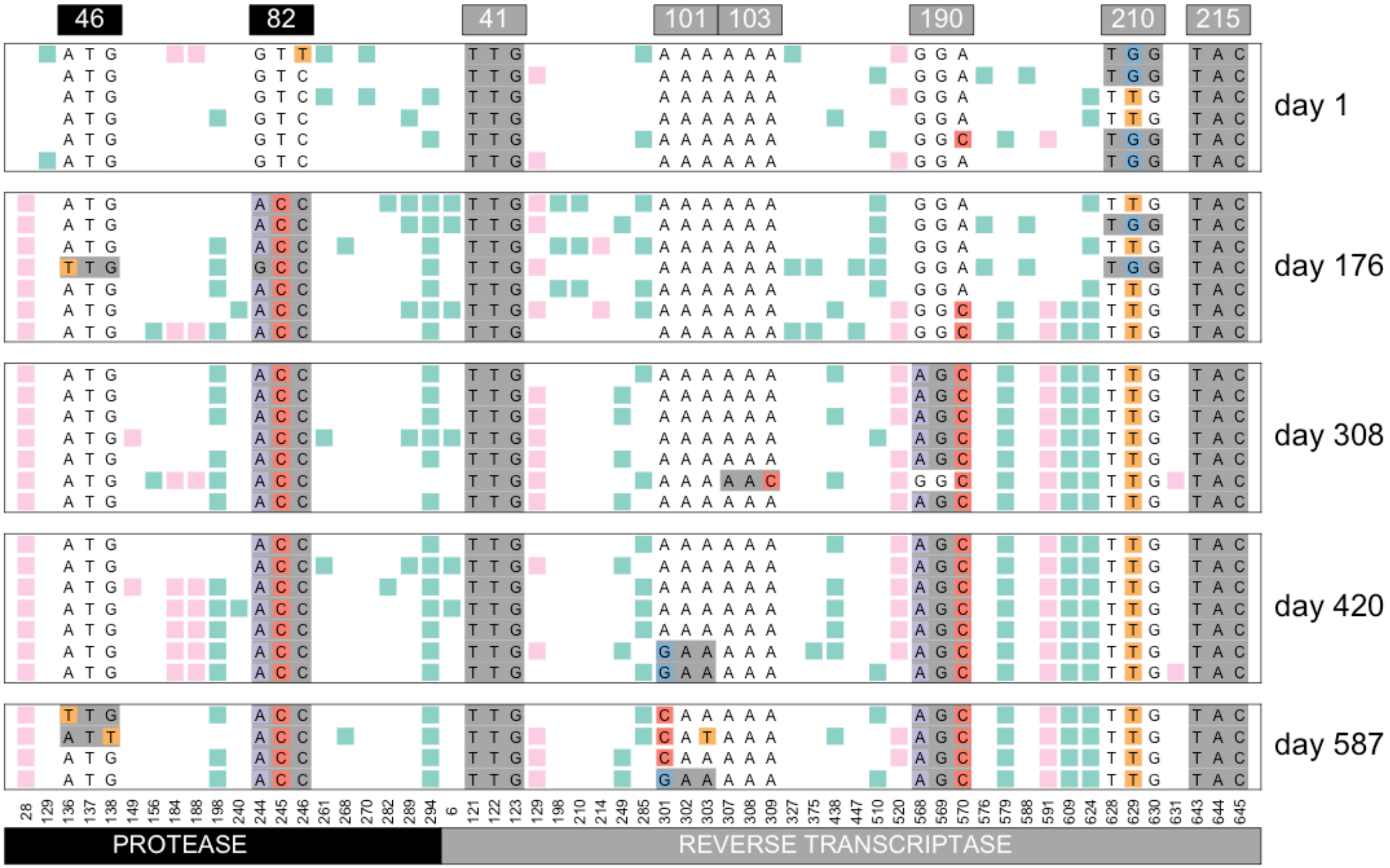


Williams, Pennings, 2018. Data from Bacheler et al 2000

### Viral sequences from patient 043

( likely treated with indinavir then later switched to efavirenz combination therapy )

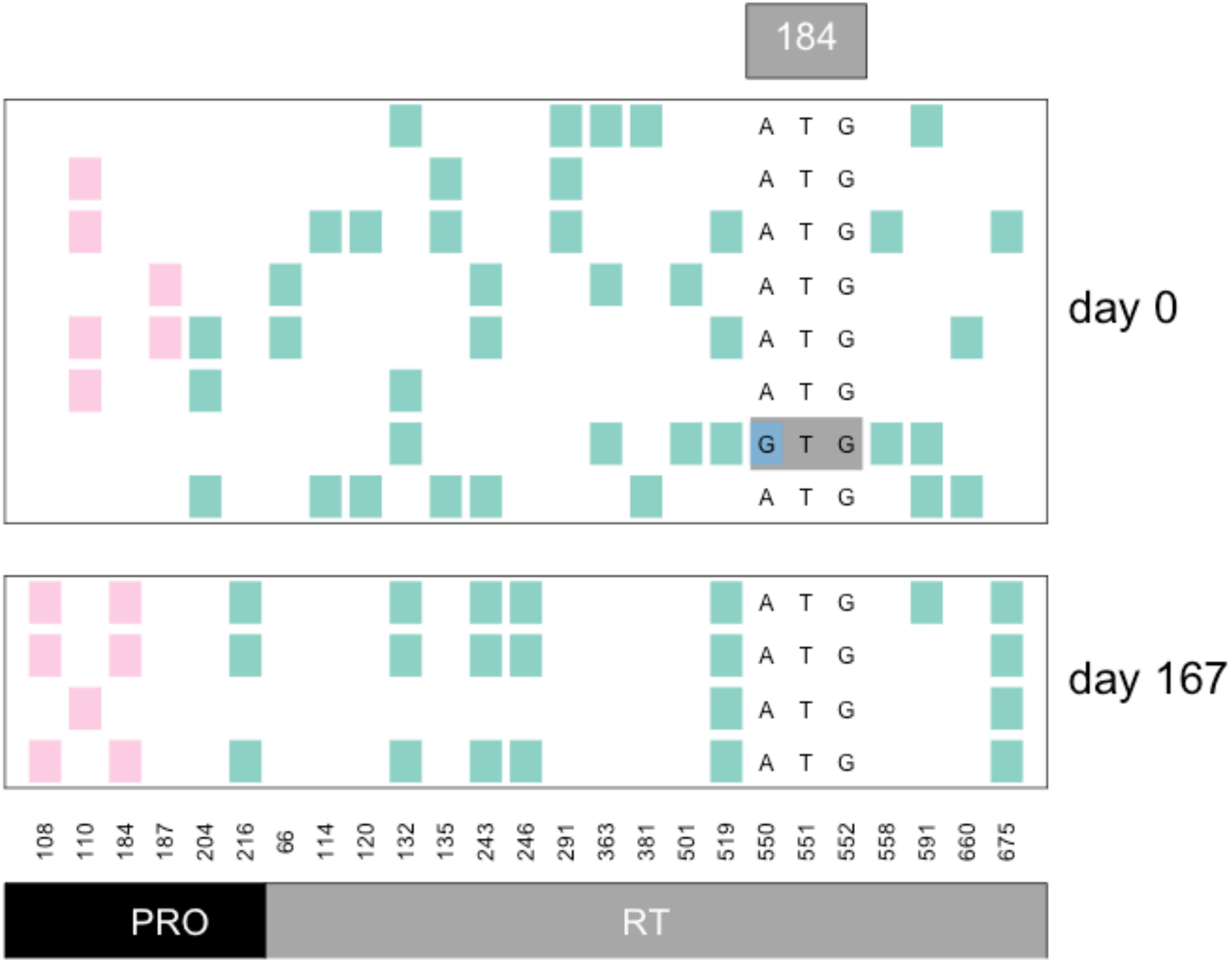


Williams, Pennings, 2018. Data from Bacheler et al 2000

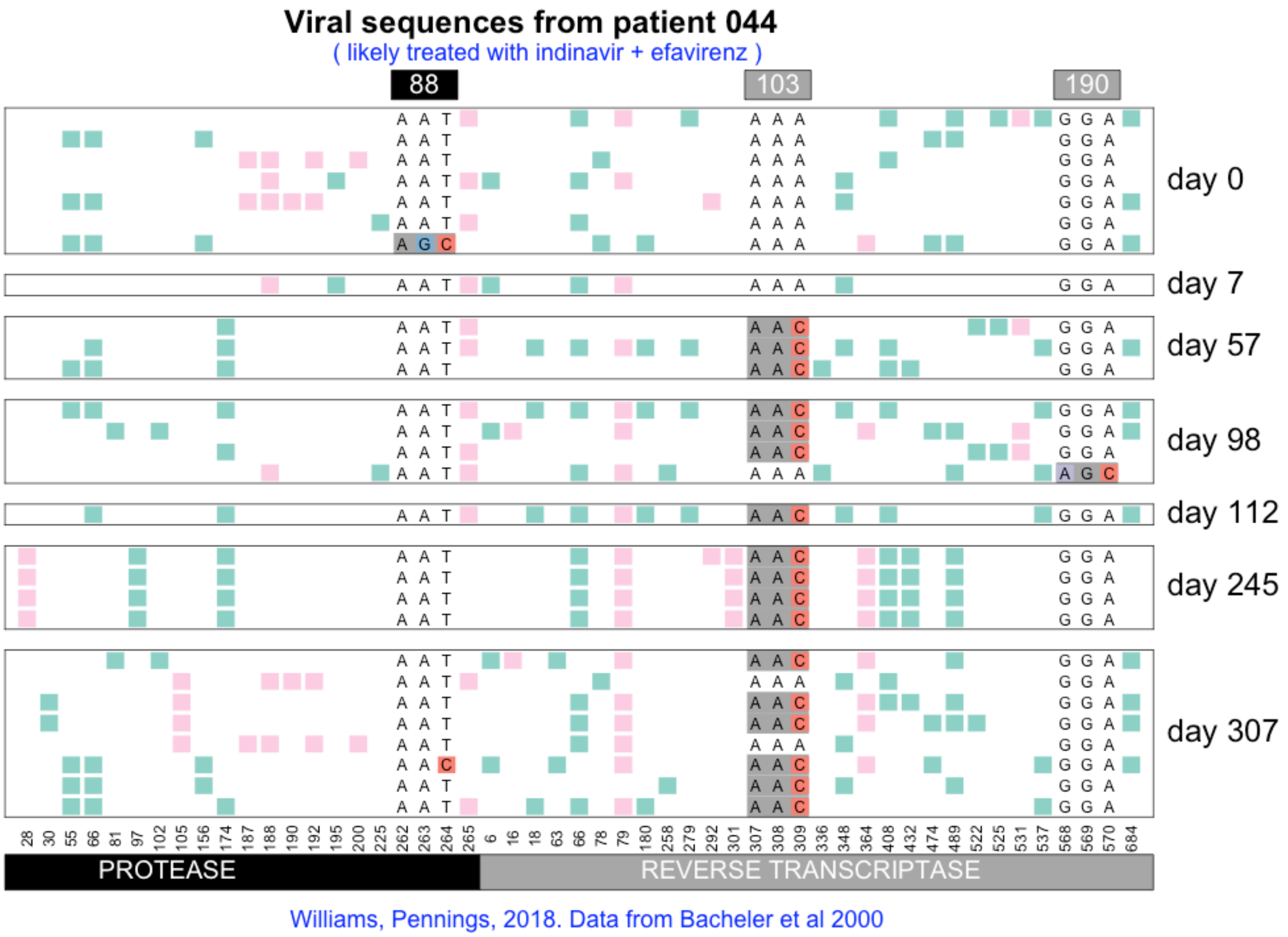
Viral sequences from patient 044
( likely treated with indinavir + efavirenz )
88
103
190
day 0
day 7
day 57
day 98
day 112
day 245
day 307
28 30 55 66 81 97 102 105 156 174 187 188 190 192 195 200 225 262 263 264 265 6 16 18 63 66 78 79 180 258 279 292 301 307 308 309 336 348 364 408 432 474 489 522 525 531 537 568 569 570 684
PROTEASE
REVERSE TRANSCRIPTASE
Williams, Pennings, 2018. Data from Bacheler et al 2000

### Viral sequences from patient 045

( likely treated with indinavir + efavirenz )

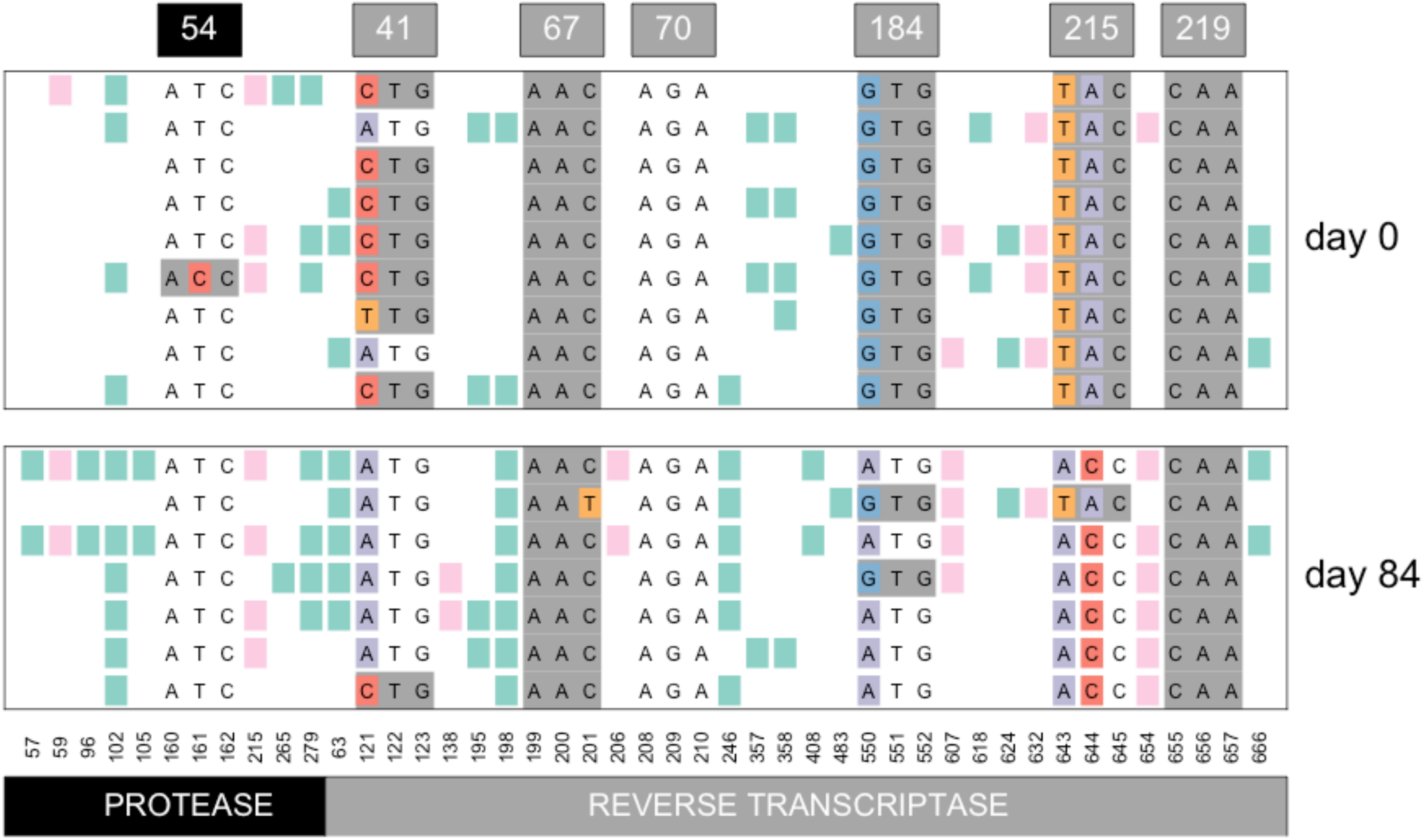


Williams, Pennings, 2018. Data from Bacheler et al 2000

### Viral sequences from patient 047

( likely treated with indinavir then later switched to efavirenz combination therapy )

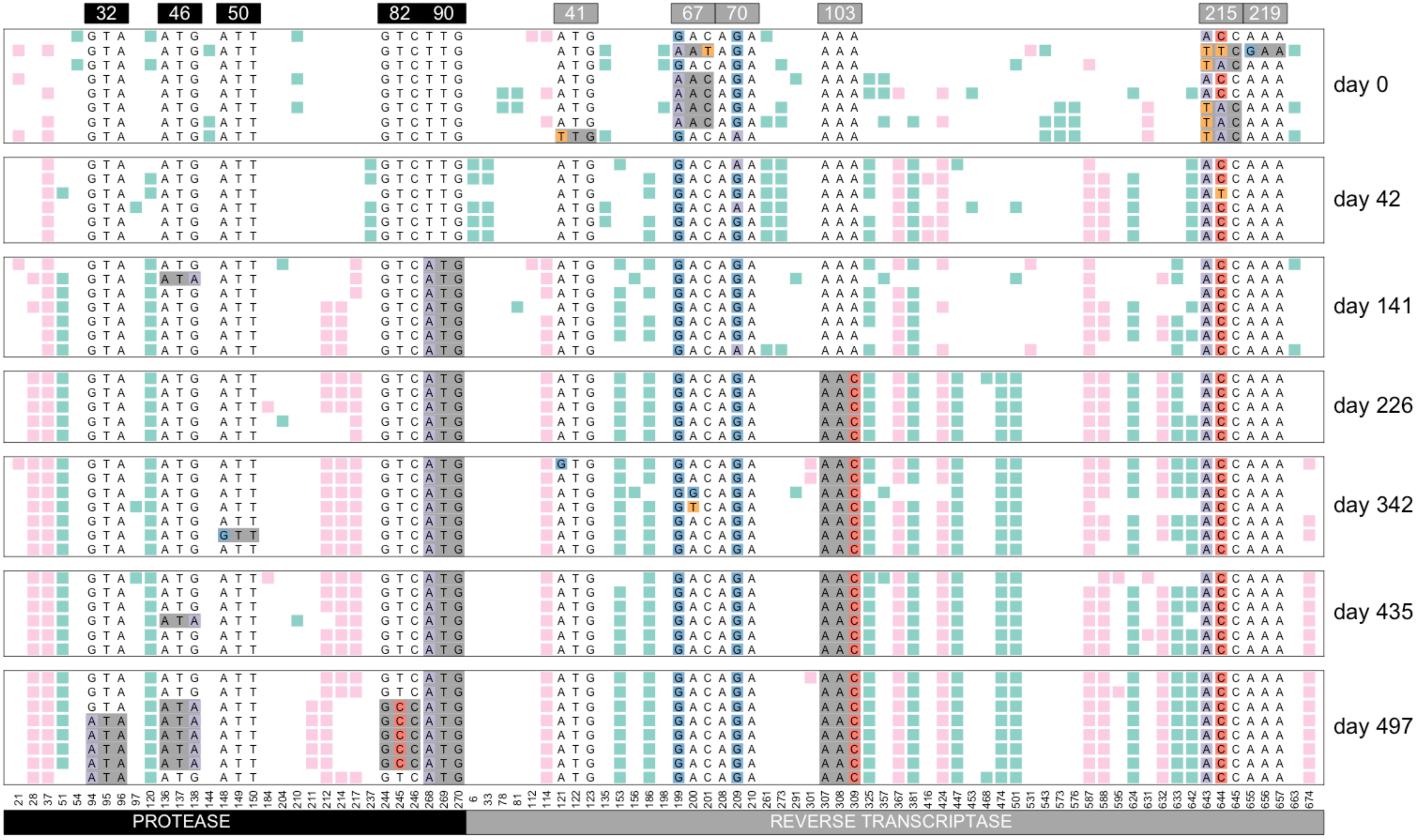


Williams, Pennings, 2018. Data from Bacheler et al 2000

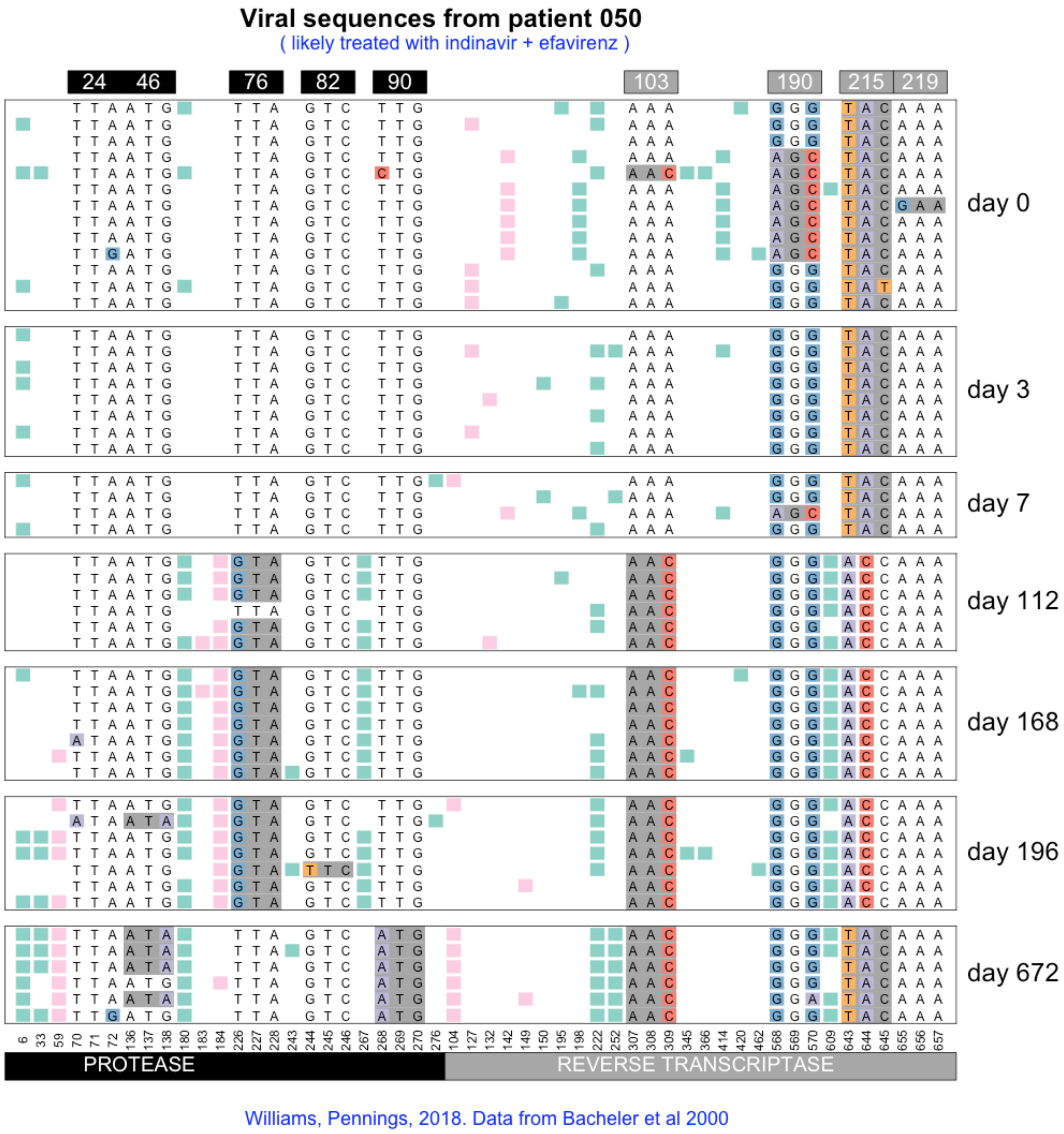
Viral sequences from patient 050
( likely treated with indinavir + efavirenz )
24 46
76
82
90
103
190
215 219
day 0
day 3
day 7
day 112
day 168
day 196
day 672
6 33 59 70 71 72 136 137 138 180 183 184 226 227 228 243 244 245 246 267 268 269 270 276 104 127 132 142 149 150 195 198 222 252 307 308 309 345 366 414 420 462 568 569 570 609 643 644 645 655 656 657
PROTEASE
REVERSE TRANSCRIPTASE
Williams, Pennings, 2018. Data from Bacheler et al 2000

#### Viral sequences from patient 055

( likely treated with indinavir + efavirenz )

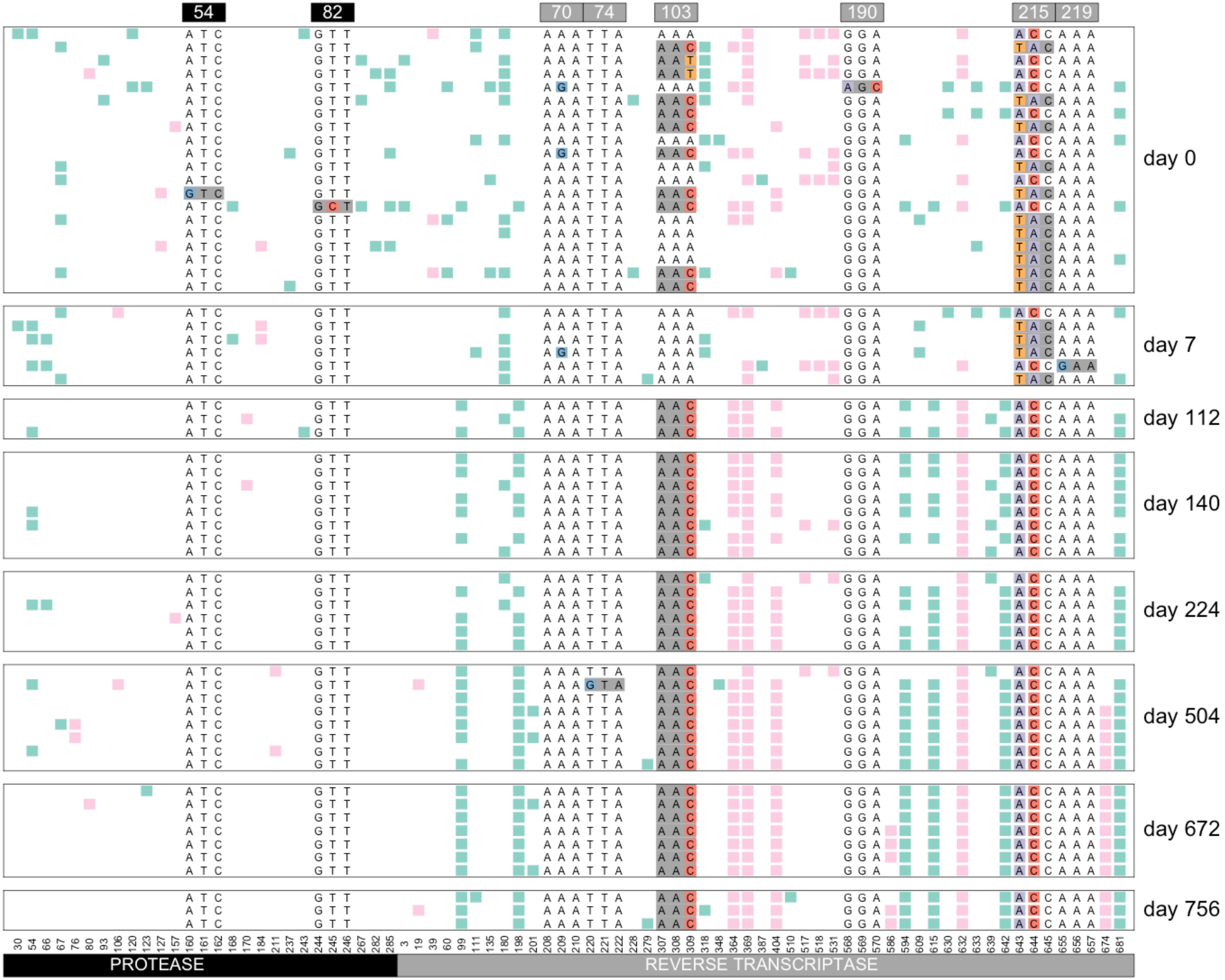


Williams, Pennings, 2018. Data from Bacheler et al 2000

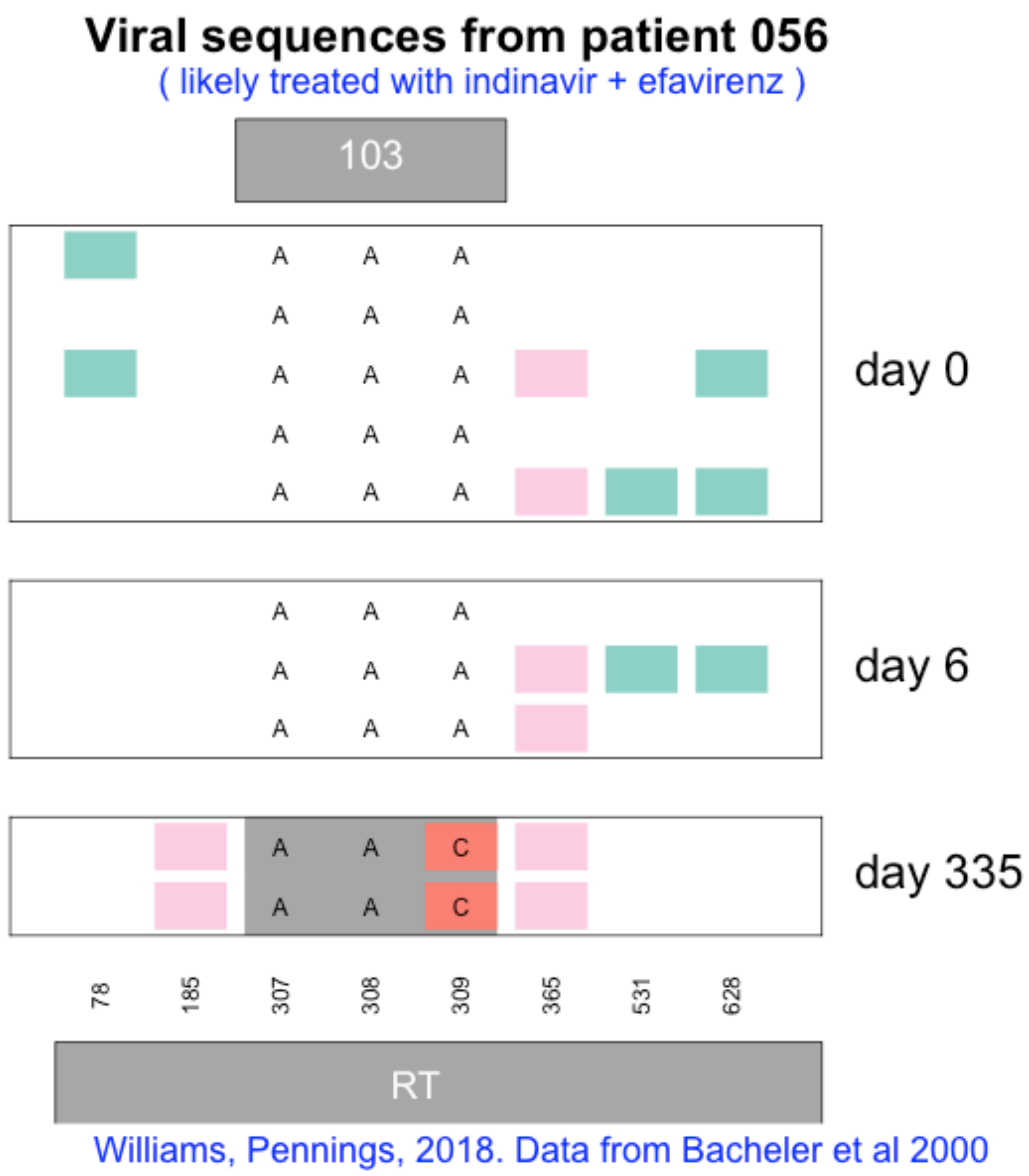
Viral sequences from patient 056
( likely treated with indinavir + efavirenz )
103
A A A
A A A
A A A
A A A
A A A
day 0
A A A
A A A
A A A
day 6
A A C
A A C
day 335
78
185
307
308
309
365
531
628
RT
Williams, Pennings, 2018. Data from Bacheler et al 2000

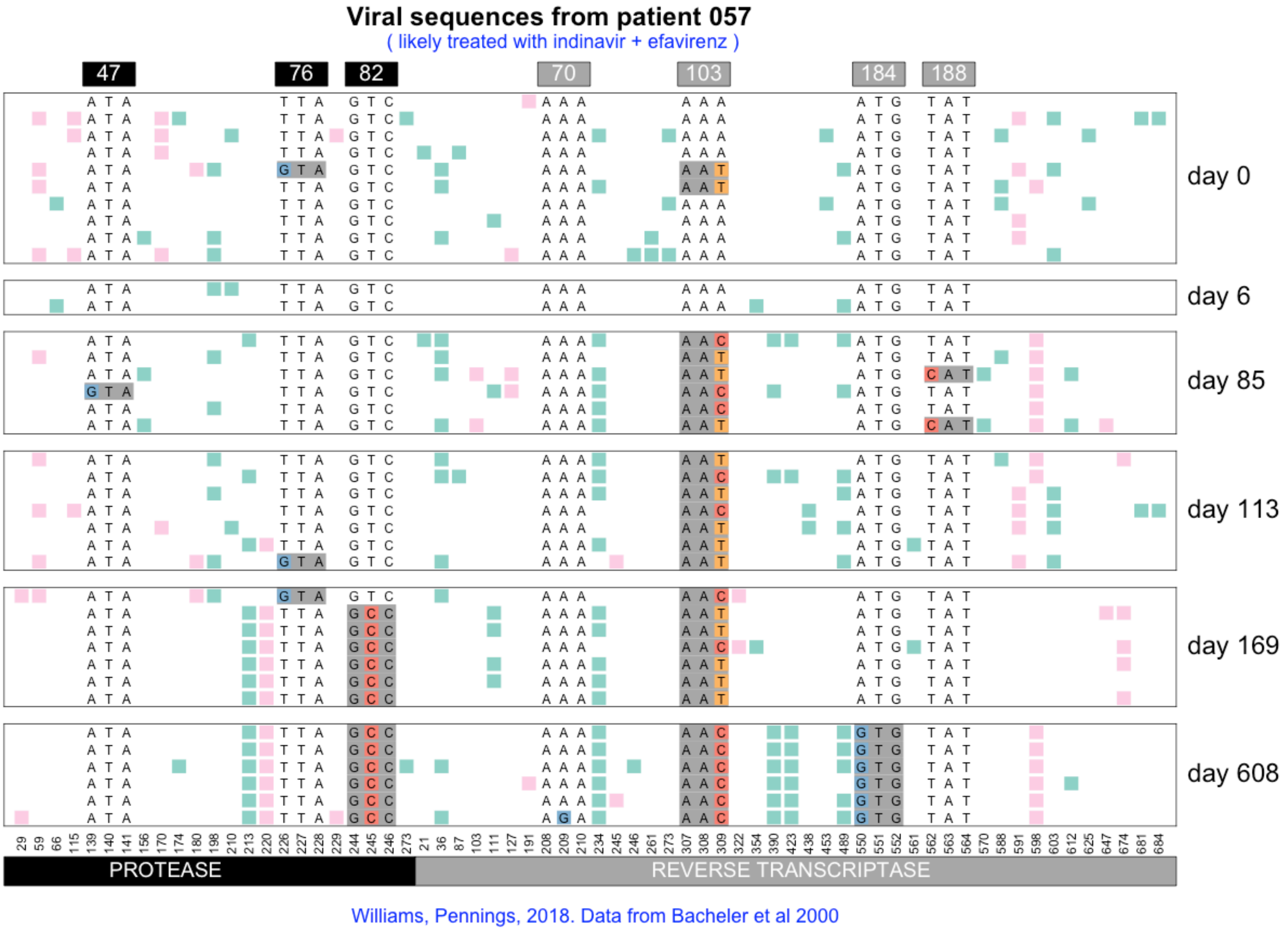
Viral sequences from patient 057
( likely treated with indinavir + efavirenz )
47
76
82
70
103
184
188
day 0
day 6
day 85
day 113
day 169
day 608
29 59 66 115 139 140 141 156 170 174 180 198 210 213 220 226 227 228 229 244 245 246 273 21 36 87 103 111 127 191 208 209 210 234 245 246 261 273 307 308 309 322 354 390 423 438 453 489 550 551 552 561 562 563 564 570 588 591 598 603 612 625 647 674 681 684
PROTEASE
REVERSE TRANSCRIPTASE
Williams, Pennings, 2018. Data from Bacheler et al 2000

### Viral sequences from patient 058

( likely treated with indinavir + efavirenz )

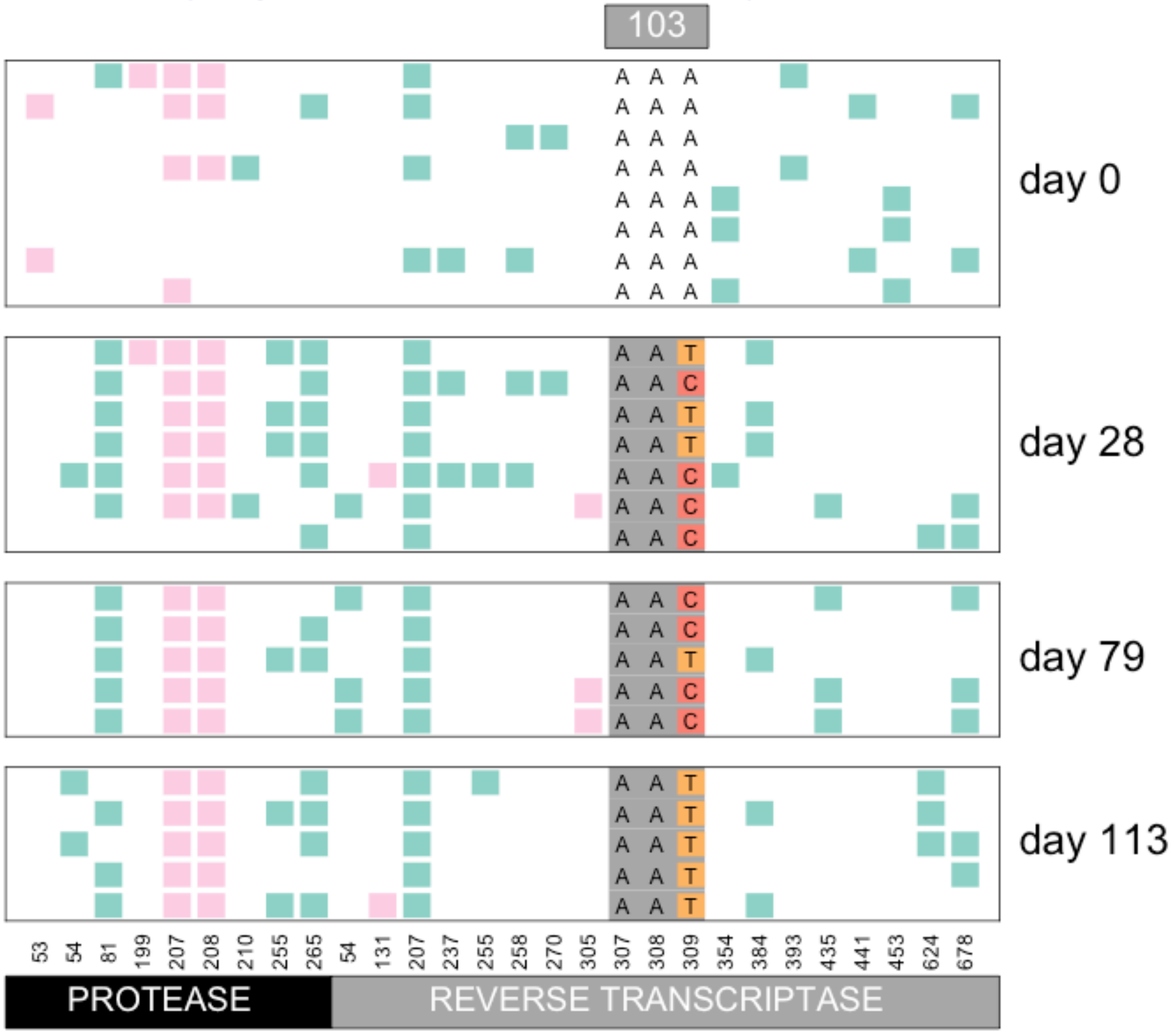


Williams, Pennings, 2018. Data from Bacheler et al 2000

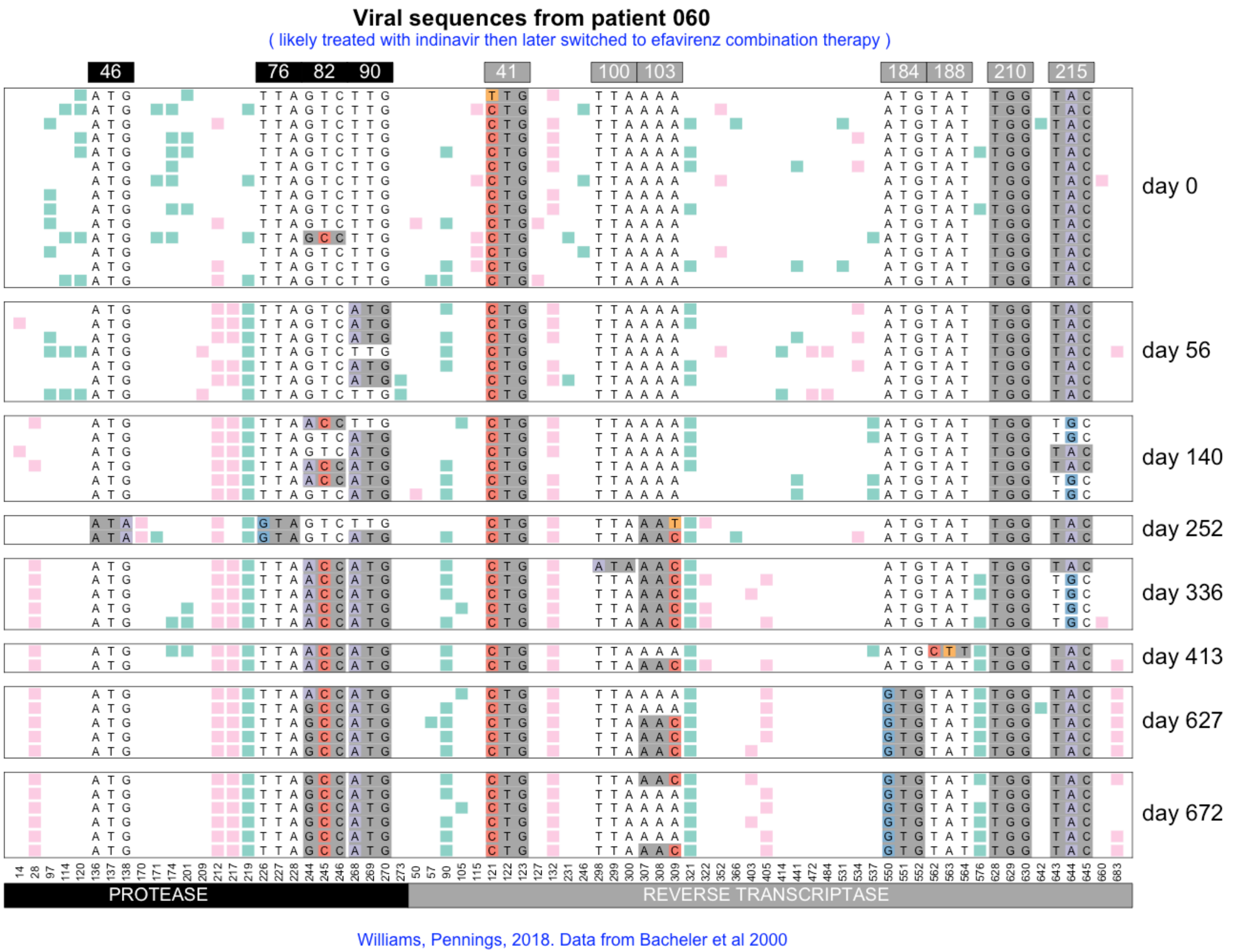
Viral sequences from patient 060
( likely treated with indinavir then later switched to efavirenz combination therapy )
46
76
82
90
41
100
103
184
188
210
215
day 0
day 56
day 140
day 252
day 336
day 413
day 627
day 672
PROTEASE
REVERSE TRANSCRIPTASE
Williams, Pennings, 2018. Data from Bacheler et al 2000

### Viral sequences from patient 062

( likely treated with indinavir then later switched to efavirenz combination therapy )

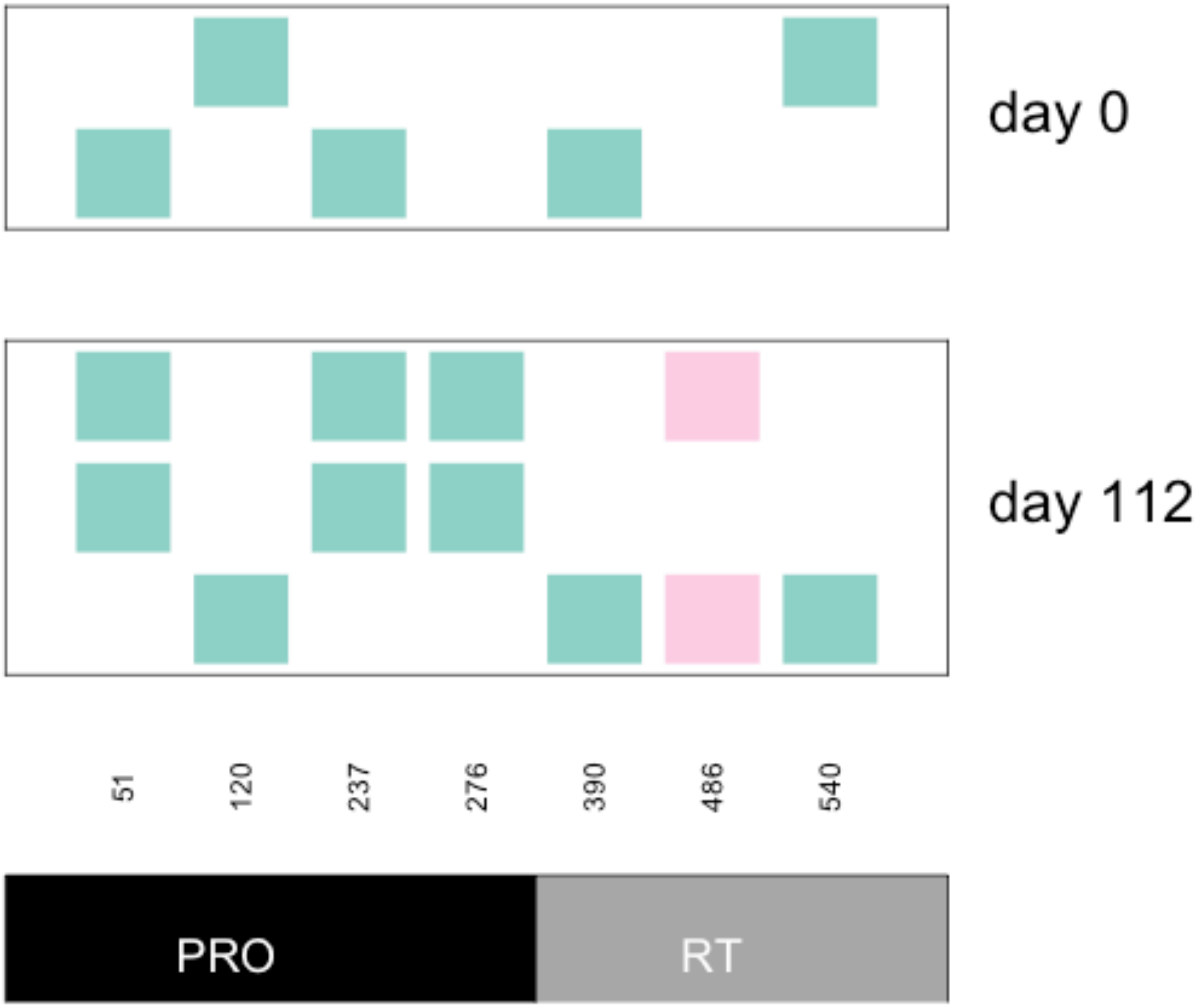


Williams, Pennings, 2018. Data from Bacheler et al 2000

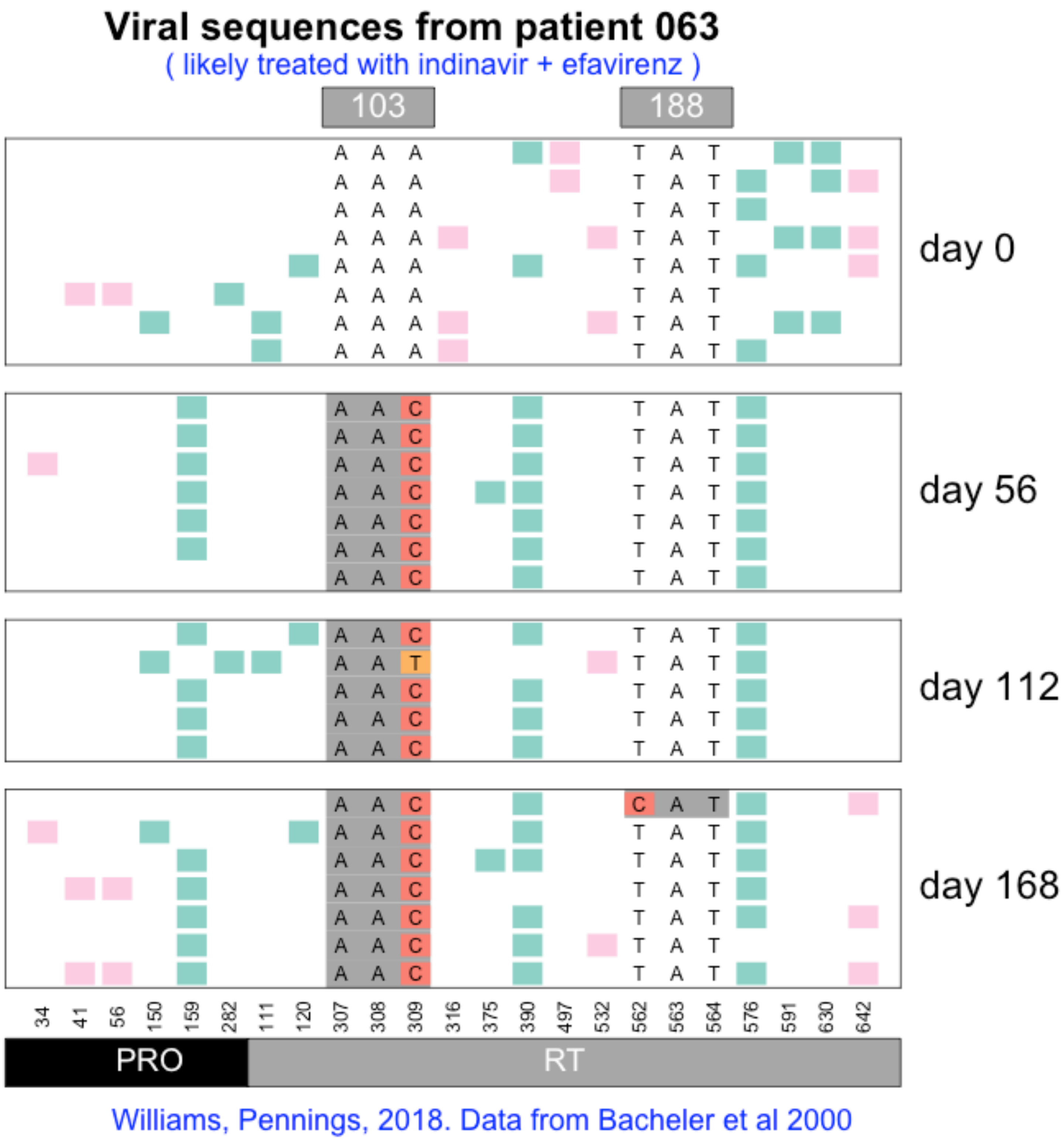

Viral sequences from patient 063
( likely treated with indinavir + efavirenz )
103
188
day 0
day 56
day 112
day 168
34 41 56 150 159 282 111 120 307 308 309 316 375 390 497 532 562 563 564 576 591 630 642
PRO
RT
Williams, Pennings, 2018. Data from Bacheler et al 2000

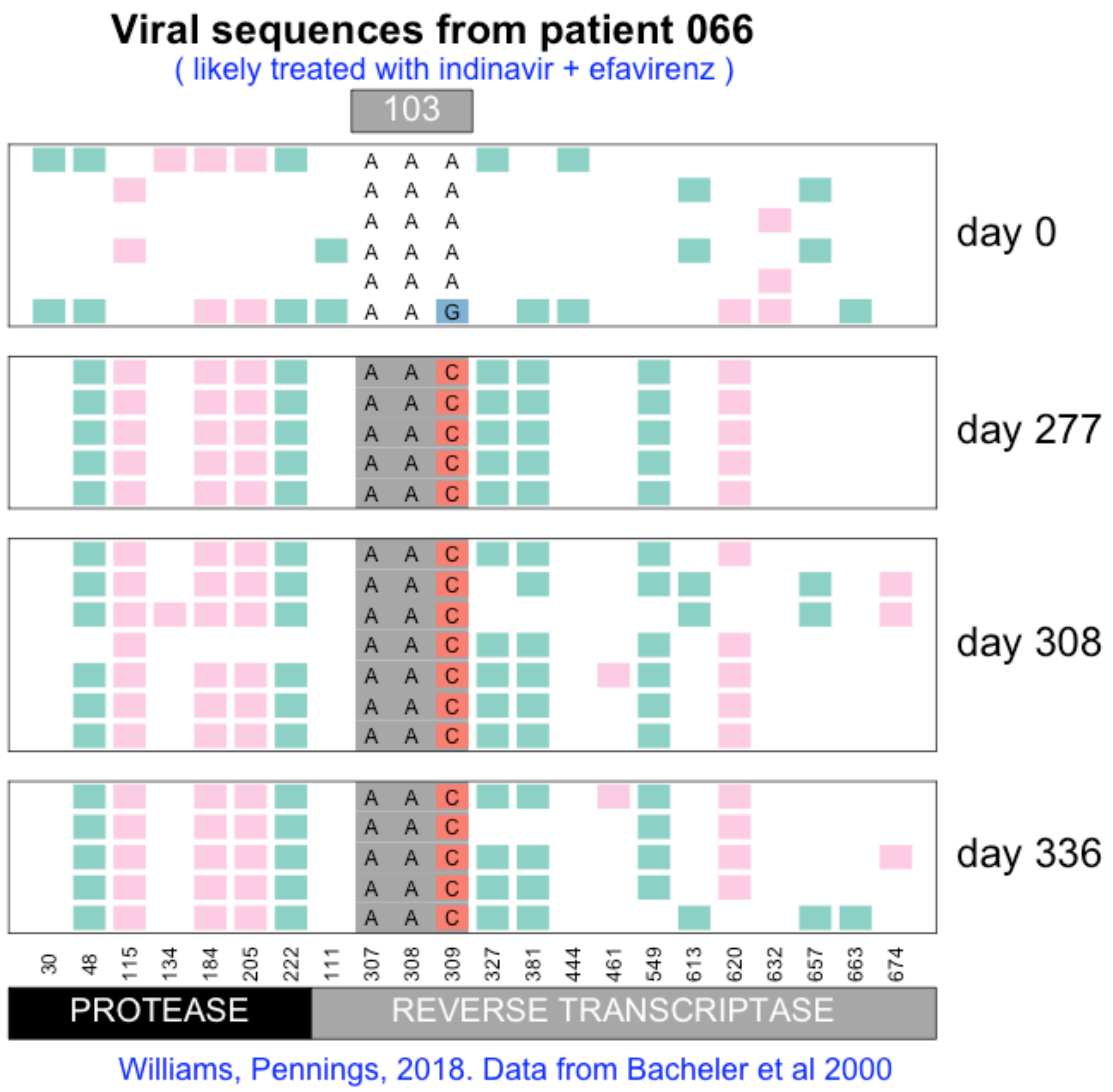

Viral sequences from patient 066
( likely treated with indinavir + efavirenz )
103
day 0
day 277
day 308
day 336
30
48
115
134
184
205
222
111
307
308
309
327
381
444
461
549
613
620
632
657
663
674
PROTEASE
REVERSE TRANSCRIPTASE
Williams, Pennings, 2018. Data from Bacheler et al 2000

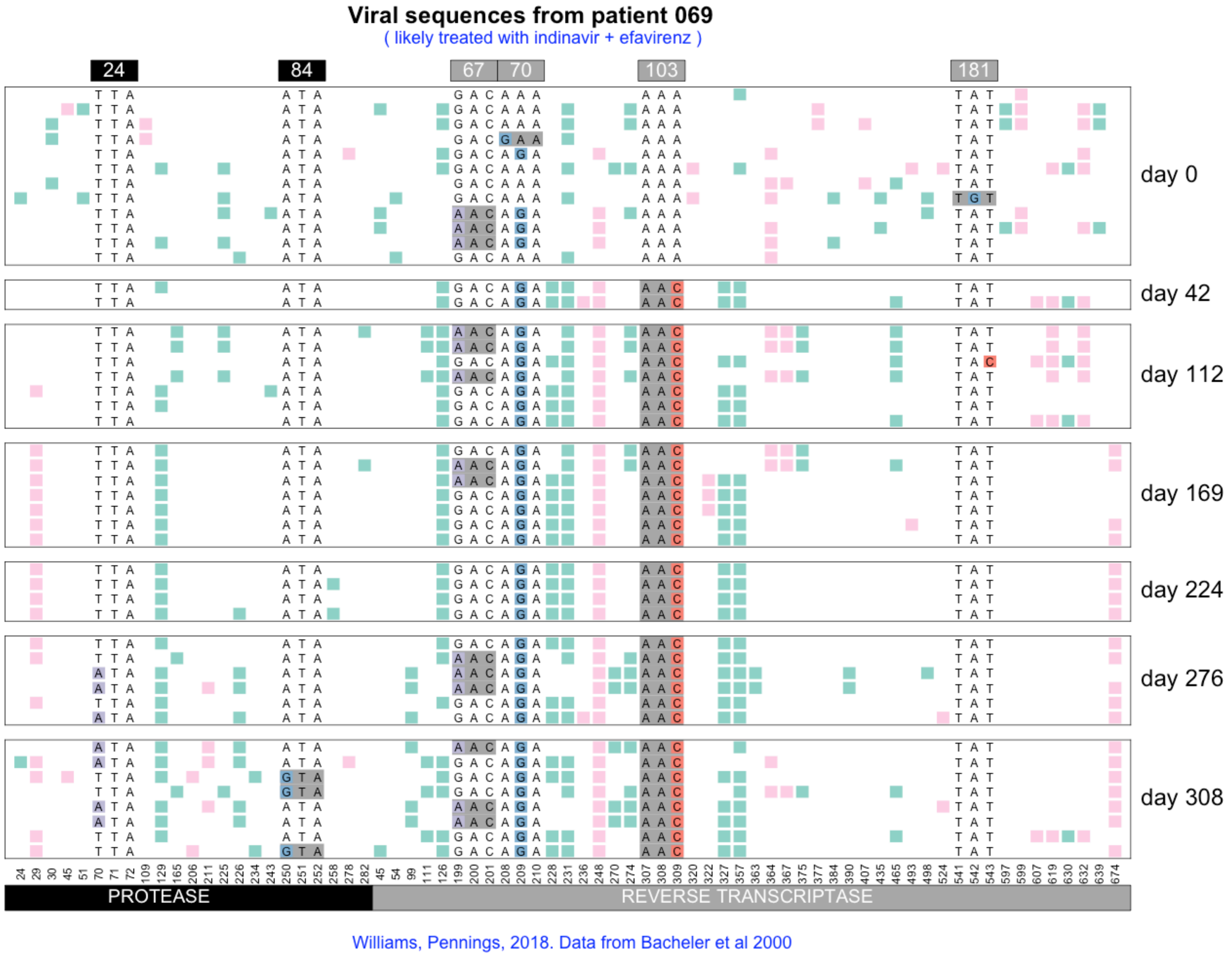

Viral sequences from patient 069
( likely treated with indinavir + efavirenz )
24
84
67
70
103
181
day 0
day 42
day 112
day 169
day 224
day 276
day 308
PROTEASE
REVERSE TRANSCRIPTASE
Williams, Pennings, 2018. Data from Bacheler et al 2000

#### Viral sequences from patient 070

( likely treated with indinavir then later switched to efavirenz combination therapy )

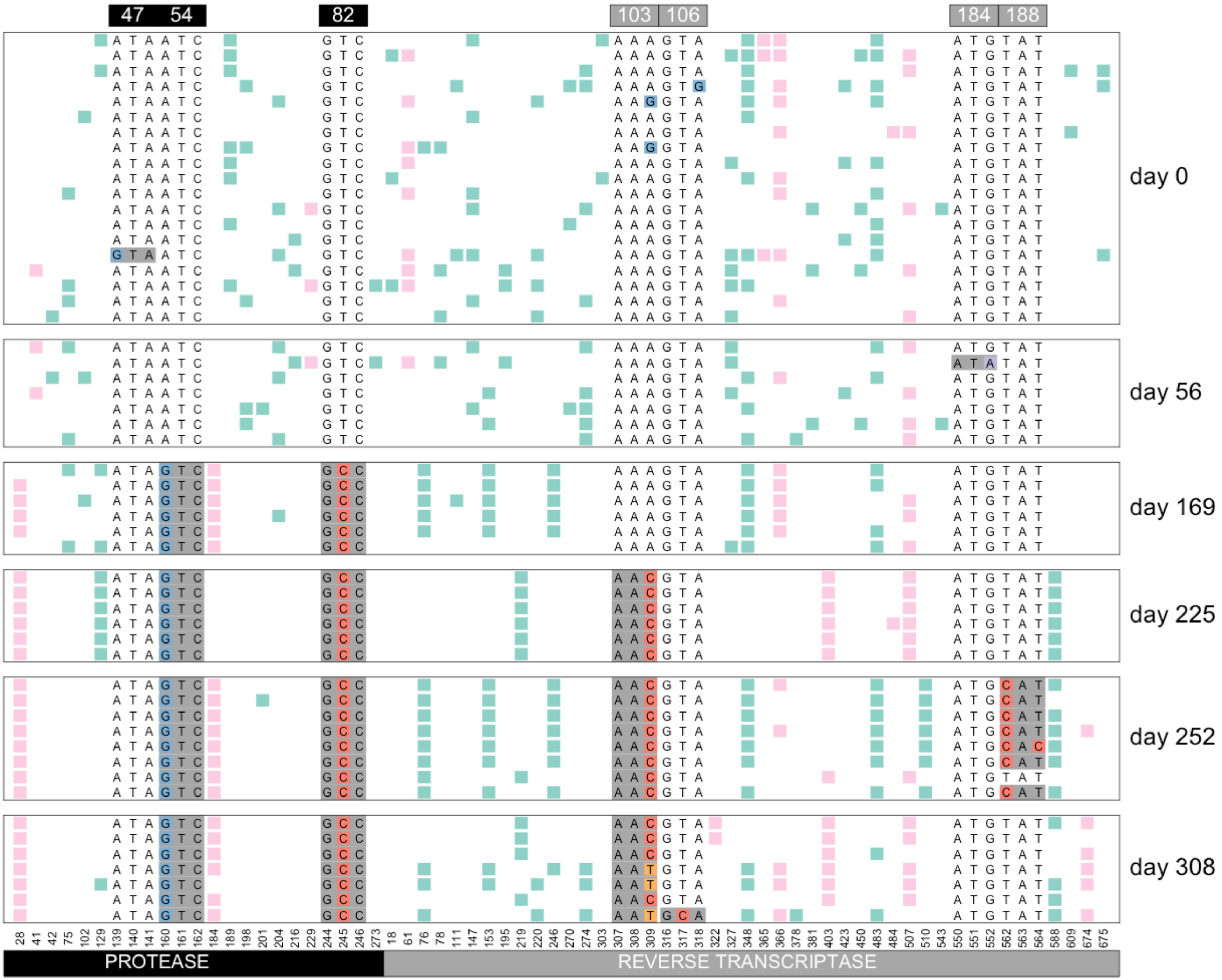


Williams, Pennings, 2018. Data from Bacheler et al 2000

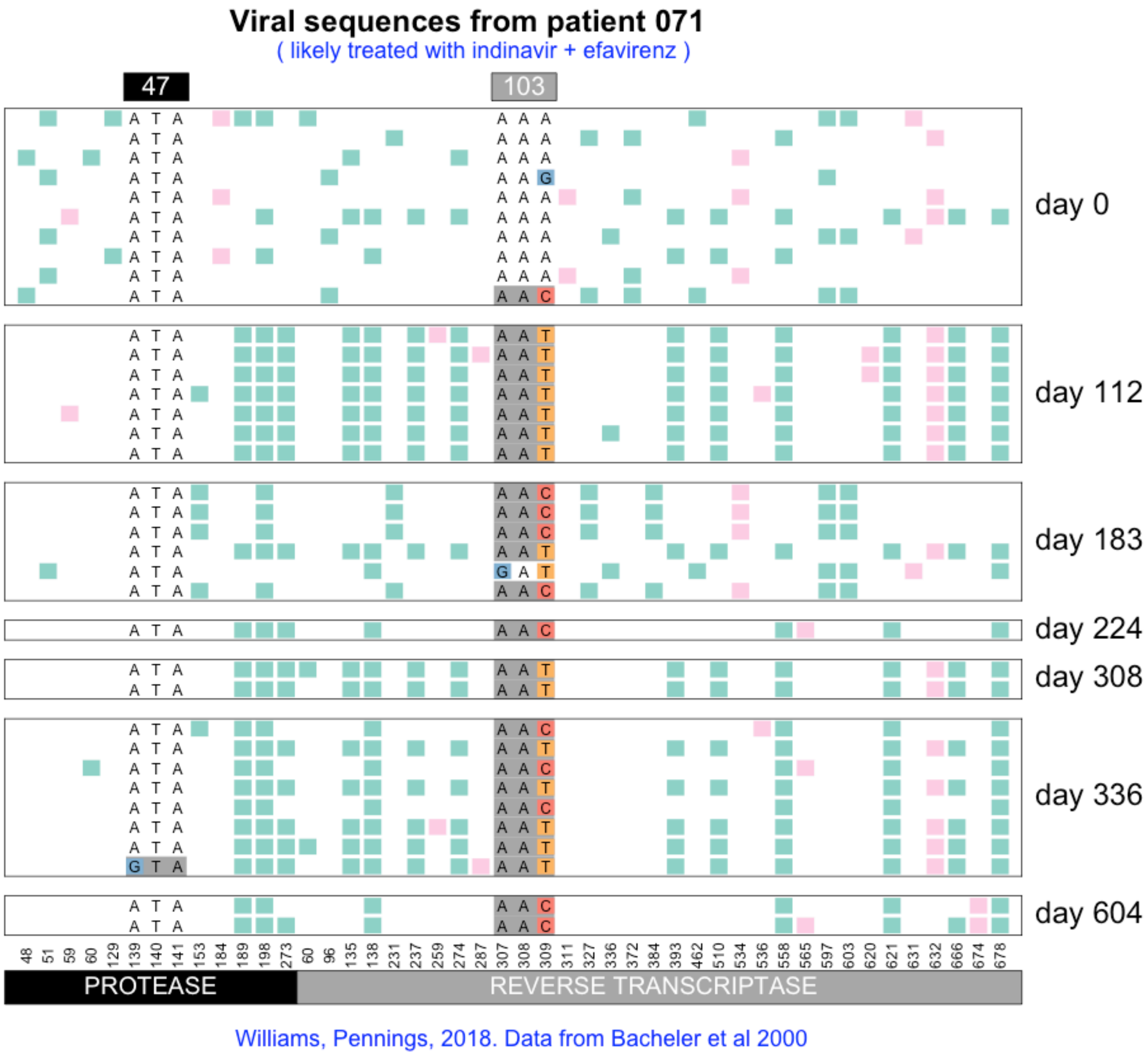
Viral sequences from patient 071
( likely treated with indinavir + efavirenz )
47
103
day 0
day 112
day 183
day 224
day 308
day 336
day 604
PROTEASE
REVERSE TRANSCRIPTASE
Williams, Pennings, 2018. Data from Bacheler et al 2000

#### Viral sequences from patient 072

( likely treated with indinavir then later switched to efavirenz combination therapy )

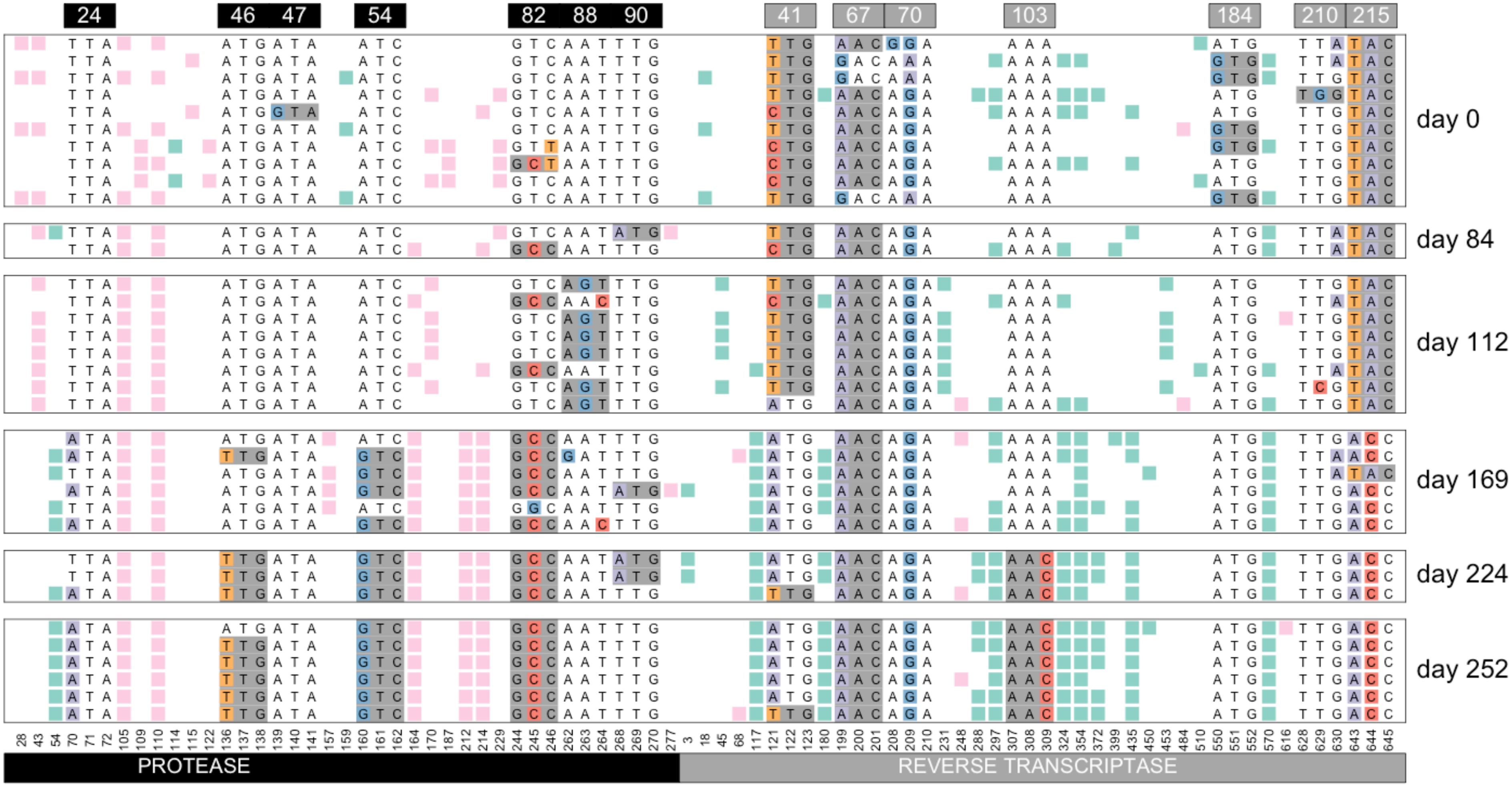


Williams, Pennings, 2018. Data from Bacheler et al 2000

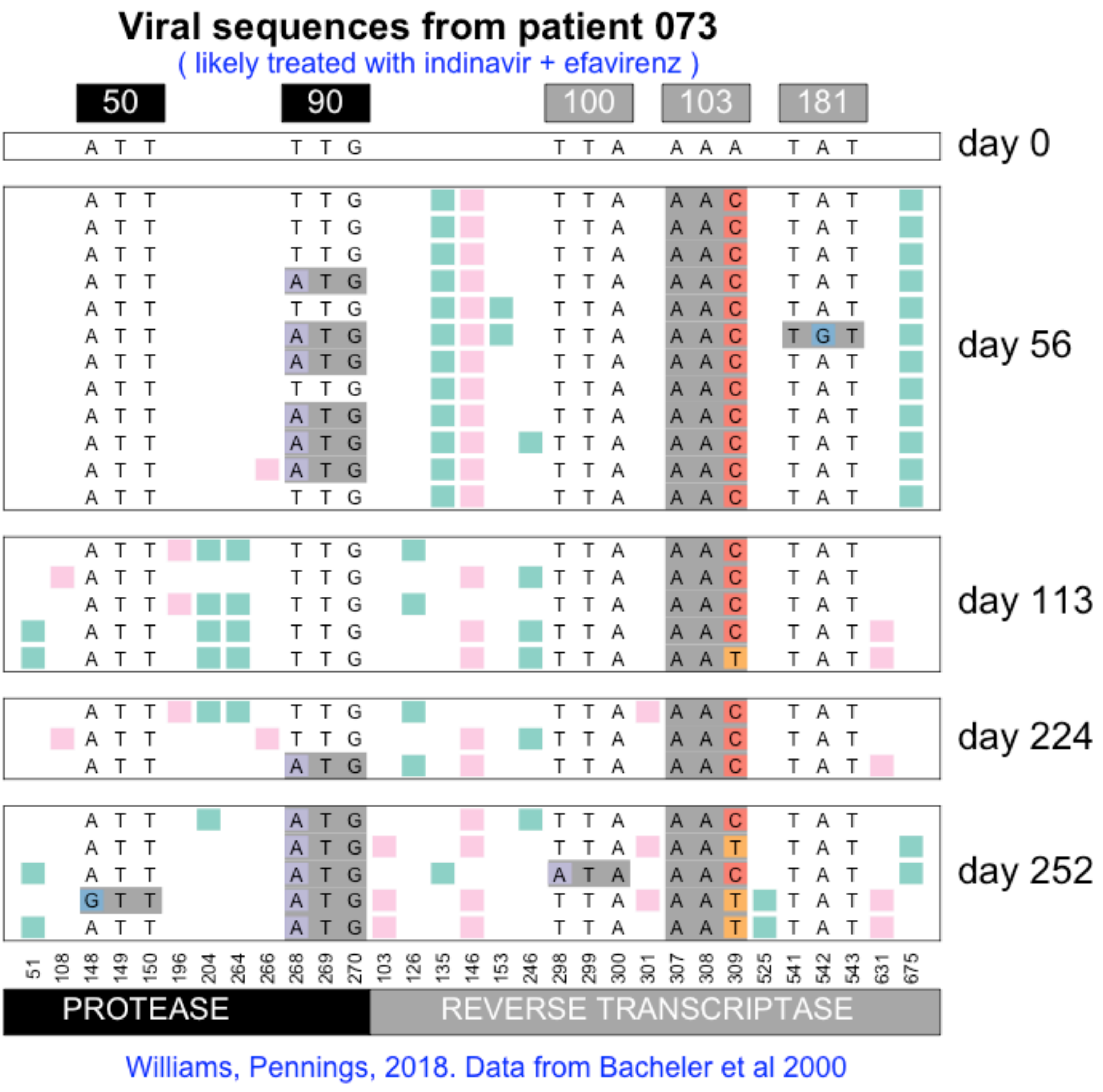

Viral sequences from patient 073
( likely treated with indinavir + efavirenz )
50
90
100
103
181
day 0
day 56
day 113
day 224
day 252
51 108 148 149 150 196 204 264 266 268 269 270 103 126 135 146 153 246 298 299 300 301 307 308 309 525 541 542 543 631 675
PROTEASE
REVERSE TRANSCRIPTASE
Williams, Pennings, 2018. Data from Bacheler et al 2000

#### Viral sequences from patient 077

( likely treated with indinavir + efavirenz )

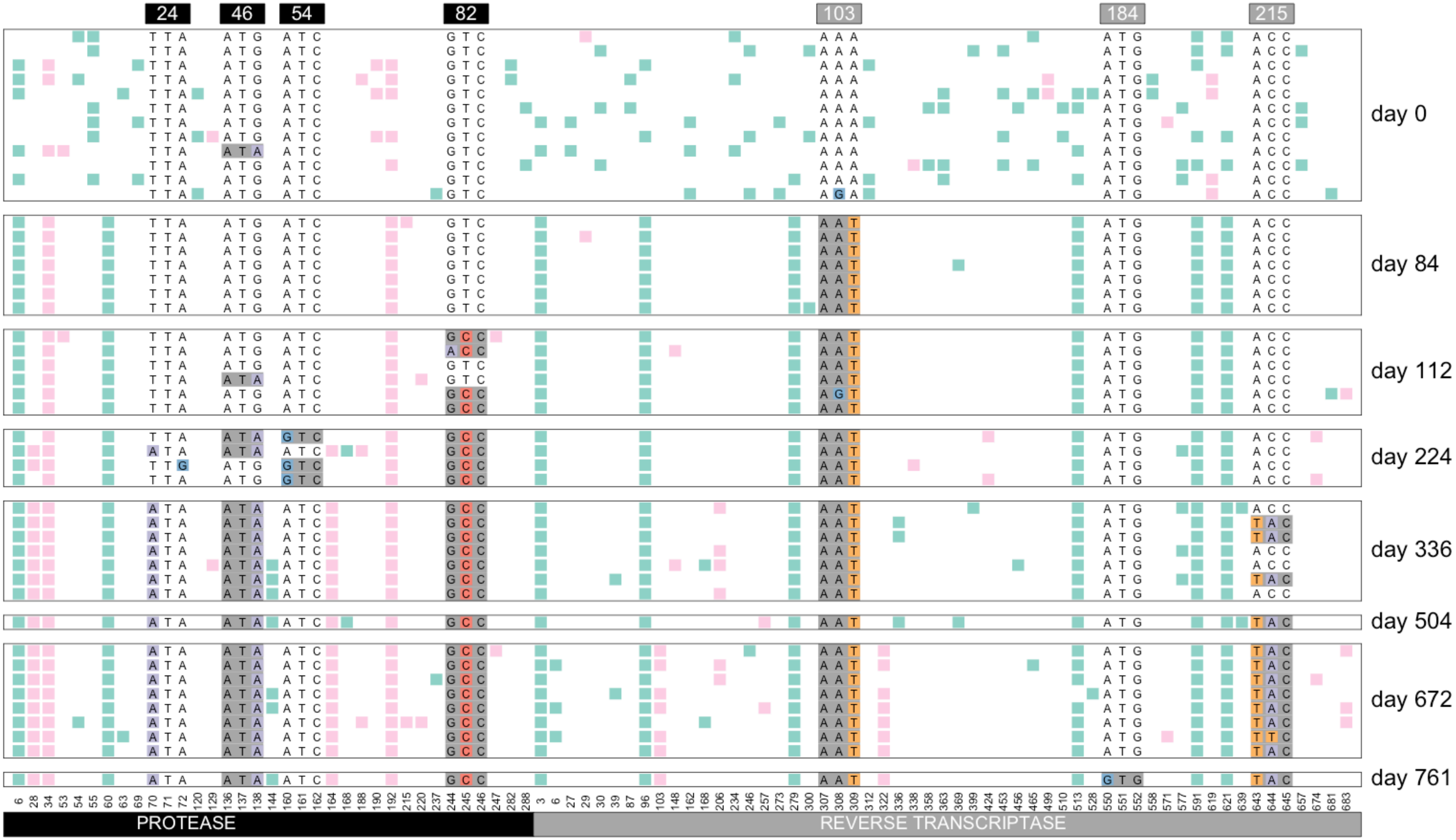


Williams, Pennings, 2018. Data from Bacheler et al 2000

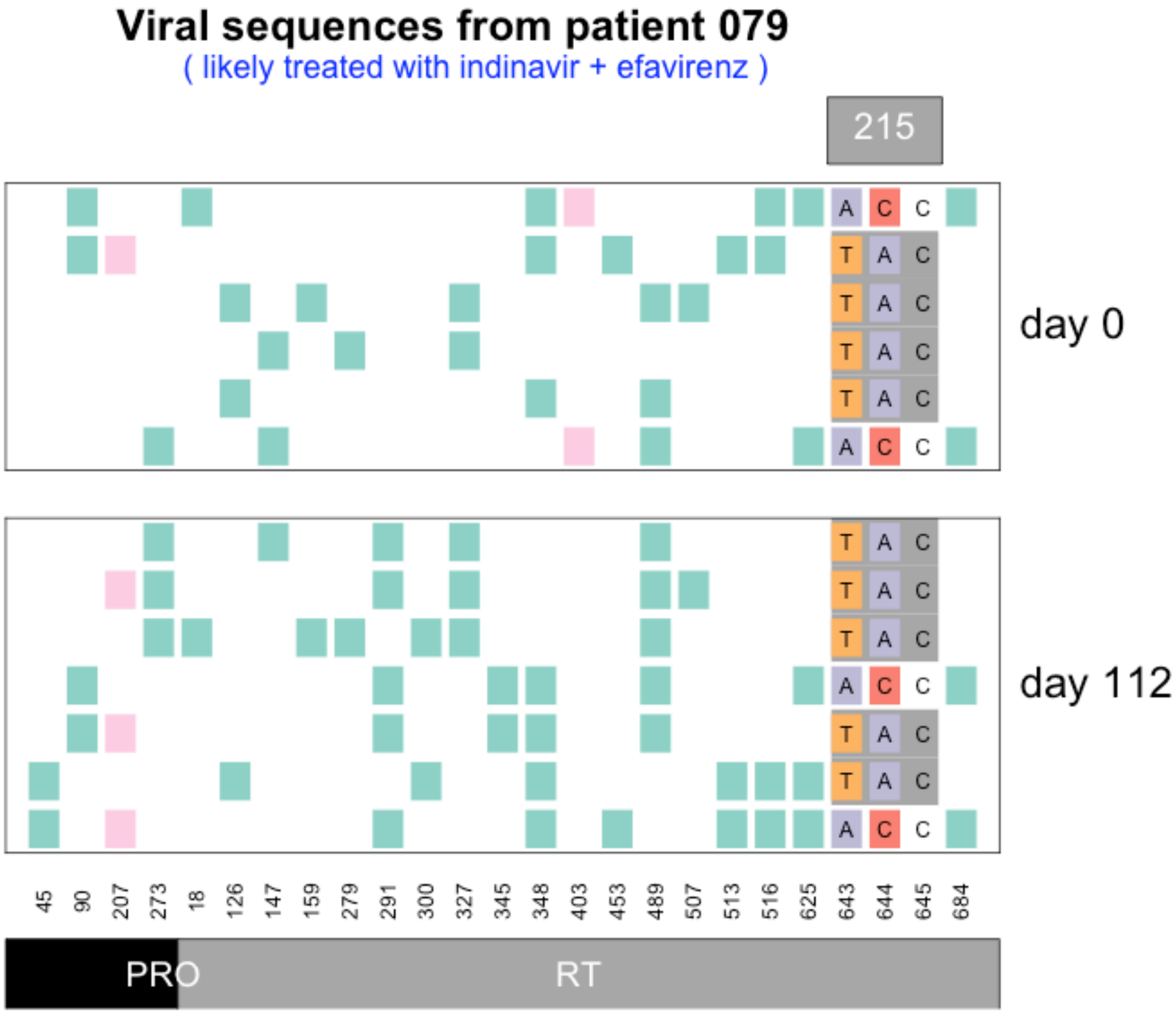


Williams, Pennings, 2018. Data from Bacheler et al 2000

#### Viral sequences from patient 080

( likely treated with indinavir then later switched to efavirenz combination therapy )

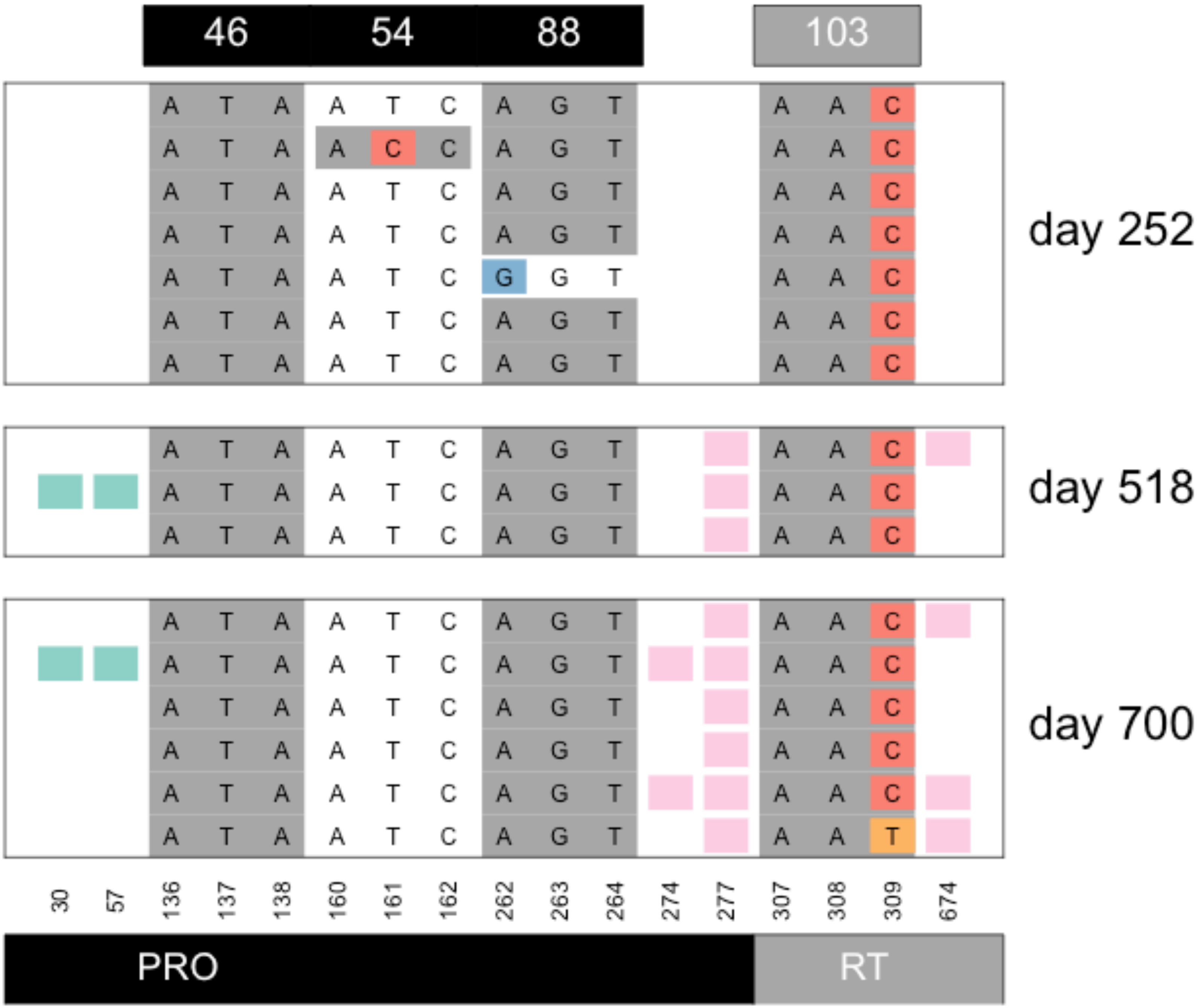


Williams, Pennings, 2018. Data from Bacheler et al 2000

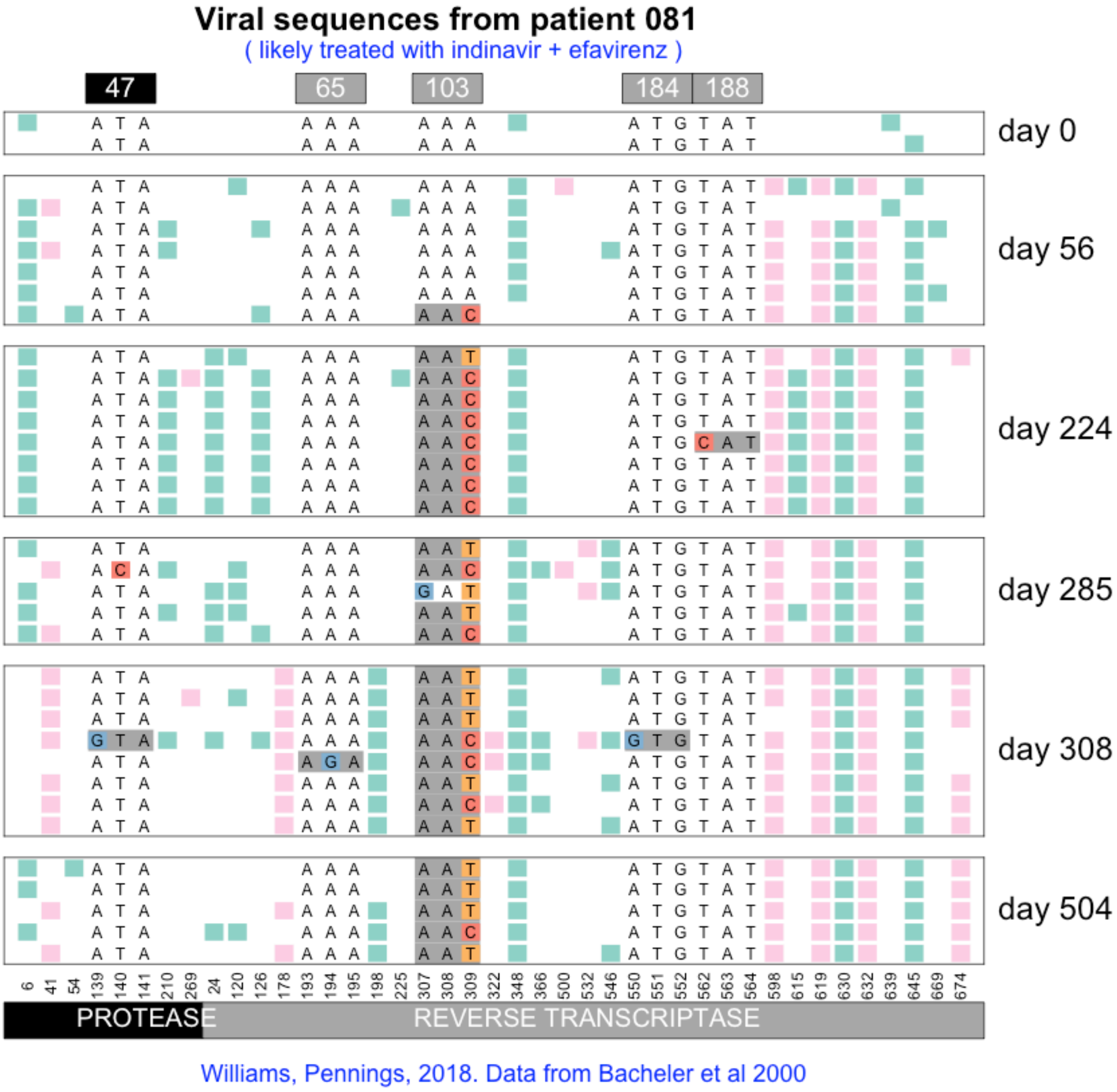

Viral sequences from patient 081
( likely treated with indinavir + efavirenz )
47
65
103
184
188
day 0
day 56
day 224
day 285
day 308
day 504
6 41 54 139 140 141 210 269 24 120 126 178 193 194 195 198 225 307 308 309 322 348 366 500 532 546 550 551 552 562 563 564 598 615 619 630 632 639 645 669 674
PROTEASE
REVERSE TRANSCRIPTASE
Williams, Pennings, 2018. Data from Bacheler et al 2000

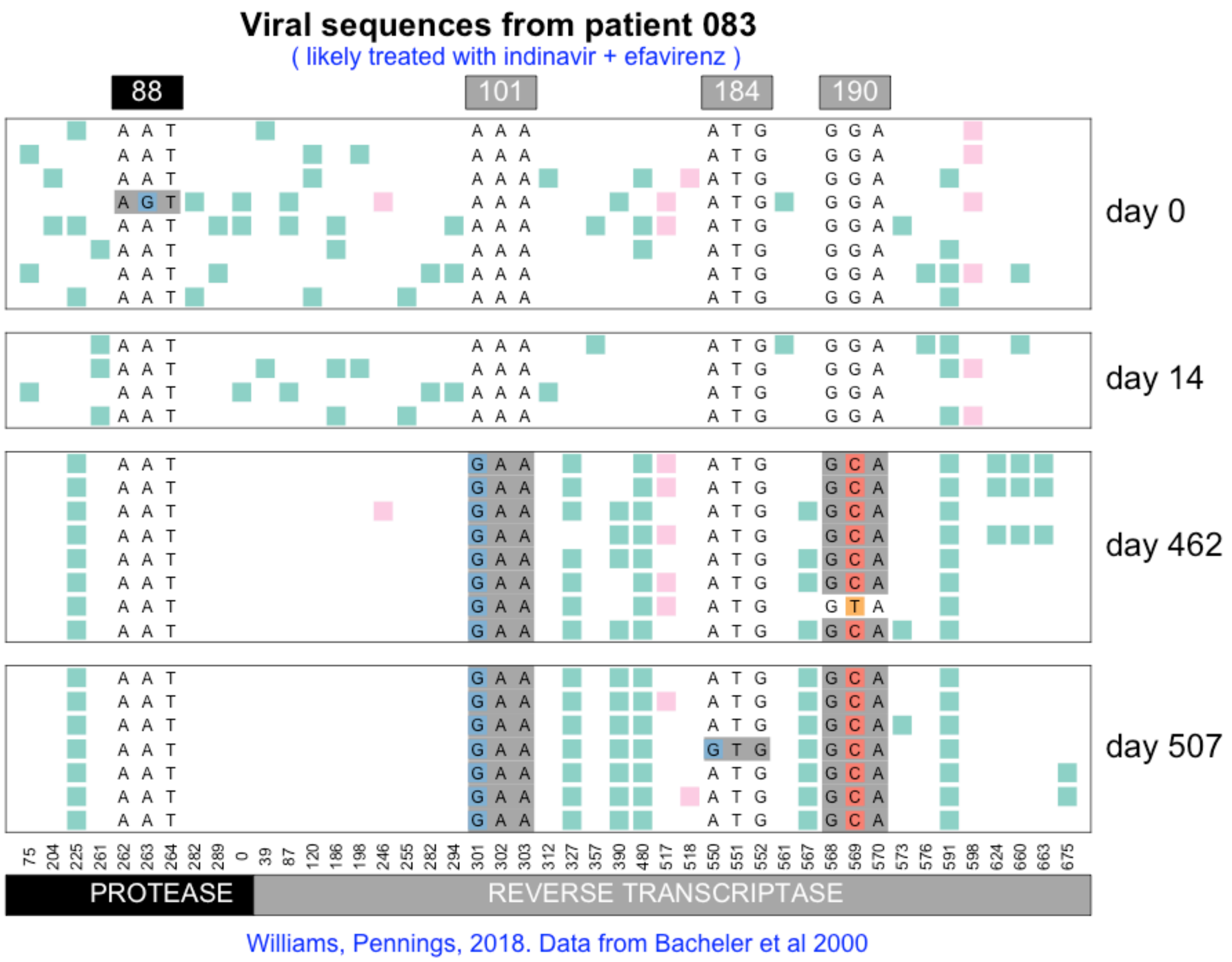
Viral sequences from patient 083
( likely treated with indinavir + efavirenz )
88
101
184
190
day 0
day 14
day 462
day 507
75 204 225 261 262 263 264 282 289 0 39 87 120 186 198 246 255 282 294 301 302 303 312 327 357 390 480 517 518 550 551 552 561 567 568 569 570 573 576 591 598 624 660 663 675
PROTEASE
REVERSE TRANSCRIPTASE
Williams, Pennings, 2018. Data from Bacheler et al 2000

#### Viral sequences from patient 084

( likely treated with indinavir then later switched to efavirenz combination therapy )

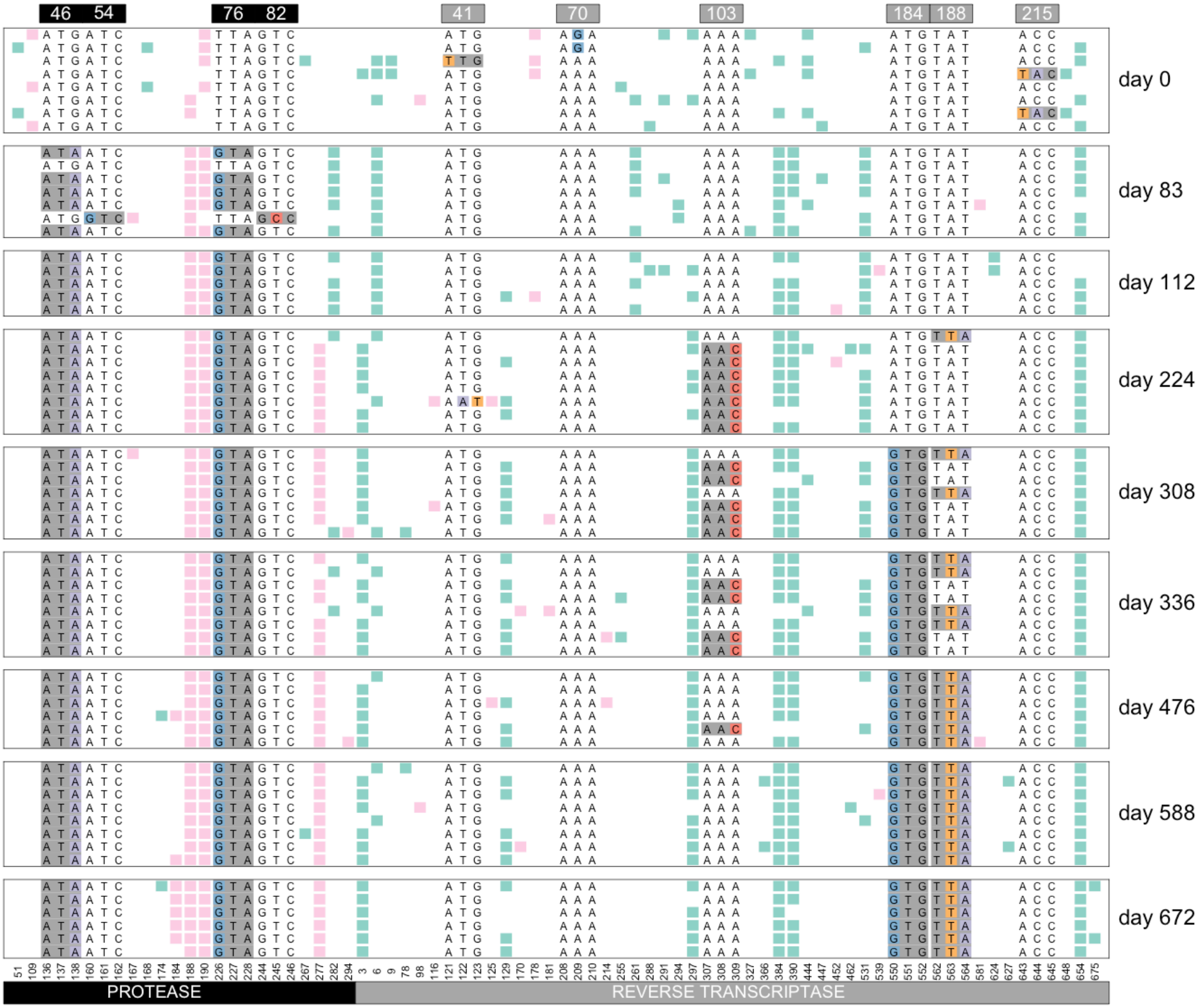


Williams, Pennings, 2018. Data from Bacheler et al 2000

### Viral sequences from patient 085

( likely treated with indinavir )

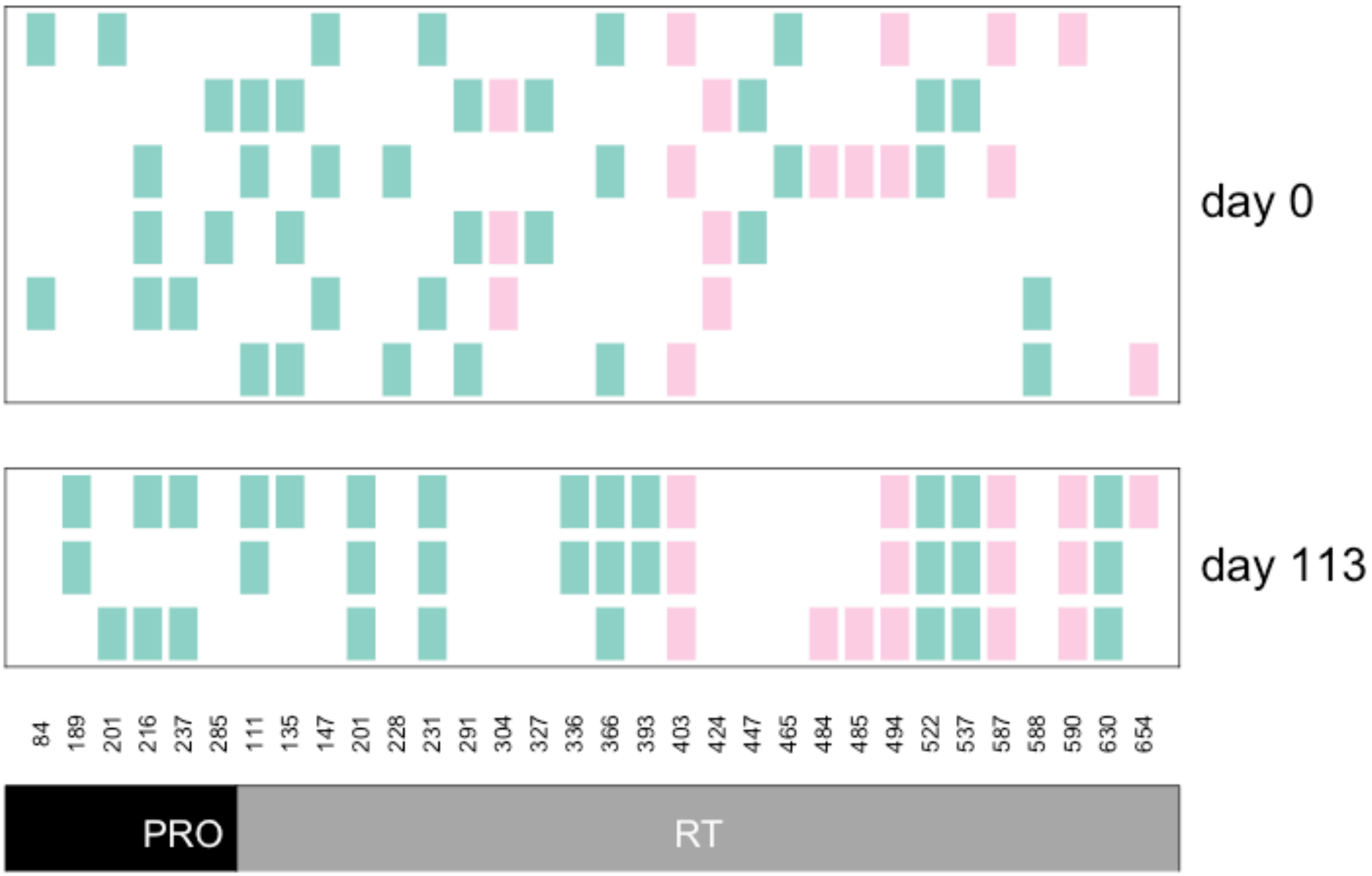


Williams, Pennings, 2018. Data from Bacheler et al 2000

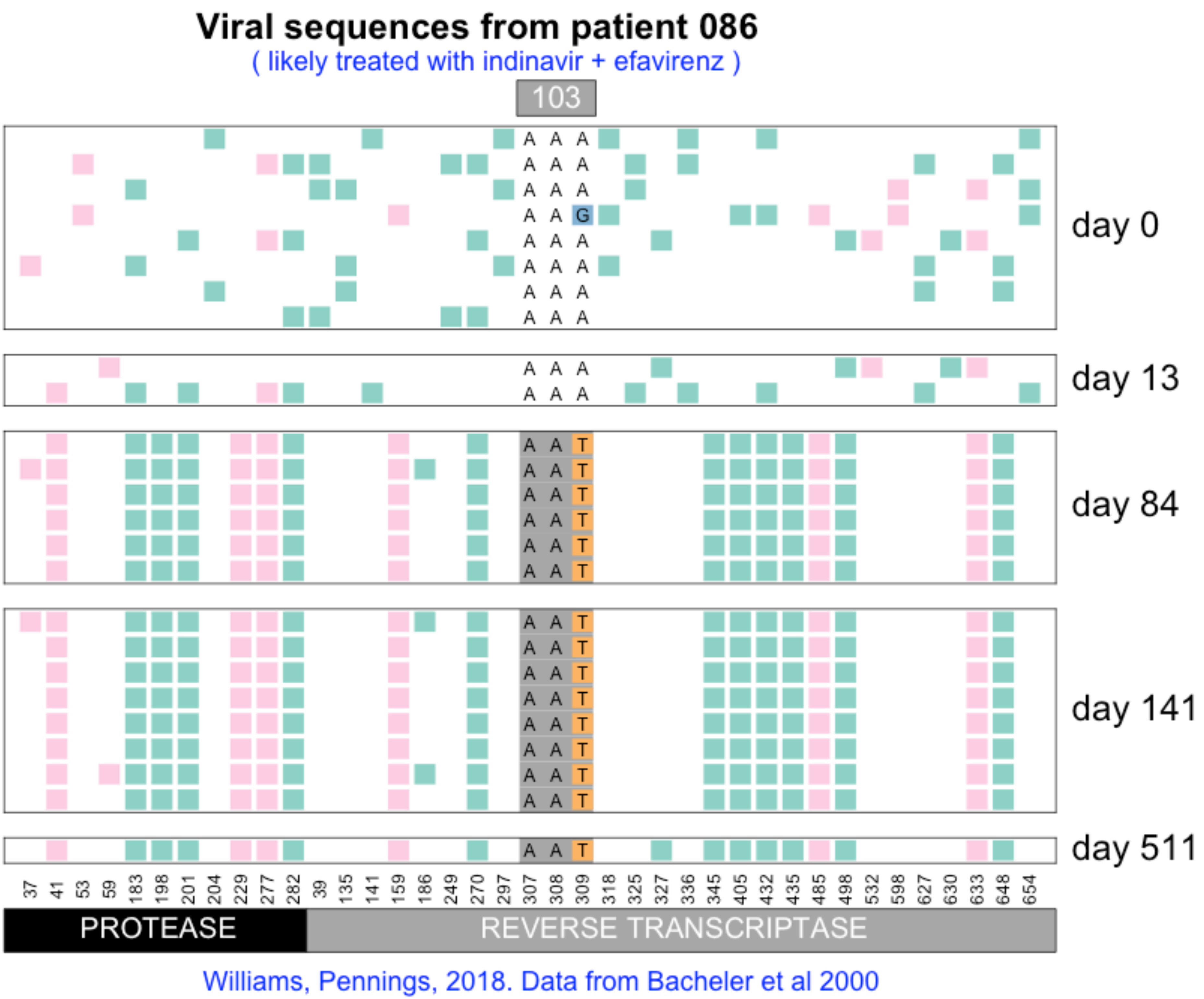

Viral sequences from patient 086
( likely treated with indinavir + efavirenz )
103
day 0
day 13
day 84
day 141
day 511
37 41 53 59 183 198 201 204 229 277 282 39 135 141 159 186 249 270 297 307 308 309 318 325 327 336 345 405 432 435 485 498 532 598 627 630 633 648 654
PROTEASE
REVERSE TRANSCRIPTASE
Williams, Pennings, 2018. Data from Bacheler et al 2000

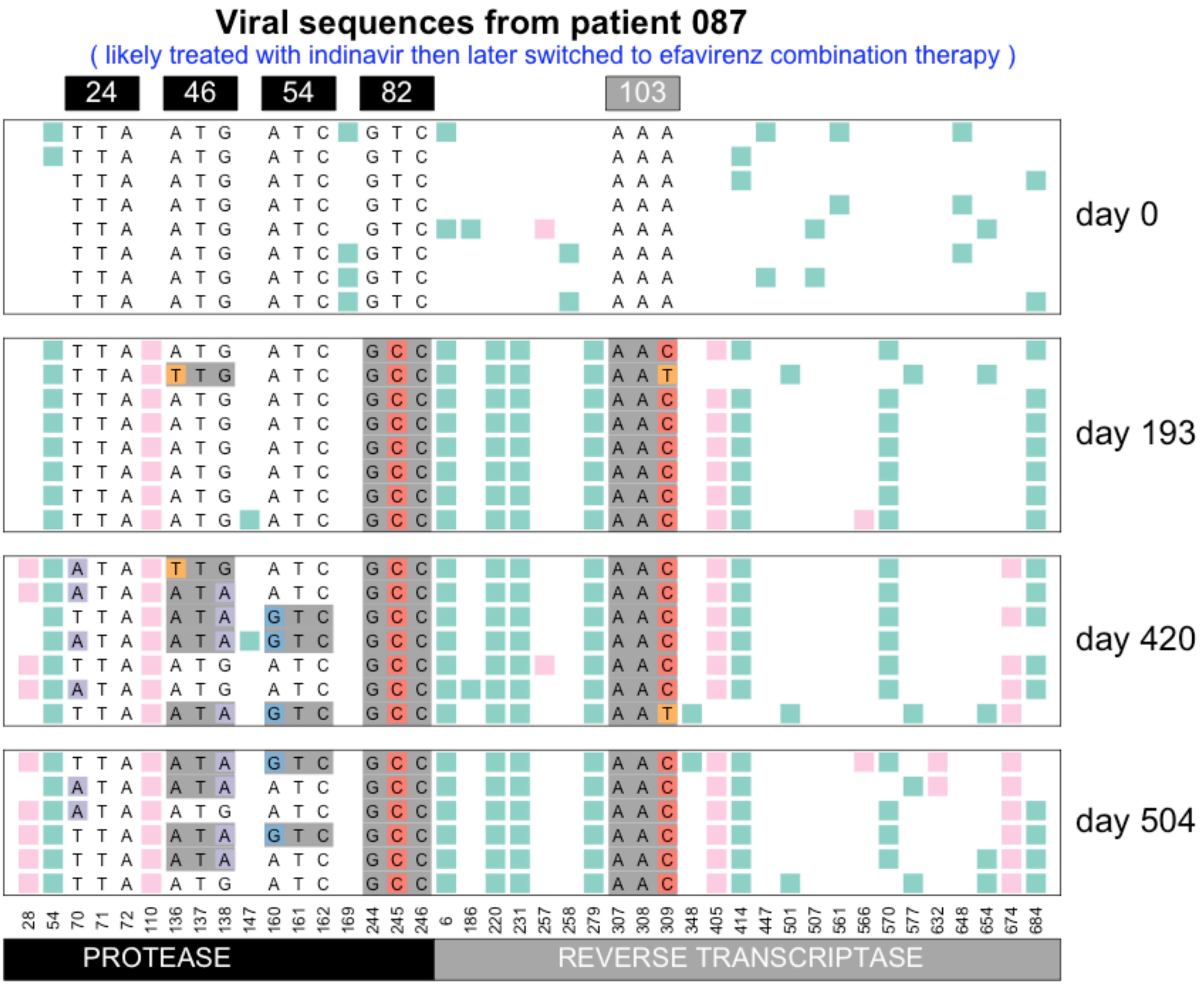


Williams, Pennings, 2018. Data from Bacheler et al 2000

#### Viral sequences from patient 089

( likely treated with indinavir + efavirenz )

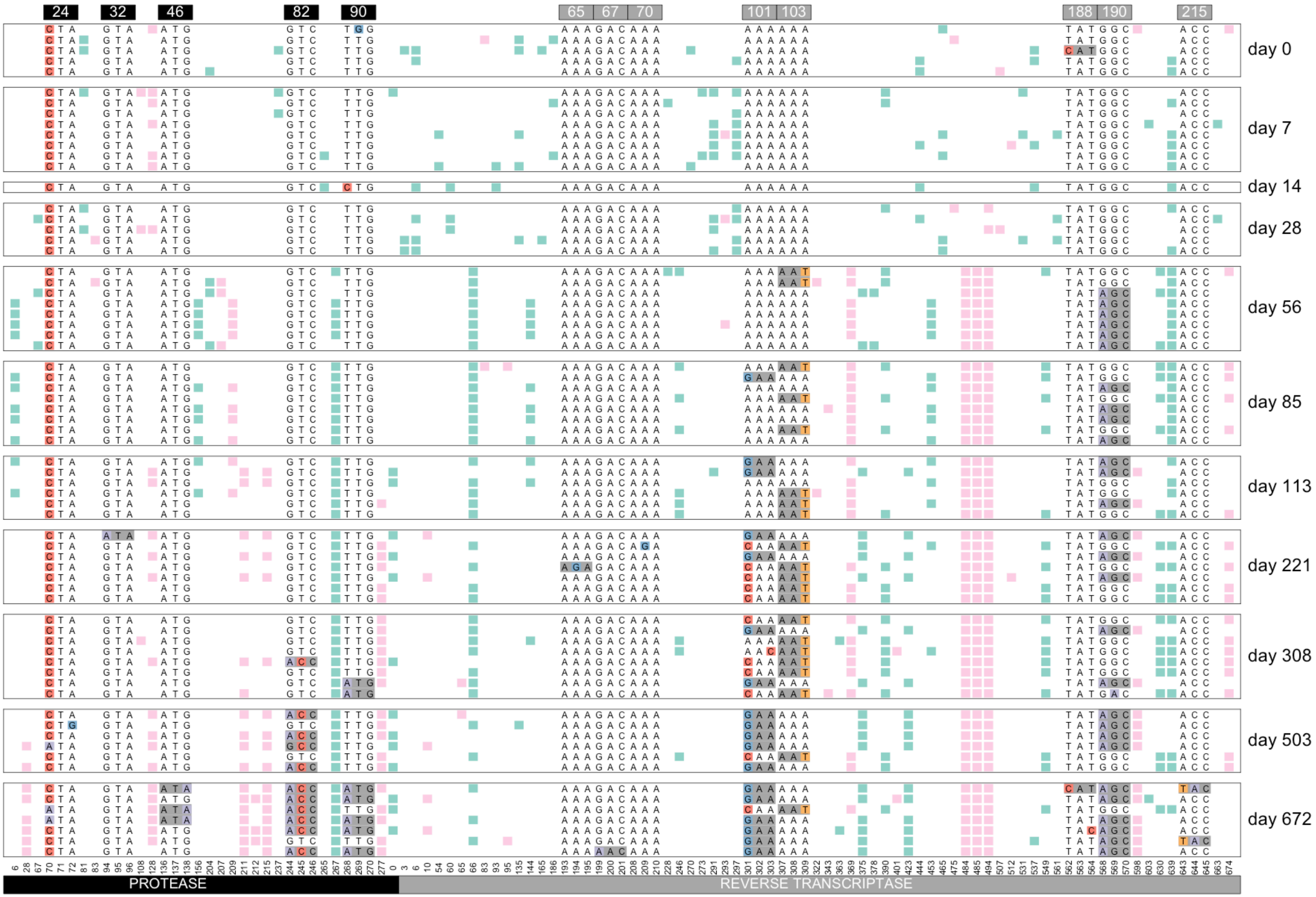


Williams, Pennings, 2018. Data from Bacheler et al 2000

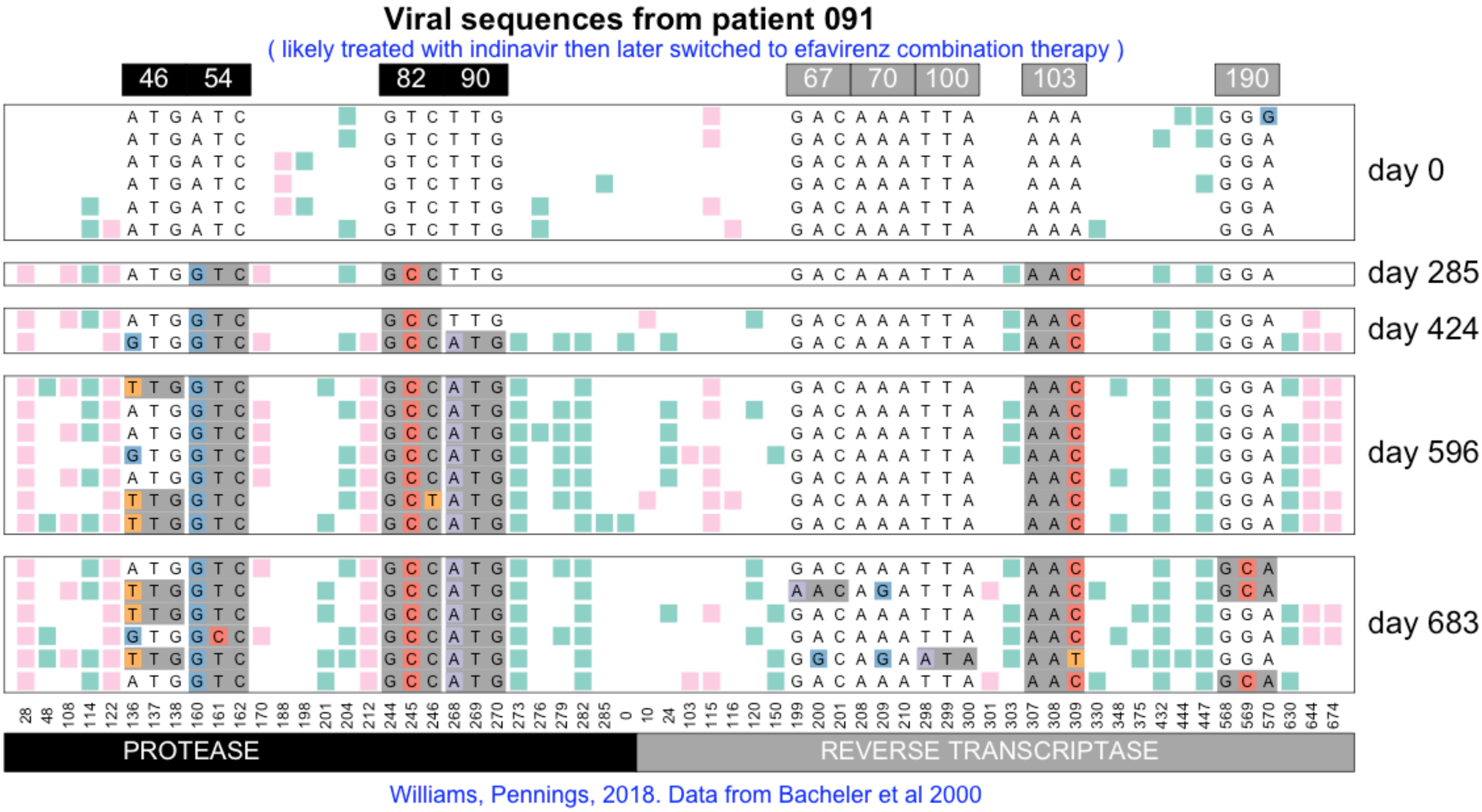
Viral sequences from patient 091
( likely treated with indinavir then later switched to efavirenz combination therapy )
46
54
82
90
67
70
100
103
190
day 0
day 285
day 424
day 596
day 683
PROTEASE
REVERSE TRANSCRIPTASE
Williams, Pennings, 2018. Data from Bacheler et al 2000

### Viral sequences from patient 093

( likely treated with indinavir then later switched to efavirenz combination therapy )

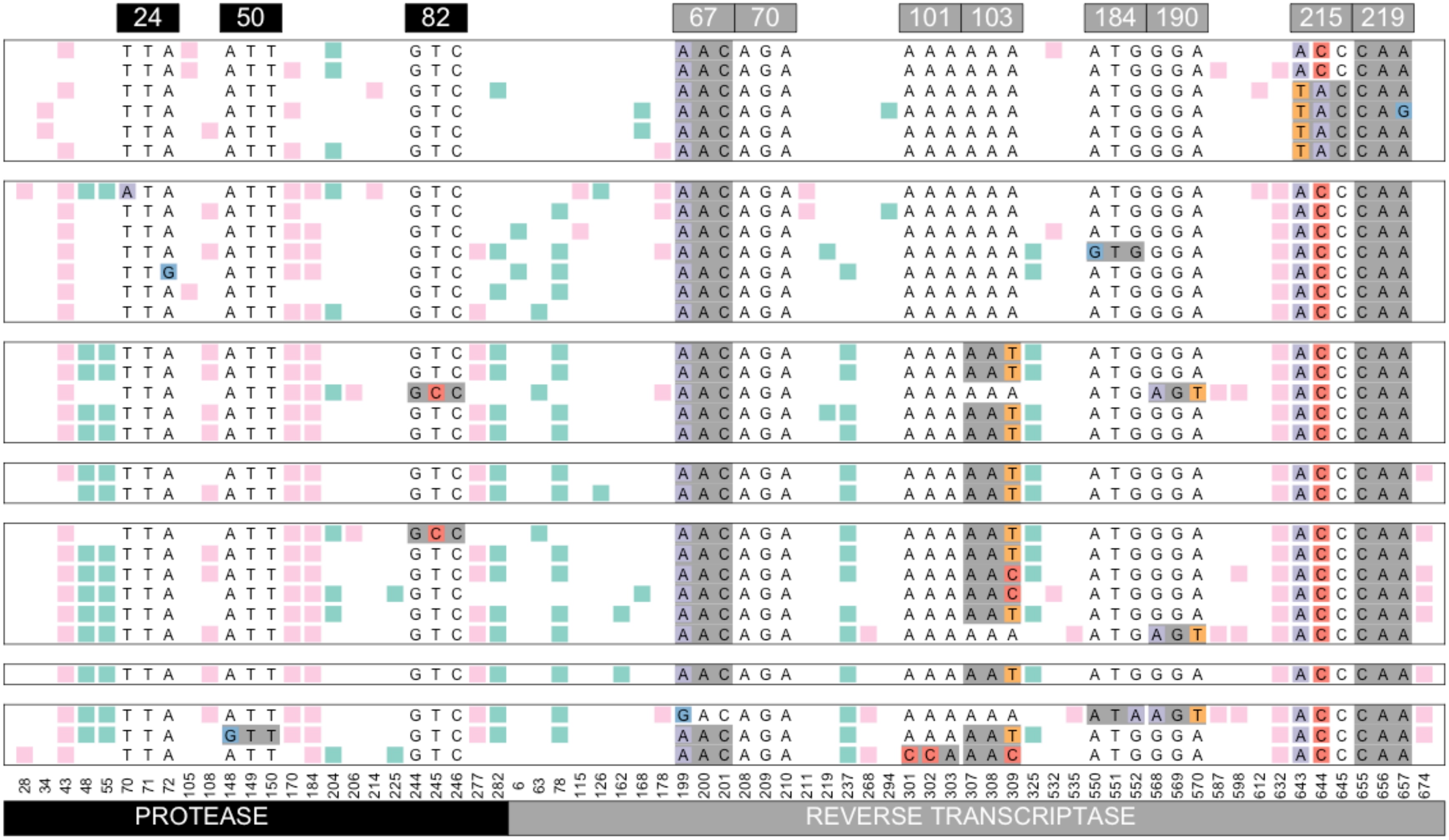


Williams, Pennings, 2018. Data from Bacheler et al 2000

#### Viral sequences from patient 094

( likely treated with indinavir + efavirenz )

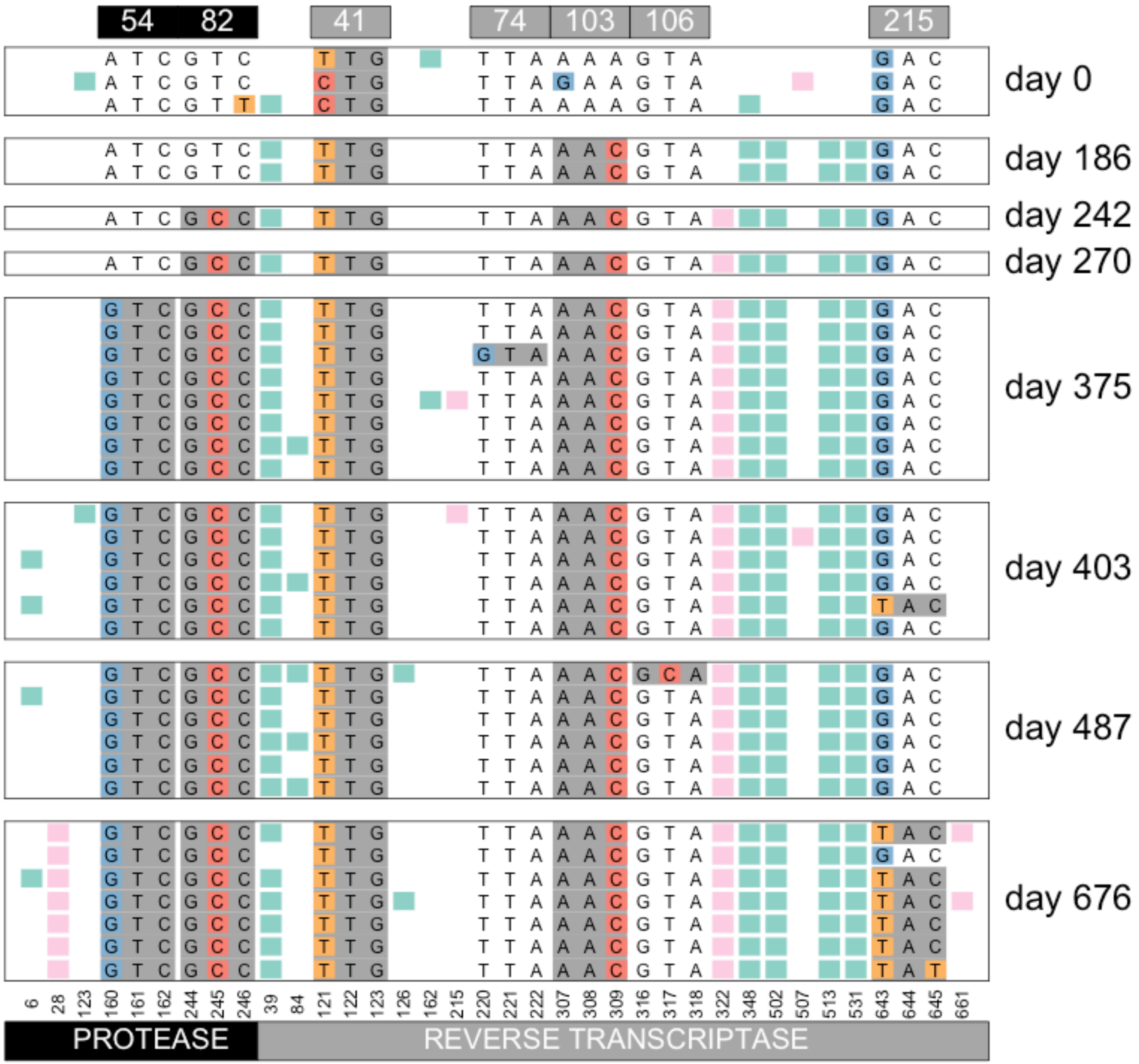


Williams, Pennings, 2018. Data from Bacheler et al 2000

#### Viral sequences from patient 095

( likely treated with indinavir then later switched to efavirenz combination therapy )

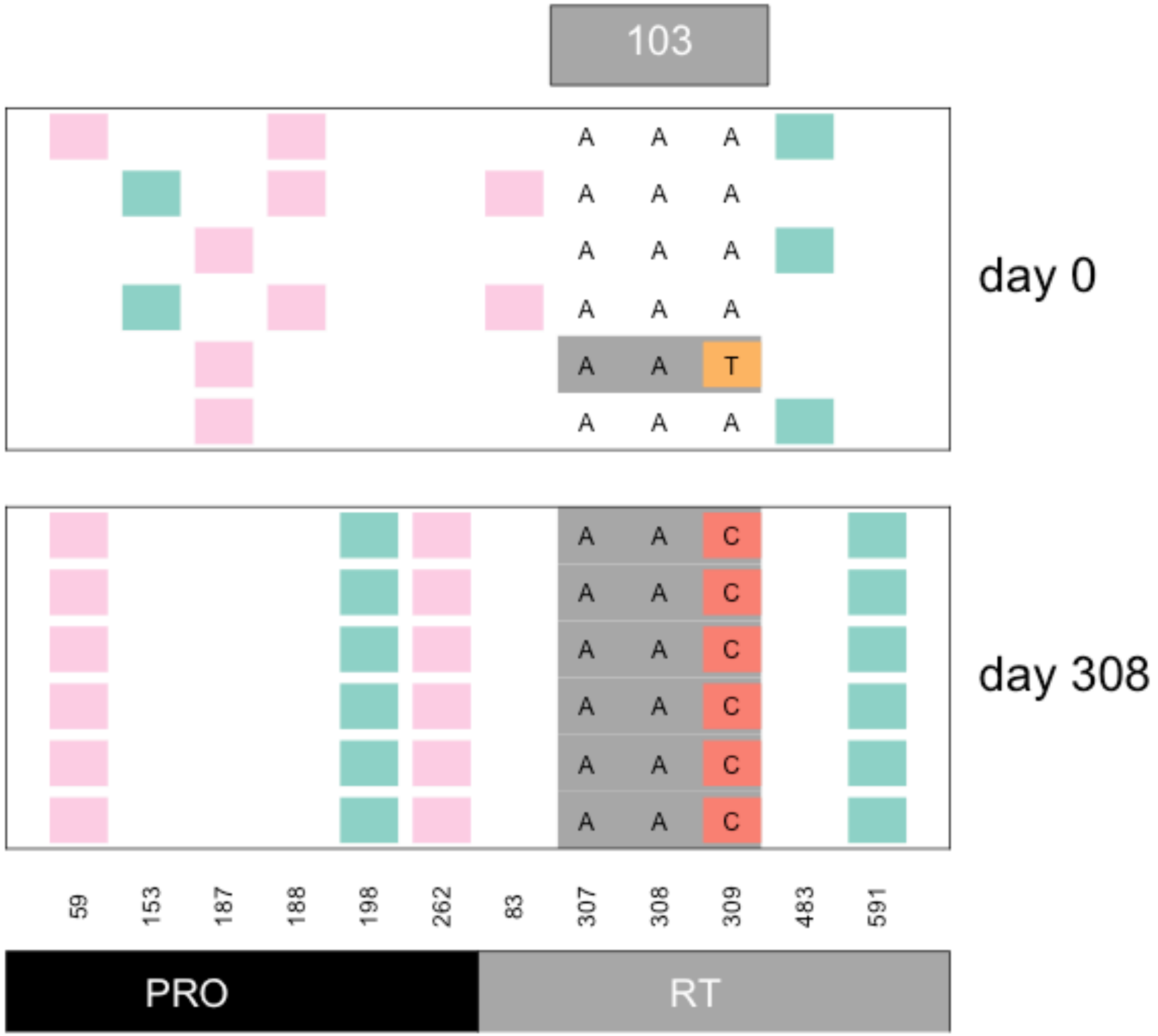


Williams, Pennings, 2018. Data from Bacheler et al 2000

### Viral sequences from patient 098

( likely treated with indinavir + efavirenz )

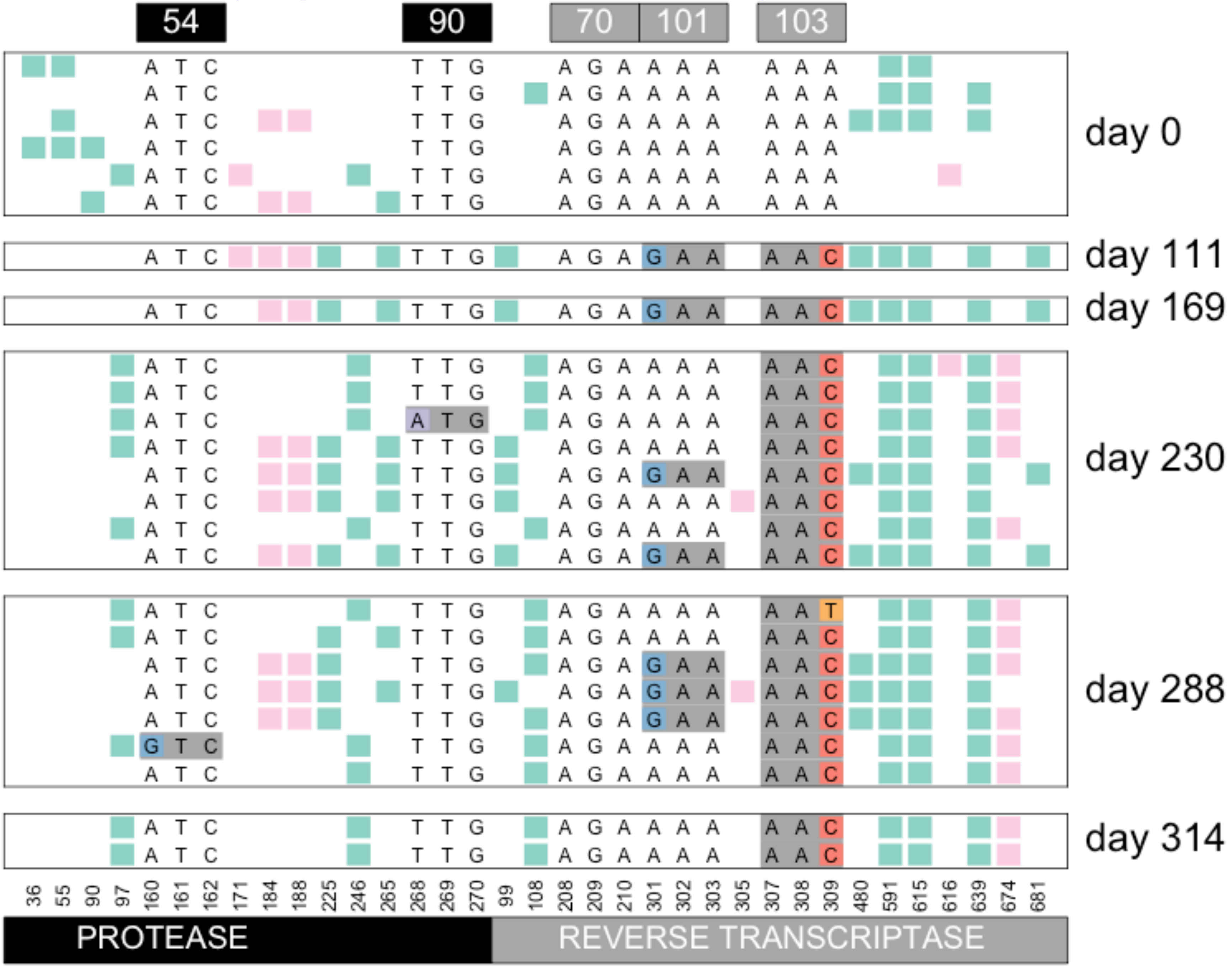


Williams, Pennings, 2018. Data from Bacheler et al 2000

#### Viral sequences from patient 099

( likely treated with indinavir + efavirenz )

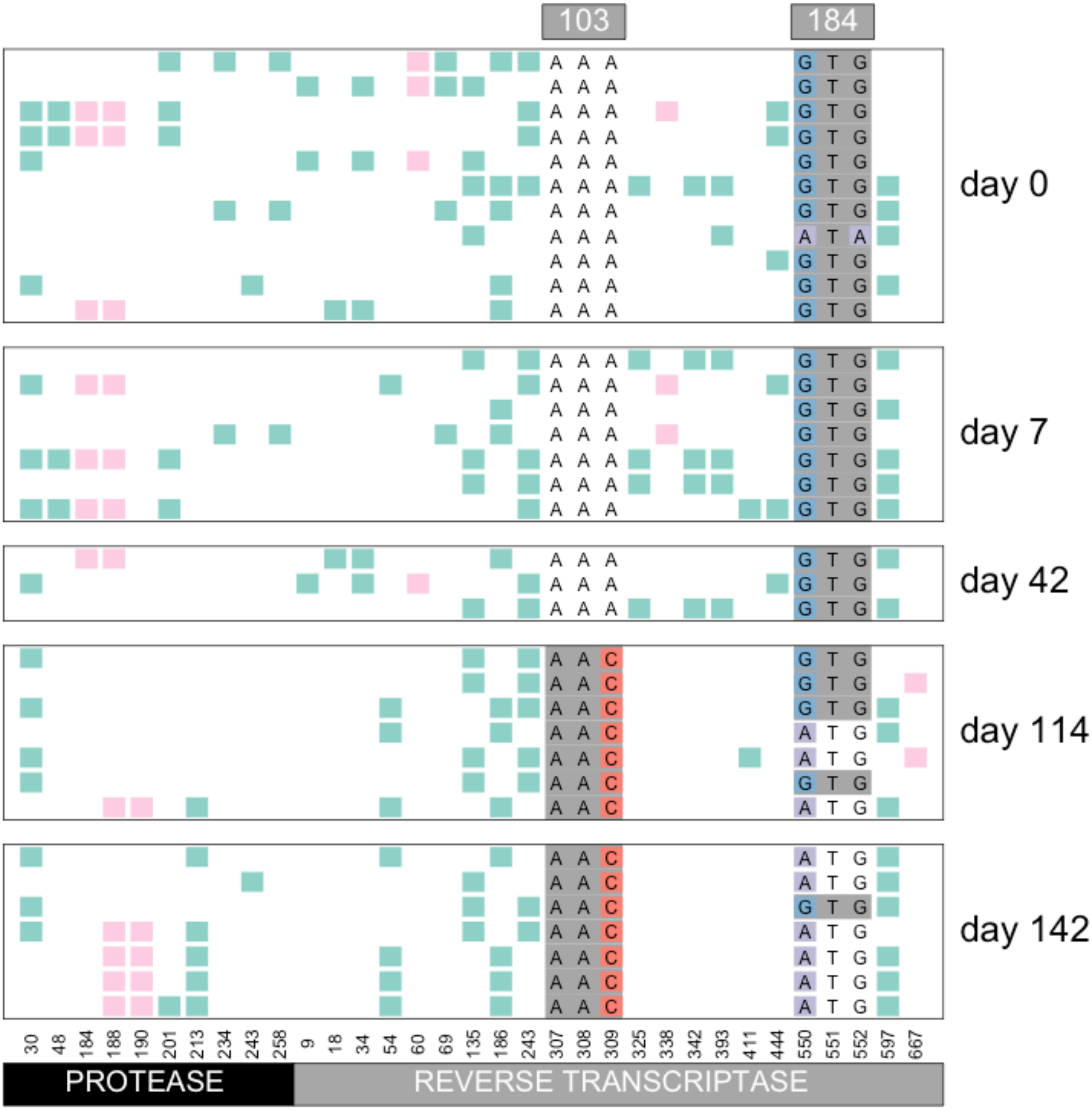


Williams, Pennings, 2018. Data from Bacheler et al 2000

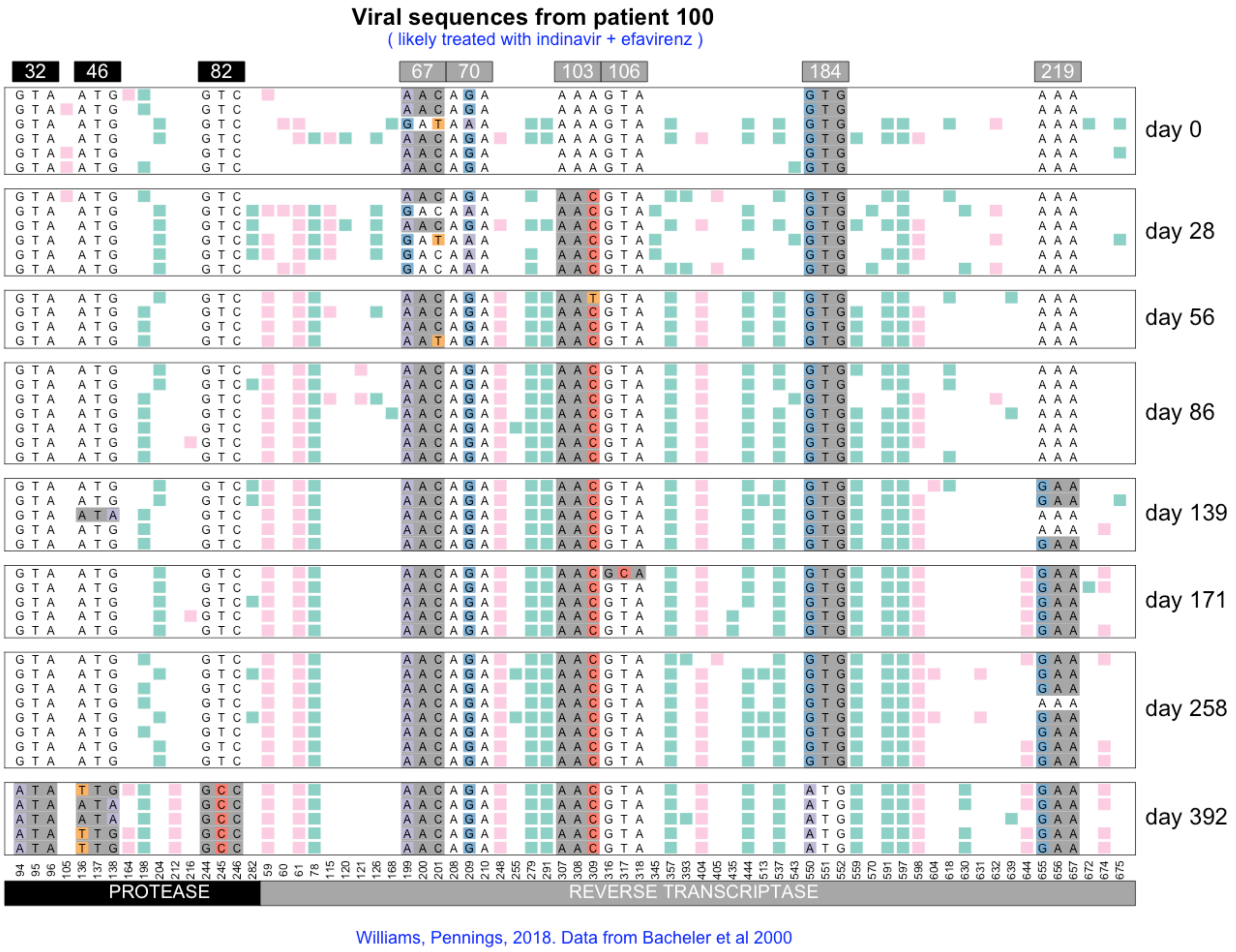
Viral sequences from patient 100
( likely treated with indinavir + efavirenz )
32
46
82
67
70
103
106
184
219
day 0
day 28
day 56
day 86
day 139
day 171
day 258
day 392
PROTEASE
REVERSE TRANSCRIPTASE
Williams, Pennings, 2018. Data from Bacheler et al 2000

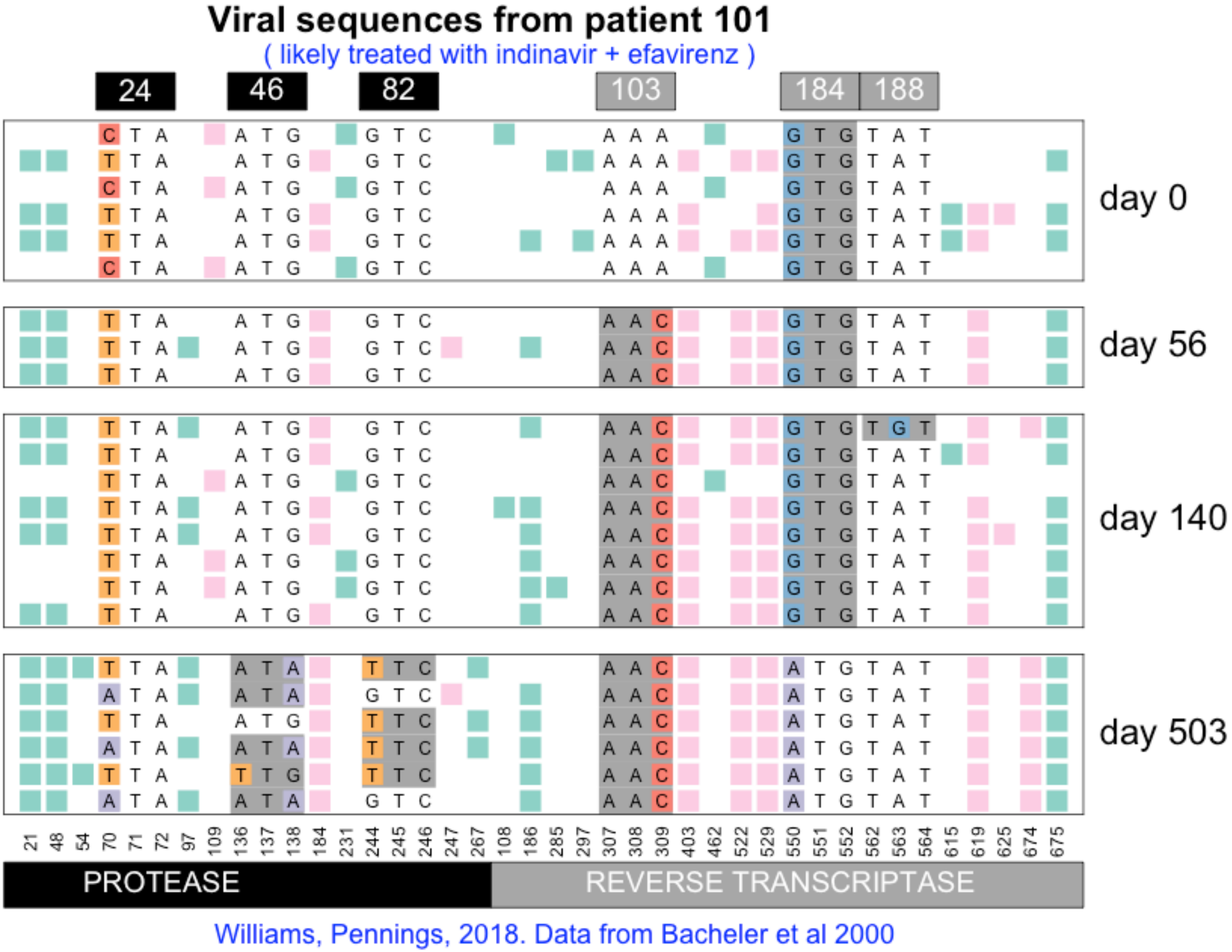

Viral sequences from patient 101
( likely treated with indinavir + efavirenz )
24
46
82
103
184
188
day 0
day 56
day 140
day 503
21 48 54 70 71 72 97 109 136 137 138 184 231 244 245 246 247 267 108 186 285 297 307 308 309 403 462 522 529 550 551 552 562 563 564 615 619 625 674 675
PROTEASE
REVERSE TRANSCRIPTASE
Williams, Pennings, 2018. Data from Bacheler et al 2000

#### Viral sequences from patient 102

( likely treated with ZDV/3TC + efavirenz )

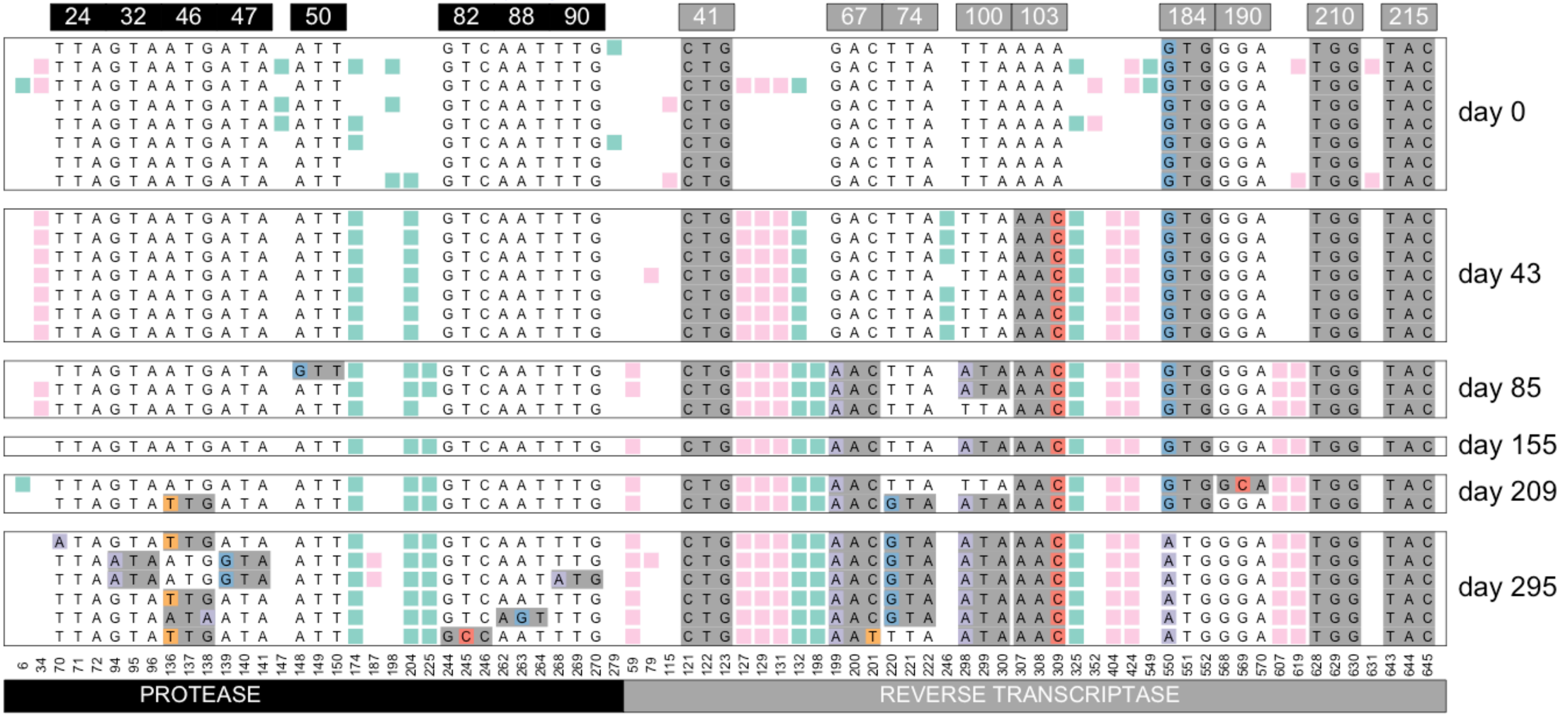


Williams, Pennings, 2018. Data from Bacheler et al 2000

**Viral sequences from patient 103**

( likely treated with ZDV/3TC + efavirenz )

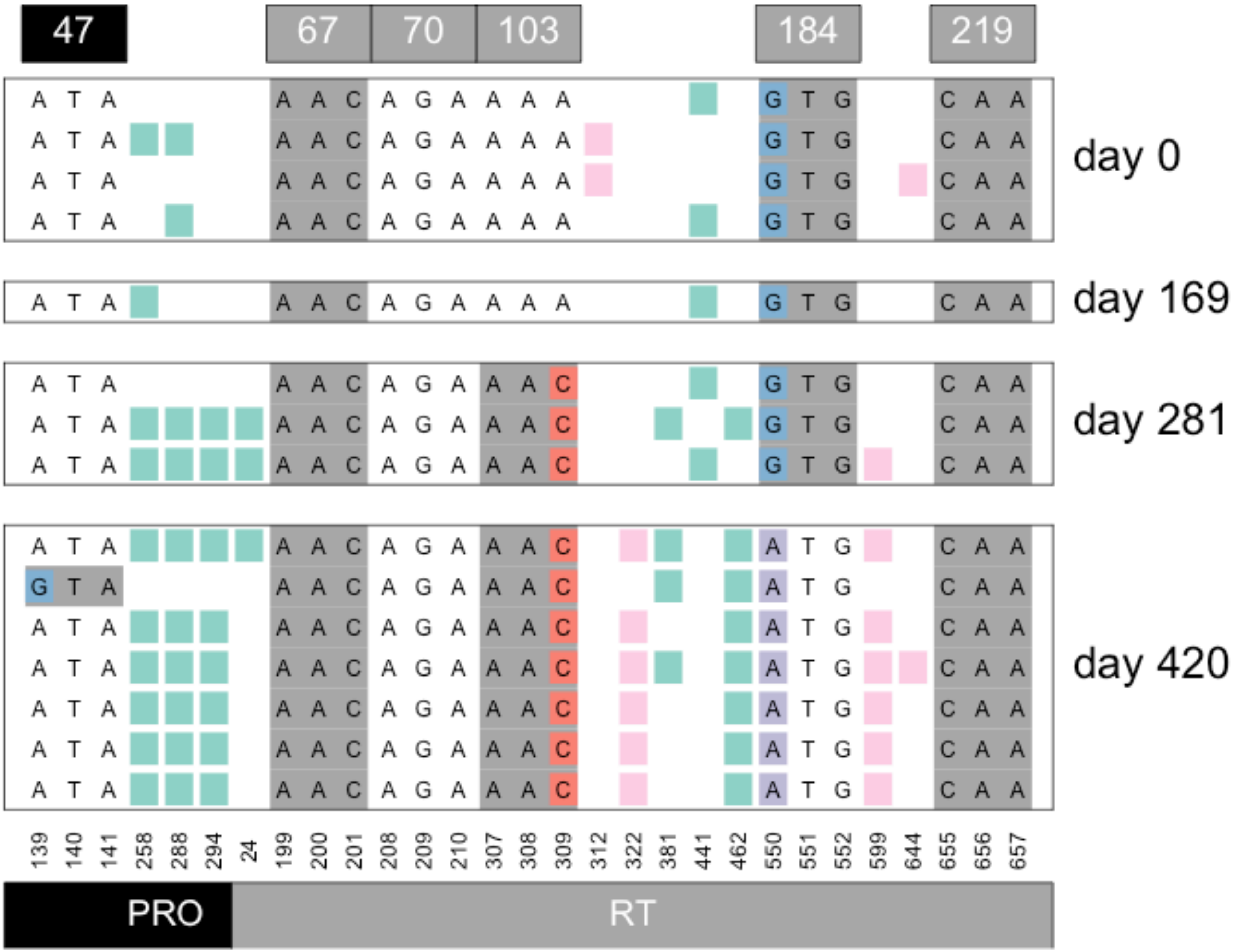


Williams, Pennings, 2018. Data from Bacheler et al 2000

### Viral sequences from patient 105

( likely treated with indinavir )

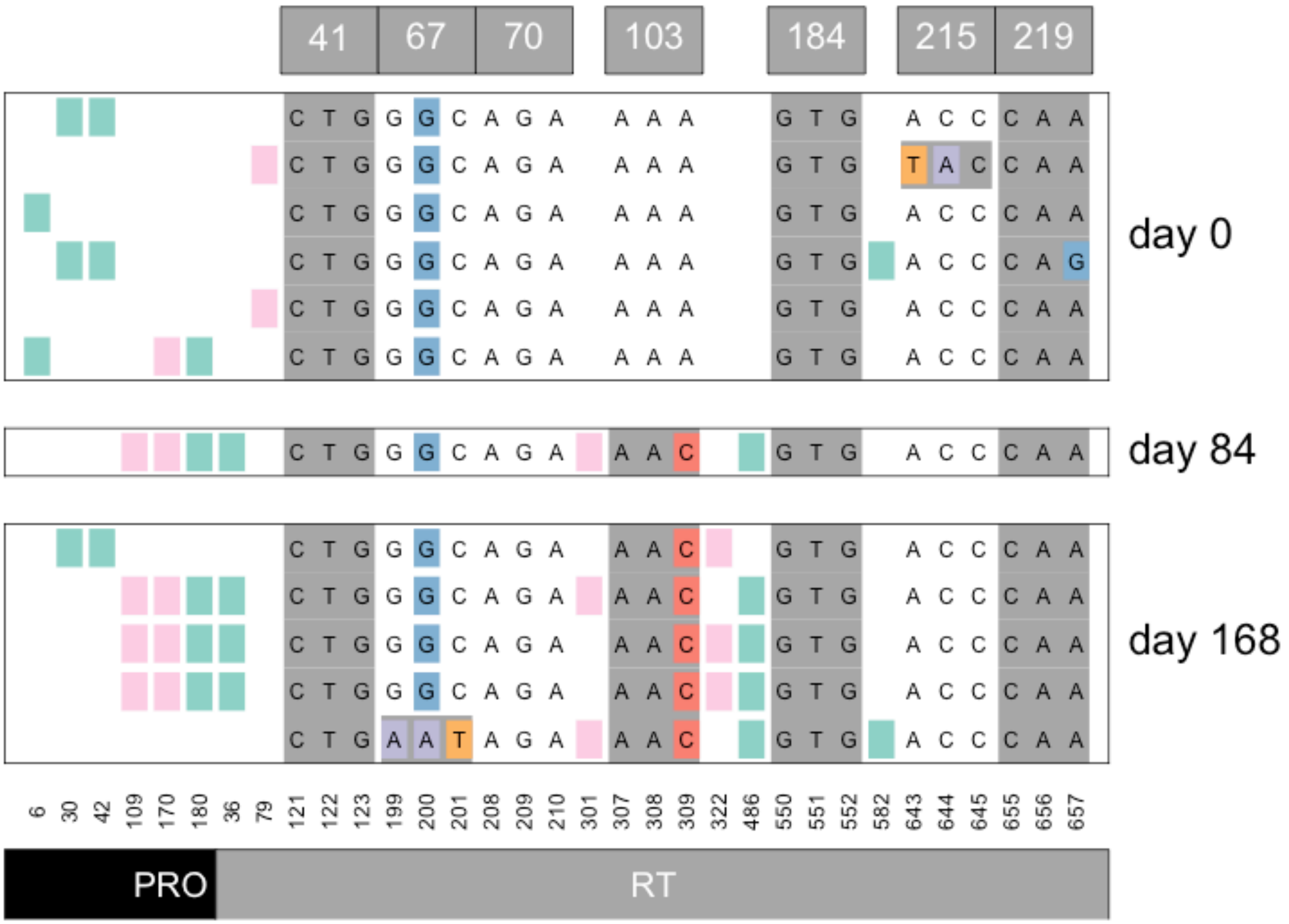


Williams, Pennings, 2018. Data from Bacheler et al 2000

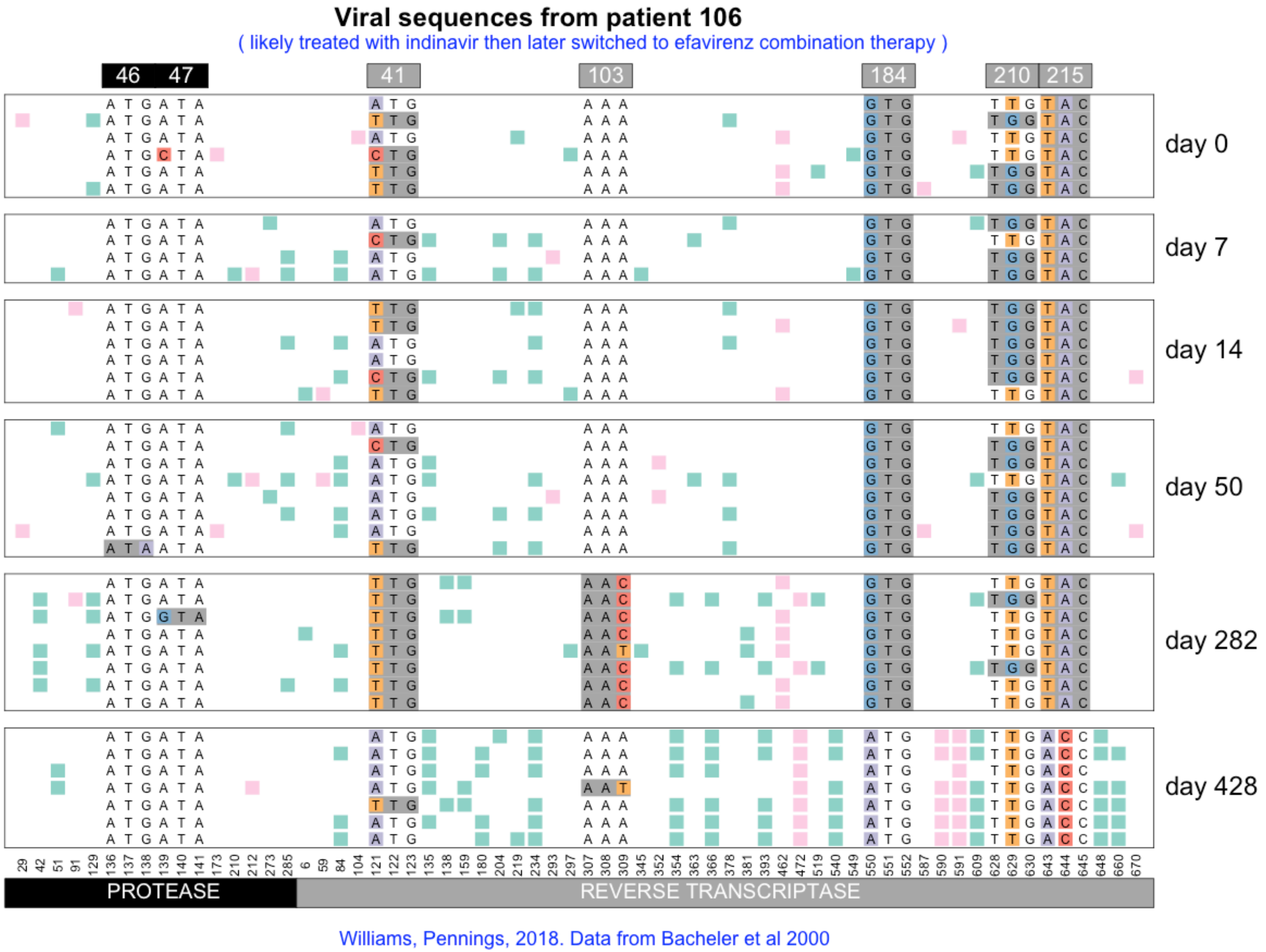
Viral sequences from patient 106
( likely treated with indinavir then later switched to efavirenz combination therapy )
46 47
41
103
184
210 215
day 0
day 7
day 14
day 50
day 282
day 428
PROTEASE
REVERSE TRANSCRIPTASE
Williams, Pennings, 2018. Data from Bacheler et al 2000

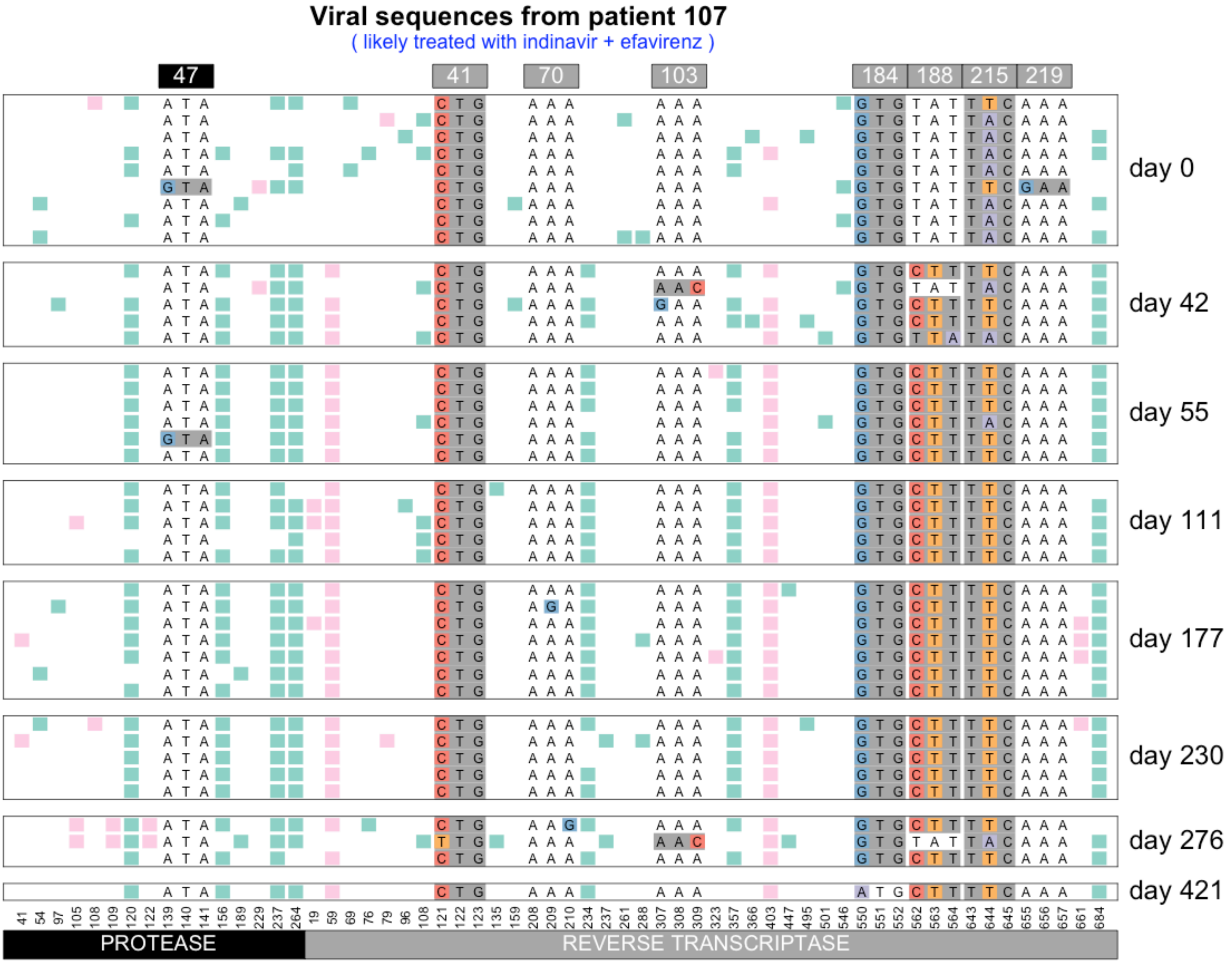


Williams, Pennings, 2018. Data from Bacheler et al 2000

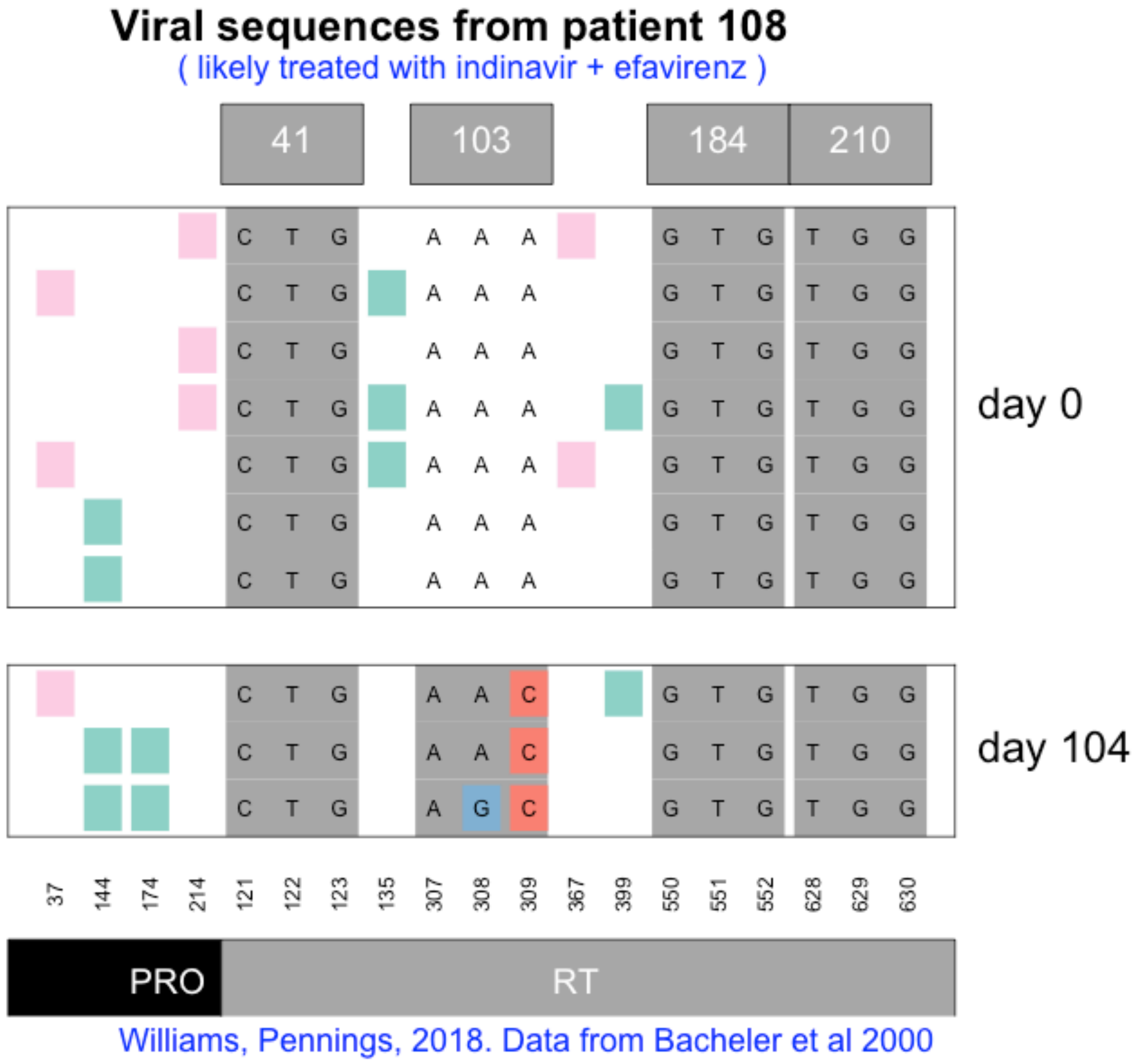

Viral sequences from patient 108
( likely treated with indinavir + efavirenz )
41
103
184
210
day 0
day 104
C T G
A A A
G T G
T G G
A A C
A G C
37
144
174
214
121
122
123
135
307
308
309
367
399
550
551
552
628
629
630
PRO
RT
Williams, Pennings, 2018. Data from Bacheler et al 2000

### Viral sequences from patient 109

( likely treated with ZDV/3TC + efavirenz )

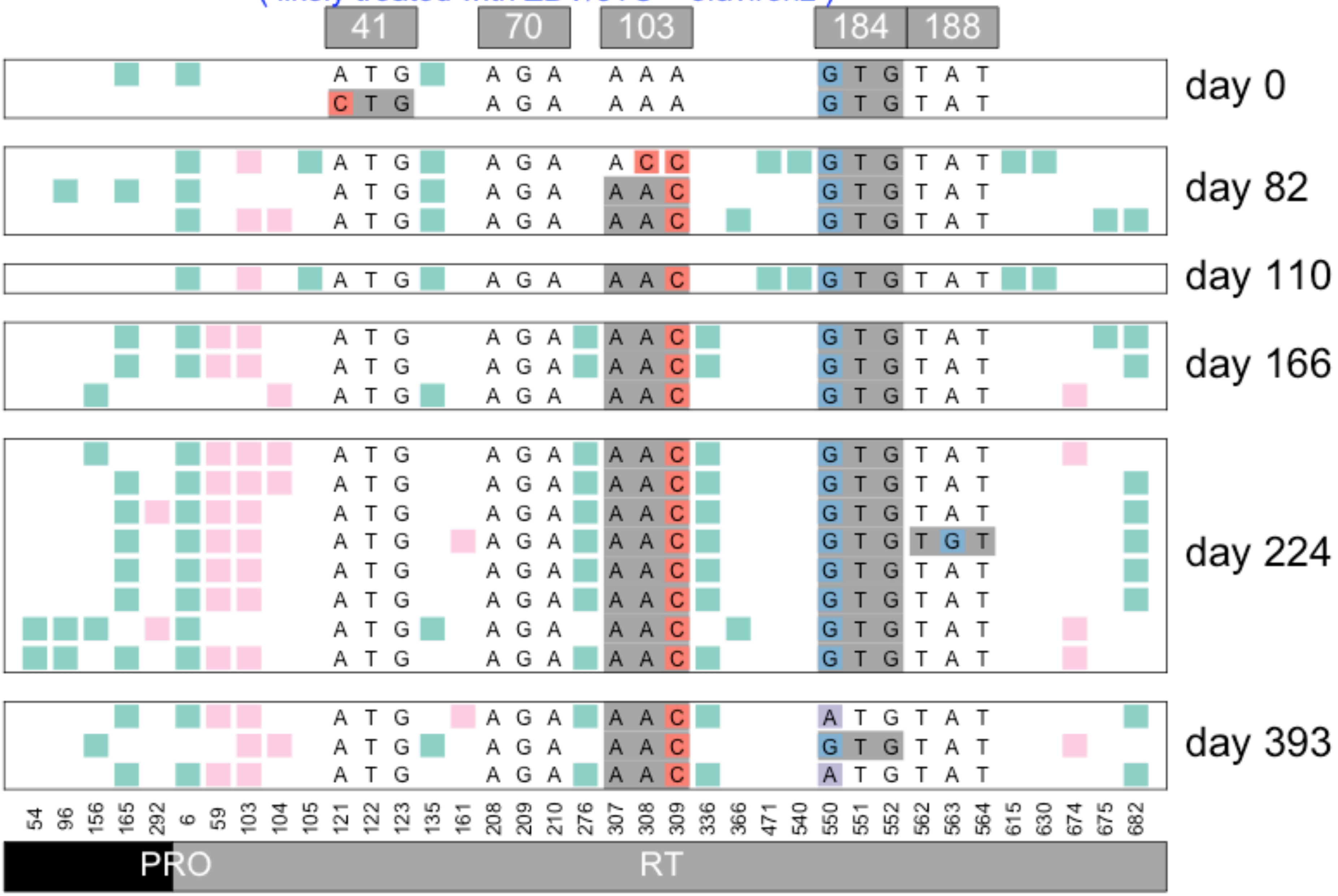


Williams, Pennings, 2018. Data from Bacheler et al 2000

#### Viral sequences from patient 110

( likely treated with indinavir then later switched to efavirenz combination therapy )

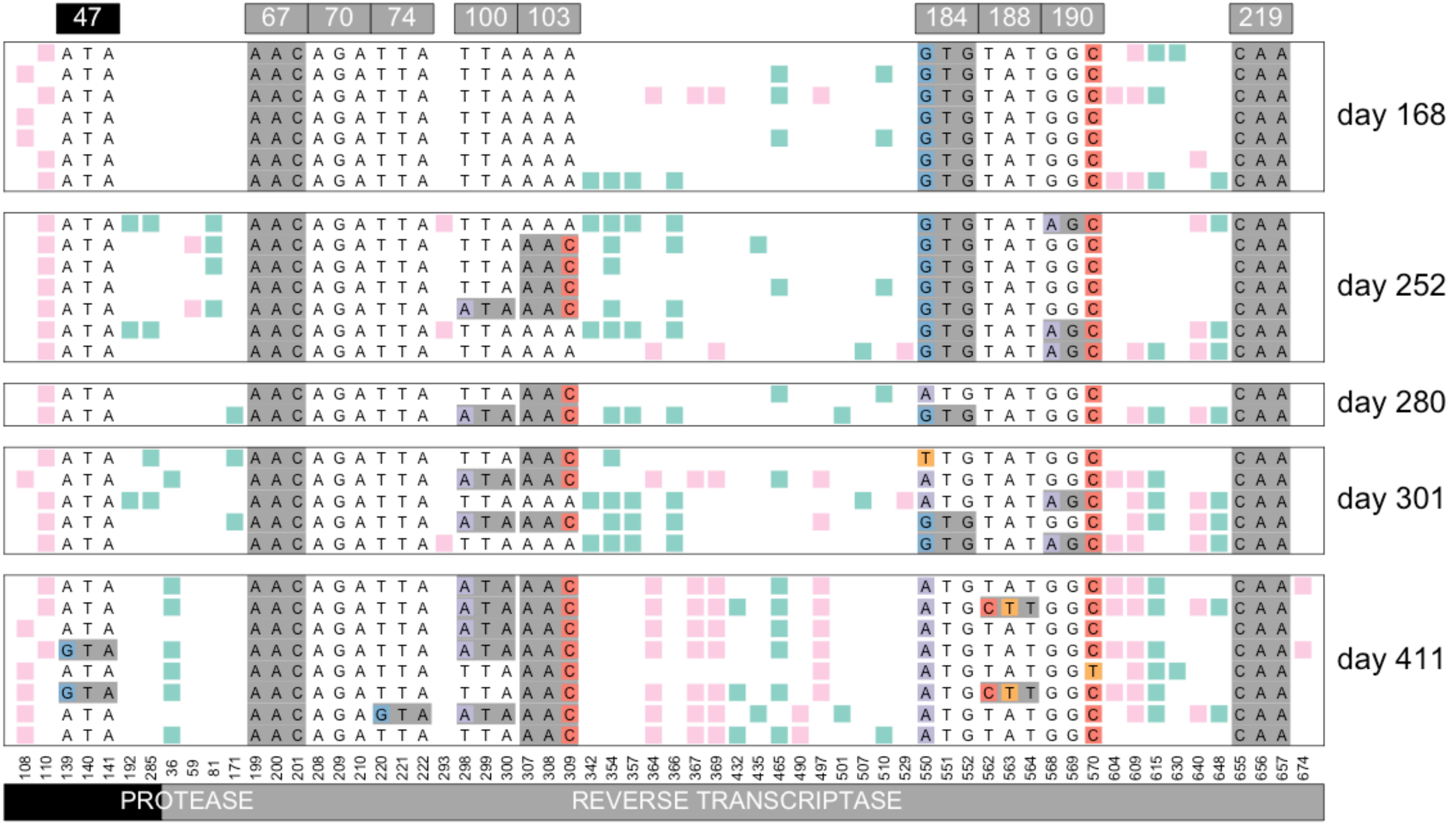


Williams, Pennings, 2018. Data from Bacheler et al 2000

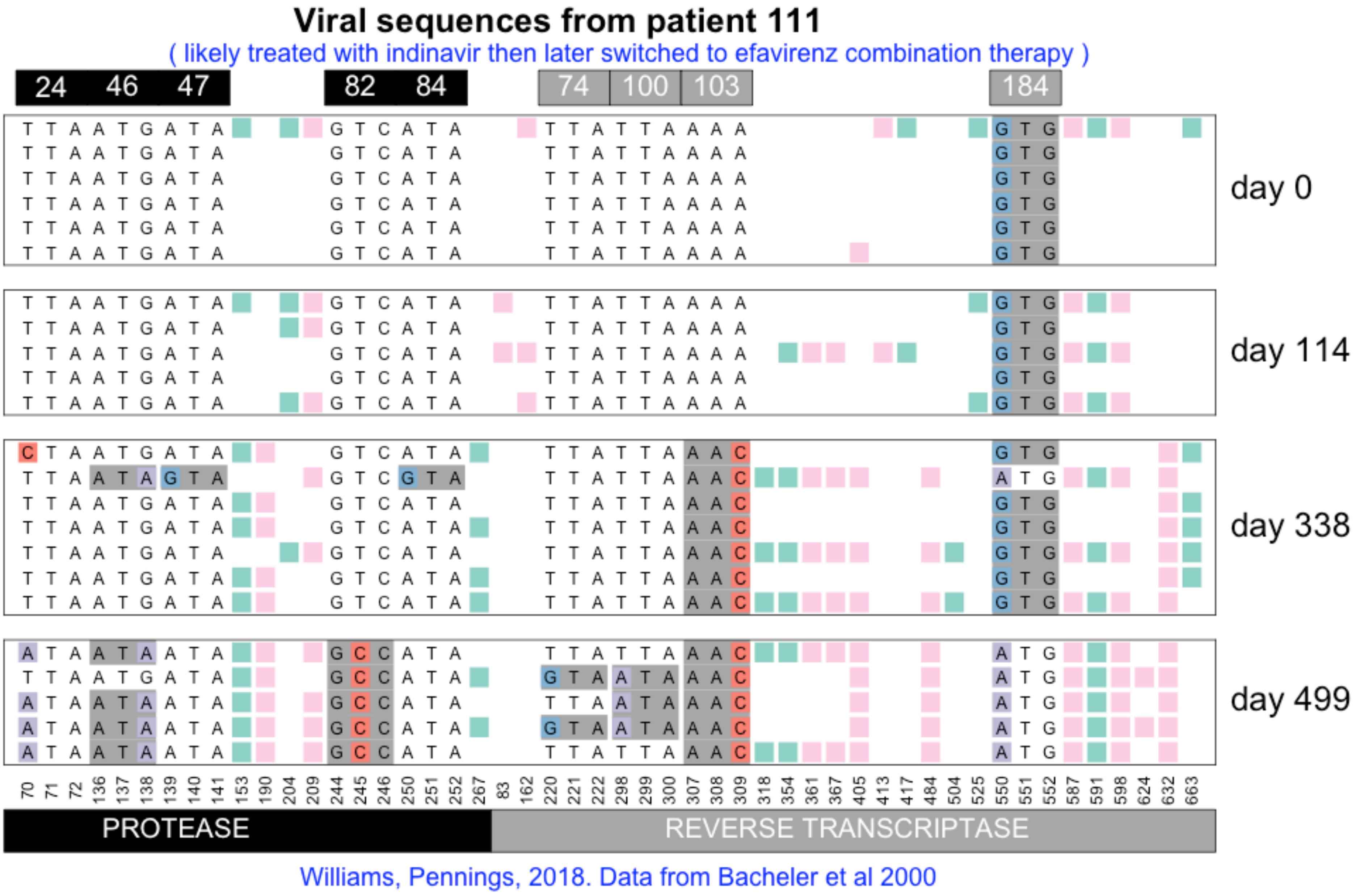

Viral sequences from patient 111
( likely treated with indinavir then later switched to efavirenz combination therapy )
24
46
47
82
84
74
100
103
184
day 0
day 114
day 338
day 499
PROTEASE
REVERSE TRANSCRIPTASE
Williams, Pennings, 2018. Data from Bacheler et al 2000

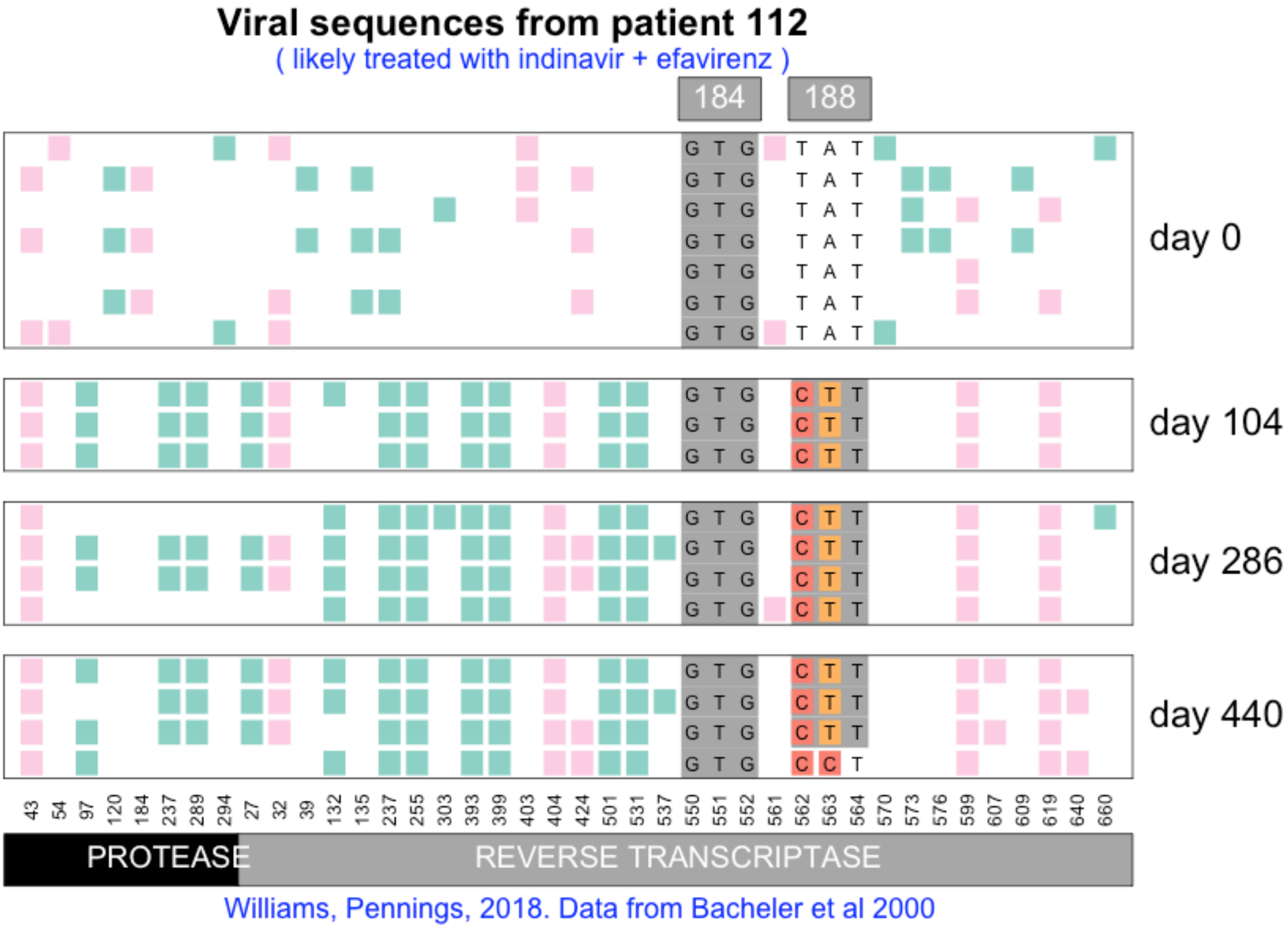

Viral sequences from patient 112
( likely treated with indinavir + efavirenz )
184
188
G T G
T A T
C T T
C C T
day 0
day 104
day 286
day 440
43 54 97 120 184 237 289 294 27 32 39 132 135 237 255 303 393 399 403 404 424 501 531 537 550 551 552 561 562 563 564 570 573 576 599 607 609 619 640 660
PROTEASE
REVERSE TRANSCRIPTASE
Williams, Pennings, 2018. Data from Bacheler et al 2000

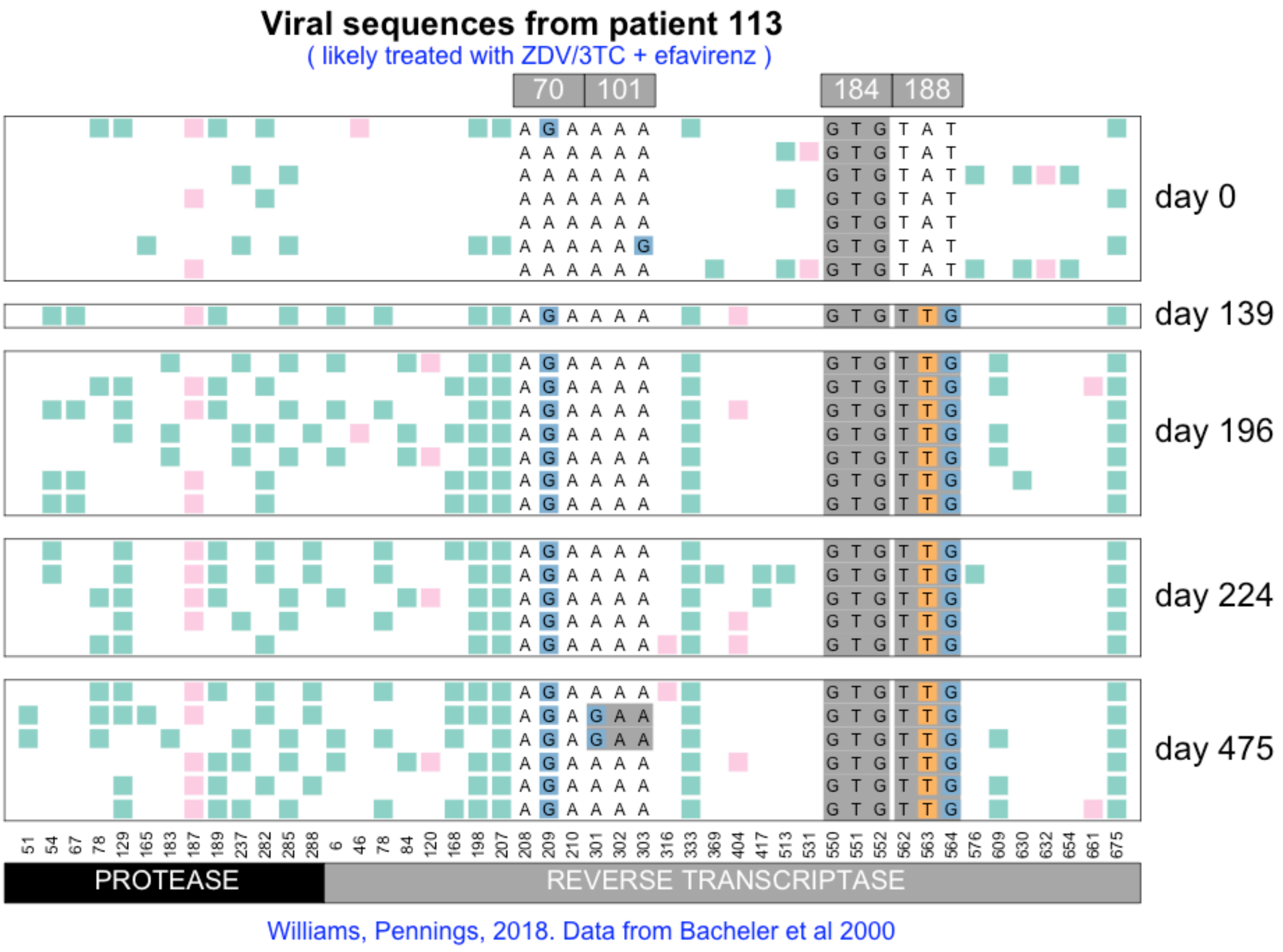
Viral sequences from patient 113
( likely treated with ZDV/3TC + efavirenz )
70
101
184
188
day 0
day 139
day 196
day 224
day 475
51 54 67 78 129 165 183 187 189 237 282 285 288 6 46 78 84 120 168 198 207 208 209 210 301 302 303 316 333 369 404 417 513 531 550 551 552 562 563 564 576 609 630 632 654 661 675
PROTEASE
REVERSE TRANSCRIPTASE
Williams, Pennings, 2018. Data from Bacheler et al 2000

### Viral sequences from patient 114

( likely treated with indinavir + efavirenz )

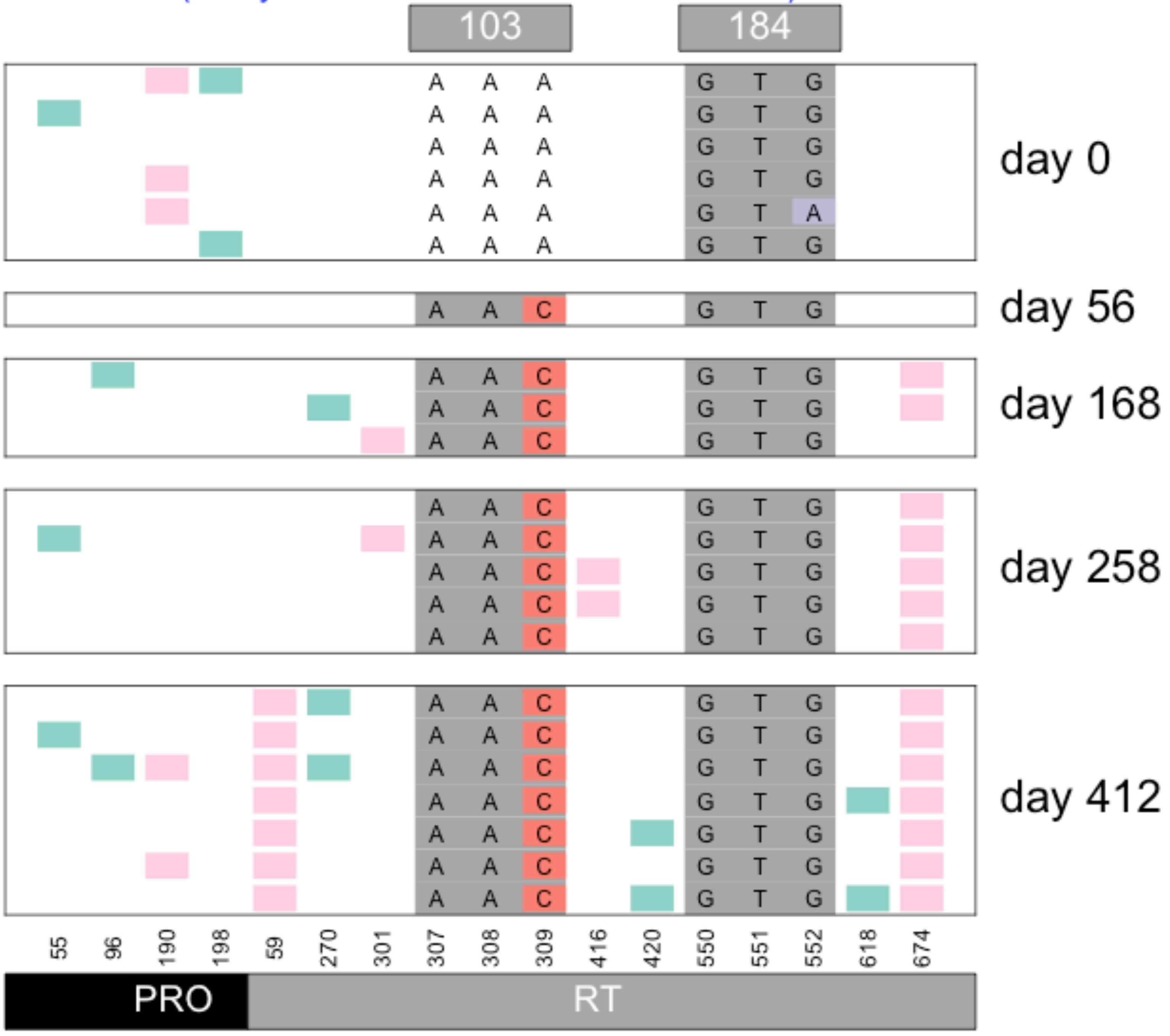


Williams, Pennings, 2018. Data from Bacheler et al 2000

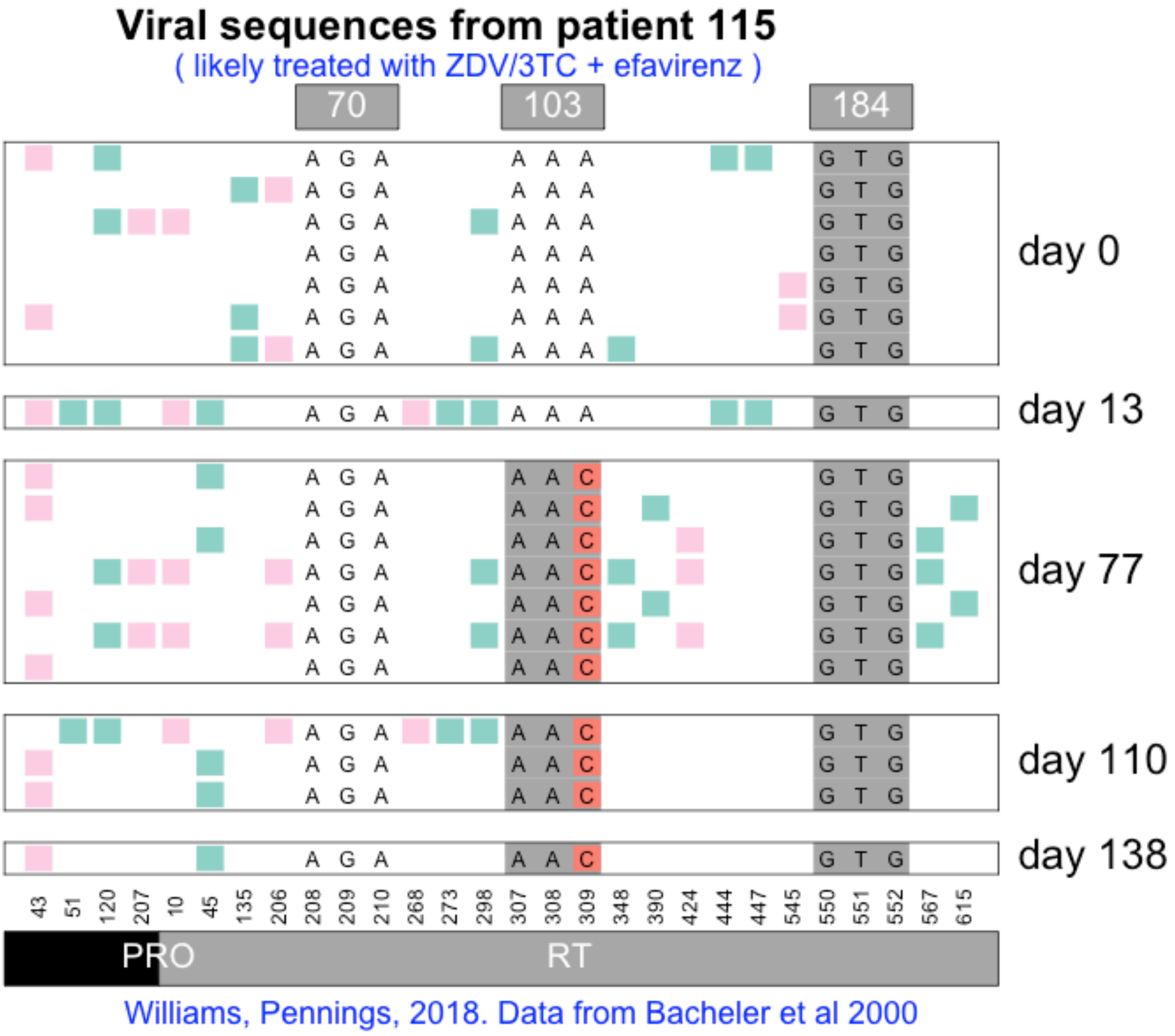
Viral sequences from patient 115
( likely treated with ZDV/3TC + efavirenz )
70
103
184
day 0
day 13
day 77
day 110
day 138
43 51 120 207 10 45 135 206 208 209 210 268 273 298 307 308 309 348 390 424 444 447 545 550 551 552 567 615
PRO
RT
Williams, Pennings, 2018. Data from Bacheler et al 2000

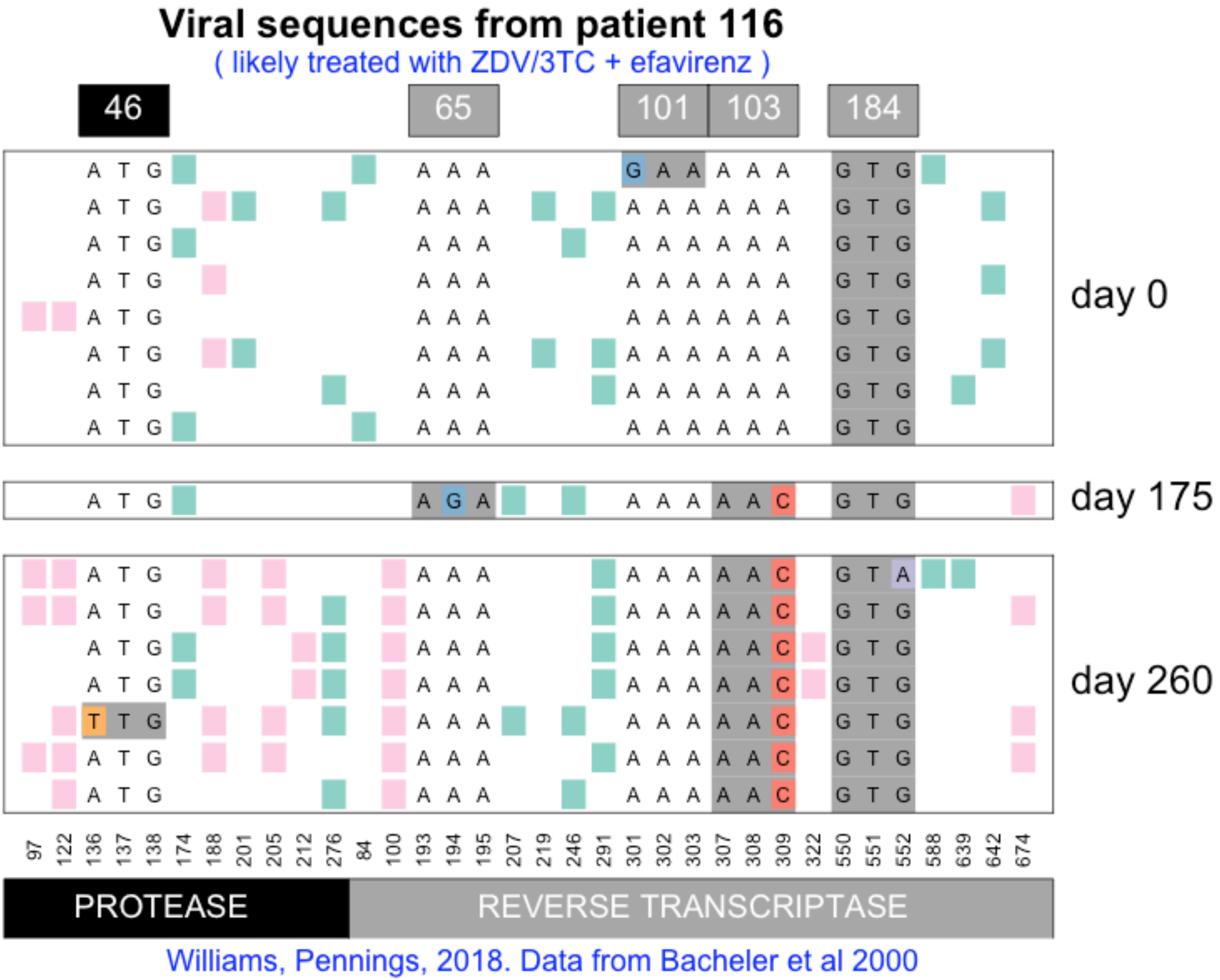
Viral sequences from patient 116
( likely treated with ZDV/3TC + efavirenz )
46
65
101
103
184
day 0
day 175
day 260
97 122 136 137 138 174 188 201 205 212 276 84 100 193 194 195 207 219 246 291 301 302 303 307 308 309 322 550 551 552 588 639 642 674
PROTEASE
REVERSE TRANSCRIPTASE
Williams, Pennings, 2018. Data from Bacheler et al 2000

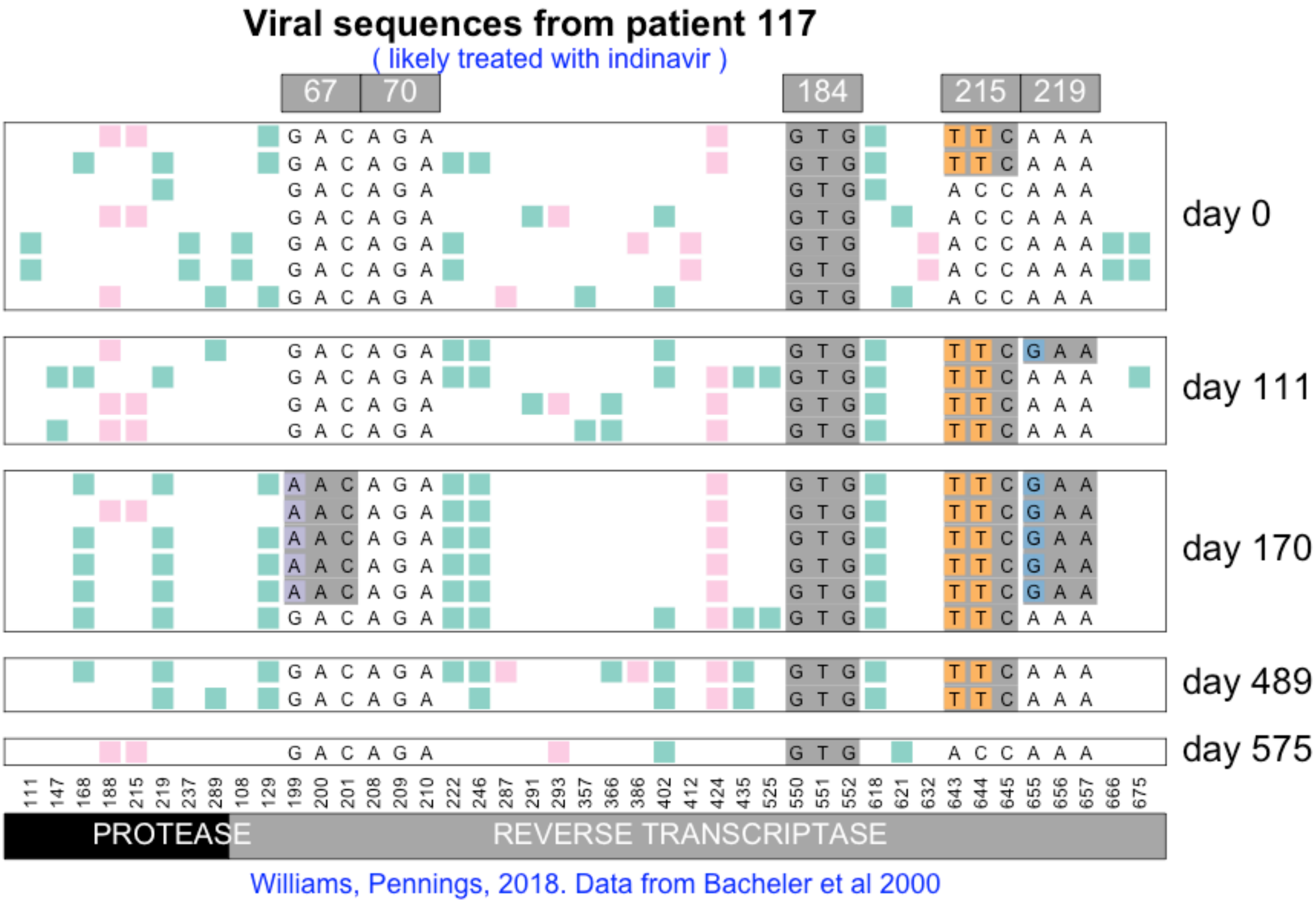
Viral sequences from patient 117
( likely treated with indinavir )
67
70
184
215
219
day 0
day 111
day 170
day 489
day 575
111 147 168 188 215 219 237 289 108 129 199 200 201 208 209 210 222 246 287 291 293 357 366 386 402 412 424 435 525 550 551 552 618 621 632 643 644 645 655 656 657 666 675
PROTEASE
REVERSE TRANSCRIPTASE
Williams, Pennings, 2018. Data from Bacheler et al 2000

### Viral sequences from patient 118

( likely treated with indinavir + efavirenz )

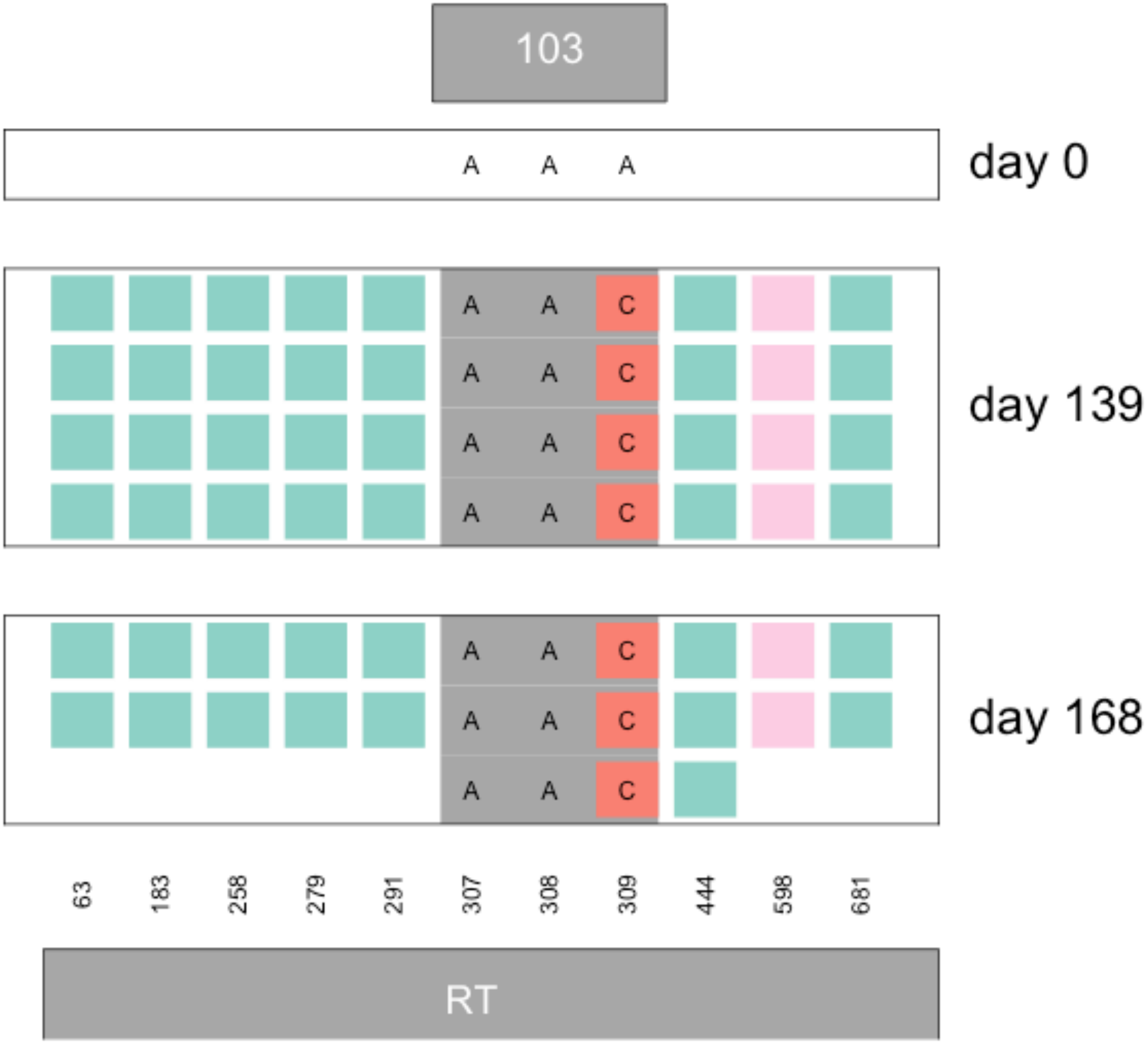


Williams, Pennings, 2018. Data from Bacheler et al 2000

#### Viral sequences from patient 119

( likely treated with indinavir + efavirenz )

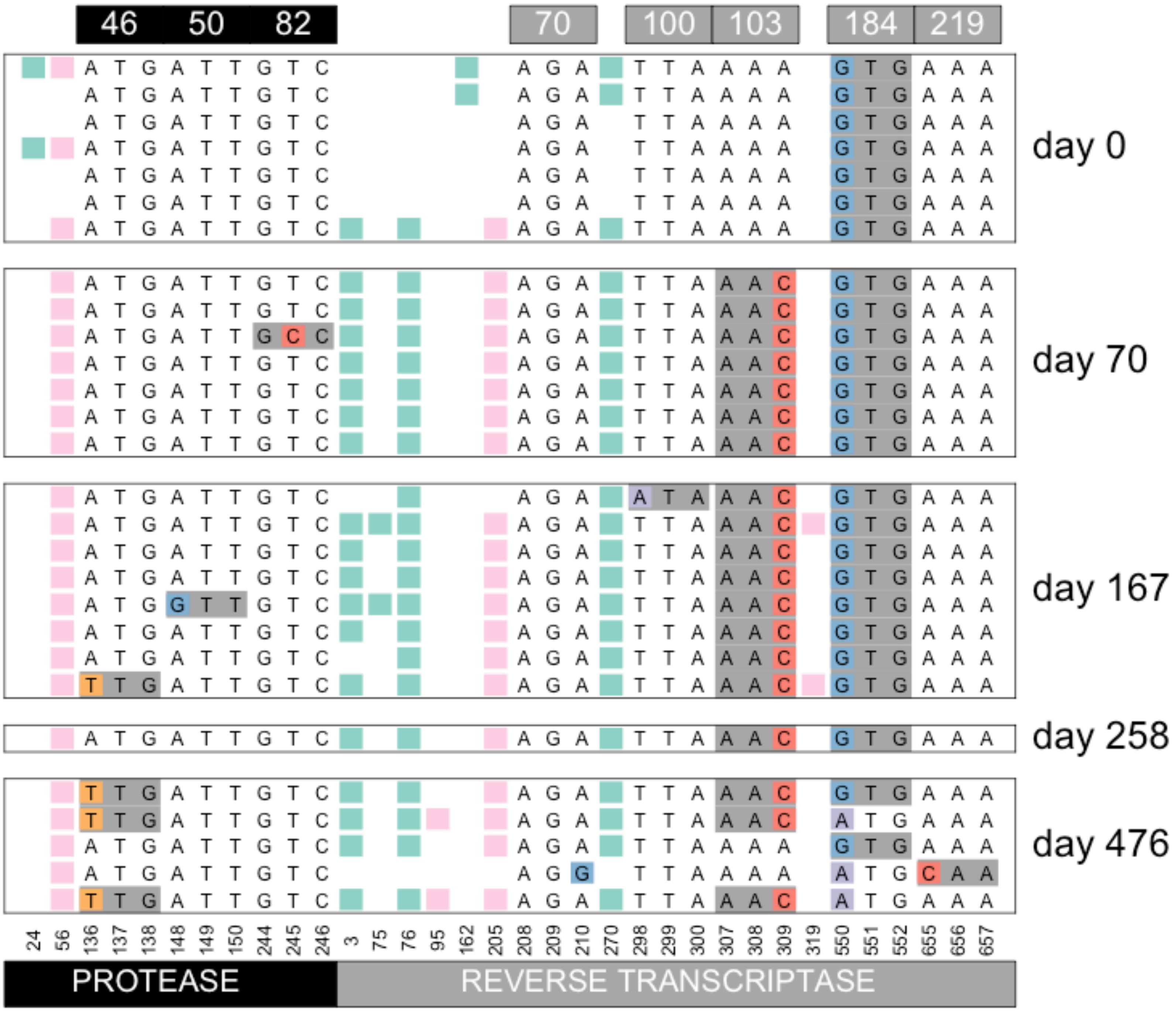


Williams, Pennings, 2018. Data from Bacheler et al 2000

### Viral sequences from patient 120

( likely treated with ZDV/3TC + efavirenz )

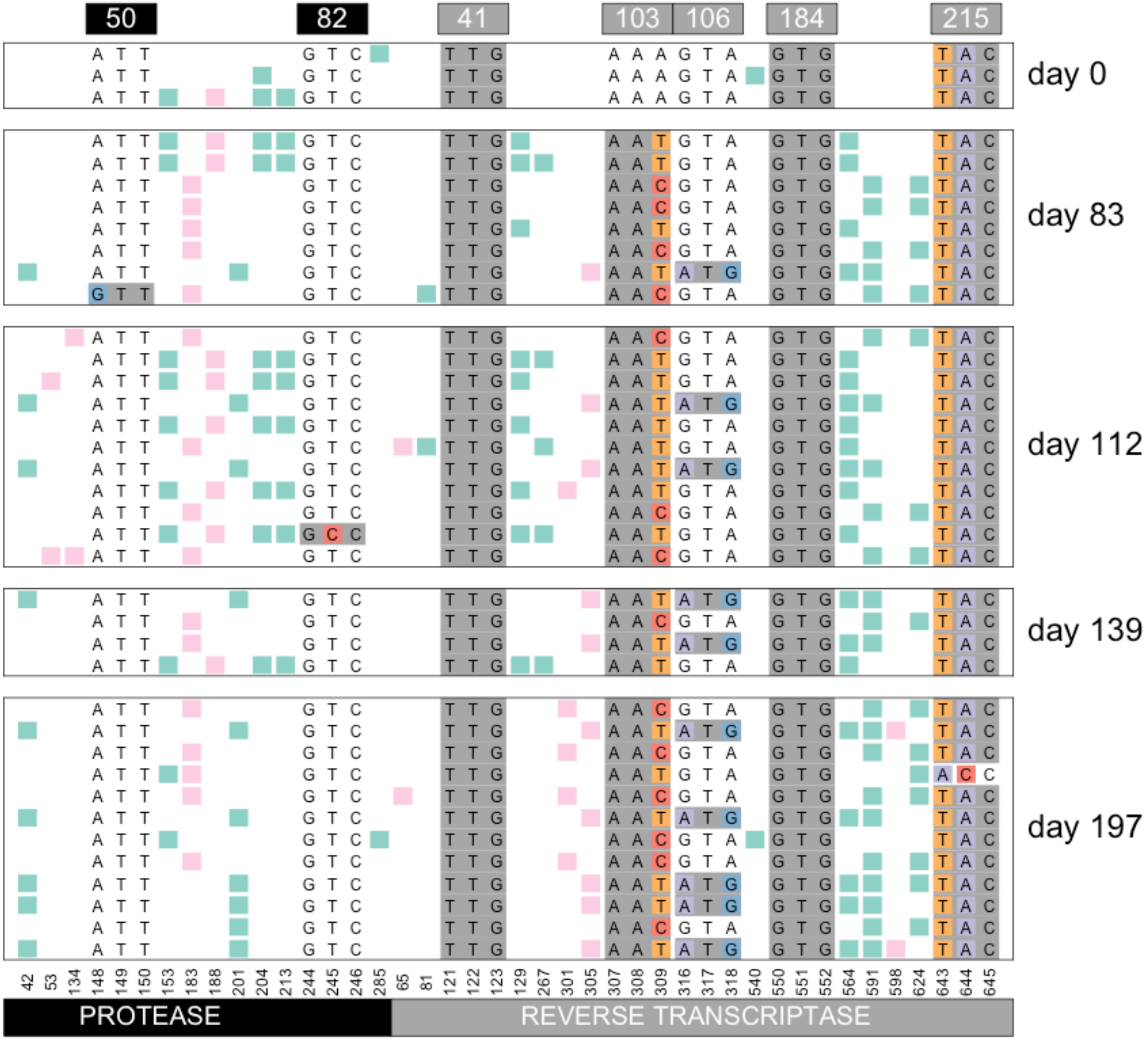


Williams, Pennings, 2018. Data from Bacheler et al 2000

#### Viral sequences from patient 121

( likely treated with indinavir + efavirenz )

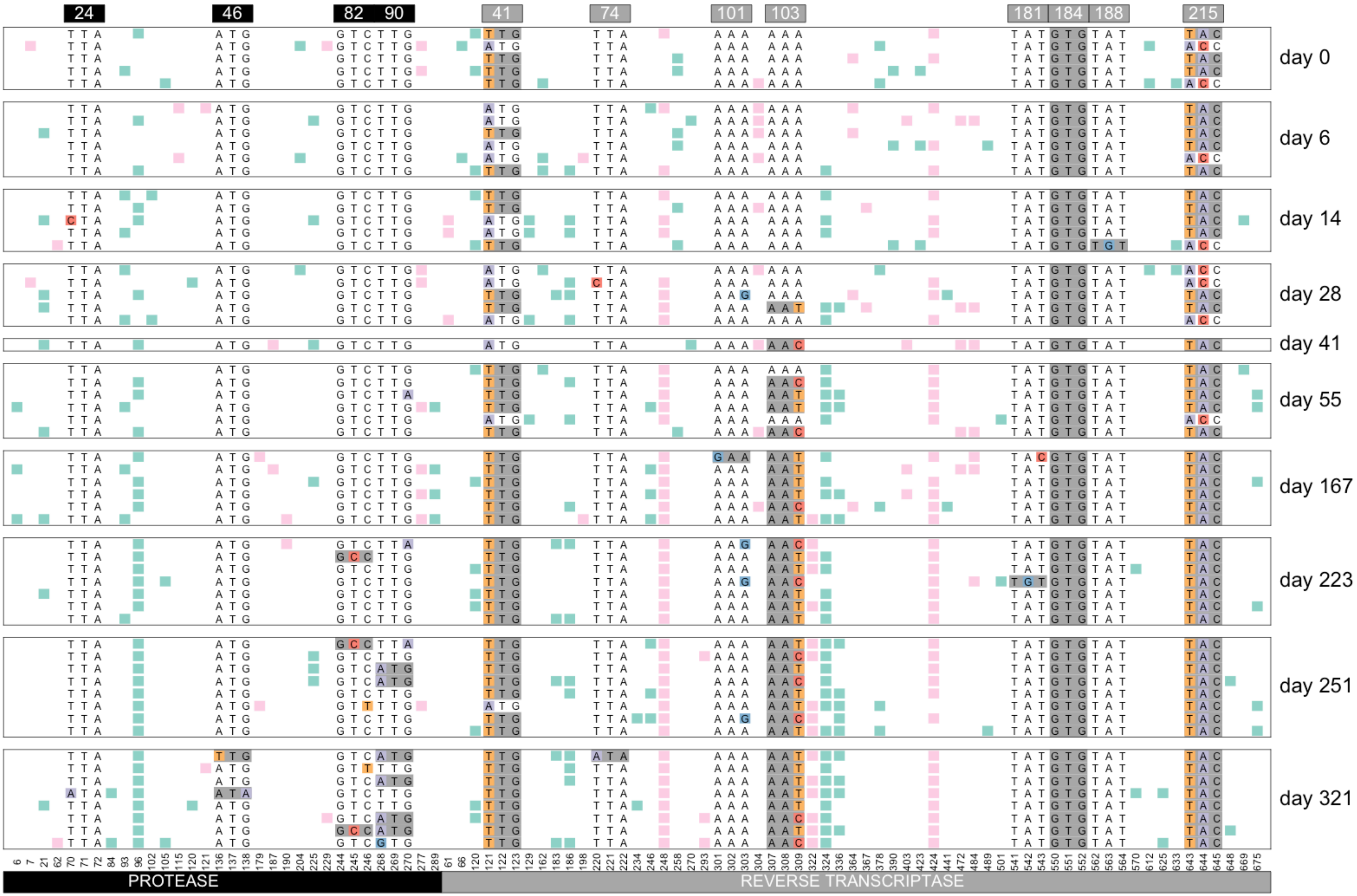


Williams, Pennings, 2018. Data from Bacheler et al 2000

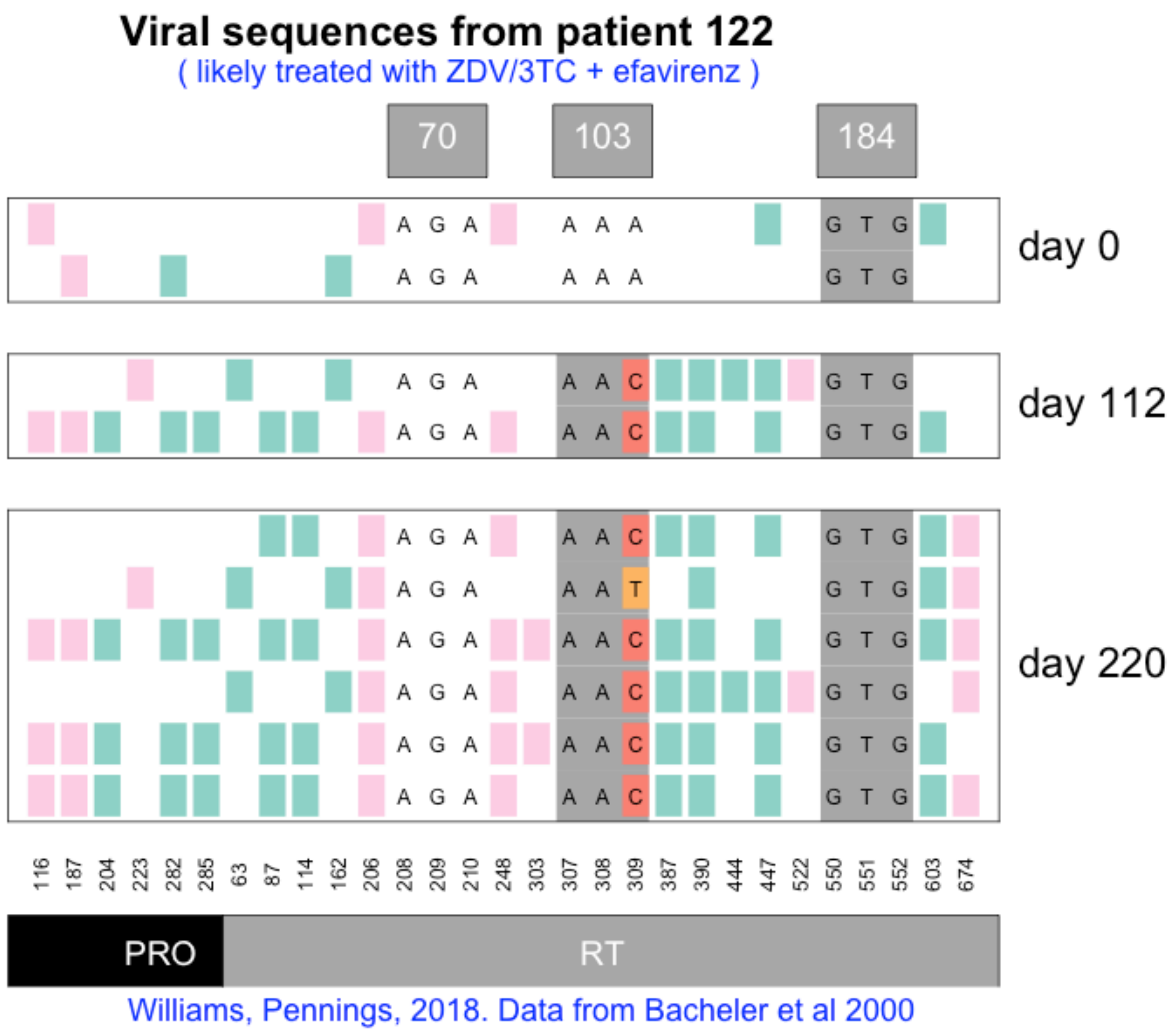
Viral sequences from patient 122
( likely treated with ZDV/3TC + efavirenz )
70
103
184
day 0
day 112
day 220
116 187 204 223 282 285 63 87 114 162 206 208 209 210 248 303 307 308 309 387 390 444 447 522 550 551 552 603 674
PRO
RT
Williams, Pennings, 2018. Data from Bacheler et al 2000

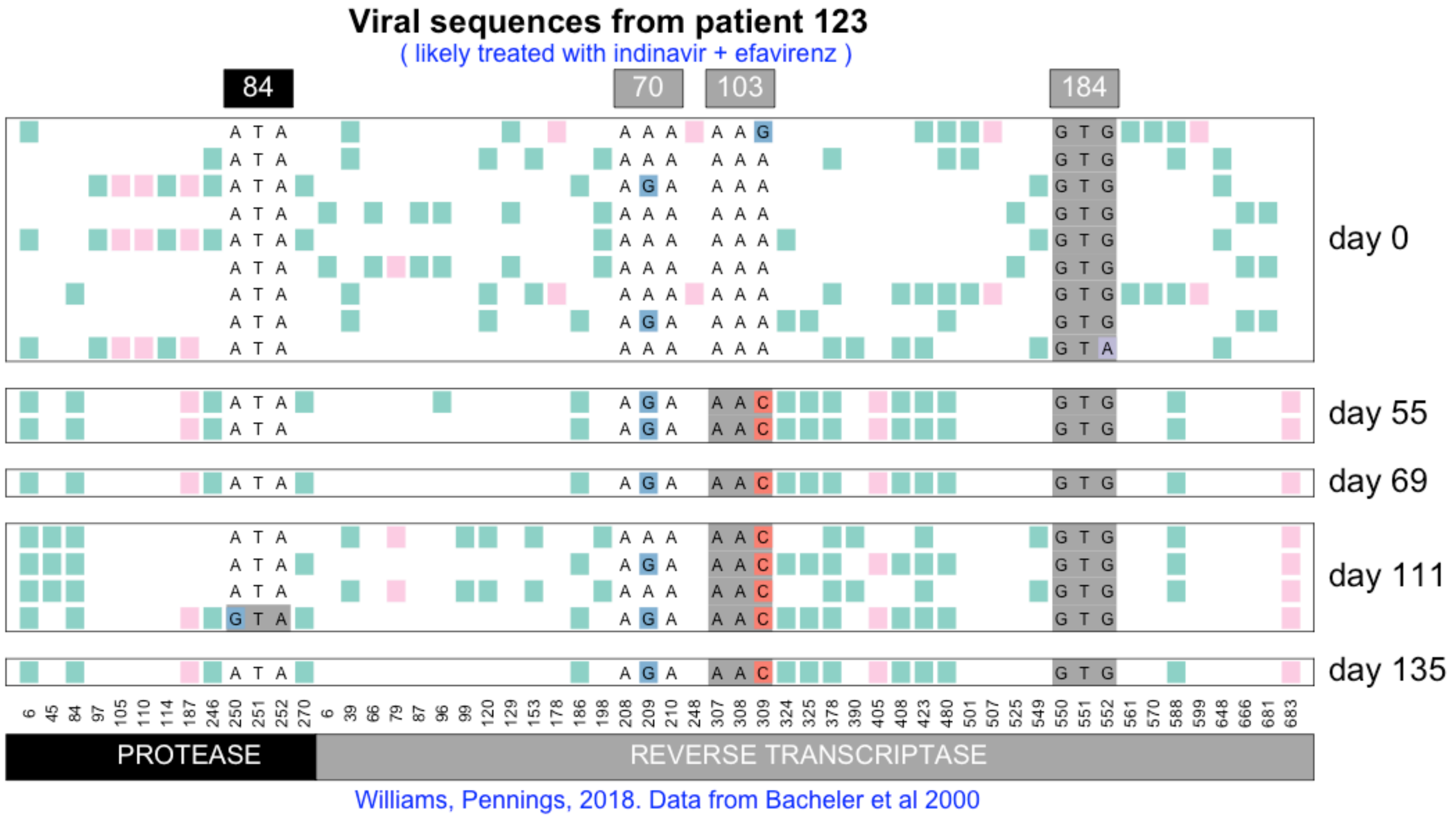
Viral sequences from patient 123
( likely treated with indinavir + efavirenz )
84
70
103
184
day 0
day 55
day 69
day 111
day 135
PROTEASE
REVERSE TRANSCRIPTASE
Williams, Pennings, 2018. Data from Bacheler et al 2000

**Viral sequences from patient 124**

( likely treated with indinavir then later switched to efavirenz combination therapy )

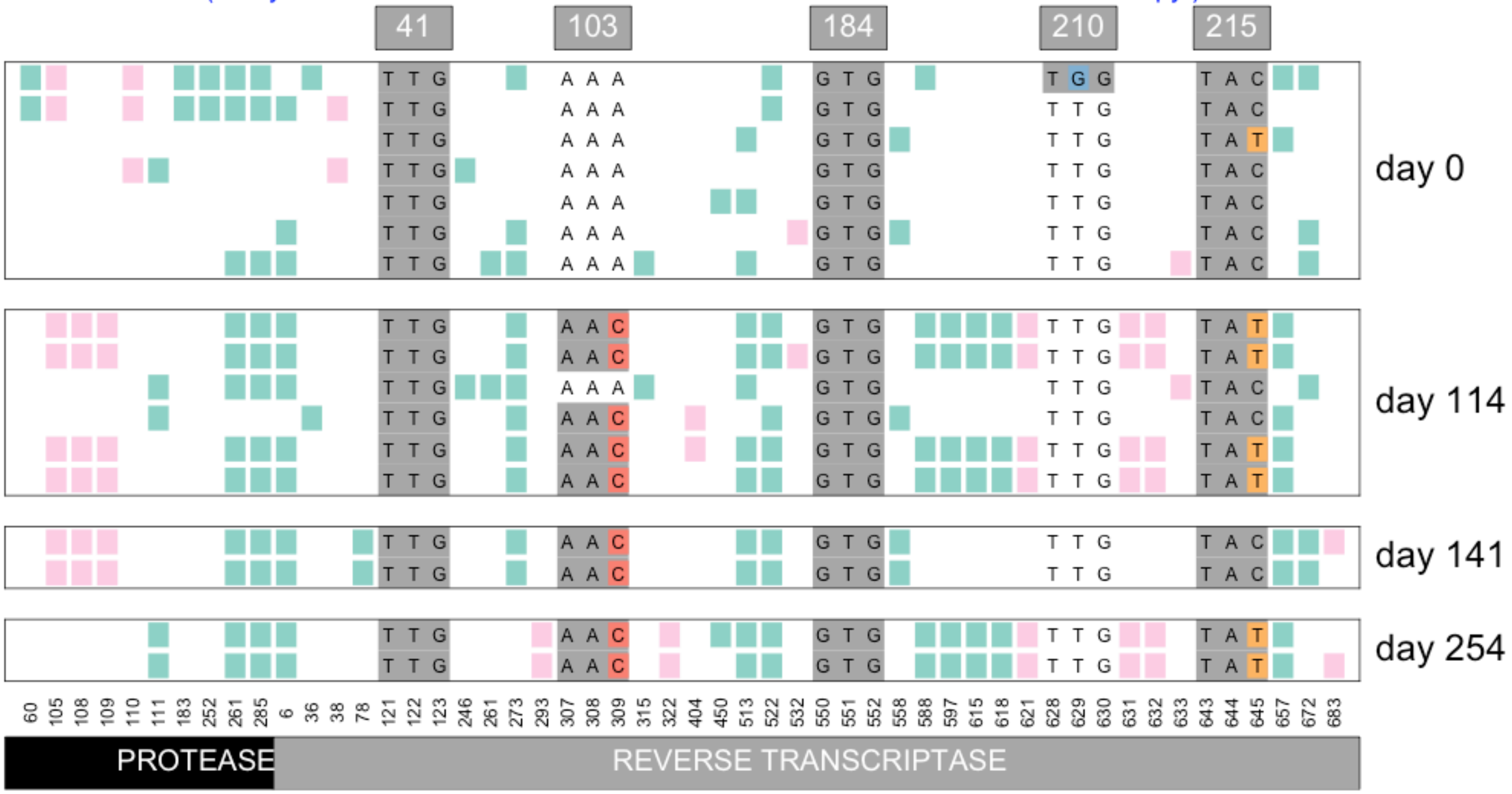


Williams, Pennings, 2018. Data from Bacheler et al 2000

#### Viral sequences from patient 126

( likely treated with indinavir + efavirenz )

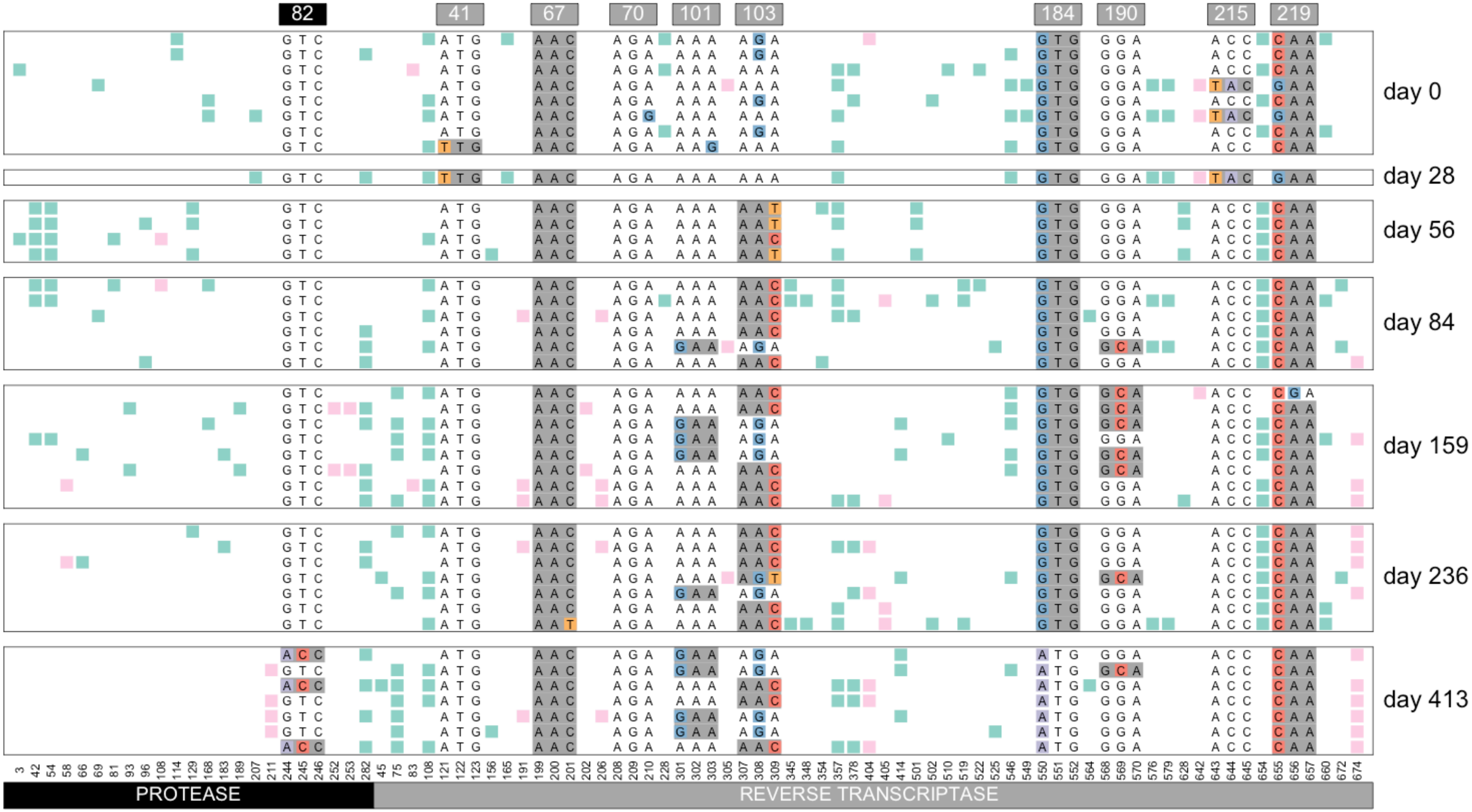


Williams, Pennings, 2018. Data from Bacheler et al 2000

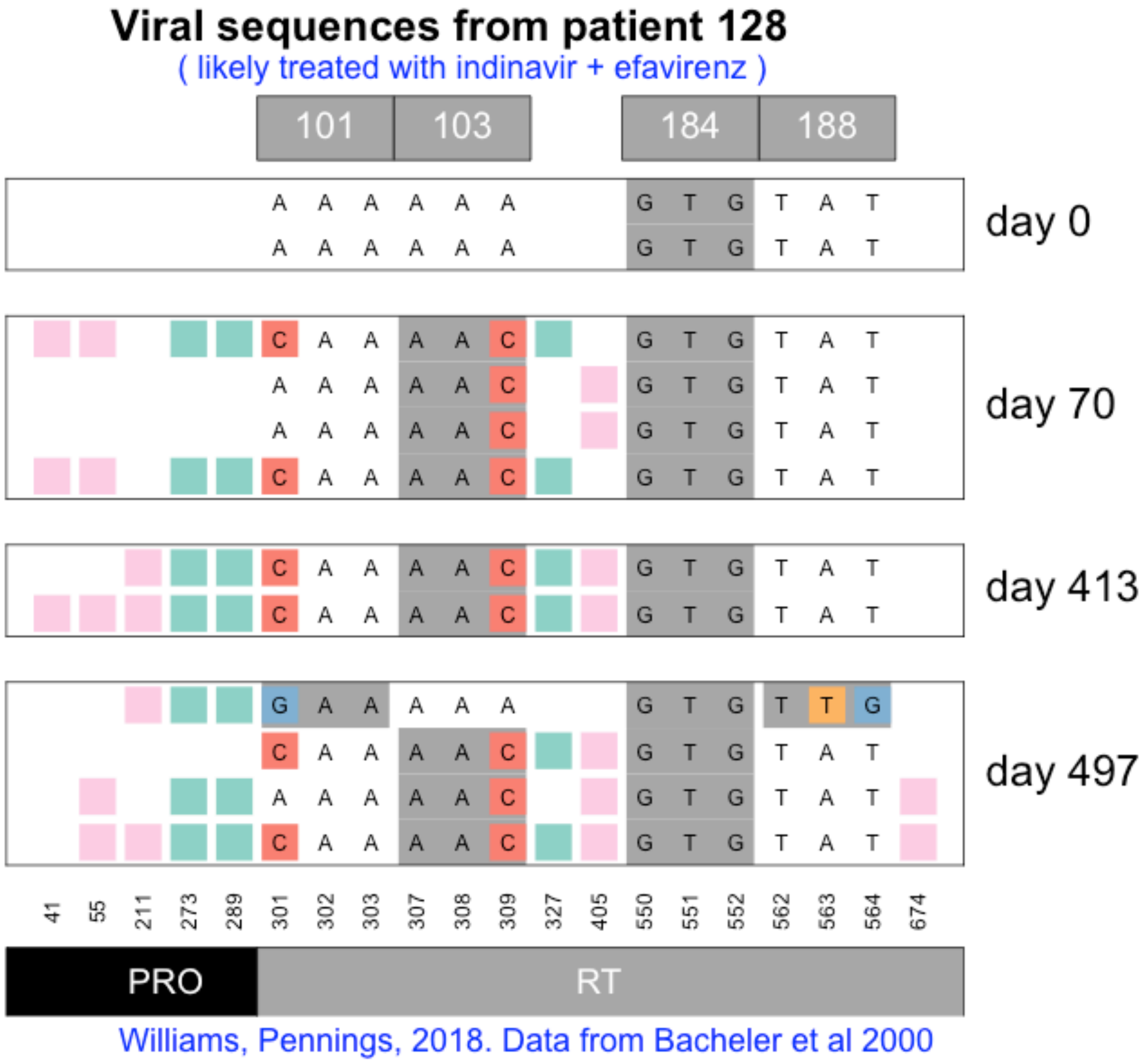

Viral sequences from patient 128
( likely treated with indinavir + efavirenz )
101
103
184
188
A A A A A A G T G T A T
A A A A A A G T G T A T
day 0
C A A A A C G T G T A T
A A A A A C G T G T A T
A A A A A C G T G T A T
C A A A A C G T G T A T
day 70
C A A A A C G T G T A T
C A A A A C G T G T A T
day 413
G A A A A A G T G T T G
C A A A A C G T G T A T
A A A A A C G T G T A T
C A A A A C G T G T A T
day 497
41 55 211 273 289 301 302 303 307 308 309 327 405 550 551 552 562 563 564 674
PRO
RT
Williams, Pennings, 2018. Data from Bacheler et al 2000

#### Viral sequences from patient 130

( likely treated with indinavir then later switched to efavirenz combination therapy )

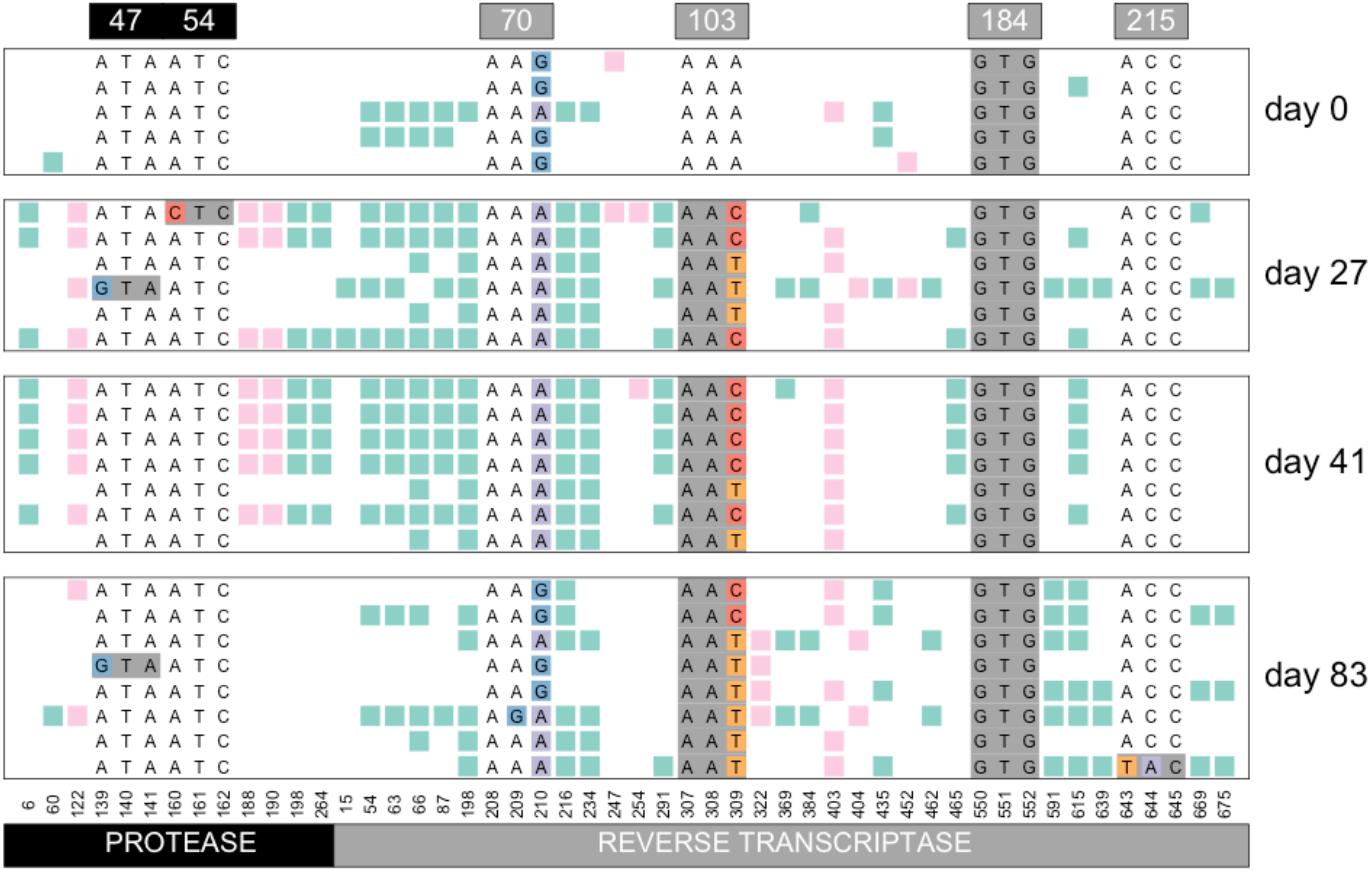


Williams, Pennings, 2018. Data from Bacheler et al 2000

#### Viral sequences from patient 131

( likely treated with indinavir + efavirenz )

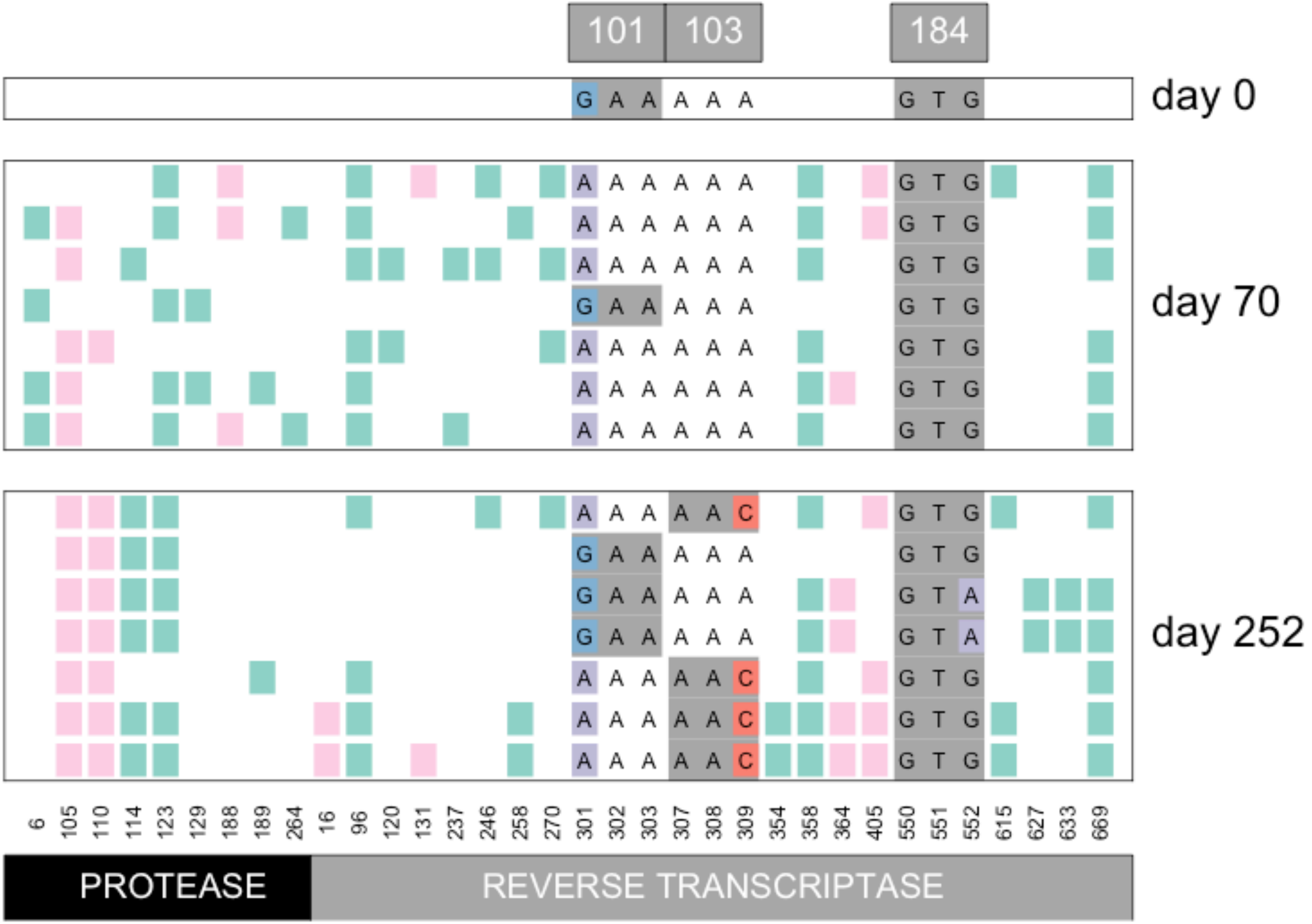


Williams, Pennings, 2018. Data from Bacheler et al 2000

#### Viral sequences from patient 132

( likely treated with indinavir + efavirenz )

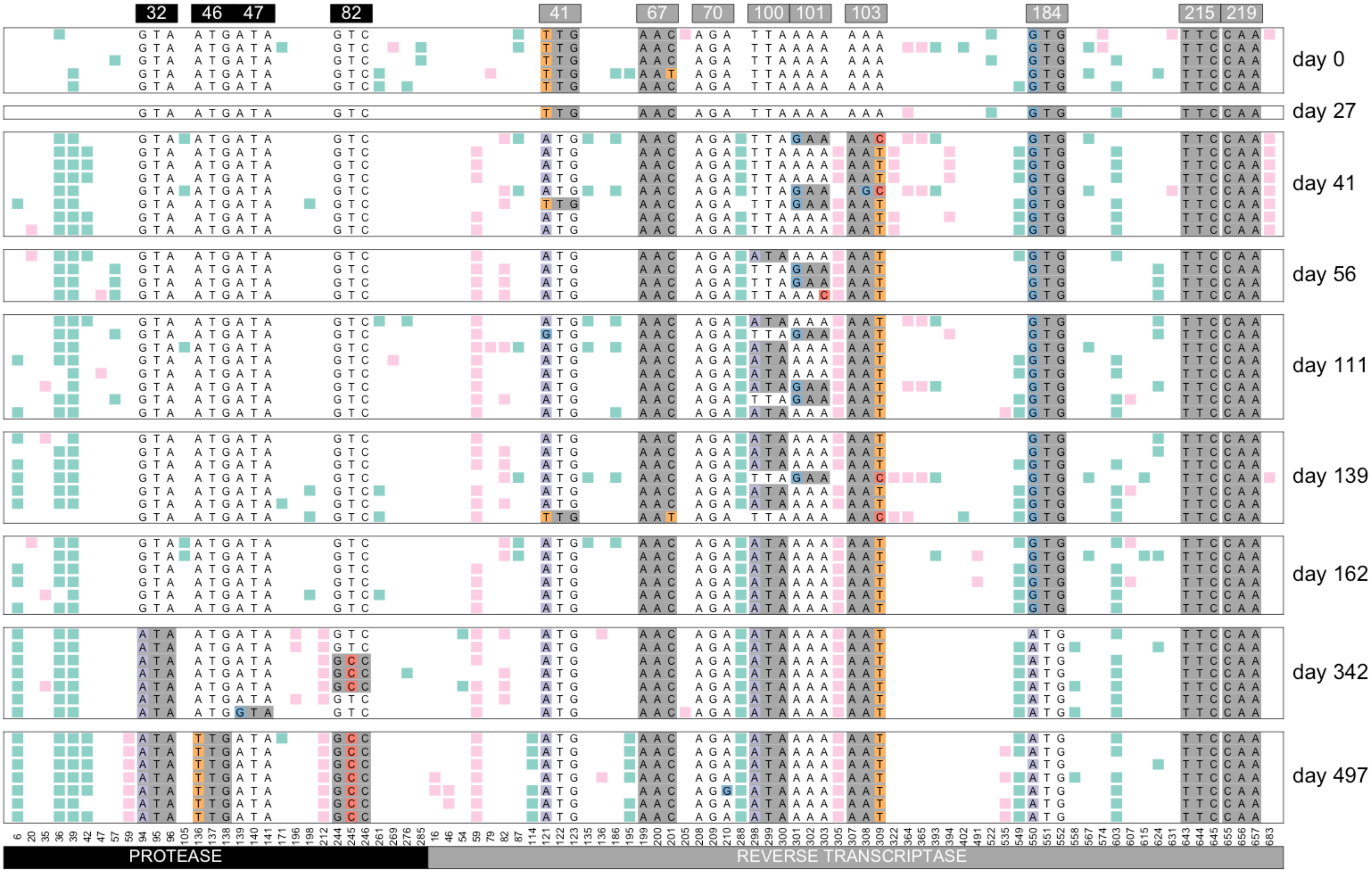


Williams, Pennings, 2018. Data from Bacheler et al 2000

### Viral sequences from patient 133

( likely treated with indinavir + efavirenz )

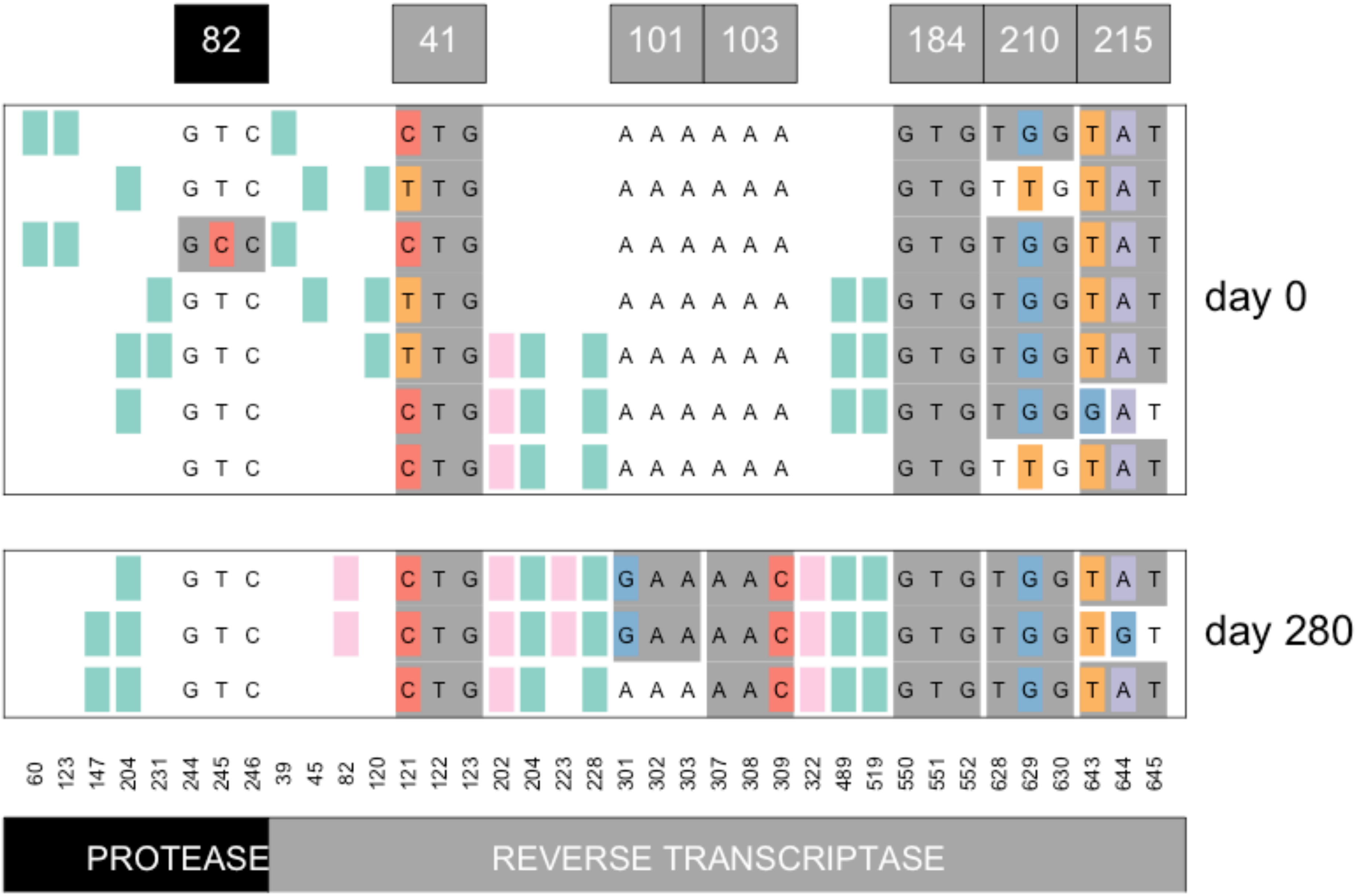


Williams, Pennings, 2018. Data from Bacheler et al 2000

### Viral sequences from patient 134

( likely treated with indinavir + efavirenz )

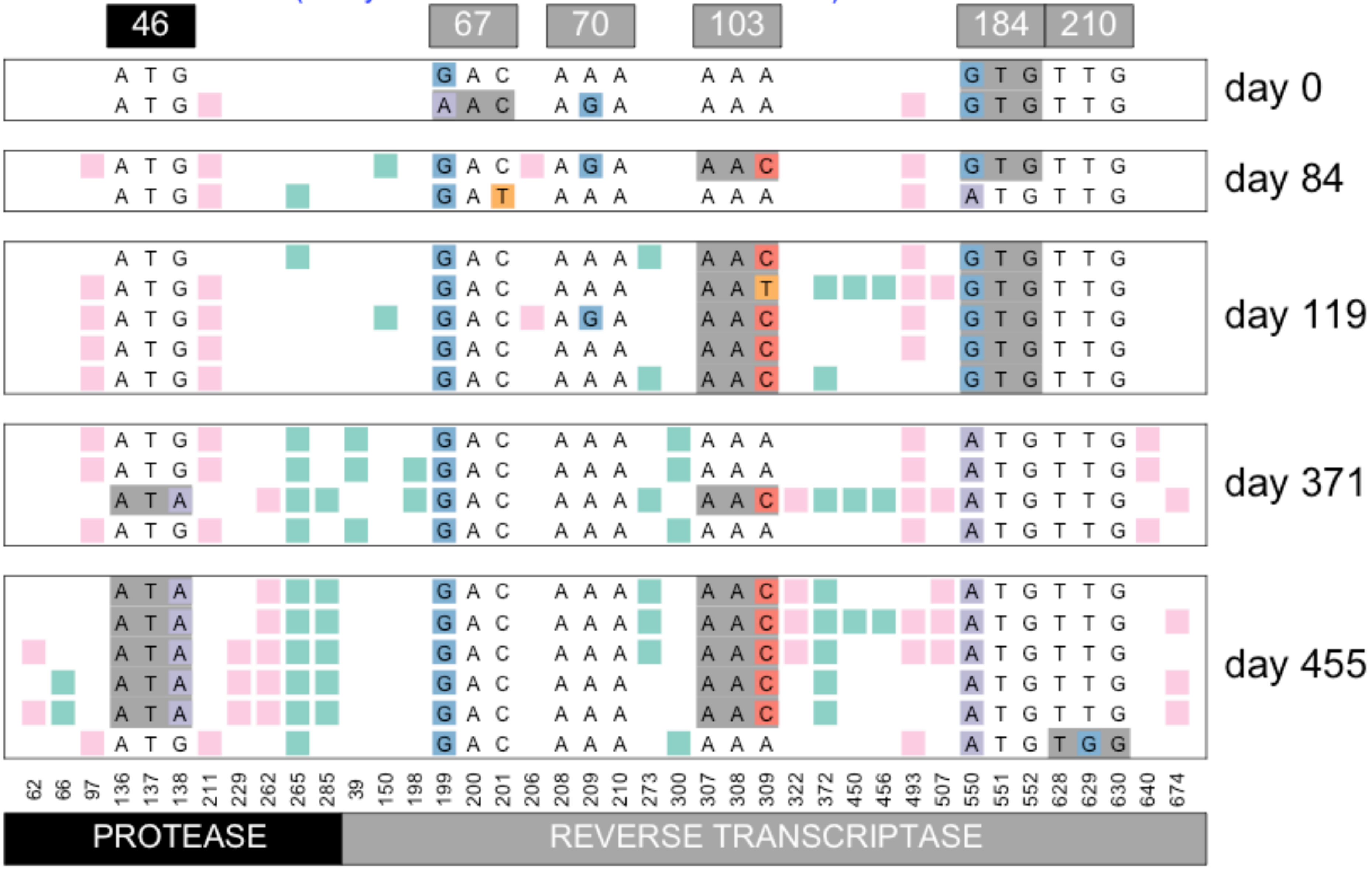


Williams, Pennings, 2018. Data from Bacheler et al 2000

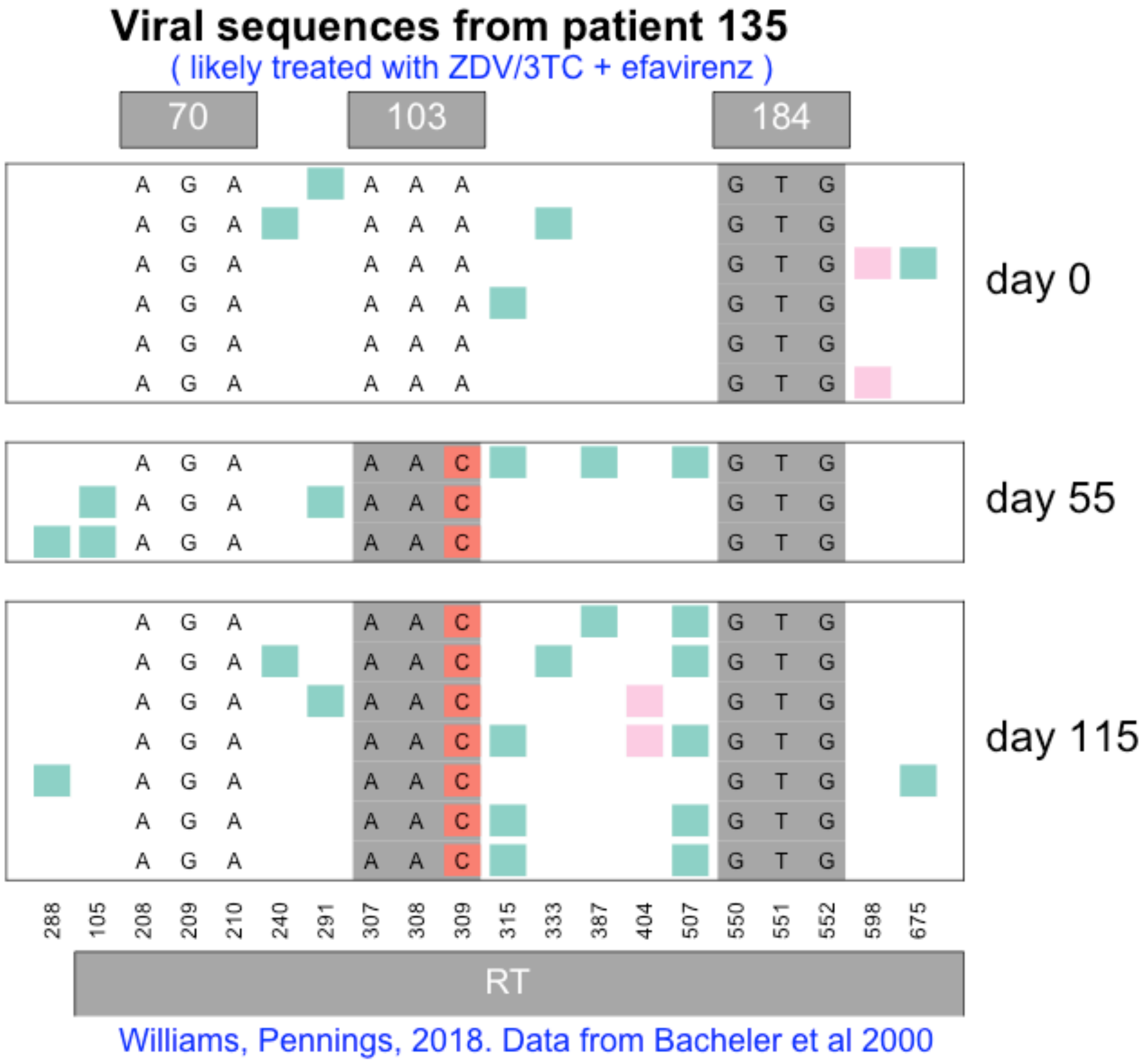

Viral sequences from patient 135
( likely treated with ZDV/3TC + efavirenz )
70
103
184
day 0
day 55
day 115
288 105 208 209 210 240 291 307 308 309 315 333 387 404 507 550 551 552 598 675
RT
Williams, Pennings, 2018. Data from Bacheler et al 2000

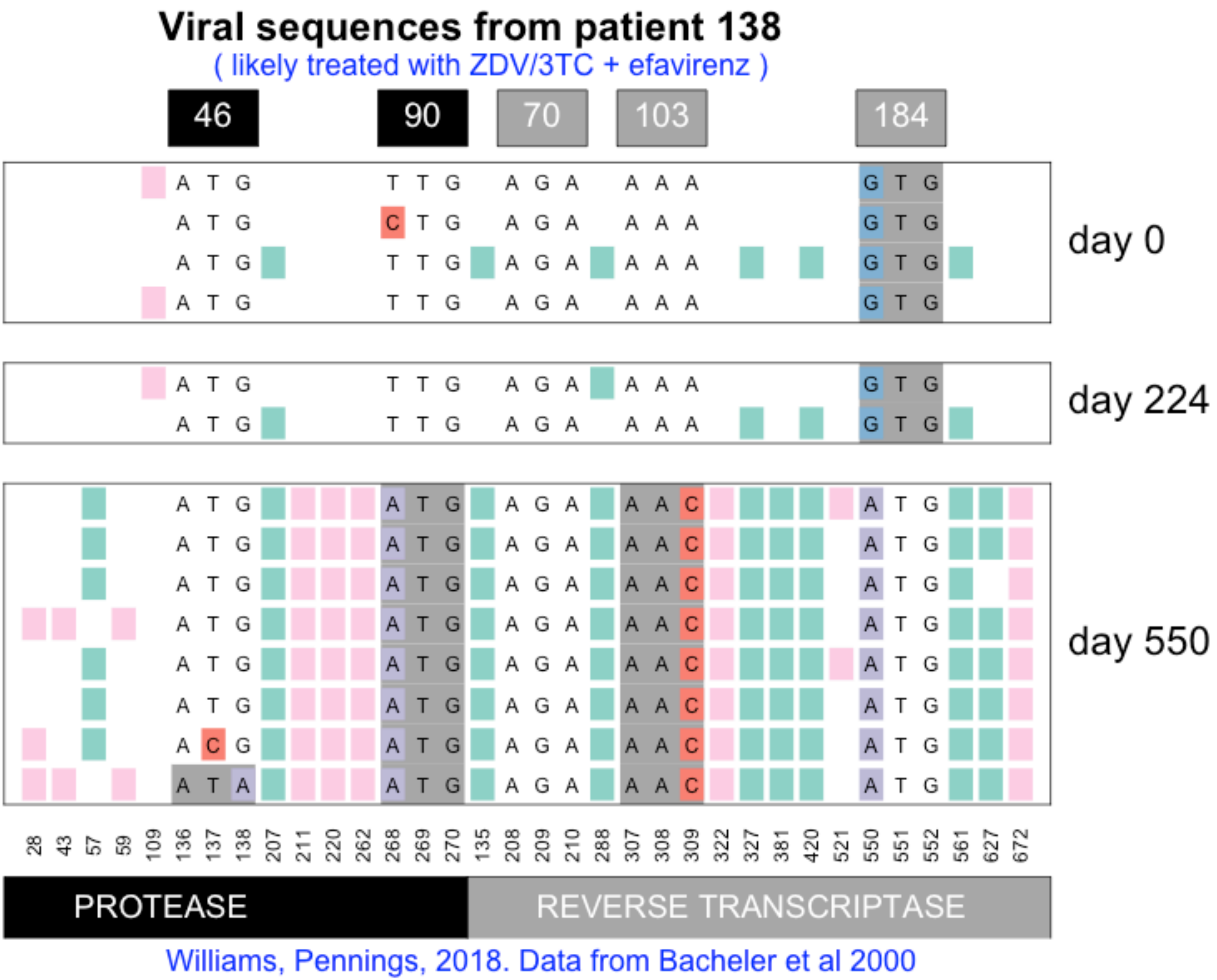
Viral sequences from patient 138
( likely treated with ZDV/3TC + efavirenz )
46
90
70
103
184
day 0
day 224
day 550
28 43 57 59 109 136 137 138 207 211 220 262 268 269 270 135 208 209 210 288 307 308 309 322 327 381 420 521 550 551 552 561 627 672
PROTEASE
REVERSE TRANSCRIPTASE
Williams, Pennings, 2018. Data from Bacheler et al 2000

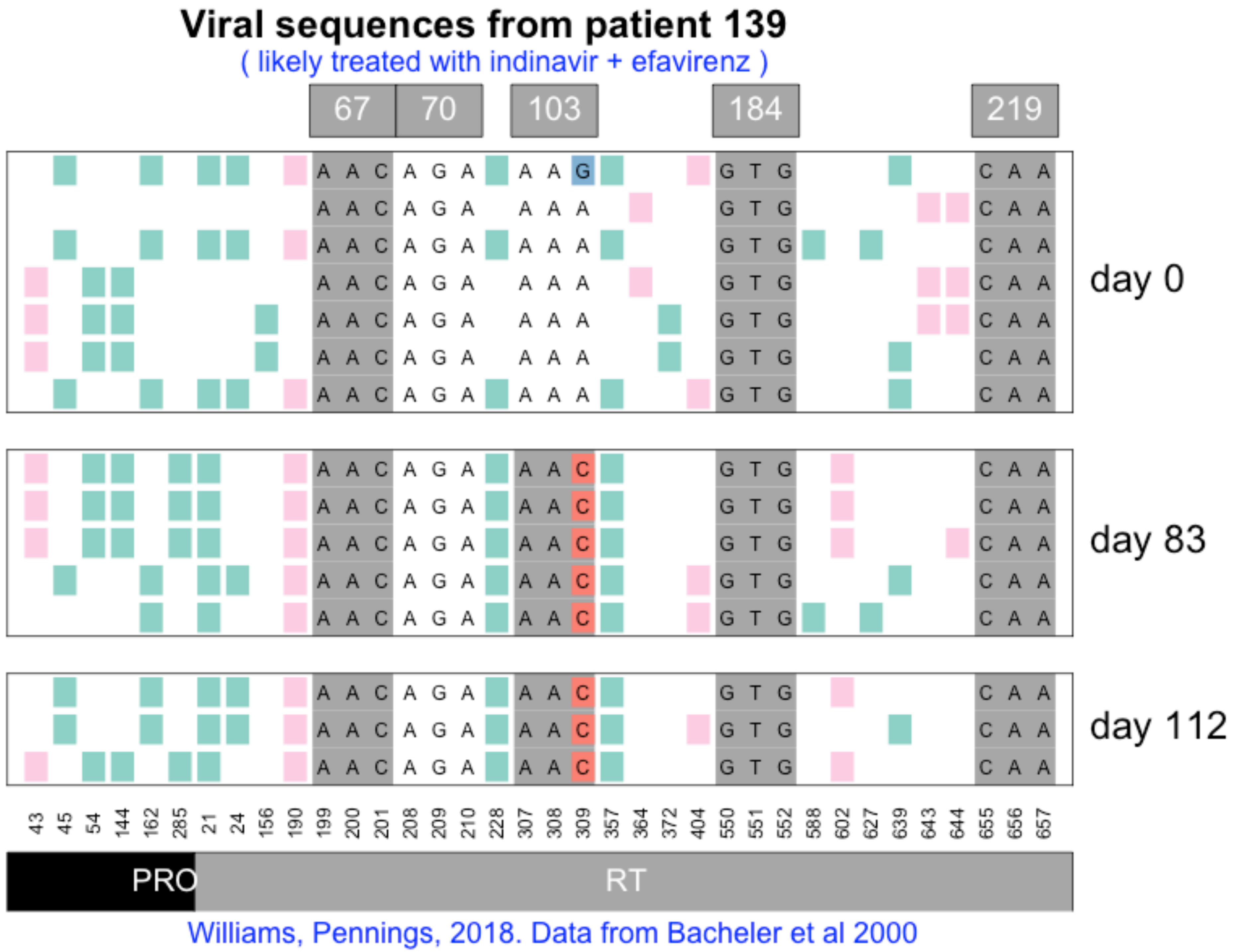
Viral sequences from patient 139
( likely treated with indinavir + efavirenz )
67
70
103
184
219
day 0
day 83
day 112
43 45 54 144 162 285 21 24 156 190 199 200 201 208 209 210 228 307 308 309 357 364 372 404 550 551 552 588 602 627 639 643 644 655 656 657
PRO
RT
Williams, Pennings, 2018. Data from Bacheler et al 2000

#### Viral sequences from patient 140

( likely treated with indinavir + efavirenz )

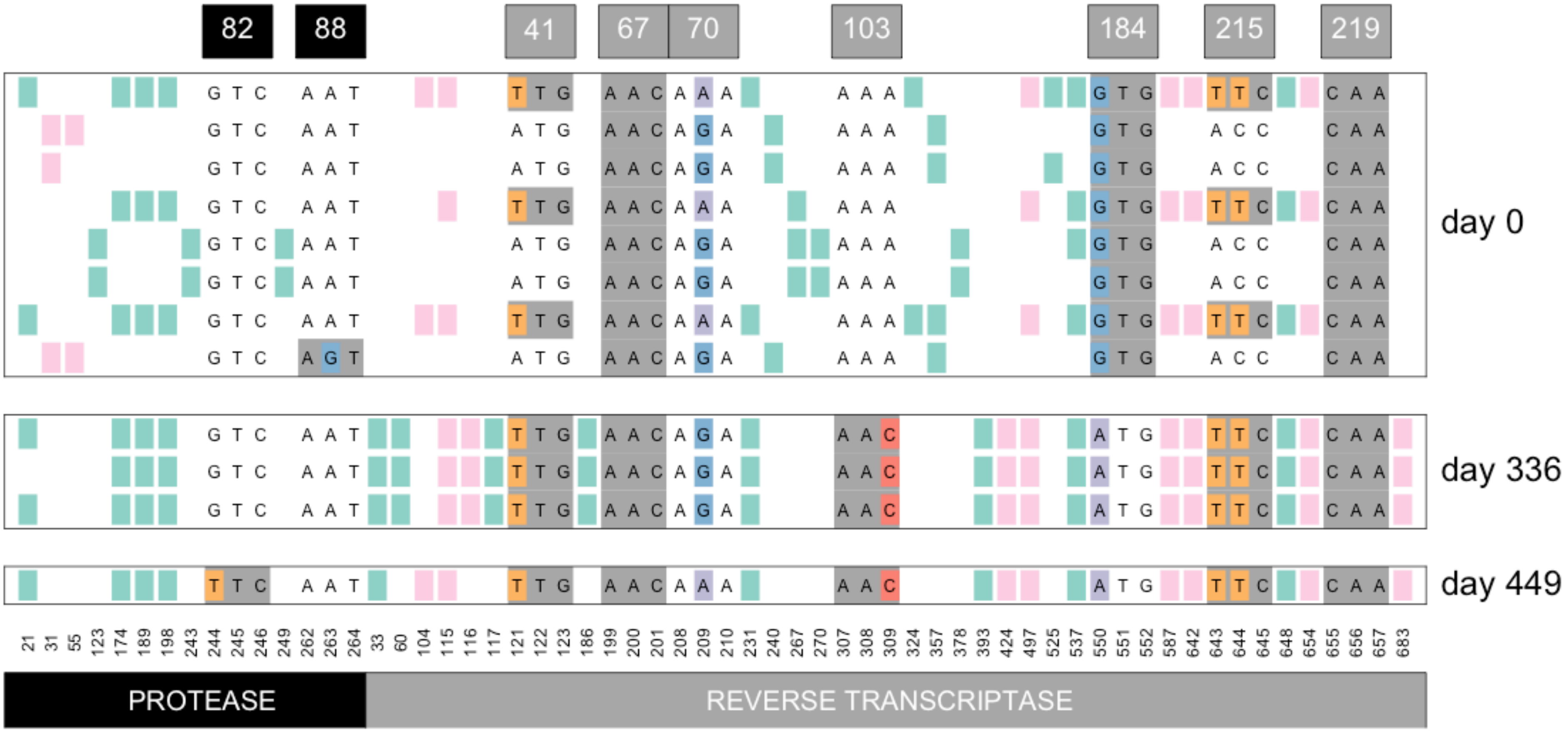


Williams, Pennings, 2018. Data from Bacheler et al 2000

#### Viral sequences from patient 141

( likely treated with indinavir then later switched to efavirenz combination therapy )

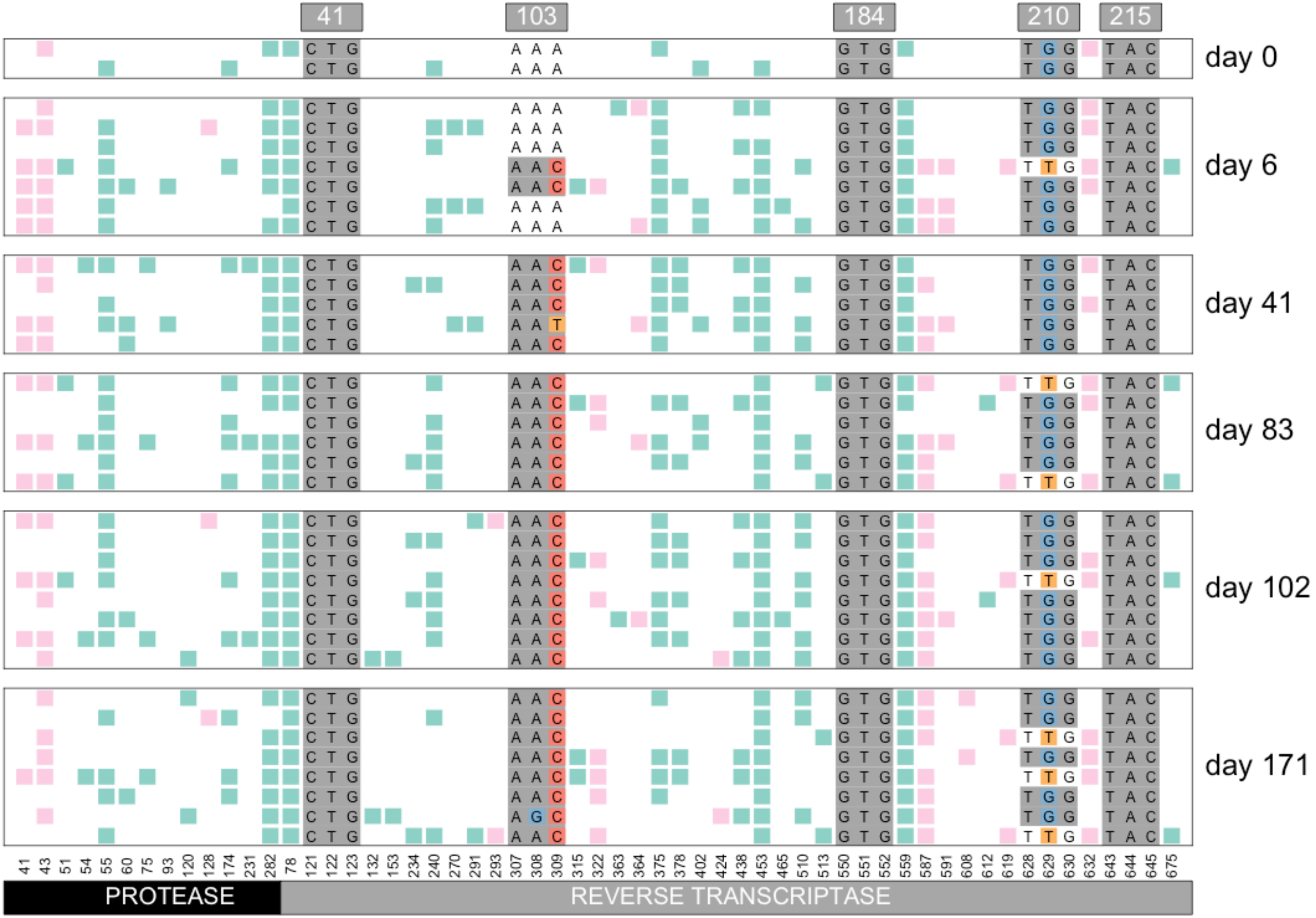


Williams, Pennings, 2018. Data from Bacheler et al 2000

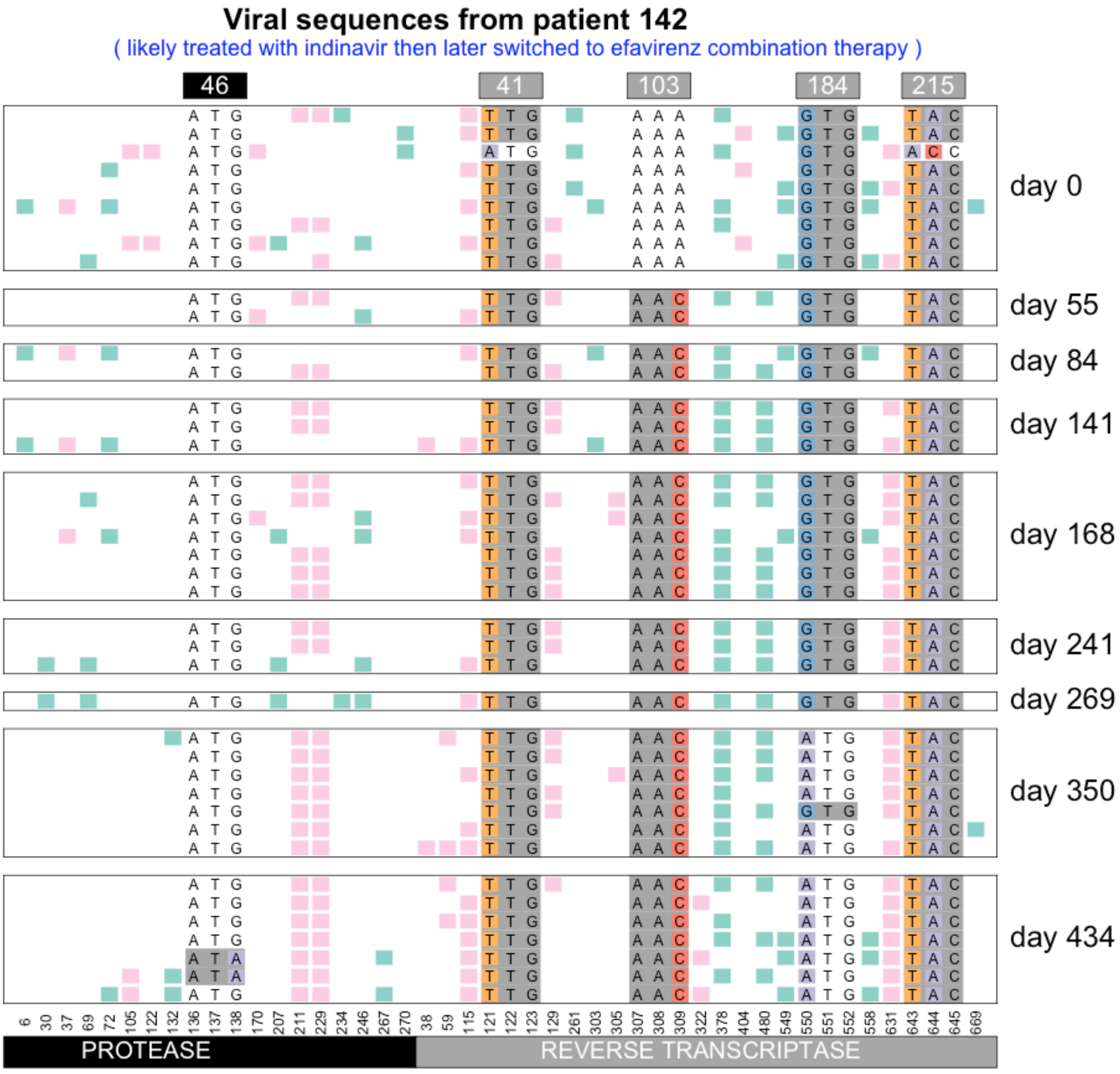


Williams, Pennings, 2018. Data from Bacheler et al 2000

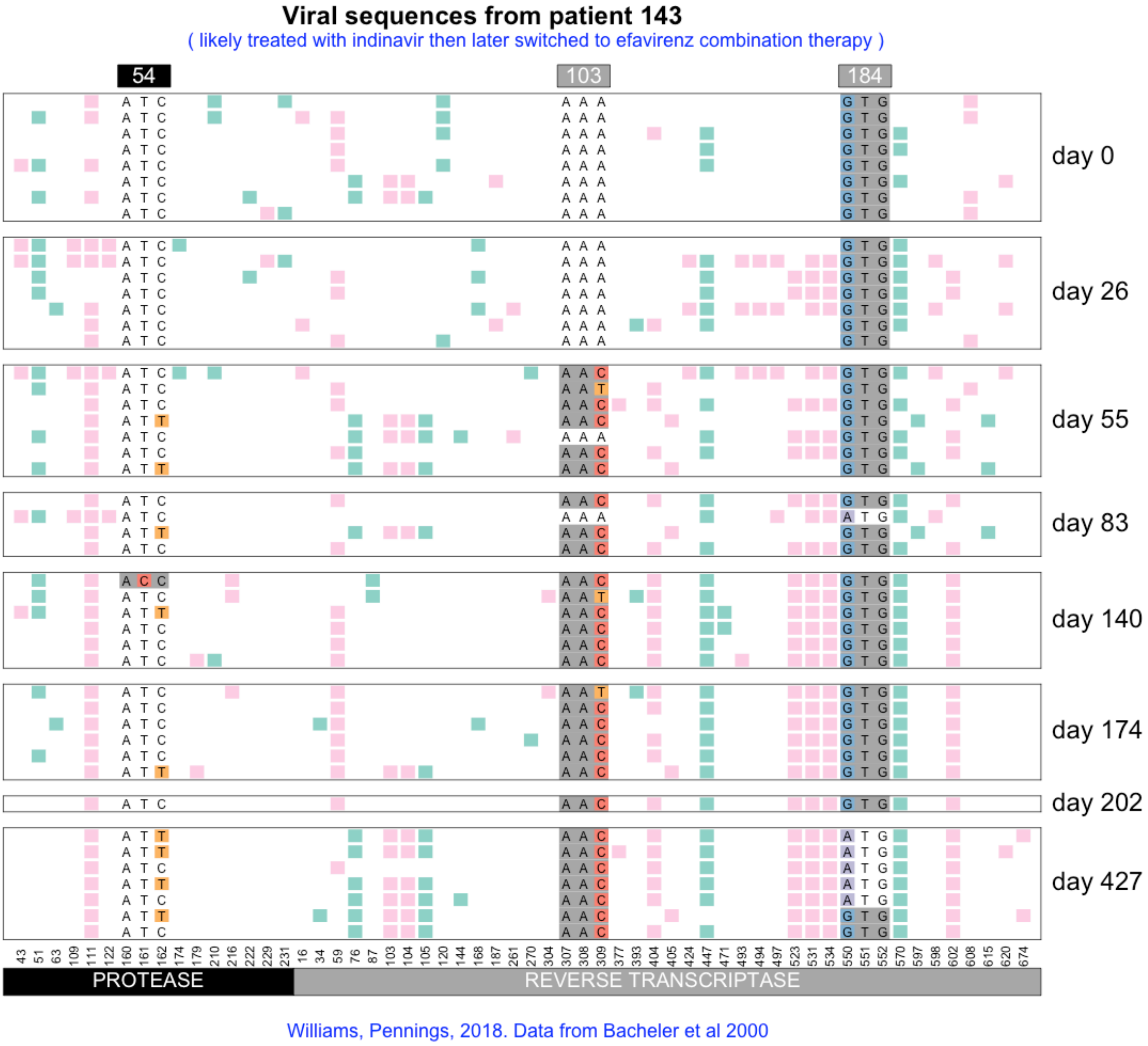

Viral sequences from patient 143
( likely treated with indinavir then later switched to efavirenz combination therapy )
54
103
184
day 0
day 26
day 55
day 83
day 140
day 174
day 202
day 427
43 51 63 109 111 122 160 161 162 174 179 210 216 222 229 231 16 34 59 76 87 103 104 105 120 144 168 187 261 270 304 307 308 309 377 393 404 405 424 447 471 493 494 497 523 531 534 550 551 552 570 597 598 602 608 615 620 674
PROTEASE
REVERSE TRANSCRIPTASE
Williams, Pennings, 2018. Data from Bacheler et al 2000

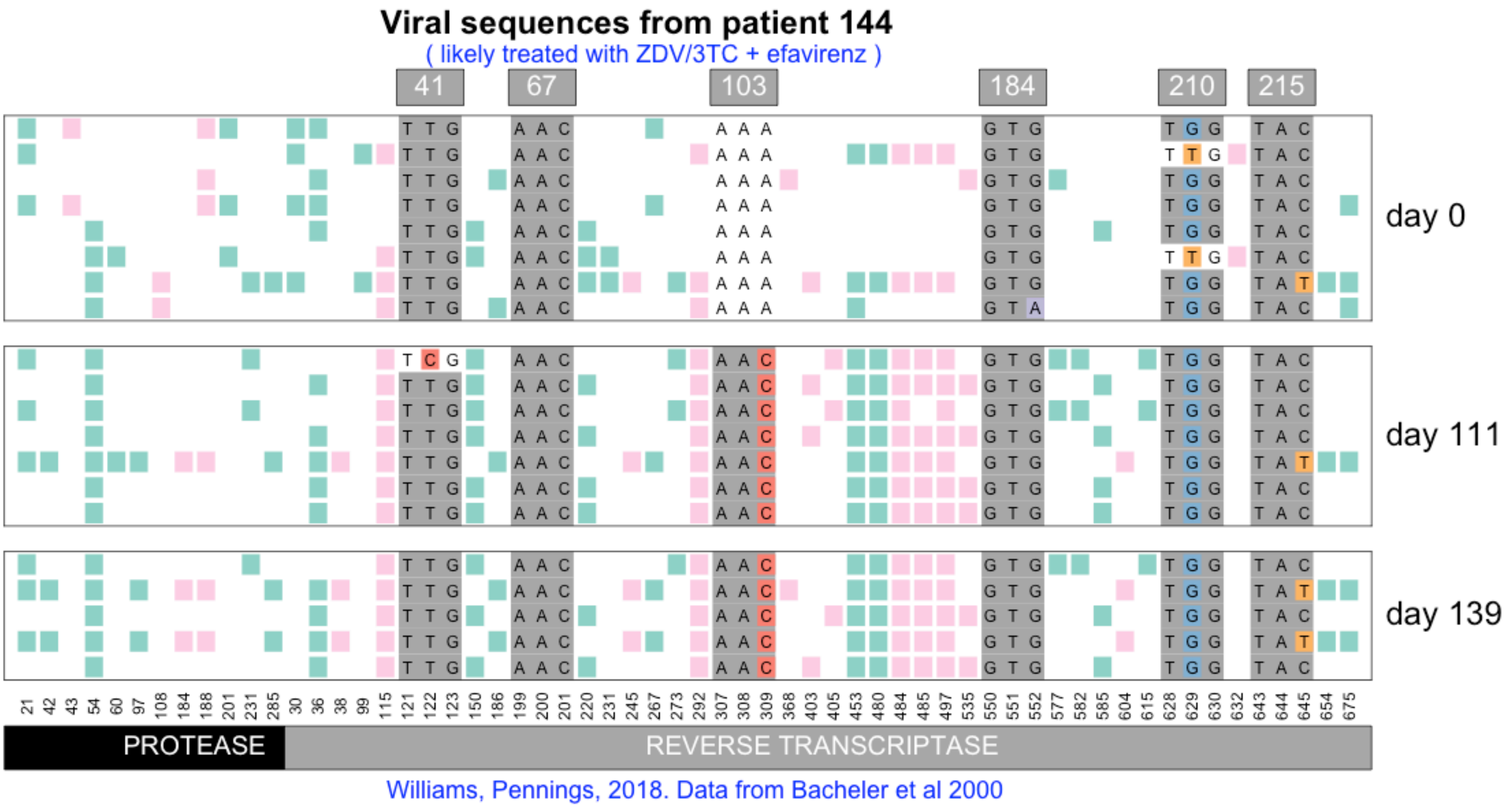
Viral sequences from patient 144
( likely treated with ZDV/3TC + efavirenz )
41
67
103
184
210
215
day 0
day 111
day 139
21 42 43 54 60 97 108 184 188 201 231 285 30 36 38 99 115 121 122 123 150 186 199 200 201 220 231 245 267 273 292 307 308 309 368 403 405 453 480 484 485 497 535 550 551 552 577 582 585 604 615 628 629 630 632 643 644 645 654 675
PROTEASE
REVERSE TRANSCRIPTASE
Williams, Pennings, 2018. Data from Bacheler et al 2000

#### Viral sequences from patient 145

( likely treated with indinavir then later switched to efavirenz combination therapy )

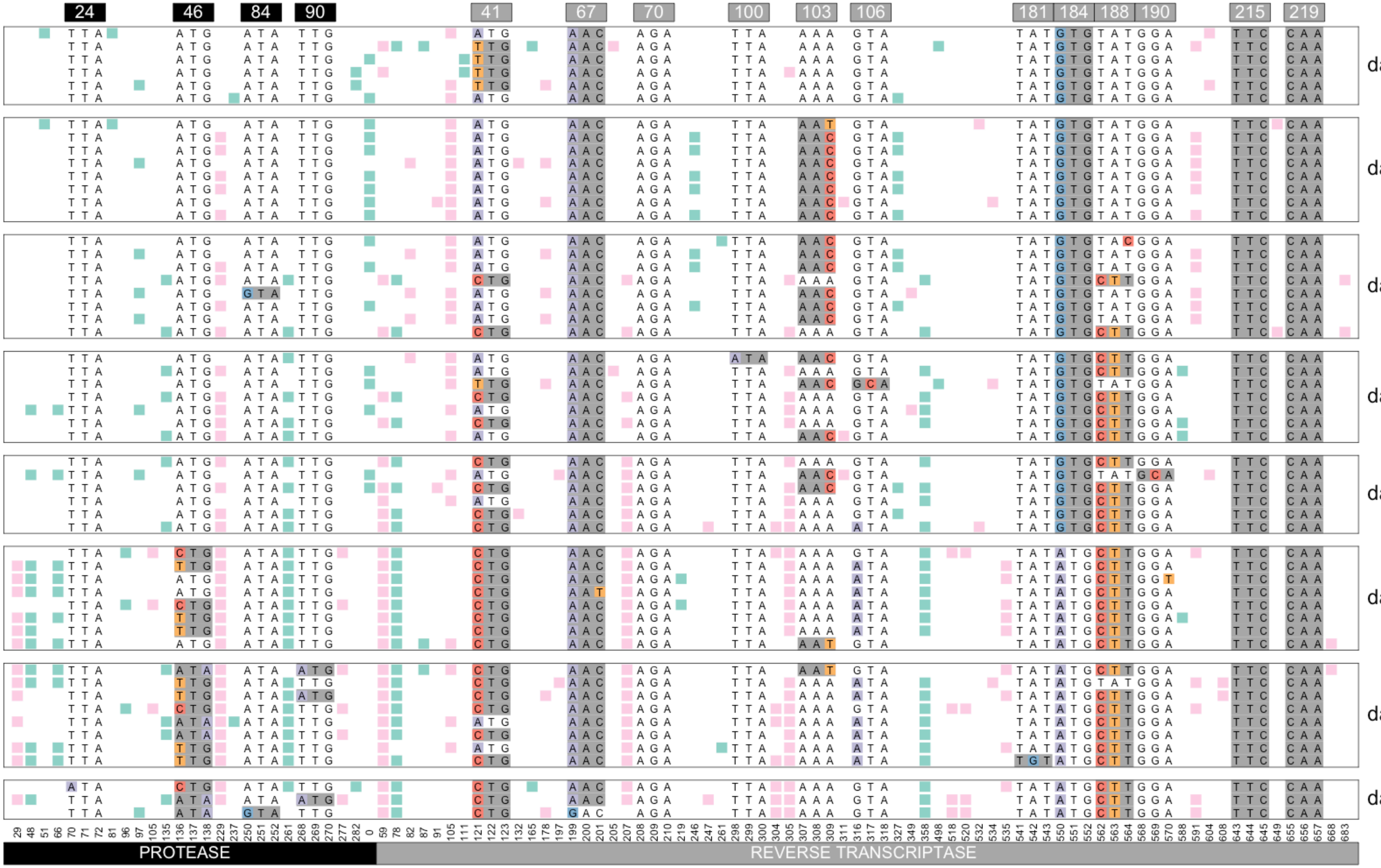


Williams, Pennings, 2018. Data from Bacheler et al 2000

#### Viral sequences from patient 146

( likely treated with indinavir then later switched to efavirenz combination therapy )

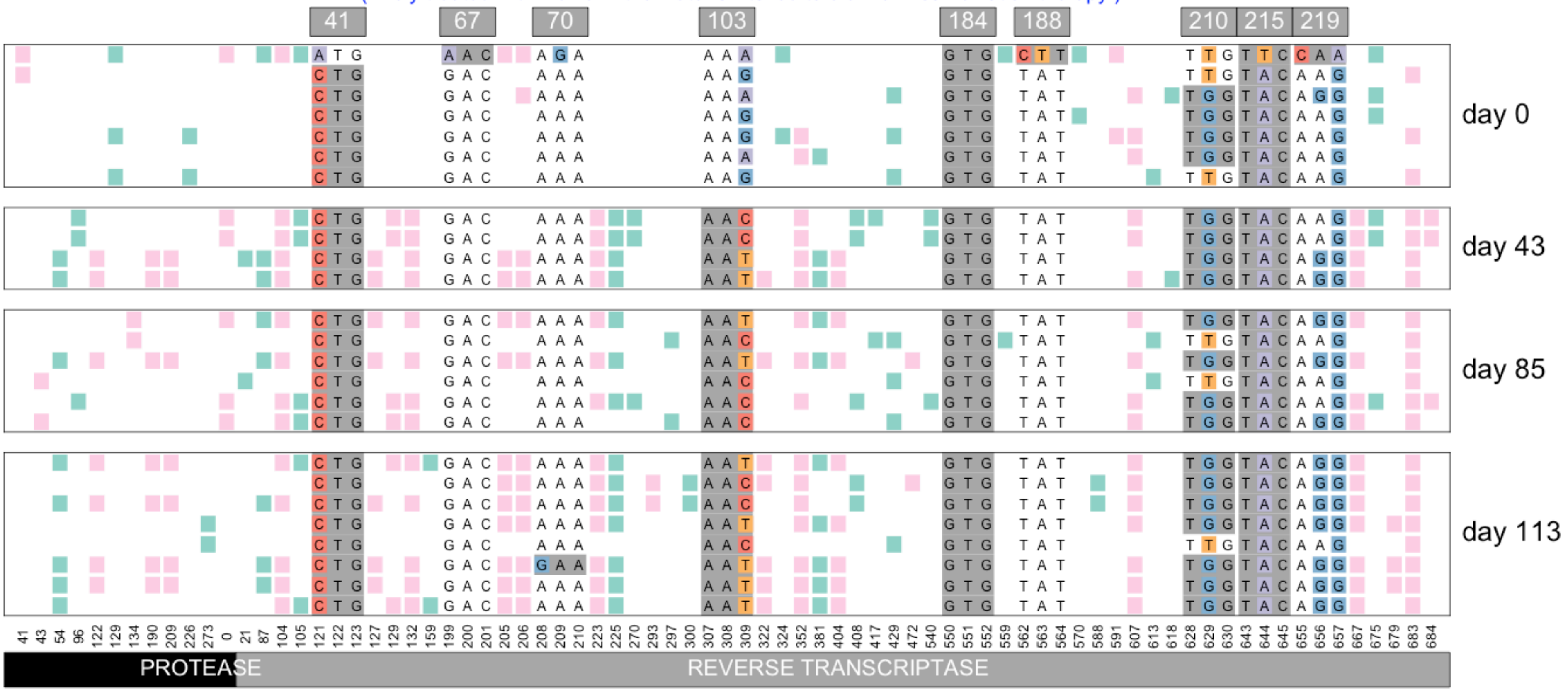


Williams, Pennings, 2018. Data from Bacheler et al 2000

#### Viral sequences from patient 147

( likely treated with indinavir then later switched to efavirenz combination therapy )

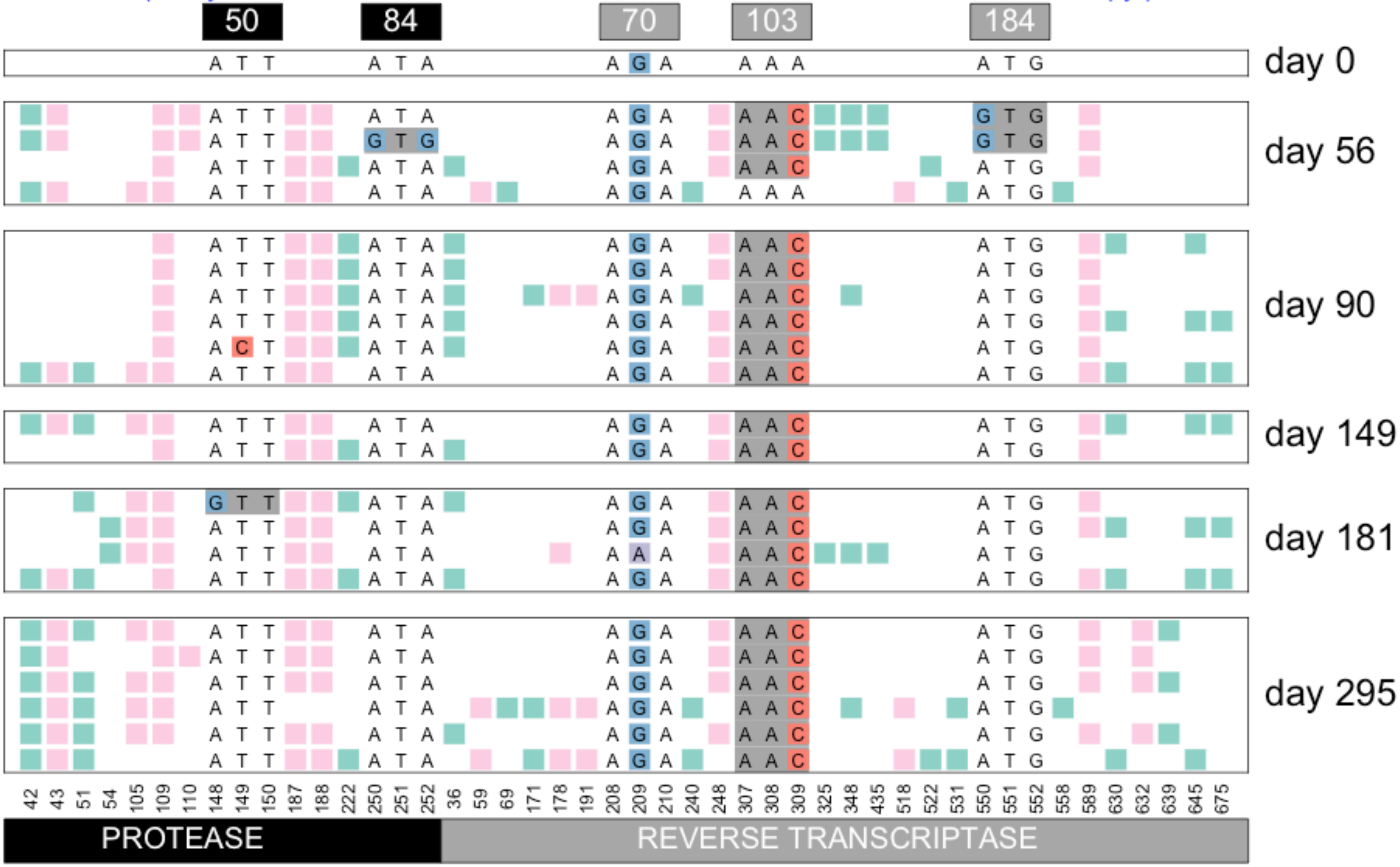


Williams, Pennings, 2018. Data from Bacheler et al 2000

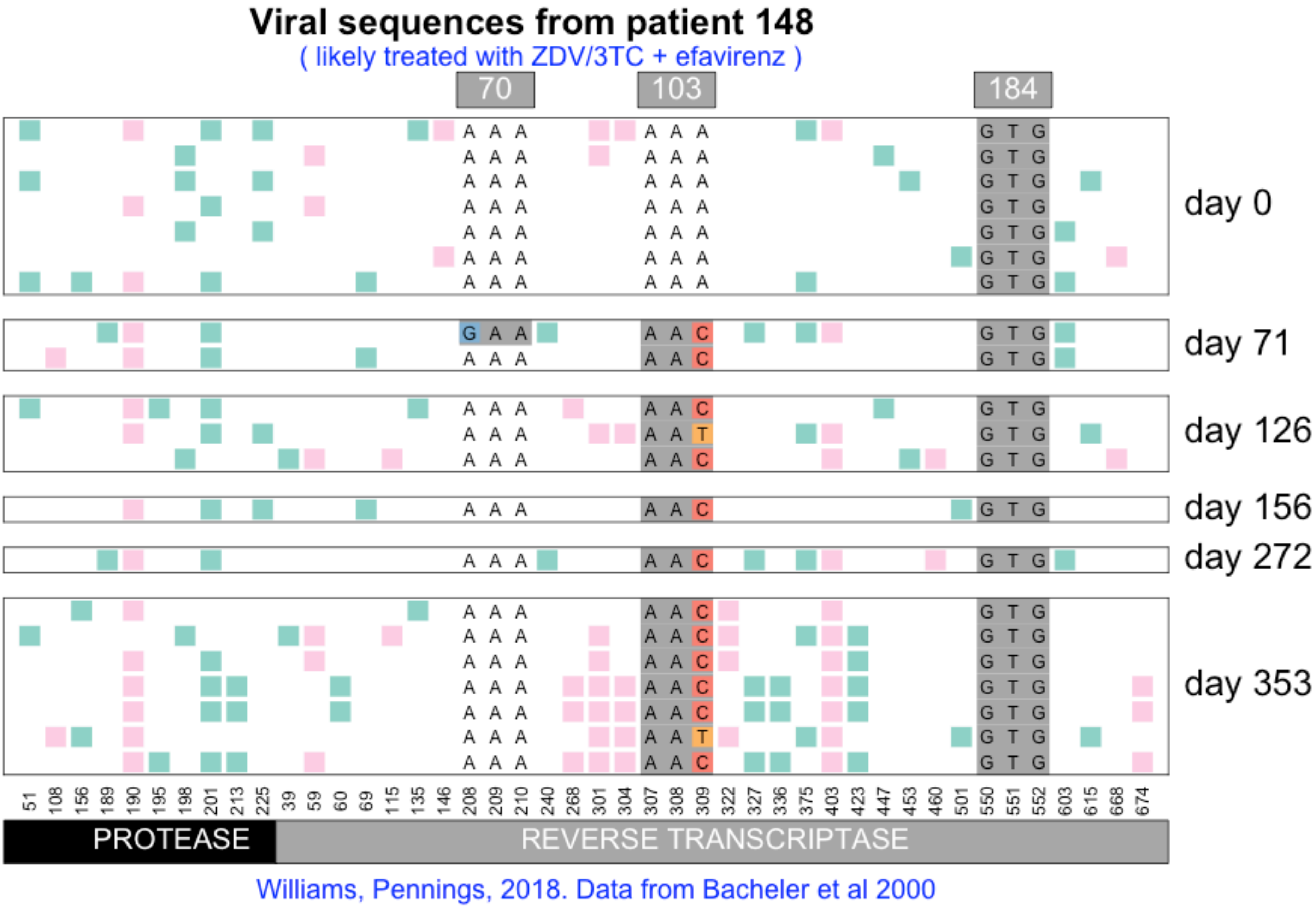
Viral sequences from patient 148
( likely treated with ZDV/3TC + efavirenz )
70
103
184
day 0
day 71
day 126
day 156
day 272
day 353
51 108 156 189 190 195 198 201 213 225 39 59 60 69 115 135 146 208 209 210 240 268 301 304 307 308 309 322 327 336 375 403 423 447 453 460 501 550 551 552 603 615 668 674
PROTEASE
REVERSE TRANSCRIPTASE
Williams, Pennings, 2018. Data from Bacheler et al 2000

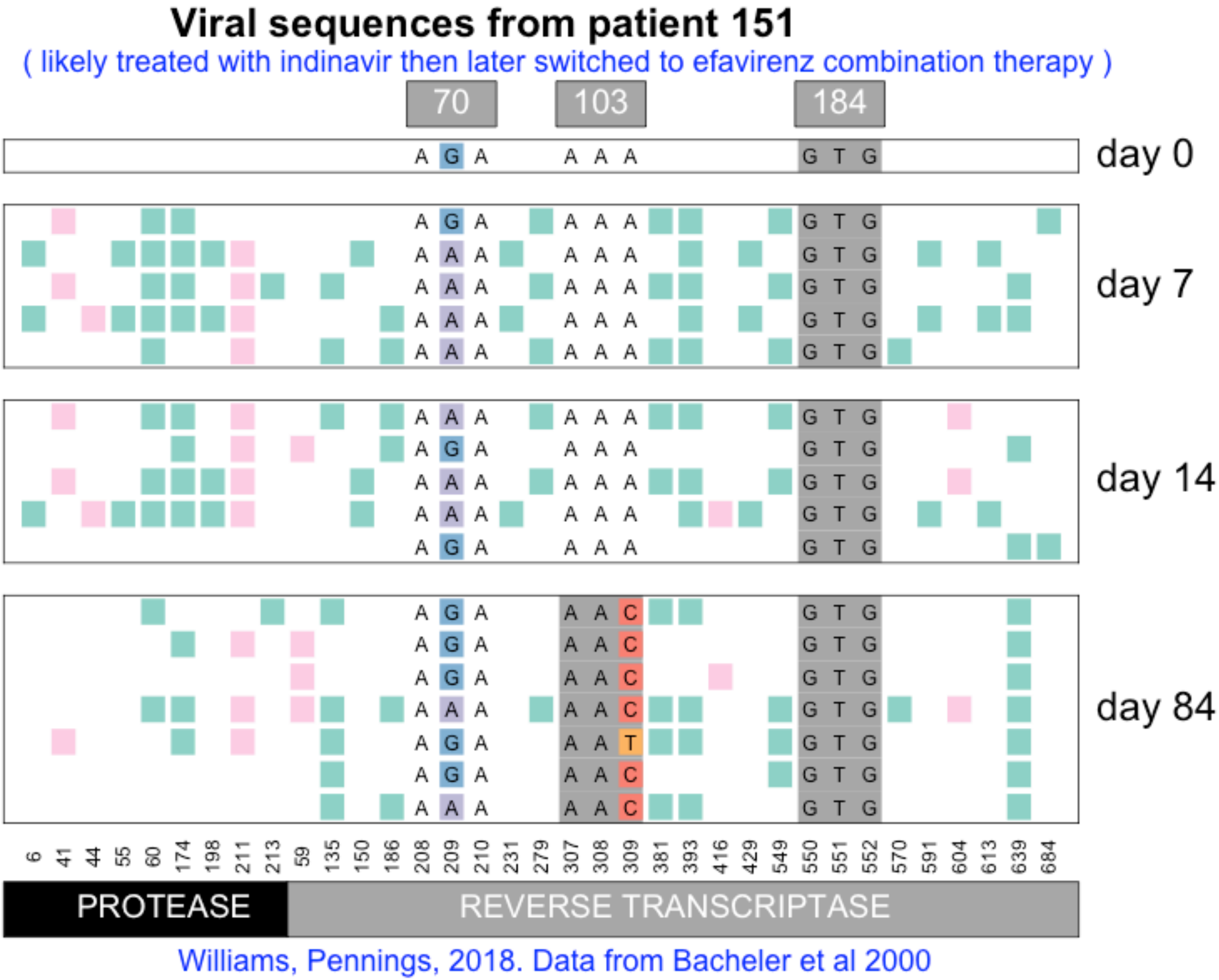
Viral sequences from patient 151
( likely treated with indinavir then later switched to efavirenz combination therapy )
70
103
184
day 0
day 7
day 14
day 84
6 41 44 55 60 174 198 211 213 59 135 150 186 208 209 210 231 279 307 308 309 381 393 416 429 549 550 551 552 570 591 604 613 639 684
PROTEASE
REVERSE TRANSCRIPTASE
Williams, Pennings, 2018. Data from Bacheler et al 2000

#### Viral sequences from patient 152

( likely treated with ZDV/3TC + efavirenz )

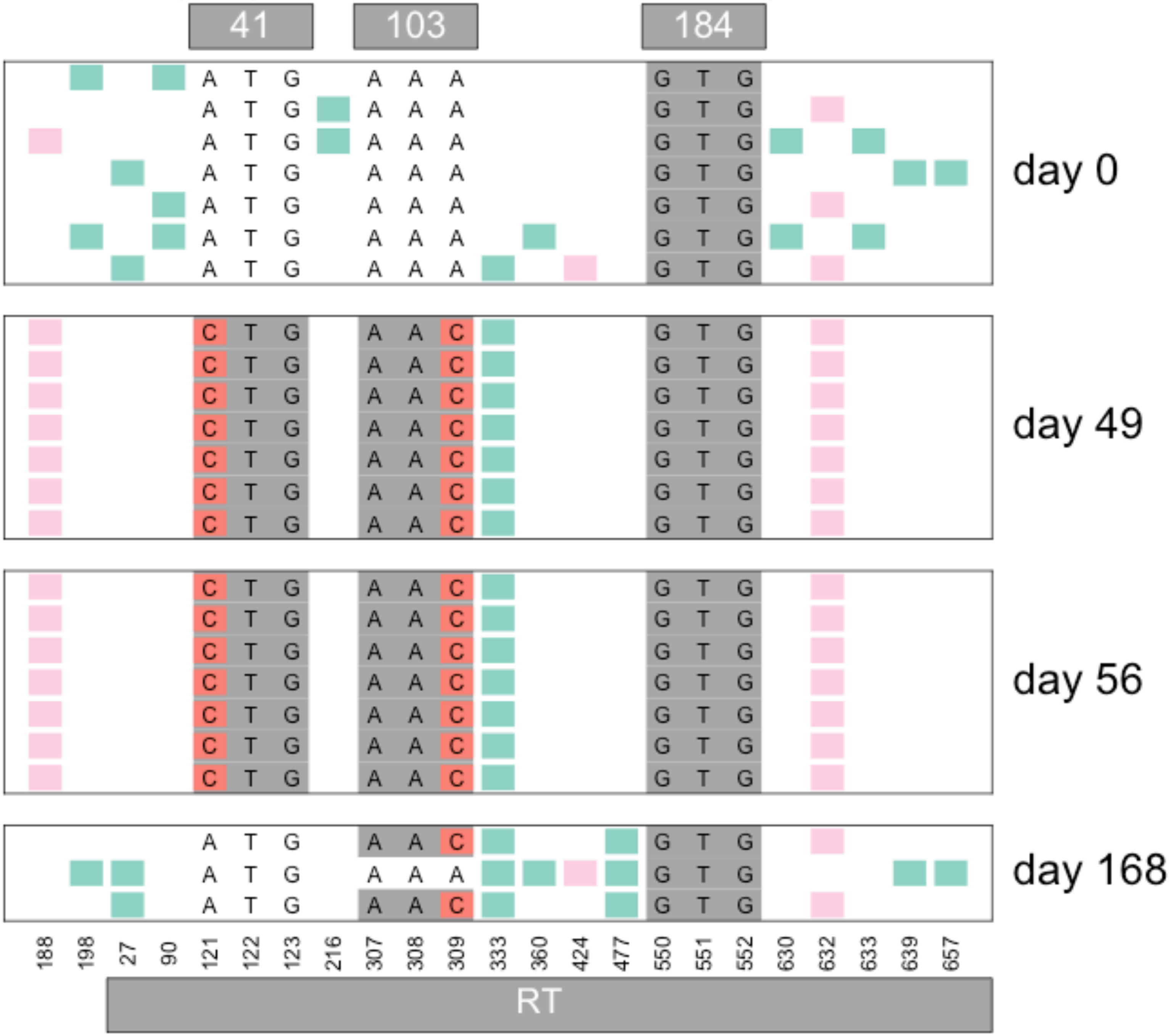


Williams, Pennings, 2018. Data from Bacheler et al 2000

### Viral sequences from patient 153

( likely treated with ZDV/3TC + efavirenz )

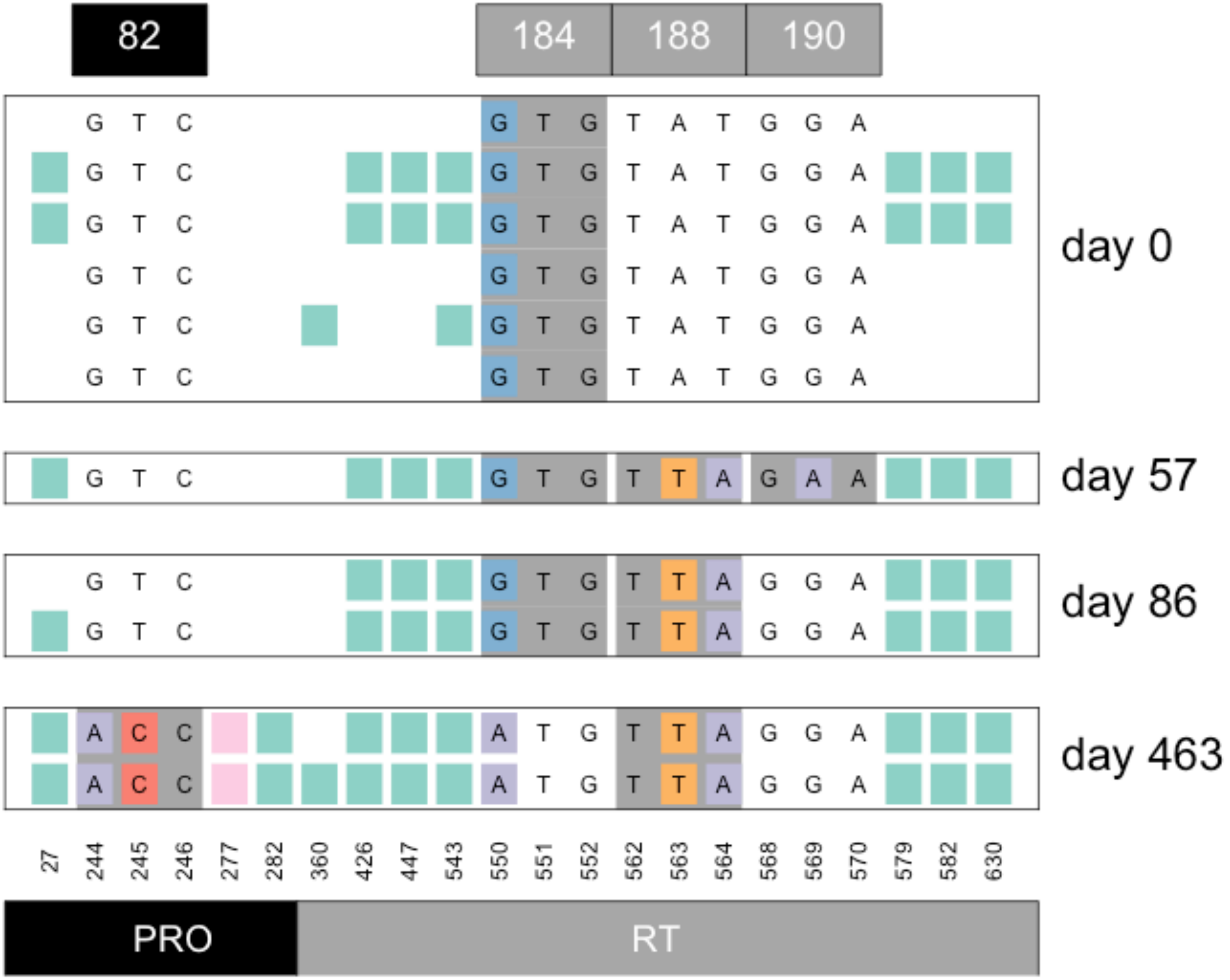


Williams, Pennings, 2018. Data from Bacheler et al 2000

#### Viral sequences from patient 154

( likely treated with ZDV/3TC + efavirenz )

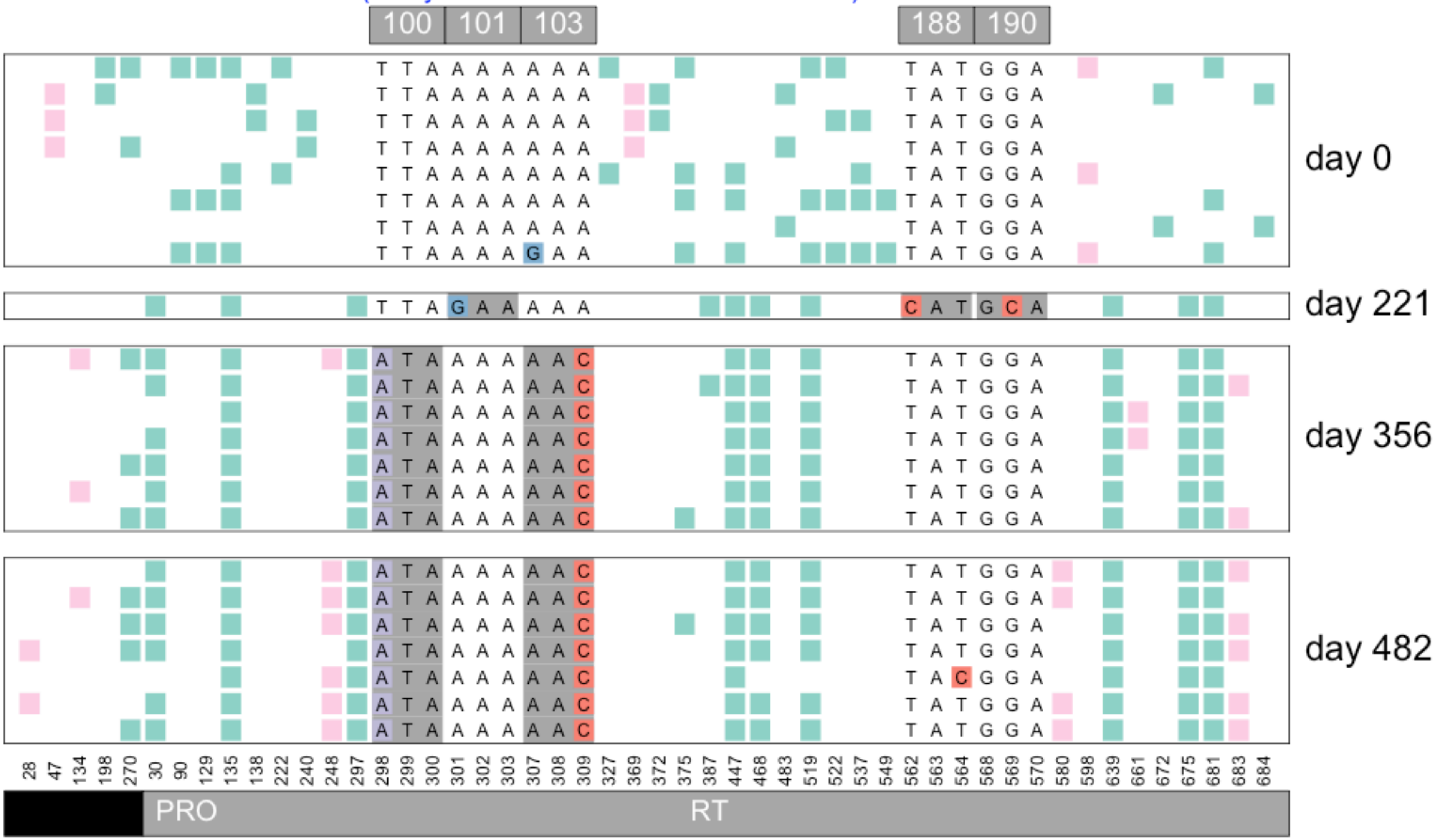


Williams, Pennings, 2018. Data from Bacheler et al 2000

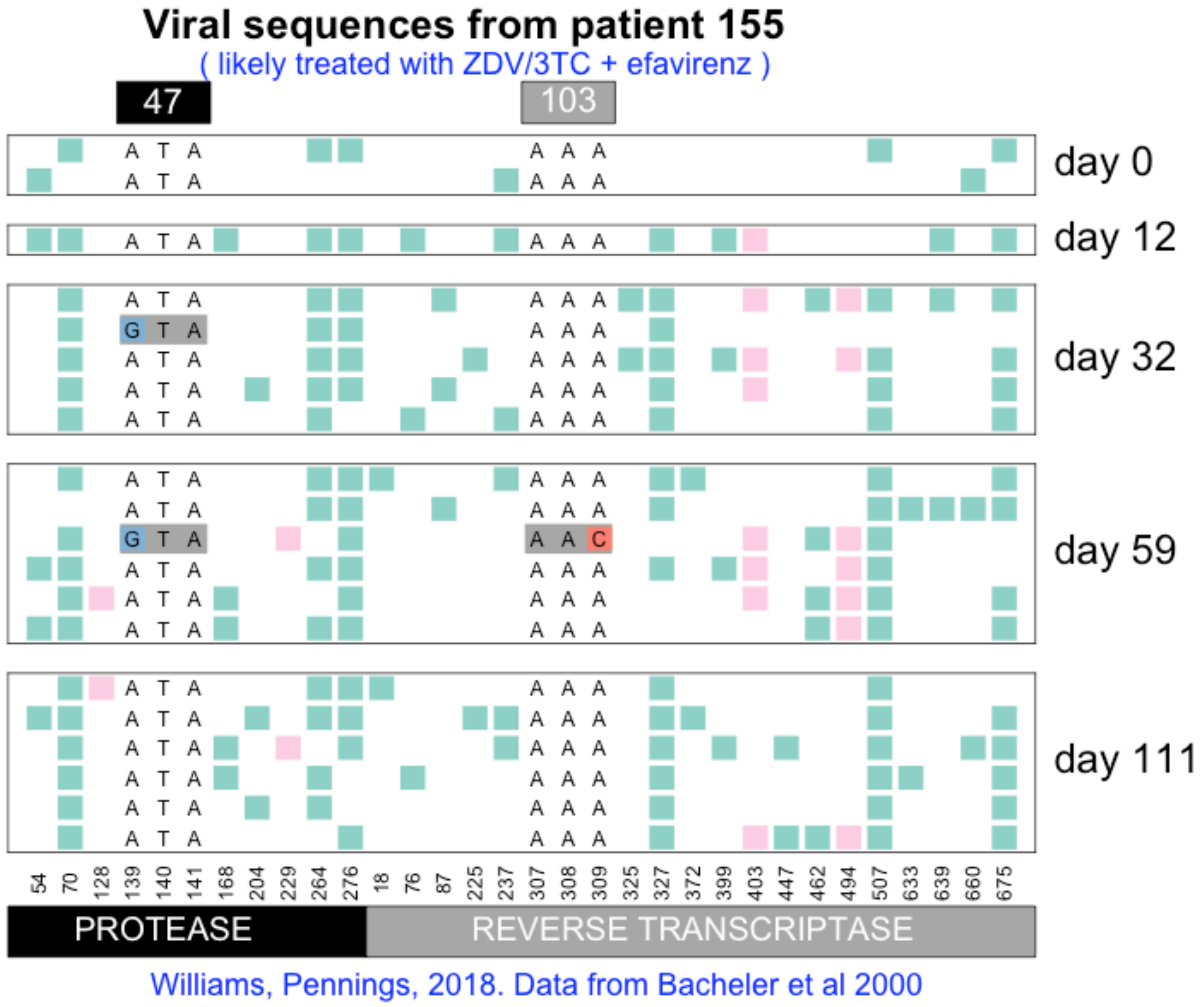

Viral sequences from patient 155
( likely treated with ZDV/3TC + efavirenz )
47
103
day 0
day 12
day 32
day 59
day 111
54 70 128 139 140 141 168 204 229 264 276 18 76 87 225 237 307 308 309 325 327 372 399 403 447 462 494 507 633 639 660 675
PROTEASE
REVERSE TRANSCRIPTASE
Williams, Pennings, 2018. Data from Bacheler et al 2000

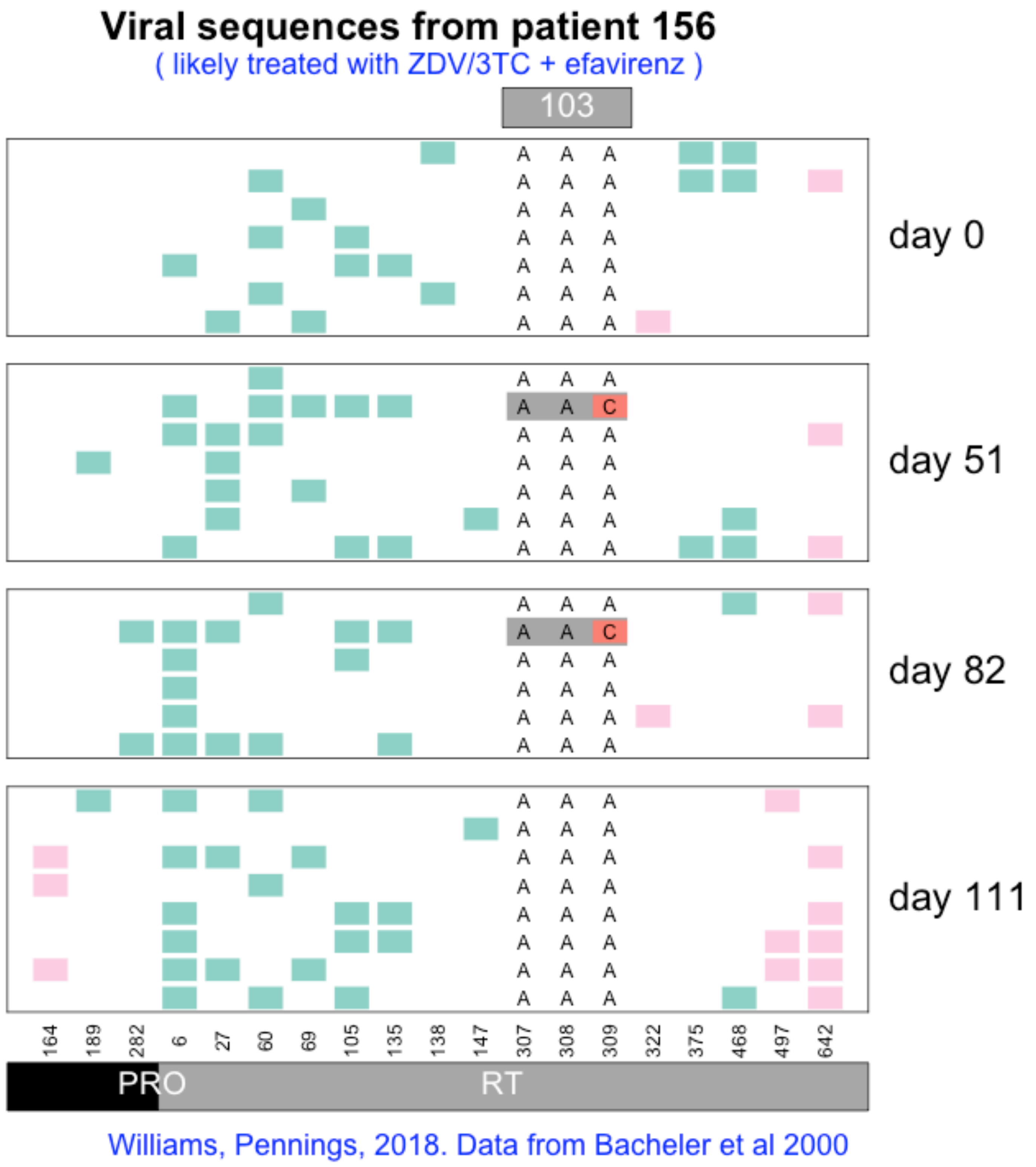

Viral sequences from patient 156
( likely treated with ZDV/3TC + efavirenz )
103
day 0
day 51
day 82
day 111
164
189
282
6
27
60
69
105
135
138
147
307
308
309
322
375
468
497
642
PRO
RT
Williams, Pennings, 2018. Data from Bacheler et al 2000

### Viral sequences from patient 157

( likely treated with ZDV/3TC + efavirenz )

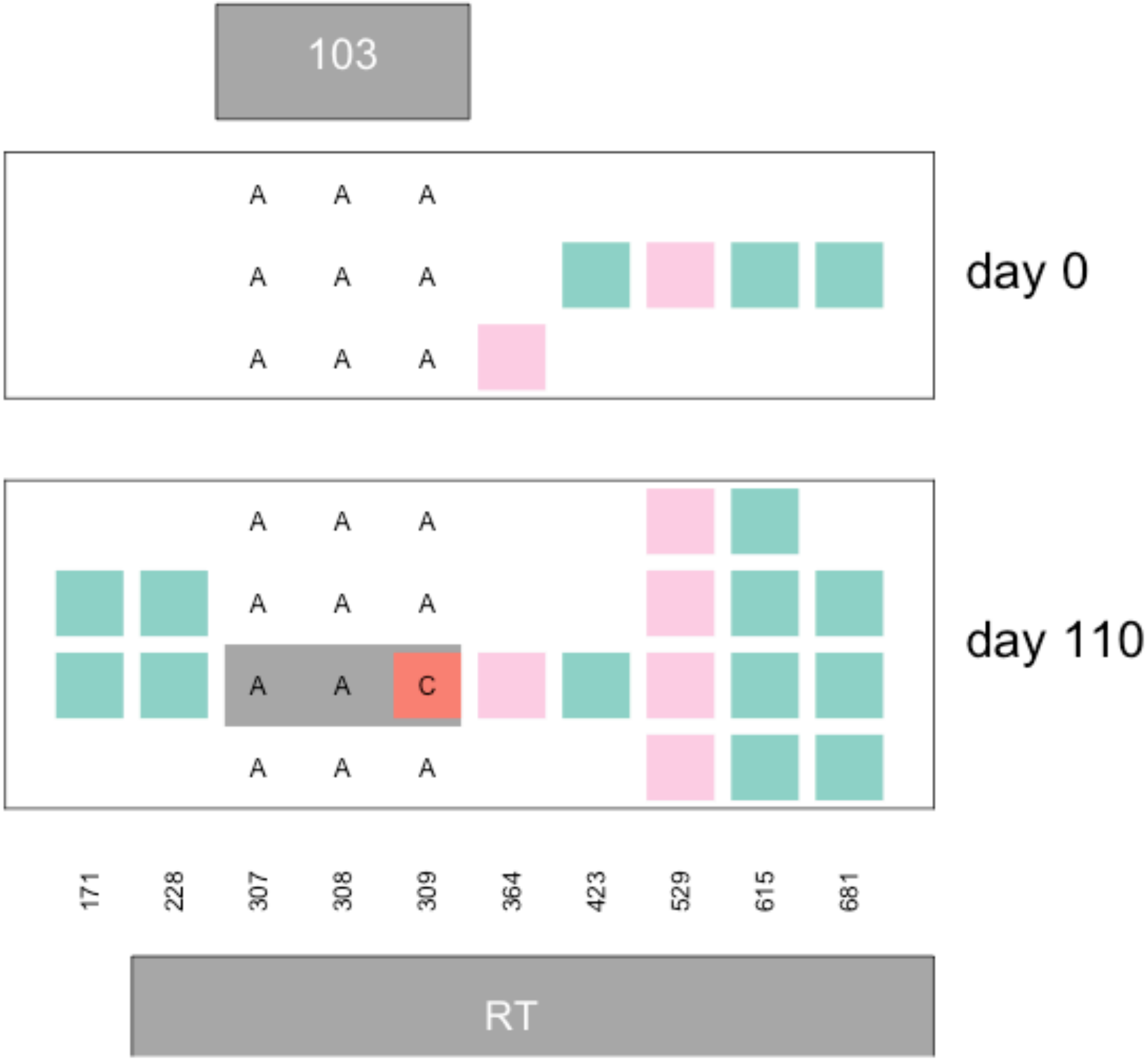


Williams, Pennings, 2018. Data from Bacheler et al 2000

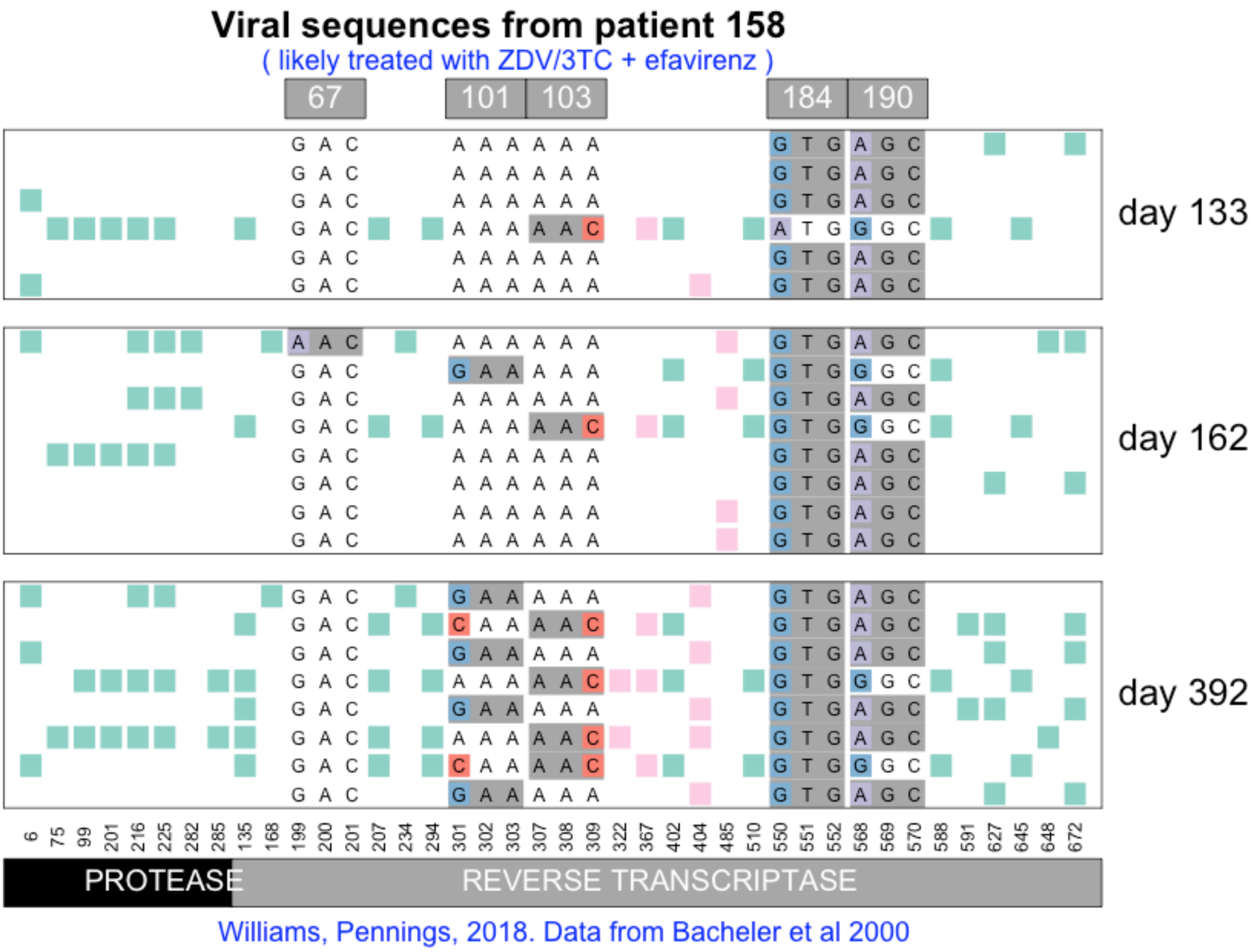

Viral sequences from patient 158
( likely treated with ZDV/3TC + efavirenz )
67
101
103
184
190
day 133
day 162
day 392
PROTEASE
REVERSE TRANSCRIPTASE
Williams, Pennings, 2018. Data from Bacheler et al 2000

#### Viral sequences from patient 159

( likely treated with ZDV/3TC + efavirenz )

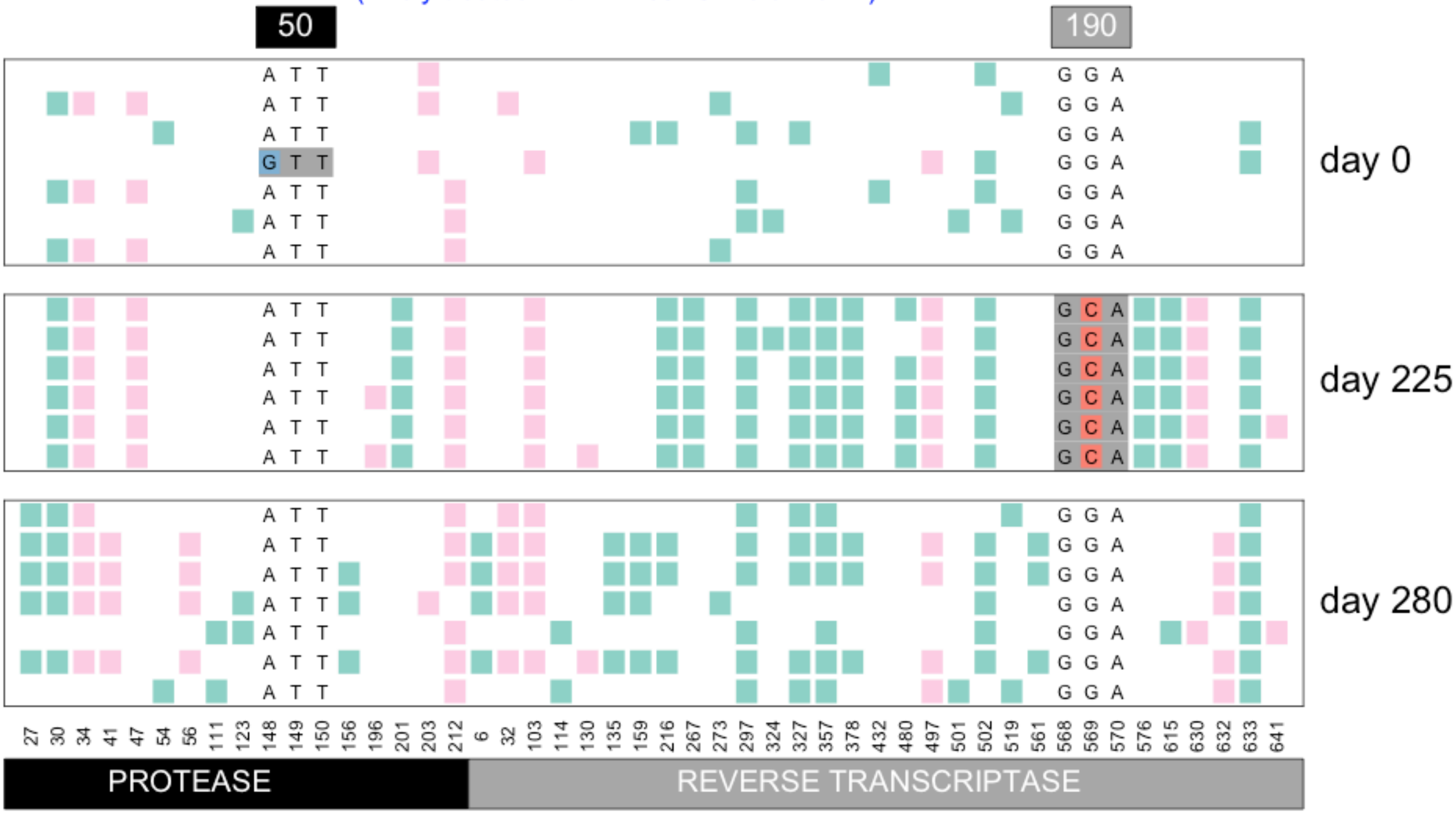


Williams, Pennings, 2018. Data from Bacheler et al 2000

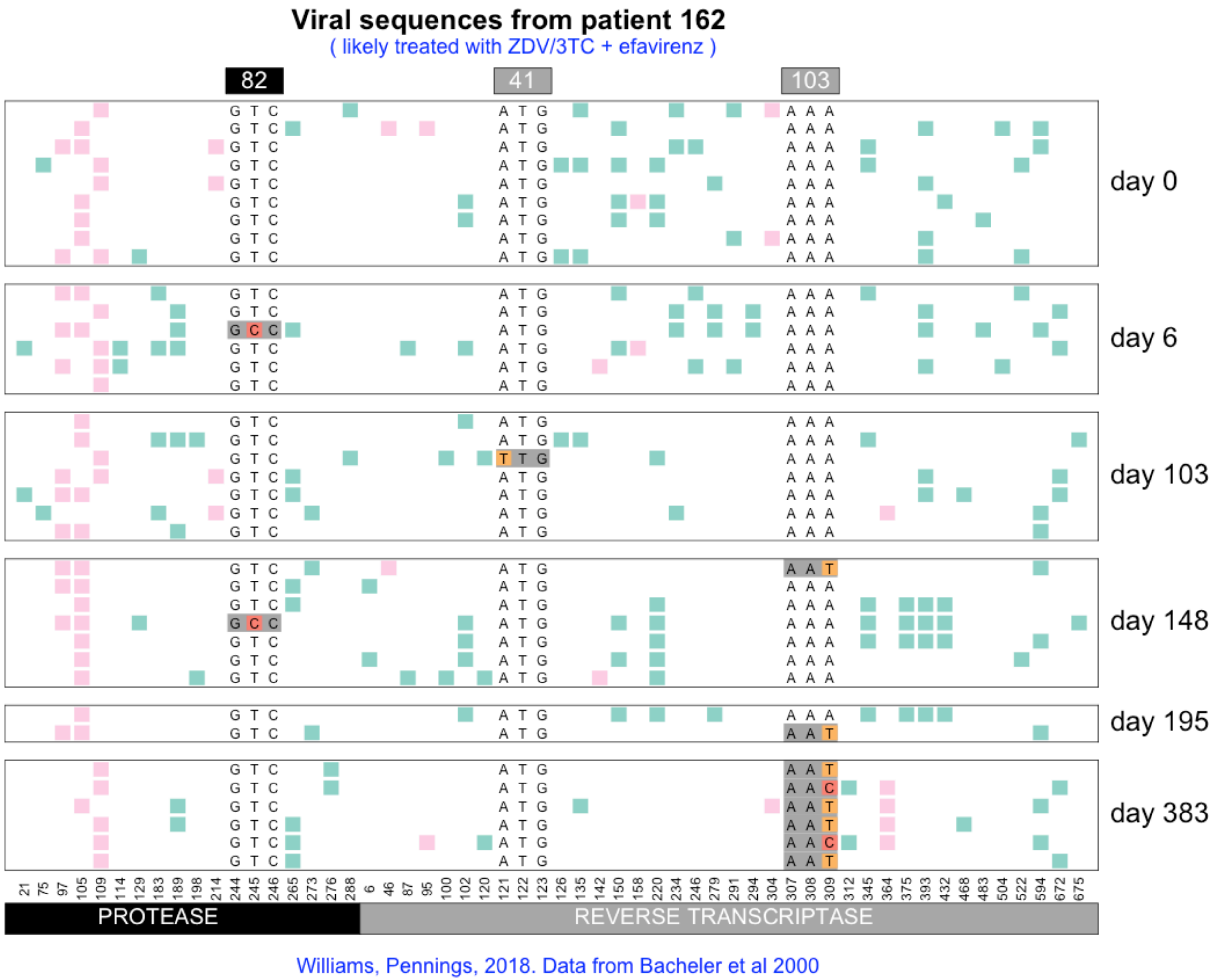
Viral sequences from patient 162
( likely treated with ZDV/3TC + efavirenz )
82
41
103
day 0
day 6
day 103
day 148
day 195
day 383
21 75 97 105 109 114 129 183 189 198 214 244 245 246 265 273 276 288 6 46 87 95 100 102 120 121 122 123 126 135 142 150 158 220 234 246 279 291 294 304 307 308 309 312 345 364 375 393 432 468 483 504 522 594 672 675
PROTEASE
REVERSE TRANSCRIPTASE
Williams, Pennings, 2018. Data from Bacheler et al 2000

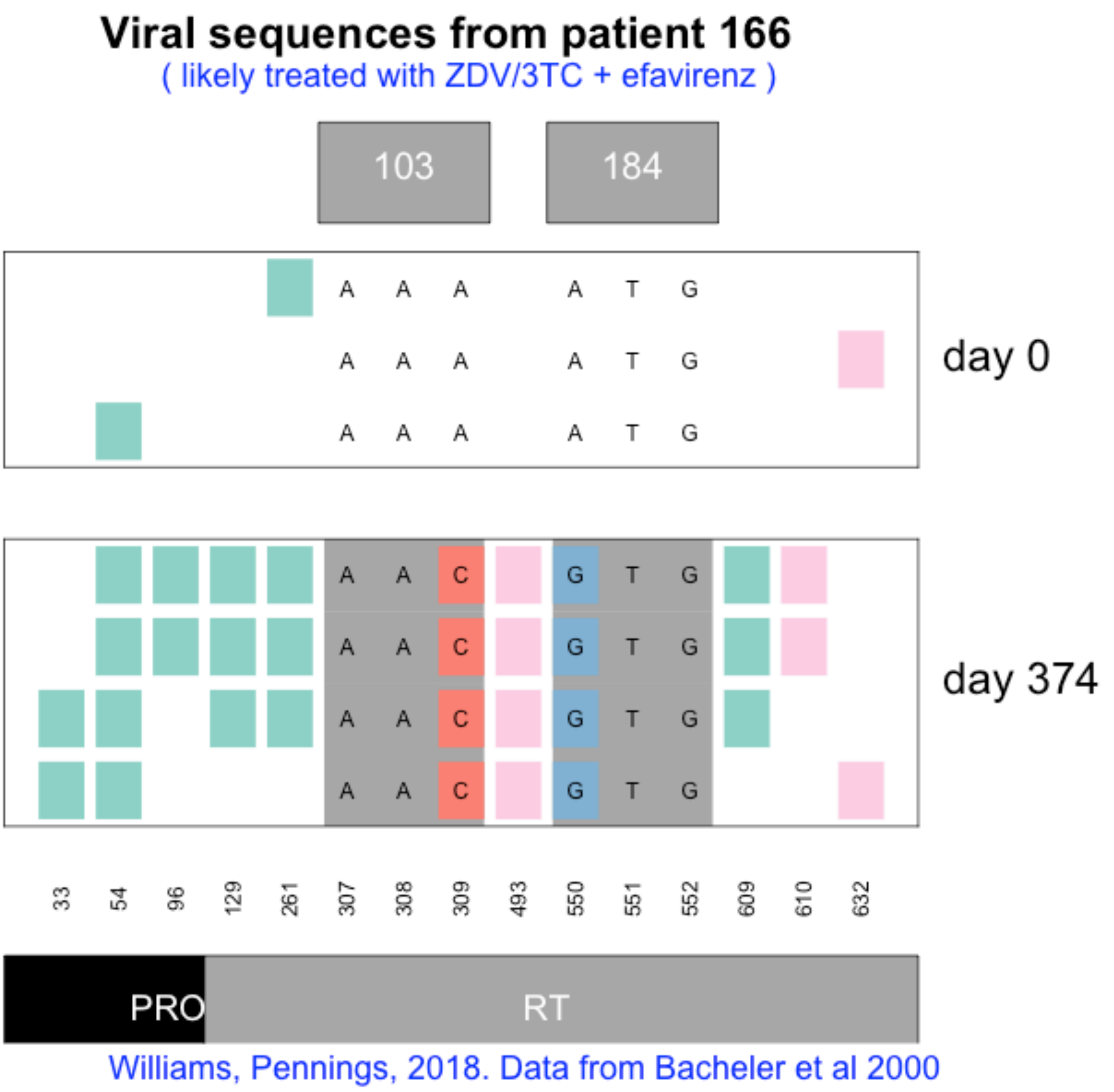
Viral sequences from patient 166
( likely treated with ZDV/3TC + efavirenz )
103
184
A A A A T G
A A A A T G
A A A A T G
day 0
A A C G T G
A A C G T G
A A C G T G
A A C G T G
day 374
33
54
96
129
261
307
308
309
493
550
551
552
609
610
632
PRO
RT
Williams, Pennings, 2018. Data from Bacheler et al 2000

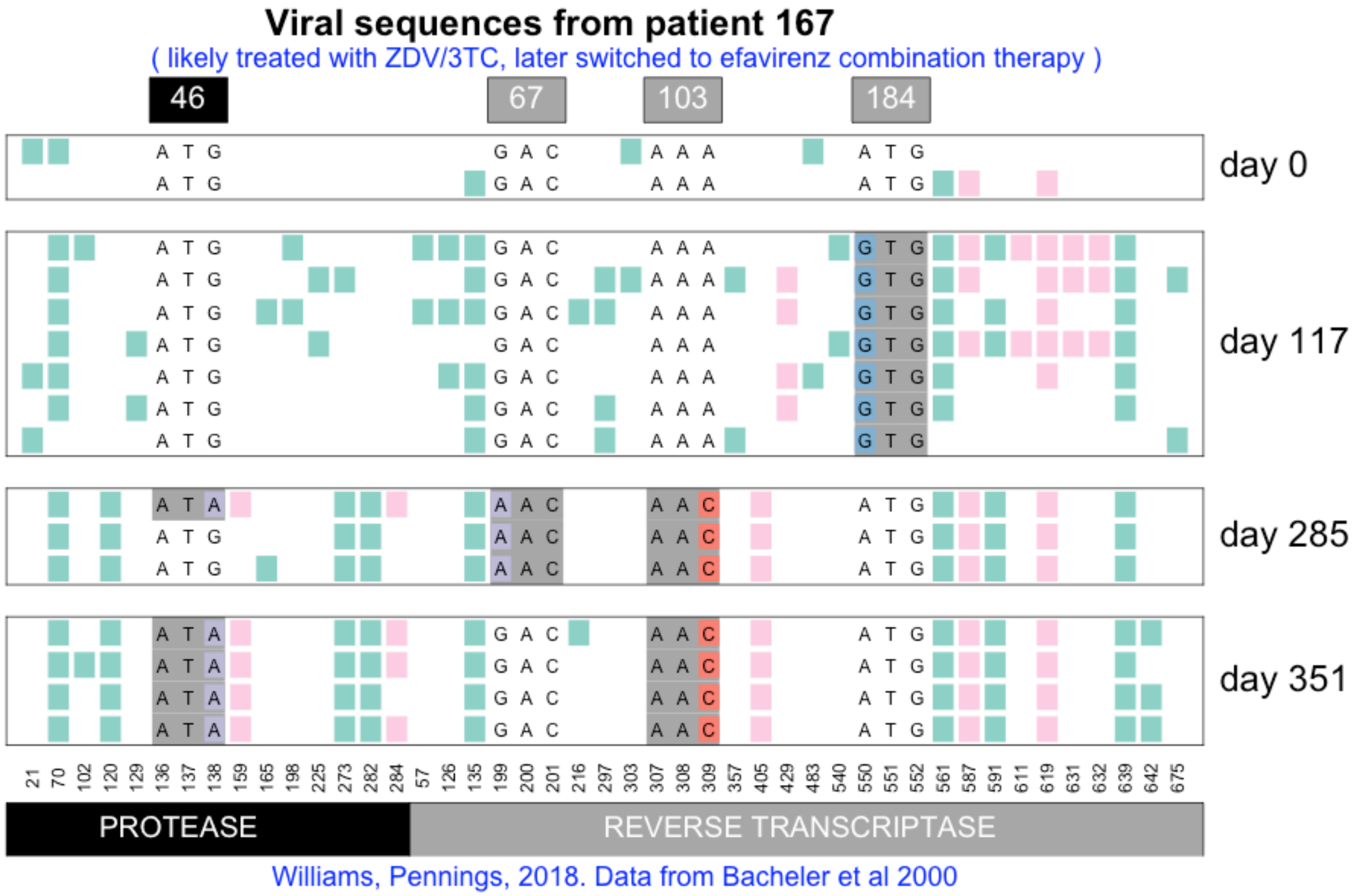

Viral sequences from patient 167
( likely treated with ZDV/3TC, later switched to efavirenz combination therapy )
46
67
103
184
day 0
day 117
day 285
day 351
21 70 102 120 129 136 137 138 159 165 198 225 273 282 284 57 126 135 199 200 201 216 297 303 307 308 309 357 405 429 483 540 550 551 552 561 587 591 611 619 631 632 639 642 675
PROTEASE
REVERSE TRANSCRIPTASE
Williams, Pennings, 2018. Data from Bacheler et al 2000

**Viral sequences from patient 168**

( likely treated with ZDV/3TC + efavirenz )

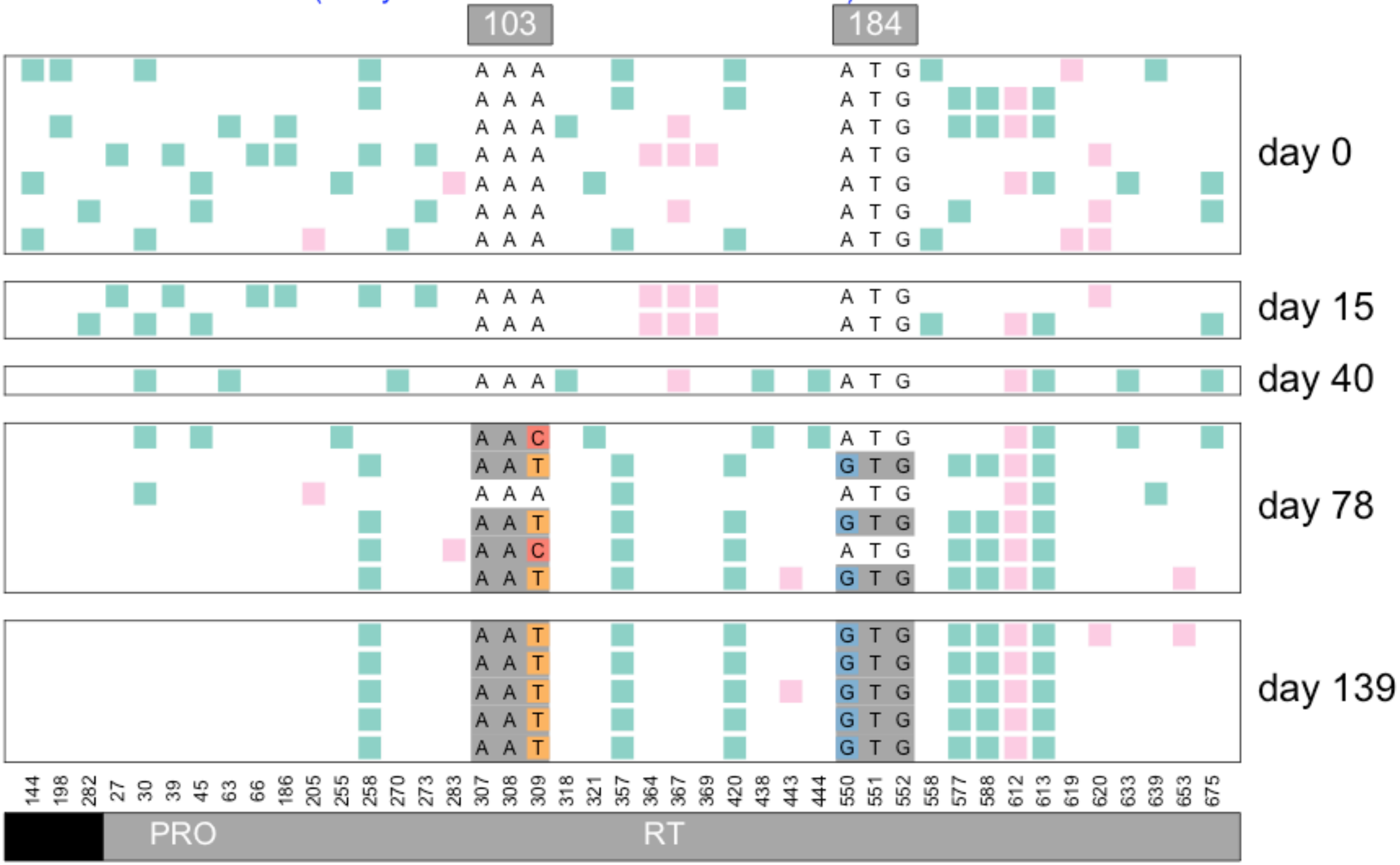


Williams, Pennings, 2018. Data from Bacheler et al 2000

#### Viral sequences from patient 169

( likely treated with ZDV/3TC )

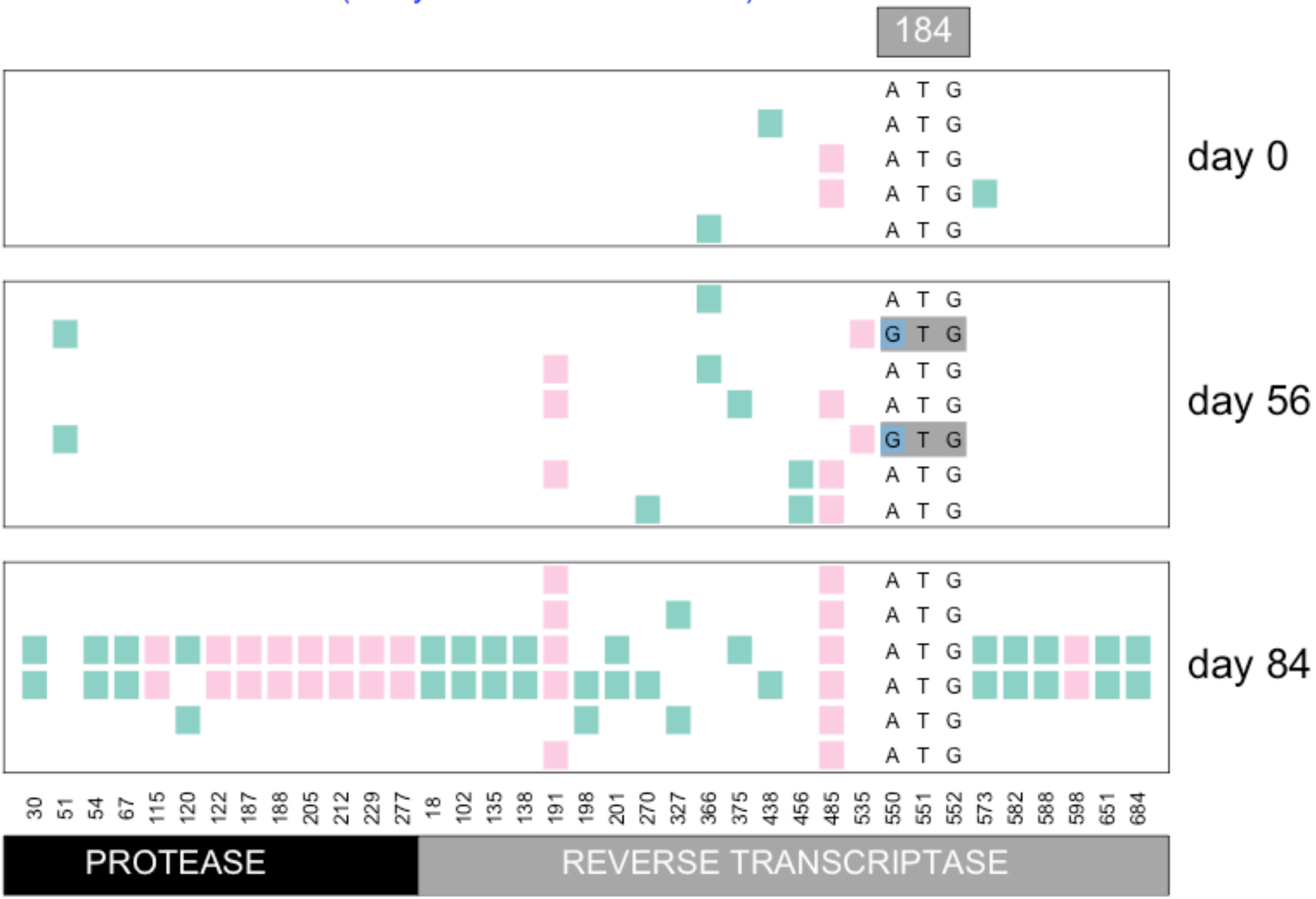


Williams, Pennings, 2018. Data from Bacheler et al 2000

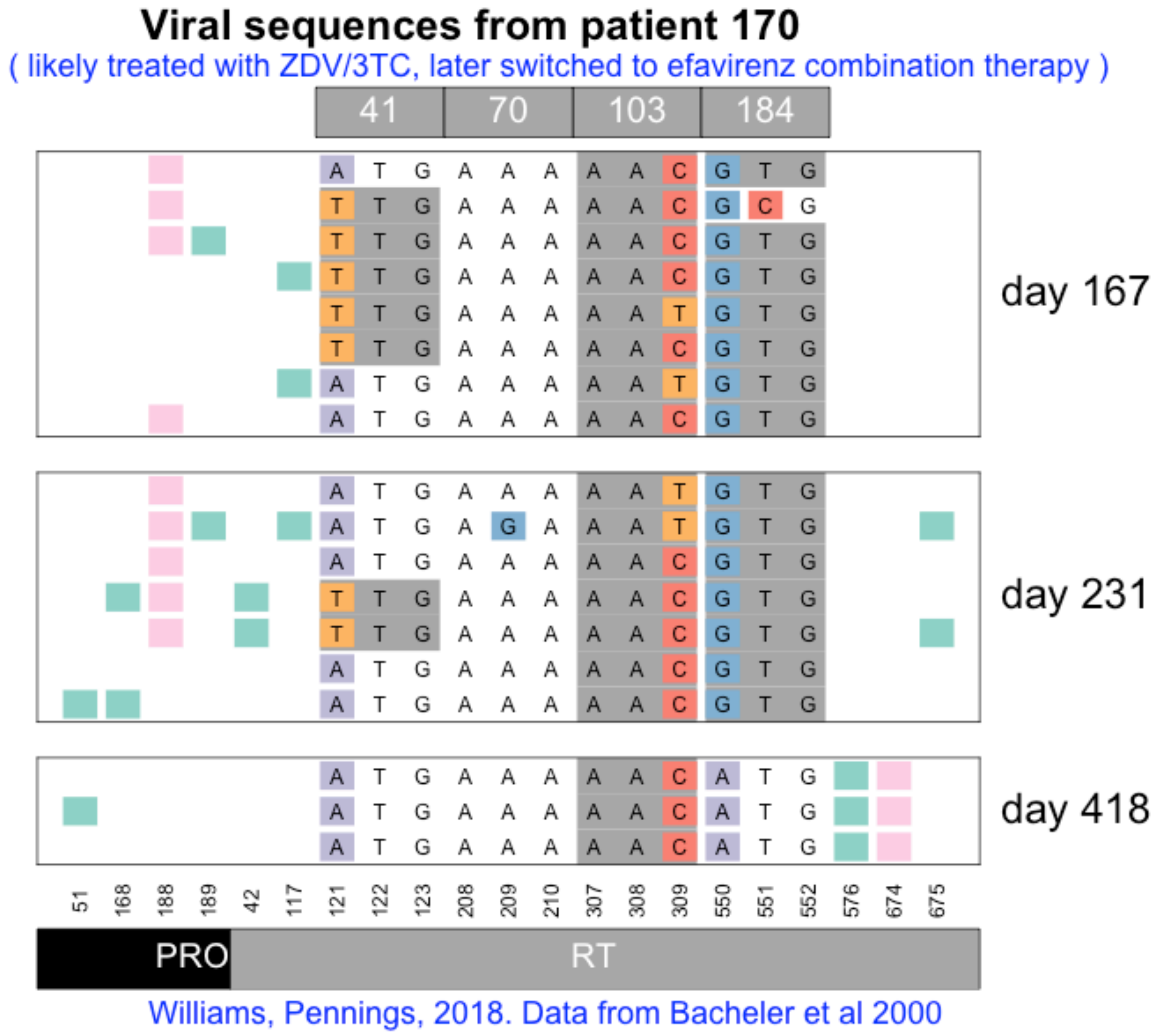
Viral sequences from patient 170
( likely treated with ZDV/3TC, later switched to efavirenz combination therapy )
41
70
103
184
day 167
day 231
day 418
51
168
188
189
42
117
121
122
123
208
209
210
307
308
309
550
551
552
576
674
675
PRO
RT
Williams, Pennings, 2018. Data from Bacheler et al 2000

**Viral sequences from patient 171**

( likely treated with ZDV/3TC + efavirenz )

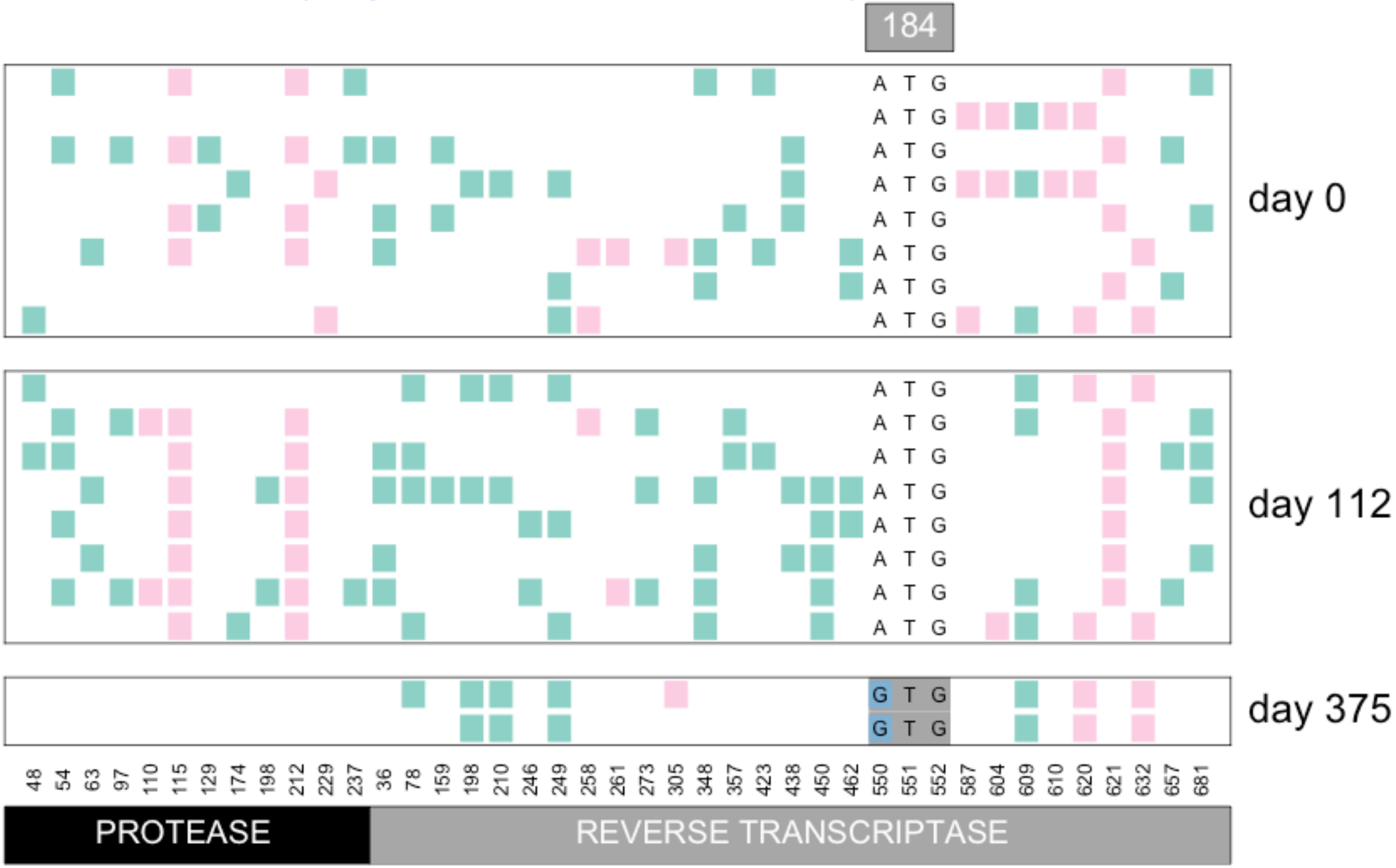


Williams, Pennings, 2018. Data from Bacheler et al 2000

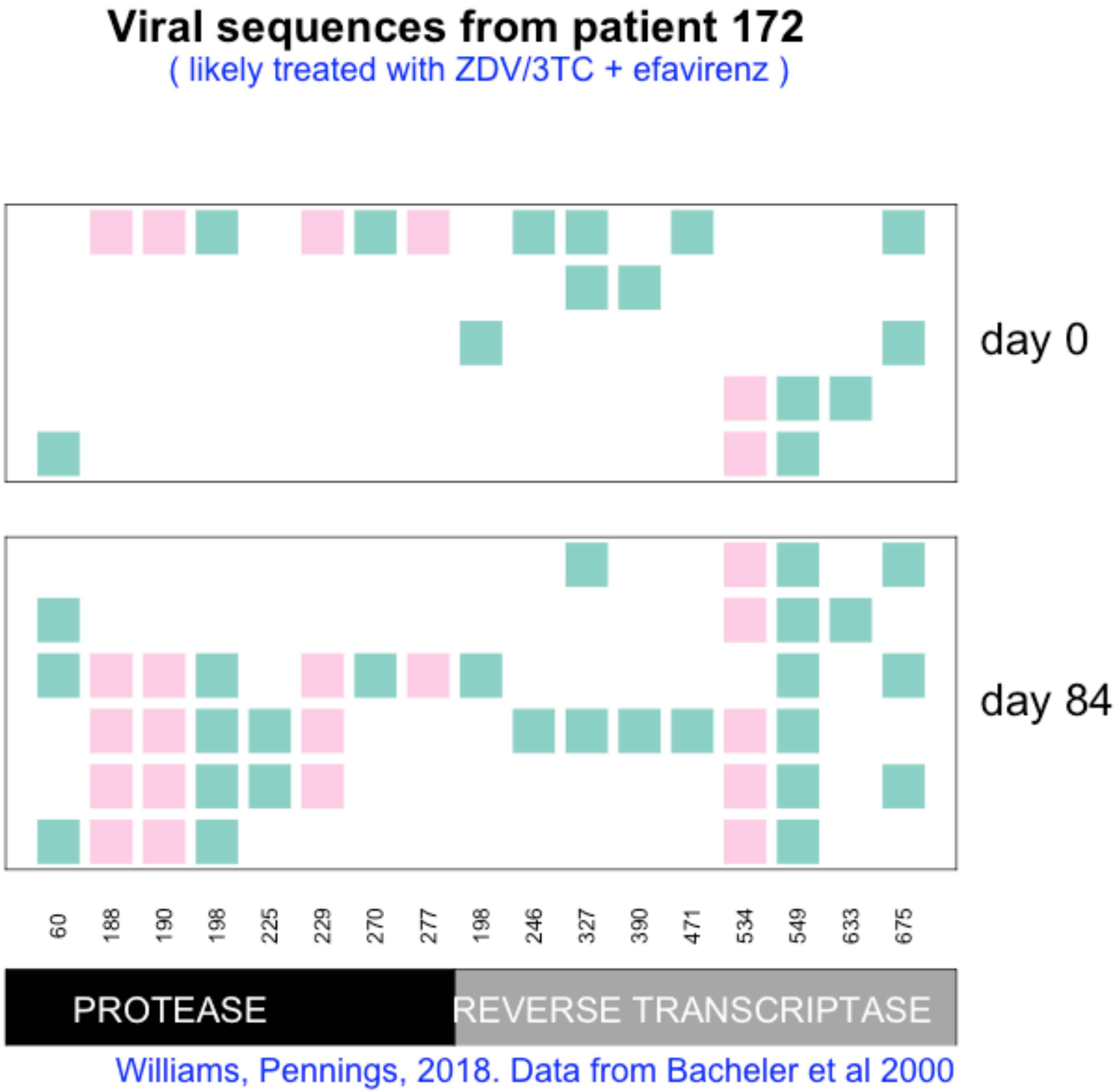
Viral sequences from patient 172
( likely treated with ZDV/3TC + efavirenz )
day 0
day 84
60
188
190
198
225
229
270
277
198
246
327
390
471
534
549
633
675
PROTEASE
REVERSE TRANSCRIPTASE
Williams, Pennings, 2018. Data from Bacheler et al 2000
